# Intratumor Heterogeneity and Evolution of Colorectal Cancer

**DOI:** 10.1101/2020.01.13.904276

**Authors:** Santasree Banerjee, Xianxiang Zhang, Shan Kuang, Jigang Wang, Lei Li, Guangyi Fan, Yonglun Luo, Shuai Sun, Peng Han, Qingyao Wu, Shujian Yang, Xiaobin Ji, Yong Li, Li Deng, Xiaofen Tian, Zhiwei Wang, Yue Zhang, Kui Wu, Shida Zhu, Lars Bolund, Huanming Yang, Xun Xu, Junnian Liu, Yun Lu, Xin Liu

**Author notes:** Correspondence Xin Liu, BGI-Qingdao, BGI-Shenzhen, Qingdao, 266555, China., Yun Lu, Department of Gastroenterology, General Surgery Center, The Affiliated Hospital of Qingdao University, Qingdao, 266555, China., Junnian Liu, BGI-Qingdao, BGI-Shenzhen, Qingdao, 266555, China. These authors contributed equally to this study (Santasree Banerjee, Xianxiang Zhang, Shan Kuang, Jigang Wang, Lei Li).

## Abstract

Intratumor heterogeneity (ITH) enable us to understand the evolution of cancer. ITH and evolution of colorectal cancer (CRC) has not been well studied. In this prospective study, we recruited different stages of 68 CRC patients with primary tumor at right-sided colon, left-sided colon and rectum. We performed high-depth whole exome sequencing of 206 multi-region tumor samples including primary tumors, lymph node metastasis (LN) and extranodal tumor deposits (ENTD). Our result showed extreme ITH with Darwinian pattern of CRC evolution, evolution pattern of left-sided CRC was more complex and divergent than right-sided CRC and both LN and ENTD were of polyclonal in origin. Extensive ITH was found in driver mutations in *KRAS* and *PIK3CA* genes, suggesting major limitations of single biopsies in clinical diagnosis for the CRC patients. In conclusion, our study showed the Darwinian pattern of CRC evolution with differences in evolution pattern between right-sided and left-sided CRC patients.

## Introduction

Identification of novel targets, and the development of target-based precision medicine for personalized cancer therapy is the biggest challenge in cancer research. Genomic instability in cancers is a continuous process involving genetic alterations at gene or chromosomal level. Extreme genetic heterogeneity drives a tumor from its benign state to malignancies. Tumor multi-region sequencing reveals ITH and evolution which play a key role in progression and metastases of the tumor, as well as identifying and developing novel targets for target-based precision medicine in personalized cancer therapy^1^. The development of effective target-based precision medicine and personalized cancer therapy is based on ITH and the pattern of clonal as well as subclonal evolution in CRC tumors^2^. Therefore, patients with CRC may respond variably to the same treatment, due to ITH, despite there being no significant differences identified in the tumor histopathology^3^. In addition, mono-sampling biopsies for clinical diagnosis are inadequate, due to ITH^4^. Hence, study of ITH is highly significant from both clinical and biological perspective, to understand the genomic changes driving the malignant process, which is fundamental to developing an effective personalized cancer therapy.

CRC is the third most common malignancy and the second leading cause of cancer death worldwide, with 18.1 million new cancer cases, and 9.6 million deaths in 2018^5^. According to the World Health Organization (WHO) GLOBOCAN database, there were 1,849,518 estimated new CRC cases and 880,792 CRC-related deaths in 2018^6^. In China, CRC is the second most common neoplasia, occupying the fifth position in mortality, accounting for an incidence of 521,490 new cases and 248,400 deaths in 2018^6^.

Amongst CRC patients, the stage of the disease is one of the most important prognostic factors which is correlated with the disease survival rate^7^. Tumor Node Metastasis (TNM)/American Joint Committee on Cancer (AJCC) Cancer Staging system is the gold standard for determining the correct cancer stage, helping us for making appropriate treatment plans. Among CRC patients, the presence of cancer cells in lymph nodes is define as stage III disease^8^. To date, the molecular signature and evolutionary relationship between LN and ENTD has not been clear. Hence, the characterization of the molecular signature and evolution of the primary tumor, LN and ENTD is very significant for TNM staging and therapeutic interventions for the patients with CRC.

The location of the primary tumor, either in the right- or left-side of colon, is also an important prognostic factor^9,10^. Clinical symptoms are also different between patients with right-sided and left-sided colon cancers. A possible explanation for this clinical heterogeneity might be due to the differences in their embryonic origin, genomic expression profiles and tumor microenvironment^11,12^. The differences in genomic expressions and subsequent alterations has not been studied well to explain the clinical heterogeneity between patients with right- and left-sided colon cancer.

Recently, tumor multi-region sequencing studies of primary tumor have demonstrated ITH^13-21^. This multiregional sequencing approach, sequencing DNA samples from geographically separated regions of a single tumor, explores ITH and cancer evolution^22-28^. Large-scale multiregional sequencing studies have systematically revealed ITH as well as cancer evolution in patients with non-small-cell lung cancer and renal cancer^22-24^. However, large-scale multiregional sequencing studies of CRC have not been well reported. In addition, multiregional sequencing studies in CRC were performed at relatively shallow sequencing depths, making it difficult to assess ITH, due to inability to detect somatic mutations with low frequencies^13-17^.

In order to overcome the drawbacks of previous studies, we have comprehensively studied the ITH and evolution of CRC, using high depth (median depth of 395×) whole exome sequencing of 206 multi-region tumor samples and 68 matched germline samples from 68 CRC tumors, determined the differences of ITH, and the evolution of CRC in patients with primary tumors in both right-sided and left-sided colon and characterized the molecular signature of the primary tumor, LN and ENTD, to define their evolutionary relationship.

## Results

Comprehensive clinical descriptions of these 68 patients were provided in Supplementary Table S1. Tumor multi-region high depth (median depth of 395×, range 179-596) whole exome sequencing (WES) was performed with 206 tumor regions (2-7 regions per tumor) including 176 primary tumor regions, 19 LN regions and 11 ENTD regions, as well as 68 matched germline samples from 68 CRC patients. WES identified 6 hypermutated (mutation rates of each tumor region were more than 10 mutations/1 Mb bases) CRC patients, of these four patients were identified with microsatellite instability (MSI). The remaining 62 CRC patients were microsatellite stable (MSS) and of these, 12 patients had right-sided colon tumors, 20 had left-sided colon tumors and 30 had rectal tumor. Hypermutated patients were analyzed separately.

Multiregion tumor tissue samples from 68 CRC patients were sequenced and analyzed (Supplementary Fig. S1). In our study, the experiments and data analysis workflow were shown in Supplementary Fig. S2.

### ITH in colorectal tumors

WES of 62 tumors with 188 tumor regions identified 19454 somatic mutations including 17560 SNVs (14361 non-silent SNVs) and 1894 INDELs (Supplementary Table S2). The mutation rate of multi-region whole-exome sequencing was significantly more than single sample sequencing due to detection of subclonal mutations (median number of mutations/1MB bases, 4.61 vs. 3.23; P=8.9×10^−9^) (Supplementary Fig. S3). In our study, the mutation rate of single sample sequencing was significantly higher than single CRC sample sequencing data from The Cancer Genome Atlas^29^ (TCGA), probably due to the higher sequencing depth in our study (median number of mutations/1 MB bases, 3.23 vs. 2.07; P=1.7×10^−22^) (Supplementary Fig. S3). Then, identified somatic mutations were divided into clonal (mutations present in all cancer cells with cancer cell fraction (CCF) > 0.9 across all the regions of a tumor) and subclonal (mutations present in only a subset of cancer cells) mutations (Fig. 1A). It is worth noting that 2 patients (CRC32 and CRC36) with left-sided colon tumors and 6 patients (CRC49, CRC42, CRC51, CRC48, CRC52 and CRC60) with rectal tumors had not identified with clonal mutations, suggesting that branched evolution was widespread in patients with left-sided colon tumors. In addition, patients with right-sided colon tumors had significantly more clonal mutations than the patients with rectal tumors (median number, 160 vs 119; P=0.035) (Supplementary Fig. S4). There were no significant differences found in the number and percentage of mutations between early (stage I and II) and late (stage III and IV) stage of patients (Supplementary Fig. S5).

**Figure 1.**
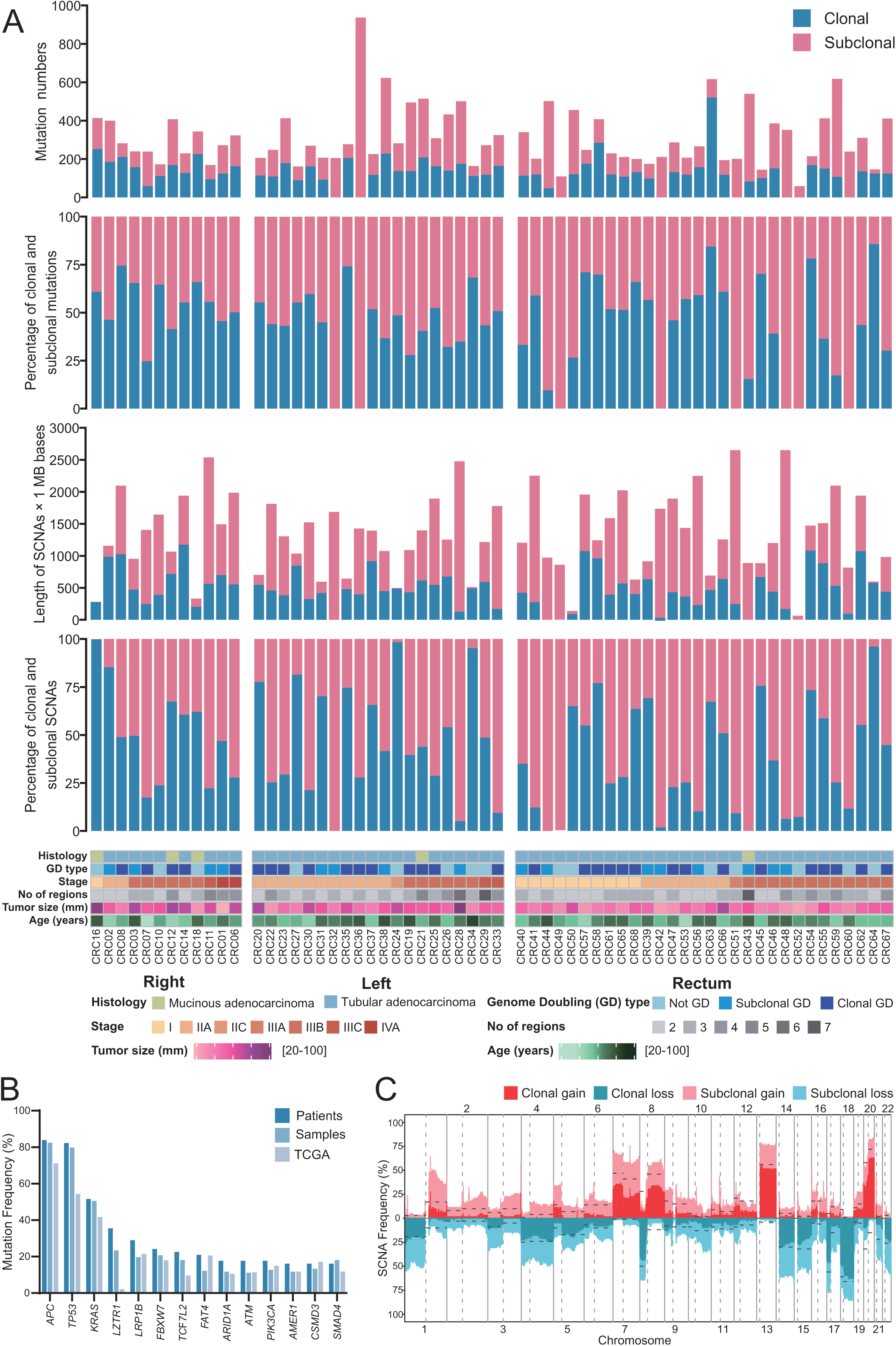
Overview of genomic heterogeneity in CRC tumors. (A) Heterogeneity of mutations and somatic copy-number alterations (SCNAs). Tumors were sorted by location and stage. (1) Number of all SNV and INDEL mutations (including coding and noncoding mutations) in CRC tumors. (2) The percentages of clonal mutations in CRC tumors. (3) Quantification of SCNAs in CRC tumors. (4) The percentages of clonal SCNAs in CRC tumors. (5) Demographic and clinical characteristics of the 62 CRC patients in this study (divided by histology; whole genome doubling status; stage; number of regions; tumor size; age and tumor location). (B) Mutation frequency of driver genes (driver mutations occurred in not less than 10 patients) and comparison with TCGA data. (C) Frequency of SCNAs in CRC tumors. The dotted lines were frequency of SCNAs in TCGA CRC samples.

Somatic copy number alterations (SCNAs) were measured as length of segments affected by either gains or losses (detailed copy number data has been given in Supplementary Table S3). We summarized the total length of the genome that subjected to SCNAs and calculated the percentage of clonal and subclonal SCNAs (Fig. 1A). Interestingly, in a patient (CRC43) with a rectal tumor, all SCNAs were subjected to subclonal SCNAs. There were no significant differences in the length and percentage of SCNAs among the patients with right-sided, left-sided and rectal tumors as well as between early and late stage of CRC tumors (Supplementary Figs. S6 and S7).

In our study, we identified that the mutation frequency of 14 driver genes (*APC, TP53, KRAS, LZTR1, LRP1B, FBXW7, TCF7L2, FAT4, ARID1A, ATM, PIK3CA, AMER1, CSMD3* and *SMAD4*) were higher at patient-level than at sample-level (Fig. 1B). In addition, we also found that the mutation frequency was higher at patient-level compared to the TCGA study^29^ except *CSMD3* (Fig. 1B). Notably, the mutation frequency of the *LZTR1* gene was much higher than TCGA study^29^ (Fig. 1B). We also identified that the frequency of SCNAs was higher than TCGA^29^ study data, probably due to the identification of subclonal SCNAs in our study (Fig. 1C).

### Clonal architecture in CRC

All the mutations (SNVs and INDELs) were clustered according to their CCF values to understand the clonal architecture and evolutionary history of 62 CRC tumors. Each colored circle in the phylogenetic tree represented one cluster of the tumor (Fig. 2). Phylogenetic trees for 62 tumors and 188 regions together with schematic diagram of 100 tumor cells representing distribution of clusters in each tumor region (Supplementary Fig. S8). Driver mutations, driver SCNAs and their clusters were annotated beside the phylogenetic trees (Supplementary Fig. S8). Detailed information of cluster numbers for each tumor was listed in Supplementary Table S4, with a median of 6 clusters per tumor (range, 1 to 13).

**Figure 2.**
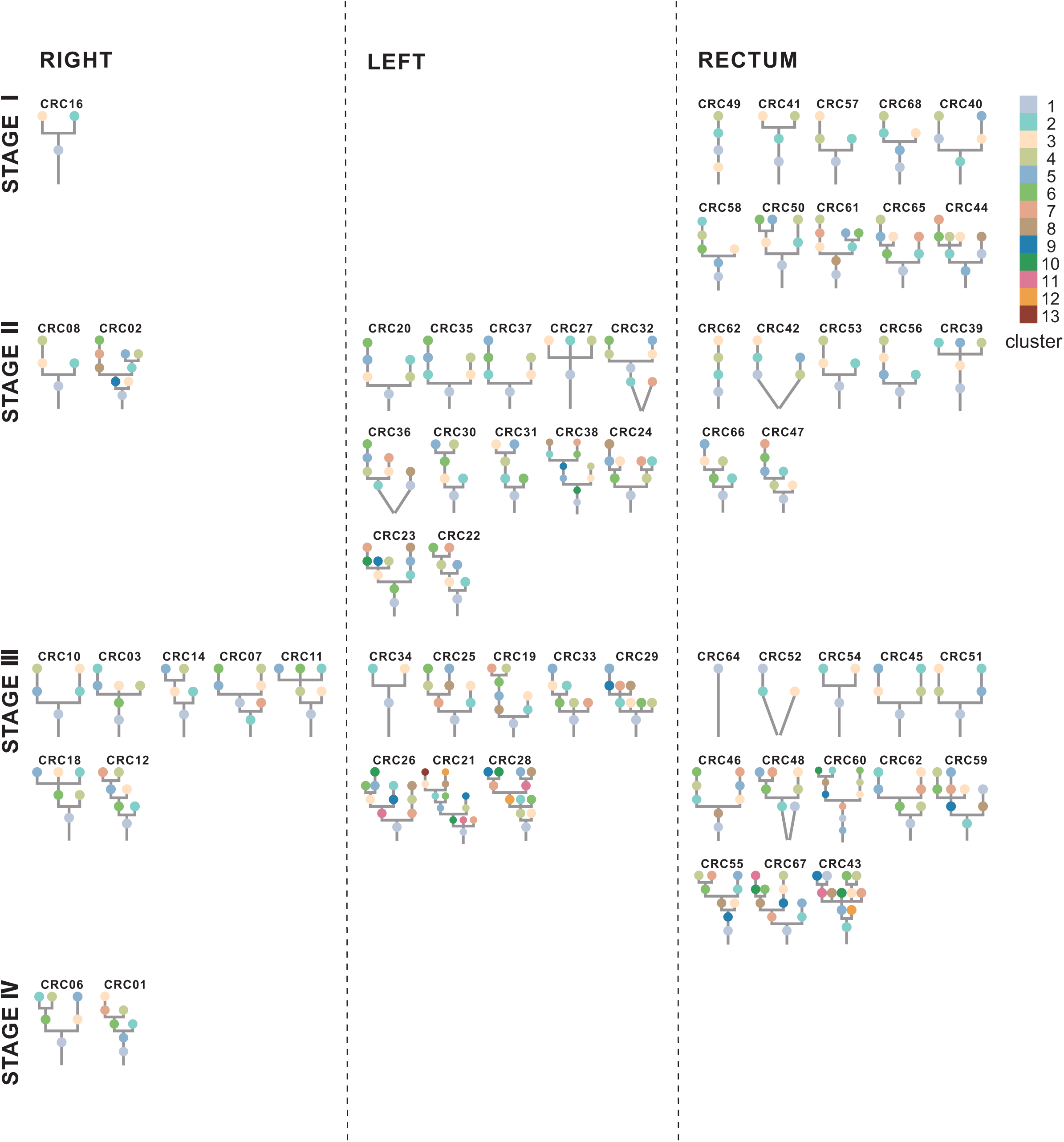
Phylogenetic trees. Phylogenetic trees for each CRC tumor were shown. The trees were ordered by overall stage (I, II, III, IV) and position (right-sided colon, left-sided colon and rectum). The cluster number corresponding to the color was displayed in the upper right corner with largest cluster labeled “1”. The lines connecting clusters does not contain any information.

Patients with left-sided colon tumors possessed significantly more cluster numbers than patients with both right-sided colon tumors (median number, 7.5 vs. 6; P=0.028) and rectal tumors (median number, 7.5 vs. 5.5; P=0.025) (Supplementary Fig. S9), which potentially reflected the more evolutional diversity in patients with left-sided colon tumors. There were no significant differences in cluster numbers between early and late stage of CRC tumors (Supplementary Fig. S9). Only 22 out of 188 tumor regions (12%) were presented with subclones in all the branches of the phylogenetic tree (Supplementary Fig. S8). This highlighted the limitations of single biopsy strategy of clinical diagnosis since mono-sampling was not enough to capture all the genetic information within a tumor.

### Driver event alterations in CRC evolution

Identifying cancer driver events and their clonality might provide important evidences for developing the target-based effective therapeutic strategies. Therefore, driver mutations, driver SCNAs, arm level SCNAs and their clonality were analyzed for CRC tumors (Fig. 3).

**Figure 3.**
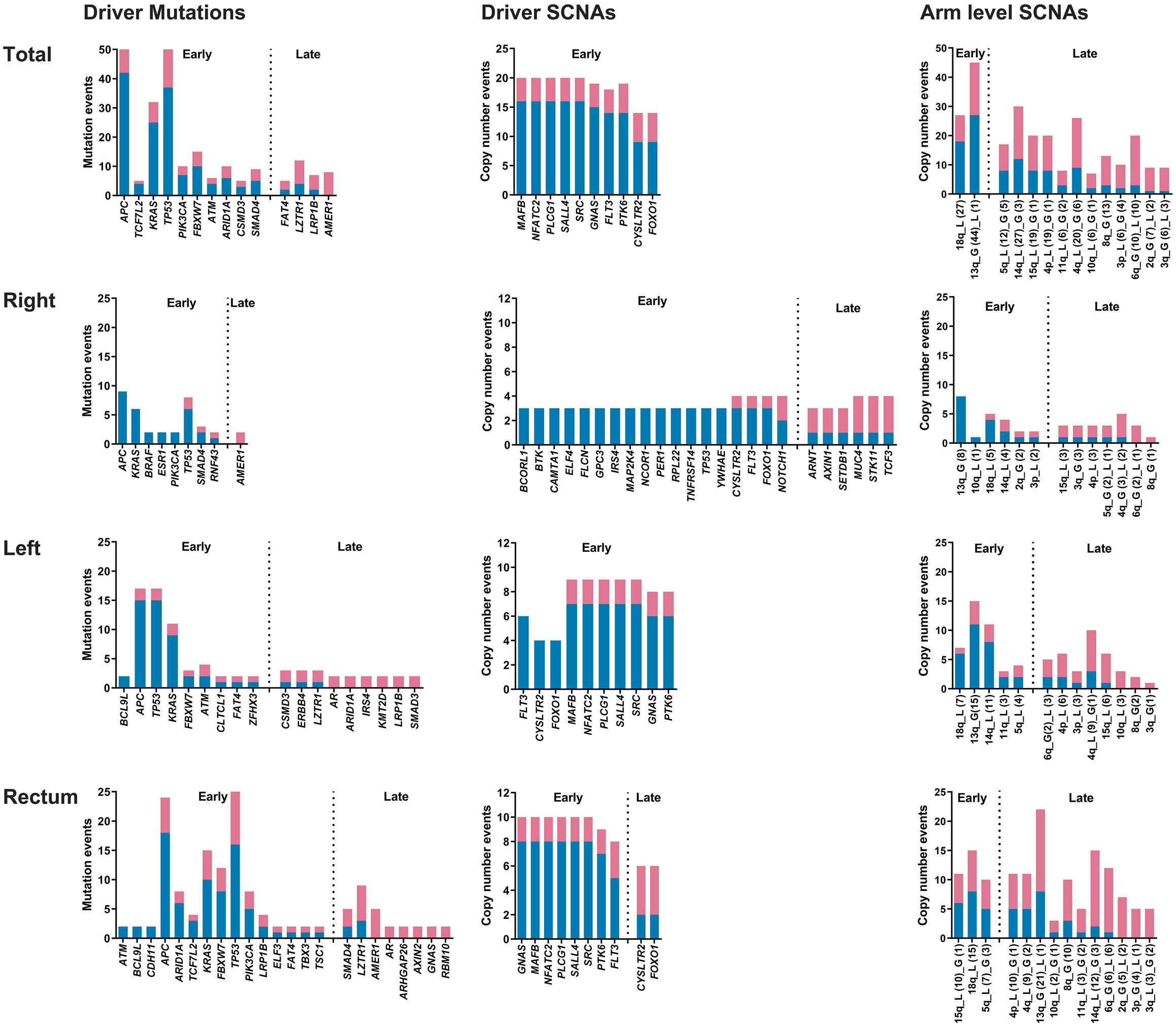
Summary of driver events in CRC evolution. Mutations and SCNAs were shown as occurrence in patients indicating whether the events are clonal (blue) or subclonal (red). Only genes that were mutated in at least five patients in total or two patients in right-sided colon/left-sided colon/rectum were shown. For SCNAs, driver SCNAs in at least 20% of the patients were shown while all the arm level SCNAs were shown. Driver events with more subclonal occurrence than clonal occurrence in tumors were late events, otherwise they were early events. In the arm level SCNAs part, “G” represented gain, “L” represented loss, and the numbers in parentheses represented the time of occurrence in tumors.

We identified 1373 driver events (405 driver mutations, 707 driver SCNAs and 261 arm level SCNAs) among 62 CRC tumors. Of these events, 44% of driver events (605 out of 1373) were found to be subclonal (41% of driver mutations, 40% of driver SCNAs and 60% of arm level SCNAs). However, significantly lower percentage of clonal driver events were identified in rectal tumors than patients with both right-sided (median percentage, 56% vs. 72%; P=0.031) and left-sided colon tumors (median percentage, 56% vs. 74%; P=0.047) (Supplementary Figs. S10 and S11), which potentially reflected the increased diversity in driver events existing amongst different tumor regions of patients with rectal tumors. Late stage CRC tumors possessed significantly more subclonal driver events than early stage CRC tumors (median number, 8 vs. 4; P=0.043) (Supplementary Figs. S12 and S13), which suggested that late stage CRC tumors were more advanced in evolution than early stage CRC tumors.

In addition, no driver events were consistently clonal among 62 CRC tumors, suggesting high ITH status and evolutionary diversity among CRC tumors, which might be the reason of low efficiency of target-based precision medicine in CRC treatment (Fig. 3). All the driver SCNAs and most of the driver mutations were early events while very few arm level SCNAs were early events, suggesting that genomic instability process occurred firstly at the driver SCNA level, then at the driver mutations level, and finally at the driver arm level SCNA level.

Driver mutations in *APC, TP53* and *KRAS* were majorly identified in CRC tumors with right-sided colon tumors, left-sided colon tumors and rectal tumors. Mutations in *APC, TP53* and *KRAS* genes were predominantly clonal and early among different locations of CRC tumors, suggesting their significance and key roles in tumor initiation. However, except for *APC, TP53* and *KRAS*, other identified driver mutations were completely different between patients with right-sided and left-sided colon tumors (Fig. 3). The genes of driver SCNAs identified were the same in patients with left-sided and rectal tumors while only 3 out of 24 genes of driver SCNAs (*CYSLTR2, FLT3* and *FOXO1*) were same in patients with right-sided and left-sided colon tumors (Fig. 3). The huge differences in both driver mutations and driver SCNAs between the patients with right-sided and left-sided colon tumors suggested that left-sided colon tumors were evolutionary closer to rectal tumors than to right-sided colon tumors. Chromosomal arm level gain of 13q and loss of 18q were mostly occurred and predominantly clonal and early in colon and rectal cancer patients except that gain of 13q were late events in rectal cancer.

### Convergent features and parallel evolution in CRC

Evidence of convergent mutations in tumor driver genes may shed light on evolutionary selection, which may provide therapeutic targets for treatment. *APC, TP53* and *KRAS* were the most frequently mutated driver genes identified in our study, 80.6 % (50 / 62), 80.6 % (50 / 62) and 51.6 % (32 / 62) respectively (Supplementary Fig. S14). Among these three genes, *APC* was the most often mutated gene in CRC patients. Among the 50 CRC tumors with *APC* mutations, 19 (38%) had 2 mutations, consistent with the two-hit hypothesis of *APC* genes in CRC tumorigenesis (Supplementary Fig. S15).

Evolutionary selection was also exemplified by parallel evolution of driver mutations, in which different driver mutations in distinct regions of the same tumor converge on the same gene. In CRC36 (left-sided colon tumor), two different nonsynonymous mutations in *TP53* were detected in tumor region 3 while another nonsynonymous mutation of *TP53* was detected in tumor region 1 and 4, indicating parallel evolution of *TP53*.

### Positive selection

In order to further evaluate evolutionary selection at the mutational level, we used the ratio of dN/dS, which could reflect the degree of enrichment of protein-altering mutations.

Evidence for positive selection (dN/dS>1 based on the 95% confidence intervals) was rejected when all the non-synonymous mutations were considered (Supplementary Table S5). However, when genes were narrowed to all driver genes identified in the COSMIC Cancer Gene Census (v88), positive selection was observed in clonal rather than subclonal of all non-synonymous mutations. These findings suggested that positive selection happened in cancer driver genes in early stage of CRC evolution.

### Mutation signature

We analyzed mutational processes for CRC evolution by using published mutational signatures^30^. We found that the age-related signature 1 was the predominant mutational process for CRC tumors, with a median percentage of age-related mutations of 70% (Supplementary Fig. S16). Interestingly, a patient (CRC66) was identified with all the mutations with age-related signature 1 while another patient (CRC63) was identified with defective DNA mismatch repair-related signature 6 and 15.

The median percentage of age-related signature 1 in CRC tumors for clonal mutations was 73%, while it dropped to 53% for subclonal mutations (Supplementary Fig. S16). This finding suggested that except for age, other mutational processes played more important roles in subclonal than clonal tumors, which accounted for ITH of CRC. Except for age, other main mutation processes were defective DNA mismatch repair-related signature 6, defective DNA double-strand break-repair-related signature 3 and defective DNA mismatch repair-related signature 15, suggesting that the main mutational process for ITH of CRC were age and defective of DNA repair system.

### Chromosome instability

Previously we analyzed the length and clonality of SCNAs relatively to ploidy (Fig. 1A), we then measured the absolute SCNAs in CRC tumors. SCNAs and ITH of SCNAs were ubiquitous, which described the continuing process of chromosome instability in CRC tumors (Supplementary Figs. S17 and S18). Left-sided colon tumor were found to have more loss type of SCNAs (total copy number = 0 or 1, or copy neutral loss of heterozygous) than rectal cancer (P=0.007) and have more SCNAs with total copy number equal to 1 than right-sided colon tumors (P=0.044) (Supplementary Fig. S17). Late stage tumors were identified with significantly fewer SCNAs than early stage CRC tumors (P=0.016) (Supplementary Fig. S18). Specifically, the late stage CRC tumors had significantly fewer SCNAs with total copy number equal to 4 than early stage CRC tumors (P=0.043) (Supplementary Fig. S18). Moreover, late stage CRC tumors had significantly more subclonal driver SCNAs than early stage CRC tumors, which suggested that the loss of random SCNAs as well as enrichment of functional SCNAs in late stage CRC tumors (Supplementary Fig. S13).

The SCNA frequency pattern in patients with left-sided colon tumors and rectal tumors were similar to each other, while right-sided colon tumors were very different (Supplementary Fig. S19). Patients with right-sided colon tumors had more 9p gain, 3q gain, 19p loss and less 20q gain, 18p loss, 8p loss than all CRC tumors (Supplementary Fig. S19). Late stage CRC tumors had more 13q gain, 9p gain, 21p gain, 11q loss, 21q loss and 12p loss than early stage CRC tumors (Supplementary Fig. S20). Interestingly, both poor prognosis location (right-sided colon tumors) and stage (late stage of CRC tumors) of CRC tumors had more 9p gain.

### Genome doubling

If the percentage of autosomal tumor genomes with a major copy number of two or more in a tumor were 50% or more than 50%, then this tumor was classified as genome doubling tumor^31^. Genome doubling events were identified in 76% of tumors (found in 47 out of 62 tumors, 9 right-sided colon tumors, 15 left-sided colon tumors, and 23 rectal tumors) and appeared to be clonal in 66% of tumors (found in 31 out of 47 tumors, 5 right-sided colon tumors, 11 left-sided colon tumors and 15 rectal tumors), which suggested that whole genome doubling was an early event in CRC evolution. In our study, we identified that the rate of whole genome doubling was much higher than 36% found in a previous study of CRC^31^, and the high rate was likely to come from multi-region sequencing because 16 out of 47 CRC were identified with subclonal whole genome doubling.

We observed a strong positive correlation between genome doubling with the ITH of both mutations and SCNAs (Supplementary Figs. S21 and S22). These findings suggested that the genome doubling was important for the progression of chromosomal instability. Tumors without genome doubling had a significantly higher percentage of clonal SCNAs than subclonal genome doubling tumors (median percentage, 62% vs. 29%, P=0.037). Moreover, tumors with clonal genome doubling had significantly more clonal SCNAs than both subclonal genome doubling tumors (median length × 1 MB, 545 vs. 420, P=0.017) and tumors without genome doubling (median length × 1 MB, 545 vs. 391, P=0.022) (Supplementary Fig. S22).

### Mirrored subclonal allelic imbalance

Recent studies identified parallel evolution of SCNA in NSCLC and renal cancer through mirrored subclonal allelic imbalance (MSAI)^22,24^, which was defined as the maternal allele was gained or lost in a subclone in one region, yet the paternal allele was gained or lost in a different subclone in another region. We identified MSAI events in 23 of 62 CRC tumors (37%, found in 5 right-sided colon tumors, 6 left-sided colon tumors and 12 rectal tumors) (Supplementary Fig. S23). MSAI parallel gain or loss events found in this study were summarized (Fig. 4A). Chromosomal regions of 7p and 13q were identified with parallel gain events in 3 tumors and chromosomal regions of 21q were identified with parallel loss events in 4 tumors. We also analyzed parallel evolution of driver SCNAs, 5 tumors (4 tumors of parallel amplification and 1 tumor of parallel deletion) were found to have driver SCNAs which overlapped with MSAI events (Figs. 4B and C). Interestingly, 2 of 5 patients (CRC12 and CRC59) were identified with parallel amplification of *FLT3* gene in chromosome 13 (Fig. 4C).

**Figure 4.**
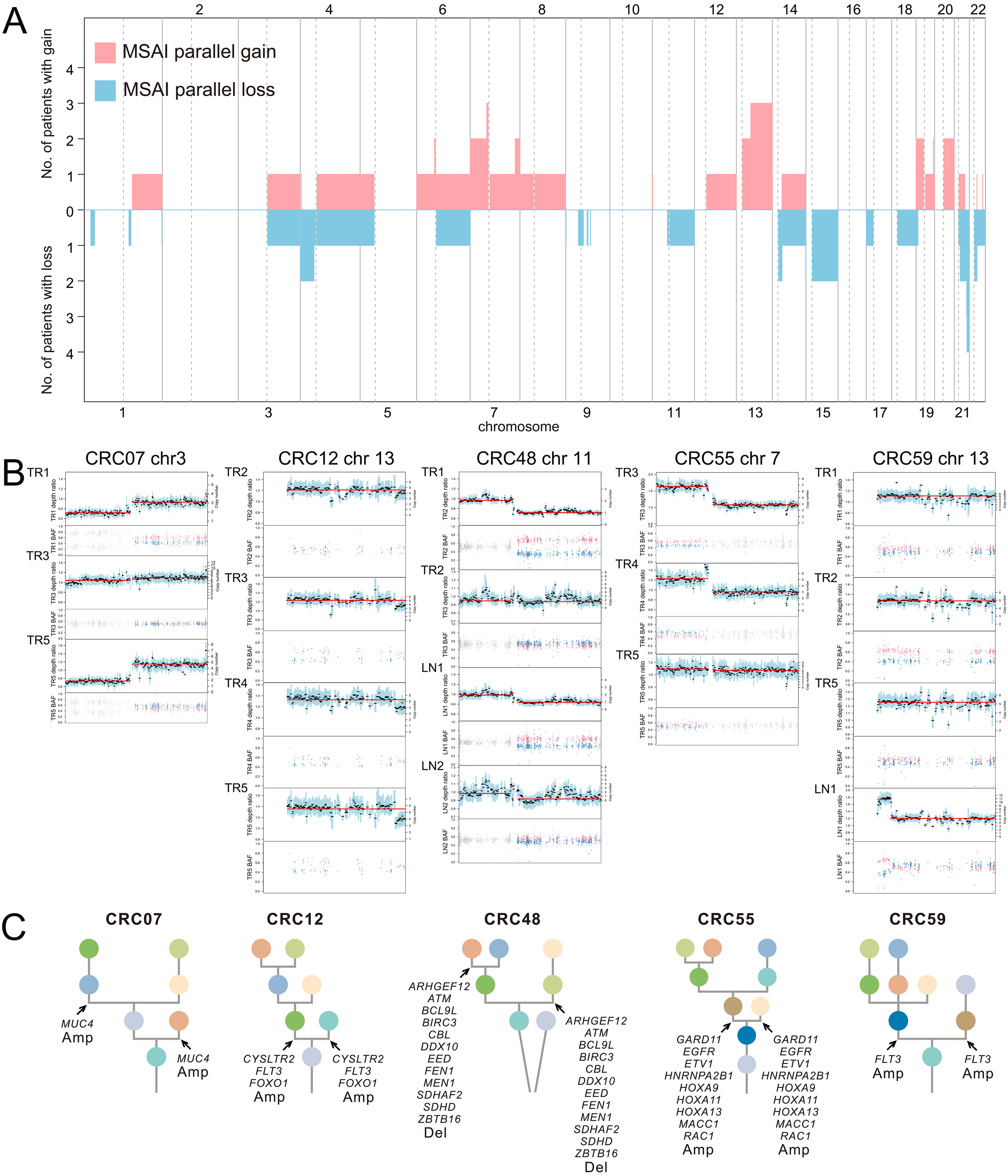
Parallel evolution. (A) Genomic position and size of all mirrored subclonal allelic imbalance (MSAI) parallel gain or loss events found in this study. This included genome-wide copy number gains and losses which was subjected to MSAI events and their occurrence in CRC tumors. (B) Parallel evolution of driver SCNAs observed in 5 CRC tumors, indicted by the depth ratio and B-allele frequency values of the same chromosome on which the driver SCNAs (C) were located. (C) Phylogenetic trees that indicated parallel evolution of driver amplifications (Amp) or deletions (Del) (Driver SCNAs) detected through the observation of MSAI (arrows).

### Conserved evolutionary features in CRC

In order to understand the constraints and features of CRC evolution, we analyzed conserved patterns of driver events by REVOLVER^32^ (Fig. 5). Evolutionary trajectories were clustered by the CCF and cluster information of all the driver events in 62 CRC tumors and four clusters (cluster red, blue, green and purple) were found (Fig. 5). In order to understand whether conserved patterns of CRC evolution correlated to distinct clinical phenotypes, clinical and genomic metrics were shown under 4 clusters (Fig. 5).

**Figure 5.**
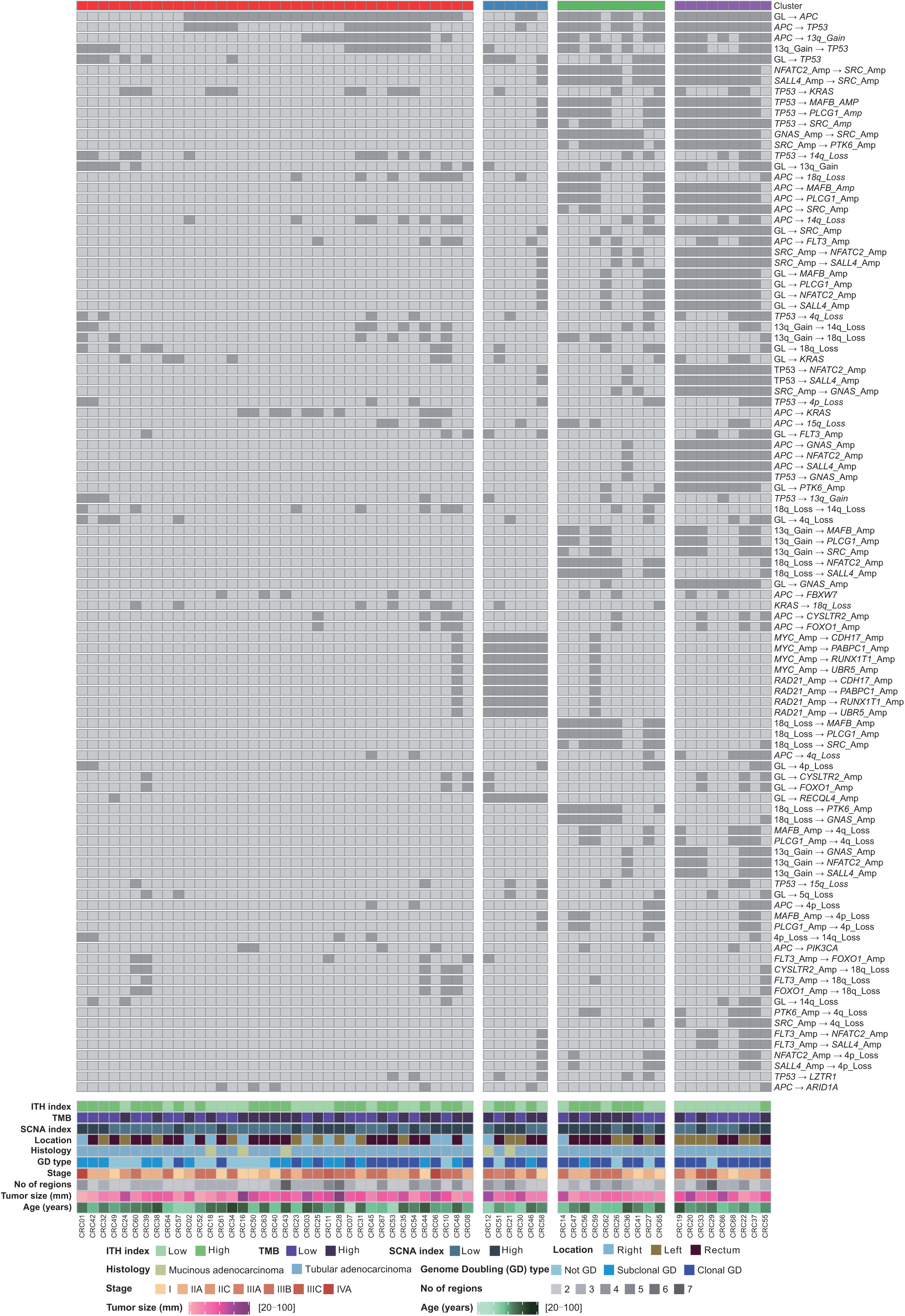
Evolutionary subtypes. Evolutionary trajectories were clustered based on CCF value and cluster information of driver mutations, driver SCNAs and arm-level SCNAs. Heat maps showed the most recurrent evolution for the most recurrent driver mutations, driver SCNAs and arm-level SCNAs. Alterations were ordered by their frequencies in CRC tumors. CRC tumors are annotated by the following parameters: ITH index (high: half of the largest ITH index value; low: the other half), TMB (high> median, low≤ median), SCNA index (high> median, low≤ median), tumor location, histology, whole genome doubling status, stage, number of regions, tumor size and age.

We found that the red and blue clusters had relatively fewer driver events than green and purple clusters. There were no specific genomic or clinical features for the tumors in red cluster. The blue, green and purple clusters had similar clinical and genomic features, which were enriched in left-sided CRC tumors and tumors with genome doubling type.

### Phylogenetic distance between LN and ENTD

We analyzed 16 non-hypermutated stage III patients to understand the phylogenetic distance and evolutionary relationship amongst primary tumor, LN and ENTD. CRC21, CRC28, CRC43 and CRC48 were identified with both LN and ENTD samples which were sequenced (Fig. 6). In CRC21, we identified that the clonal evolution of LN and ENTD was similar, while ENTD appeared evolutionarily later than LN (Supplementary Fig. S8). In CRC28, two ENTD were clustered together while LN were far away from them, which indicated that the LN and ENTD were polyclonal in origin (Fig. 6). In CRC 43 and CRC48, we identified that the ENTD were not clustered together with LN and evolved separately (Figs. 6 and S8). In tumors with more than one LN sequenced (CRC01, CRC11, CRC29 and CRC33), some LN were clustered together while some LN were not (Fig. 6). In tumors with two ENTD sequenced (CRC60), the two ENTD were far away from each other in the phylogenetic tree (Fig. 6). These findings suggested that both LN and ENTD were polyclonal in origin.

**Figure 6.**
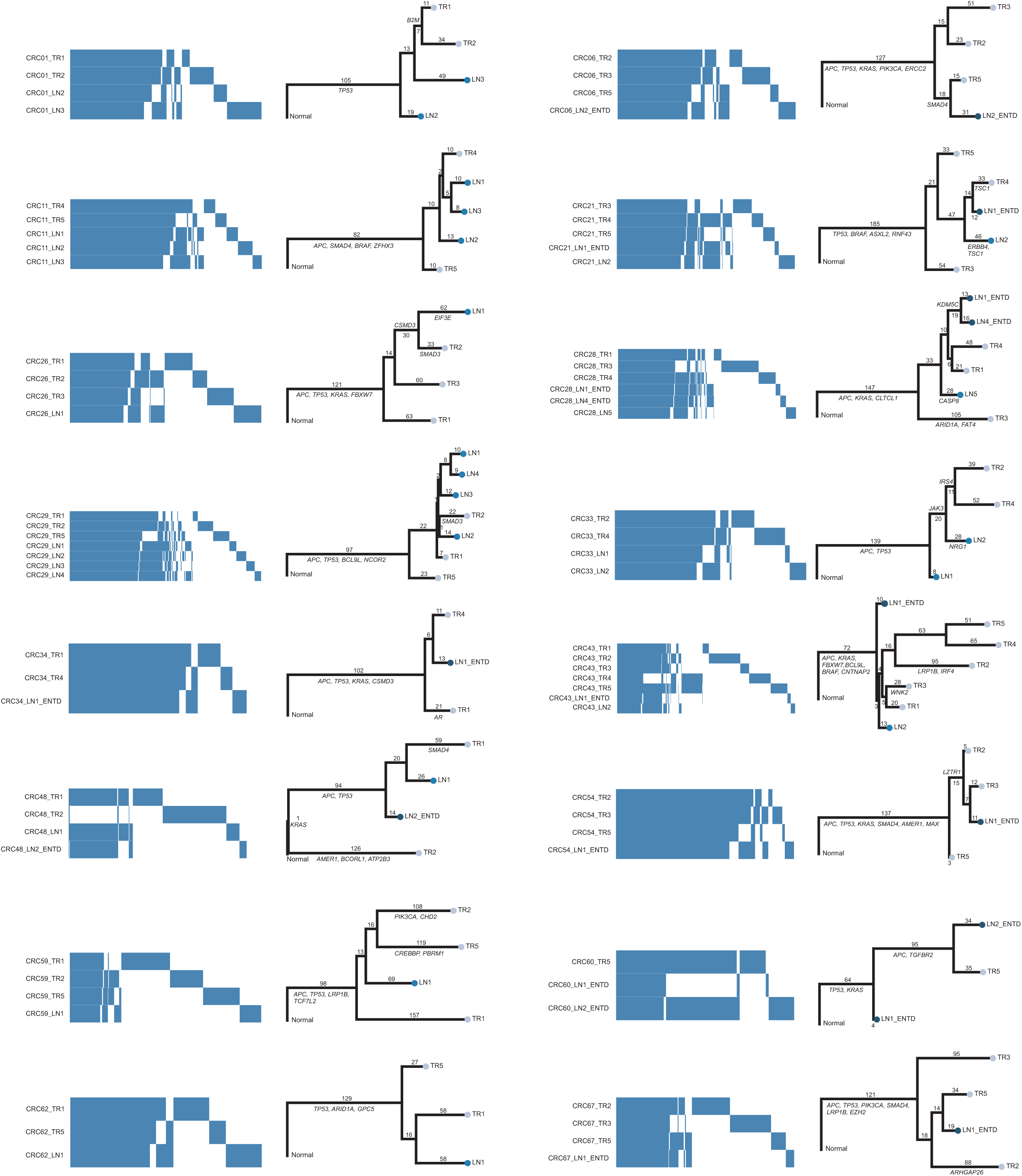
Phylogenetic distance between primary tumor, lymph node metastasis and ENTD. Heatmap showed the presence (blue) and absence (white) of all the mutations (SNVs and INDELs) among different tumor regions of the patients with lymph node metastasis or ENTD. Phylogeny reconstruction using maximum parsimony based on mutational presence or absence of all the mutations were shown beside heatmap. Driver genes were labeled in the phylogenetic trees.

LN were identified with significantly less TMB than primary tumors (P=0.035) (Supplementary Fig. S24). LN were also presented with significantly more loss type of SCNAs (P=0.043) than primary tumors (Supplementary Fig. S25). Regarding SCNA frequency, the biggest difference of gain events existed between LN and ENTD while the biggest difference of loss events existed between primary tumors and ENTD (Supplementary Fig. S26). In conclusion, LN and ENTD were different in both mutation and SCNA level.

### Evolution landscape of hypermutated CRC tumors

All 6 (CRC04, CRC05, CRC09, CRC13, CRC15 and CRC17) hypermutated CRC patients were identified with right-sided colon tumors, of these two (CRC09 and CRC13) were with MSS and remaining four (CRC04, CRC05, CRC15, CRC17) were with MSI tumors (Supplementary Fig. S27A). All of the 6 hypermutated tumors had mutations in mismatch-repair genes, *POLE* or *POLD* gene family (Supplementary Fig. S27A). CRC09 had one missense mutation and one nonsense mutation of *POLE*. CRC13 had one missense mutation of *POLE* (Supplementary Fig. S27A). These findings were consistent with that the predominant mutational process in these two MSS tumors was *POLE*-related signature 10 (Supplementary Fig. S27B). Defective DNA mismatch repair-related signature 6, 15, or 26 contributed to the mutational process of 4 MSI tumors (Supplementary Fig. S27B).

We next analyzed evolution landscape of hypermutated tumors in SCNA level. None of the tumor regions in 6 hypermutated tumors had genome doubling. Absolute SCNAs of hypermutated CRC tumors were less compared with non-hypermutated CRC tumors (Supplementary Figs. S27C and S17), which suggested that hypermutated tumors were mainly mutation driven tumors. Interestingly, CRC04 had MSAI events in X-chromosome (Supplementary Fig. S28).

## Discussion

In this study, we performed high-depth whole exome sequencing and analyzed 206 multi-region tumor samples from 68 patients with CRC. Our result showed very clear evidence of ITH in respect of both mutations and somatic copy number alterations. Our result showed the specific temporal and spatial features of evolution of CRC, following a Darwinian pattern of evolution. In addition, left-sided CRC was structurally and functionally more complex and divergent in terms of evolutionary perspective. We also identified that both ENTD and LN were polyclonal in origin and ENTD was a distinctive entity from LN, which appeared later in tumor evolution.

### Evolution pattern: Darwinian pattern of evolution and neutral evolution

In this present study, we found predominantly Darwinian pattern of evolution (59 out of 62 tumors) and a small portion of linear evolution (3 out of 62 tumors). Previous studies proposed neutral evolution model for colorectal cancers^13,33,34^, whilst our conclusion was different from them, based on two reasons. Firstly, clonal events of both mutations (SNVs and INDELs) and SCNAs were widespread, with a median percentage of 47% and 43% respectively. Secondly, our study demonstrated a clear selection for functional mutations rather than non-functional mutations in colorectal cancer. In addition, 59% of driver mutations were clonal while only 41% of non-driver mutations were clonal, which indicated the enrichment of clonal driver mutations in course of evolution. Furthermore, the dN/dS value was 1.03 (95% confidence interval, 0.983 to 1.07) for all non-synonymous mutations while it reached to 1.37 (95% confidence interval, 1.16 to 1.61) when we consider only cancer driver genes based on COSMIC database. These findings indicated that positive selection existed only for cancer related gene mutations. Thirdly, convergent and parallel events were also present for driver genes in both mutational and SCNA level, especially for genes *APC, TP53* and *KRAS*. Some studies showed that the evolution pattern for colorectal cancer was Darwinian evolution followed by neutral evolution^17,35^. In our study, according to the existence of mutations in different tumor regions, 28% of subclonal mutations were shared by tumor regions (branch or trunk mutations), which suggested the importance of branches in phylogenetic trees.

### Right-sided colon, left-sided colon and rectal cancer: In the light of genomic evolution

Previous studies^10,36^ had shown remarkable differences between right-sided, left-sided colon cancer and rectal cancer, based on histology, MSI status, genetic subtype and prognosis. Almost no research has been done to date for understanding the differences between different locations of CRC cancer from an evolutionary perspective. Our study demonstrated that ITH and evolution in different location of CRC were different in the following aspects: mutation, SCNA, structure of polygenetic tree and driver events. Firstly, rectal cancers had shown fewer clonal mutations than right-sided colon cancers, indicating higher ITH in rectal cancer at mutational level. Secondly, the pattern and clonality of SCNA frequency in right-sided colon cancer were different from left-sided colon cancer, which addressed the evolutionary difference between them at SCNA level. Thirdly, the structure of phylogenetic trees in left-sided colon cancer were more complicated and branched than that of the right-sided colon cancer. Specifically, left-sided colon cancer had the most complicated structure of the phylogenetic tree, reflected by the more cluster numbers. In addition, only left-sided colon cancer had polyclonal origin. Fourthly, left-sided colon cancer was enriched in clusters (blue and purple clusters) which had more driver events. These findings indicated that left-sided colon cancer had more functional diversity in the course of evolution. Specifically, rectal cancer had less percentage of clonal driver events than colon cancer, indicating more functional diversity occurred in the process of evolution in rectal cancer. In conclusion, our data showed that left-sided colon cancer were more divergent and complicated in terms of evolution than right-sided colon cancer, not only structurally but also functionally, which indicated that the evolutionary diversity might play an important role in the initiation and progression of left-sided colon cancer. Moreover, the frequency and clonality of SCNA was a potential and significant biomarker to distinguish right-sided colon cancer from the left-sided colon cancer.

### Primary tumor, LN and ENTD: In evolutionary perspective

To date, no systematic research studies have been done to understand the similarities and differences between ENTD and LN. In this study, we found that ENTD were later events in the evolution of the tumor than LN, which could be distinguished at mutational, SCNA and evolutionary levels. At the mutational level, the TMB of LN was significantly less than primary tumors. Also, LN and ENTD could not be clustered together in the polygenetic tree according to the occurrence of mutations. Unlike in previous studies^14,37,38^, different LN or ENTD in the same tumor did not cluster together in all cases, indicating the polyclonal origin of both LN and ENTD. We also identified here that LN were different from ENTD at SCNA level. In addition, the biggest difference between LN and ENTD was gain of events in SCNA frequency. ENTD thus appeared to be as later event than LN according to the clonal evolution history in CRC21. In conclusion, our present study provided comprehensive evidences to prove that ENTD and LN were two distinctive entities, which support the 7th and 8th editions of TMN staging.

### Stage and ITH in CRC

There was no significant difference between early and late stage of CRC at both mutational and evolution (cluster number) level. It was well known that chromosomal instability was associated with a worse prognosis. Therefore, we analyzed the difference between early and late stage of CRC at SCNA level. There was no significant difference in length and clonality of SCNA relatively to ploidy between early and late stage of CRC in our study. However, early stage CRC were found to have significantly more absolute SCNAs than late stage CRC in our study. These two distinctive conclusions suggested the definition of chromosomal instability was important. Interestingly, late stage CRC had significantly more subclonal driver SCNAs than early stage CRC, which suggested ITH of functional SCNAs rather than total absolute SCNAs might be more closely related with CRC progress.

### Germline mutations and ITH in CRC

Five patients in our study were found to have pathogenic or likely pathogenic germline mutations in DNA mismatch repair genes. Among them, two (CRC05 and CRC15) patients were hypermutated MSI patients identified with pathogenic and likely pathogenic mutations in *MLH1*. In addition, CRC33 were found to carry a likely pathogenic mutation in the *MLH1* gene, CRC37 were presented with a pathogenic mutation in *PMS2* gene and CRC44 were identified with a pathogenic mutation in *MLH3*. Interestingly, we found these 3 (CRC33, CRC37 and CRC44) non-hypermutated MSS patients had relatively divergent and complex clonal evolution with cluster number of 7, 6 and 8 respectively. These findings reminded us that germline mutations in MSI genes might accelerate the evolution process in CRC.

### Personalized therapy: Target-based Precision medicine for CRC

Presently, the principle of treatment for CRC has shifted from ‘one gene, one drug approach’ paradigm to a ‘multi-gene, multi-drug’ model when making decisions for precision medicine^39,40^. It is of great clinical importance to define the mutation status, clonality of genes and biomarkers in CRC. Among 62 CRC tumors, 32 were identified with driver mutations in *KRAS* while 7 were subclonal. Similarly, 7 CRC tumors harbored clonal driver mutations in *PIK3CA*, while 3 were carrying subclonal driver mutations. Driver mutations in *NRAS* (1 tumor), *BRAF* (4 tumors) and *ERBB2* (1 tumor) were all clonal in origin. In addition, among 4 driver mutations in *BRAF*, 2 were V600E. Furthermore, we identified 5 CRC tumors with *EGFR* amplification whilst only 1 was clonal in origin. No amplification of *ERBB2* was found in our study. However, the occurrence of subclonal driver mutations in biomarker genes emphasized the limitations of the single biopsy strategy for the clinical diagnosis of CRC. For example, in CRC59, we found all 4 tumor regions had wildtype *KRAS, BRAF* and *NRAS* genes, whilst only 1 region had the driver mutation in *PIK3CA*.

## Materials and methods

### Patient recruitment

The study was approved by the Ethics committee of the Affiliated Hospital of Qingdao University. All the samples were collected after obtaining written informed consent from the patients. Patients were recruited based on the following criteria. (i). age over 18 years, (ii) patients clinically diagnosed with CRC by enteroscopy, imaging, biopsy and followed by surgery, and histopathology performed with the resected tumor tissues. Patients with sufficient tissue were available for the study.

### Sample collection

A pathologist performed macroscopic examination of all surgically resected specimens to guide the multi-region sampling in this study. Firstly, the pathologist performed routine pathological sampling for clinical diagnosis, and then multi-region sampling was performed by using the remaining samples. At least 2 regions of each tumor, which were at least 3 mm apart, were collected. Areas with significant necrosis, fibrosis, or hemorrhage were avoided to maximize the viability of tumor cells. Normal colorectal mucosa tissues were also sampled from areas remote from the primary tumor (at least 2 cm distant from the tumor edge). Peri-intestinal nodules including lymph nodes present in the resected specimen were sampled. If there was malignancy appearance (the cut section appeared tan-gray and hard), after confirming the malignancy, a portion of the lymph nodes was sampled for diagnostic requirements. The remaining part was taken for this study. Each selected tissue block was split into two for snap freezing and formalin fixing respectively (mirrored FFPE sample). Fresh samples were placed in a 2 ml cryotube, and snap frozen with immediate immersion into liquid nitrogen before transferred to -80°C freezer for storage. Peripheral blood was collected and processed into EDTA anticoagulation tube. The tumor tissue samples from 68 patients were sequenced and analyzed after filtering according to the filtering pipeline, schematically presented in the CONSORT diagram (CONSORT flowchart, Supplementary Fig. S1). The workflow summarizing experiments and data analysis in our study was shown in Supplementary Fig. S2.

### Sample processing

Approximately 50 mm^3^ of tumor tissue from each region was used for genomic DNA extraction using the QIAamp DNA Mini Kit (Qiagen, Germany) according to the manufacturer’s instructions. 2 ml of peripheral blood was used for germline DNA extraction using the QIAamp DNA Blood midi kit (Qiagen, Germany) according to the manufacturer’s instructions. DNA was quantified by the Qubit Fluorometric Quantitation (Thermo Fisher Scientific, USA) and the quality of DNA was assessed by agarose gel electrophoresis.

### Pathology diagnoses and review

Pathological diagnoses were established according to the WHO classification and independently reviewed by two pathologists. Clinical details were summarized in Supplementary Table S1. Hematoxylin-eosin sections of mirrored FFPE samples for each region in every case (387 sections from 70 patients) were evaluated. Only primary tumor regions with more than 30% tumor component and pathological heterogeneity were considered for sequencing. In addition, pathologist distinguished LN and ENTD by reviewing hematoxylin-eosin sections of their mirrored FFPE samples in this study were also sent for sequencing.

### Whole exome library construction and sequencing

Tumor tissues and matched germline tissues were subjected to whole exome sequencing. Exome capture was performed on 1 μg of genomic DNA. Covaris (LE220) was used to randomly fragmented DNA into 150-250 bp. These fragments were purified and connected through a PE Index Adaptor designed by BGI, and then captured by using the the MGIeasy Exome Capture V4 probe set (∼ 59 Mb; MGI Tech Co., Ltd, China). All constructed libraries were loaded onto BGISEQ-500 (MGI Tech Co., Ltd, China) and the sequences were generated as 100-bp paired-end reads.

Sequencing reads containing sequencing adapters, more than 10% of unknown bases and low-quality bases (> 50% bases with quality <5) were removed by SOAPnuke (v1.5.6)^41^. The processed sequencing reads were then aligned to UCSC human reference genome (hg19) using BWA-MEM (v0.7.12)^42^. Picard (v1.137) (https://broadinstitute.github.io/picard/) was used to generate chromosomal coordinate-sorted bam files to remove PCR duplicates. Then, the median sequencing depth of the generated data for the tumor area were reached 391 (range 179-537), and the matched germline tissues were reached 414.5 (range 243-596). We then used the Genomic Analysis Toolkit (GATK v3.8.0)^43^ to perform base quality score recalibration and local realignment of the aligned reads to improve alignment accuracy.

### Quality control to prevent contamination, inter-patient sample swaps and removal of regions with extremely low mutation occurrence

ContEst^44^, a GATK module, was used to estimate the cross-individual contamination level. Samples with contamination level more than 1% were deleted (3 samples failed the QC due to contamination as shown in Supplementary Fig. S1. In order to avoid sample swaps between patients, we used BAM-matcher^45^.

The number of mutations in each tumor region was called independently. The median number of mutations across all regions for each tumor was calculated. A region in one tumor was removed if less than 20% of the median mutation count of that tumor was identified in that region.

### Somatic mutation detection and filtering

After processed the sequencing data, SAMtools (v1.2)^46^ mpileup was used to locate non-reference locations in tumor and germline samples. Bases with phred scores less than 20 or reads with mapping quality (MAPQ) values less than 20 were deleted. Base-alignment quality (BAQ) computation was disabled with adjust mapping quality coefficient set of 50. Both VarScan 2 (v2.4.3)^47^ and MuTect (v1.1.7)^48^ were used to call somatic mutations. The somatic variants called by VarScan 2 were filtered and the minimum coverage of the germline sample was set to 10, the minimum variant frequency was changed to 0.01, and tumor purity was set to 0.5. We further filtered the resulting single nucleotide variant (SNV) calls for false positives using Varscan 2 associated fpfilter.pl script. We used bam-readcount (v0.8.0) (https://github.com/genome/bam-readcount) to prepare input files for fpfilter and min-var-freq was set to 0.02. All insertions/deletions (INDELs) called in reads that VarScan 2 processSomatic classified as “high confidence” were recorded for further downstream filtering. MuTect was used to detect SNVs using annotation files contained in GATK bundle (v2.8) and variants were filtered according to the filter parameter ‘PASS’.

Additional filtering was performed to reduce false positive mutation calls. If the variant allele frequency (VAF) is greater than 2%, and both VarScan 2 (with a somatic p-value <= 0.01) and MuTect called the mutation, then a SNV was considered as truly positive. Alternatively, if a SNV was called only in VarScan 2 with a somatic p-value <=0.01, a frequency of 5% was required. In addition, the sequencing depth supporting the variant call in each region required >= 30, and the sequence reads required >= 5. In contrast, the VAF value of the variant in the germline should be <= 1%. We filtered the INDEL using the same parameters as above, except that reads >= 10 were required to support mutation calls, somatic p-values <= 0.001 and sequencing depth >= 50.

ANNOVAR^49^ was used to annotate mutations with COSMIC (v88)^50^, SIFT^51^, PolyPhen-2^52^ and MutationTaster^53^ databases. All mutations used in the analysis can be found in Supplementary Table S2. Mutations were classified as clonal or subclonal using PyClone (v0.13.1)^54^. PyClone CCF (cancer cell fraction) value were calculated as described in the subclonal deconstruction section. Mutations with CCF>0.9 across all regions of a tumor were considered as clonal mutations, otherwise they were considered as subclonal mutations.

### Driver mutation identification

All variants were compared with all genes identified and enlisted in the COSMIC Cancer Gene Census (v88)^50^. Then, three types of mutations were classified as a driver mutation according to the following criteria. Firstly, if the gene was annotated as TSG (tumor suppressor gene) by COSMIC, and the non-silent variant was considered deleterious: either *loss of function* (stop-gain/stop-loss, frameshift deletion/insertion or non-frameshift insertion/deletion) or predicted deleterious in two of these three computational approaches applied – SIFT^51^, PolyPhen-2^52^ and MutationTaster^53^, then the specific variant would be classified as a driver mutation. Secondly, if the variant was annotated as oncogene by COSMIC, then we tried to identify exact matches to non-silent variants in COSMIC. If an exact match was found ≥ 3 times, the variant was categorized as a driver mutation. Thirdly, if the gene was annotated as TSG by COSMIC, and the variant is located at the canonical splice site, then the specific variant would be classified as a driver mutation. Finally, we compared all these three types of driver mutations to the CpG island location file on UCSC Genome Bioinformatics website (http://genome.ucsc.edu). We then deleted all mutations that occurred on the CpG island and finally got all driver mutations.

### Copy number analysis

Sequenza (v3.0.0)^55^ was used to detect the somatic copy number alterations (SCNAs) and evaluate the purity and ploidy of tumor cells as follows. Firstly, we used SAMtools (v1.2)^46^ mpileup to convert the Bam file to Pileup format. Secondly, paired tumors and normal Pileup files were processed by sequenza-utils to extract the sequencing depth, determine the homozygous and heterozygous positions of variants in normal samples, and calculate the variant alleles and allelic frequencies from tumor samples. The sequenza-utils output was further processed by using Sequenza R package to provide segmented copy number data, cellularity and estimated ploidy for each sample. All segmented copy number data has been given in Supplementary Table S3. Heatmap of genome-wide SCNAs is visualized by R package Copynumber (v1.24.0)^56^.

The driver gene copy number variations (driver SCNAs) of all genes enlisted in the COSMIC cancer gene census were analyzed as follows. Firstly, if the gene was annotated as oncogene by COSMIC, gene level amplification was called if gene copy number >2 × ploidy of that sample. Secondly, if the gene was annotated as TSG by COSMIC, gene level deletion was called if gene copy number = 0. To determine the ITH status of driver SCNAs, we called driver SCNAs across all regions from each tumor. If at least one region showed an amplified SCNA, we called a gene as clonal amplification if all other regions of this gene showed copy number > ploidy + 1. If at least one region showed a deleted SCNA, we called a gene as clonal deletion if all other regions of this gene showed copy number < ploidy -1. All other driver SCNAs were defined as subclonal amplification or deletion. In 8 polyclonally originated tumors (CRC32, CRC36, CRC42, CRC48, CRC49, CRC51, CRC52 and CRC60) without founder clusters (cluster with CCF > 0.9 across all regions of a tumor), all their driver SCNAs were subclonal. To correlate driver SCNAs with specific mutation clusters of PyClone, we first identified all clusters where >= 50% CCF was present in each tumor region. We then identified all the clusters present in the same regions as a given driver SCNAs. We called a gene as clonally amplified if all the regions of this gene showed copy number >2 × ploidy while we called a gene as clonally deleted if all the regions of this gene showed copy number = 0. Then we repeated the association test above. If an SCNA still could not be associated with a mutant cluster, it was annotated as a subclone associated with no known cluster (NA cluster).

To determine the ITH status of global SCNA, all parts of the genome were considered independently and divided into the smallest contiguous segments that overlap in all the regions within each tumor. The gains and losses of segment were determined as follows. Firstly, copy number data for each segment was divided by the sample mean ploidy and then converted to log2. Secondly, gain and loss were defined as log2 (2.5/2) and log2 (1.5/2), respectively. Thirdly, any segment of gain or loss that spanned across all the regions was defined as clonal and all other segments of SCNA were defined as subclonal. Within each tumor, we summarized the length of the genome that subjected to SCNA in any region (total SCNA), the length of the genome that subjected to clonal SCNA (clonal gain or clonal loss), and the length of the genome that subjected to subclonal SCNA (subclonal gain, subclonal loss or subclonal undetermined). The proportion of subclonal SCNAs were then defined as the percentage of genomes subjected to subclonal SCNA divided by the percentage of genomes subjected to total SCNAs.

Chromosomal arm level SCNAs were determined if at least one region has shown an increase or decrease of at least 97% in chromosomal arm. To determine the ITH status of chromosome arm gain and loss, we called clonal arm gain or loss if the same chromosomal arm showed at least 75% gain or loss in all the remaining regions. While we called subclonal arm gain or loss if at least one of the remaining regions showed less than 75% gain or loss. In 8 polyclonally originated tumors, all their arm level SCNAs were subclonal. As previously described in the driver SCNAs part, we correlated arm level SCNAs with specific mutation clusters of PyClone in the same way.

### Mirrored sub-clonal allelic imbalance analysis

Single nucleotide polymorphisms (SNPs) were called by using Platypus (v0.8.1)^57^ and only SNPs with a minimum coverage of 20× were analyzed. The B allele frequency (BAF) of each SNP was calculated as the ratio of reads of reference base to variant. Heterozygous SNPs and BAFs were used as input and mirror subclone allelic imbalances (MSAI) were analyzed and visualized by RECUR^58^.

Parallel evolution events for driver SCNAs were identified as follows. Firstly, driver SCNAs were identified as described in the “copy number analysis” section. Secondly, we annotated the regions of MSAI events in each tumor to the events of driver SCNAs. If two events coincided with each other, then these driver SCNAs undergone parallel evolution.

### Sub-clonal deconstruction

In order to estimate whether mutations were clonal or subclonal, and the phylogenetic trees of each tumor, the following formula were used^22,59^:

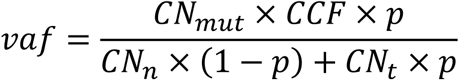

Where *vaf* is the mutated allele frequency of the mutated base; *p* is the estimated tumor purity; *CNt* is tumor locus specific copy number; *CNn* is normal locus specific copy number, assuming 2 for autosomal chromosomes; *CCF* is the fraction of tumor cells carrying mutations. Considering that *CNmut* is the copy number of the chromosome harboring the mutation, the possible *CNmut* range is from 1 to *CNt* (integer). We then assigned one of the possible values to *CCF*: 0.01, 0.02, …, 1, together with every possible *CNmut* to find the best fit *CCF* using maximum likelihood. In detail, for point mutations with alternative reads as “*a*” and sequencing coverage as “*N*”, we used Bayesian probability theory and binomial distribution to estimate the probability of a given *CCF*:

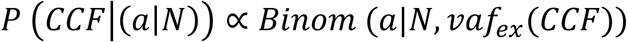

Then, the distribution of *CCF* was obtained by calculating *P (CCF)* on 100 uniform grids with *CCF* values from 0.01 to 1 and dividing by their sum.

Then, we used PyClone (v0.12.9)^54^ Dirichlet process clustering to cluster all the mutations (SNVs and INDELs). For each mutation, we used the observed mutation count and set the reference count so that vaf equal to half of the CCF value calculated by maximum likelihood previously. We set the major allele copy number to 2, the minor allele copy number to 0 and the purity to 0.5 since they had been modified.

Since the vaf values of INDELs were potentially unreliable, we multiplied each estimated INDEL CCF with a region-specific correction factor, which was calculated by dividing the median mutation CCF of the ubiquitous mutations (mutations presented in all regions) in that region by the median INDEL CCF of the ubiquitous INDELs (INDELs presented in all regions) in that region. We ran PyClone with 10,000 iterations and a burn-in of 1000.

### Phylogenetic tree construction

Phylogenetic trees were constructed using the published tool CITUP (v0.1.0)^60^. As input, CITUP requires mutation clusters and their mean cancer cell prevalence values which were collected from PyClone. All clusters with at least 5 mutations were used as input to CITUP. Clusters for phylogenetic tree construction were summarized in Supplementary Table S4. The optimal phylogenetic trees for each patient from CITUP were illustrated using MapScape (v1.8.0)^61^.

### Evolution subtype analysis

Evolutionary subtypes were clustered and visualized by REVOLVER (v0.2.0)^32^. CCF values and cluster information of driver events were processed as previously described, which were used as input to REVOLVER. REVOLVER requires a founder cluster for all the input tumors. Therefore, we artificially defined a founder cluster for 8 polyclonally originated tumors. ITH index was calculated as the numbers of subclonal driver events divided by the numbers of clonal driver events, and SCNA index was indicated by the length of total SCNA.

### Phylogenetic analysis

Phylogenetic distance between primary tumor, LN and ENTD were analyzed by using the binary matrix of mutations present or absent in each region of tumors with LN or ENTD. Private mutations of each region were discarded from phylogenetic tree building due to lack of information. Fake outgroups with no mutations were generated for each individual as a root. Phylogenies were constructed using the PHYLIP (v3.697)^62^ suite of tools. For each tumor, we used seqboot to generate 100 bootstrap replicates by resampling of the mutations with replacement.

Phylogenetic trees were then constructed for each bootstrap replicate by maximum parsimony using the Mix programme in Wagner method. The jumble = 10 option was used and the order of the input samples was randomized 10 times for each bootstrap replicate. Finally, the Consense program was used to build a consensus of all the phylogenetic trees by using the majority rule (extended) option. Phylogenetic trees were redrawn by FigTree (v1.4.4)^63^ with the length of trunks and branches, proportional to the number of mutations.

### dN/dS analysis

Values of dN/dS were analysed for different types of mutations: missense (wmis), nonsense (wnon), essential splice site substitution (wspl), non-synonymous substitutions (wall) and truncating substitutions (wtru). Values of dN/dS were analyzed by dNdScv R package^64^.

### Mutation signature analysis

Mutation signatures were estimated by using the DeconstructSigs (v1.8.0)^65^ package in R. Mutational signature analysis was applied only in the presence of at least 15 mutations.

### Statistical analysis

All analyses were performed in R statistical environment version >= 3.5.0. All statistical comparisons of two distributions used the Wilcoxon test (wilcox.test function in R).

### Data availability

The sequencing data as been deposited at the CNGB Nucleotide Sequence Archive (CNSA: https://db.cngb.org/cnsa), under accession number CNP0000594.

## Acknowledgement

We are thankful to the proband and all the family members for participating in our study and we are thankful to the China National GeneBank.

## Author contributions

Study concept and design: Santasree Banerjee, Shan Kuang, Junnian Liu, Yun Lu and Xin Liu; Patient recruitment and clinical sample collection: Xianxiang Zhang, Qingyao Wu, Shujian Yang; Histology and histopathology: Jigang Wang and Xiaobin Ji; Experiments (DNA extraction and whole exome sequencing): Peng Han, Yong Li, Xiaofen Tian and Zhiwei Wang; Analysis and interpretation of data: Santasree Banerjee, Shan Kuang, Lei Li, Shui Shun, Li Deng and Yue Zhang; Drafting of the manuscript: Santasree Banerjee, Shan Kuang, Lei Li, Xianxiang Zhang and Jigang Wang; Critical revision of the manuscript for important intellectual content: Huanming Yang, Lars Bolund, Yonglun Luo, Kui Wu, Shida Zhu, Guangyi Fan and Xun Xu; Supervision of the study: Santasree Banerjee, Shan Kuang, Junnian Liu, Yun Lu and Xin Liu.

## References

1. Alizadeh, A.A. et al. Toward understanding and exploiting tumor heterogeneity. Nature medicine 21, 846 (2015).

2. Turner, N.C. & Reis-Filho, J.S. Genetic heterogeneity and cancer drug resistance. The lancet oncology 13, e178–e185 (2012).

3. Waddell, N. et al. Whole genomes redefine the mutational landscape of pancreatic cancer. Nature 518, 495–501 (2015).

4. Burrell, R.A., McGranahan, N., Bartek, J. & Swanton, C. The causes and consequences of genetic heterogeneity in cancer evolution. Nature 501, 338 (2013).

5. World Health Organization. Cancer. https://www.who.int/news-room/fact-sheets/detail/cancer. Accessed January 21, 2019. (2019).

6. International Agency for Research on Cancer. Cancer Today. https://gco.iarc.fr/today/. Accessed January 21, 2019. (2019).

7. O’Connell, J.B., Maggard, M.A. & Ko, C.Y. Colon cancer survival rates with the new American Joint Committee on Cancer sixth edition staging. Journal of the National Cancer Institute 96, 1420–1425 (2004).

8. Adjuvant therapy for patients with colon and rectal cancer. JAMA 264, 1444–1450 (1990).

9. Loupakis, F. et al. Primary tumor location as a prognostic factor in metastatic colorectal cancer. JNCI: Journal of the National Cancer Institute 107 (2015).

10. Petrelli, F. et al. Prognostic survival associated with left-sided vs right-sided colon cancer: a systematic review and meta-analysis. JAMA oncology 3, 211–219 (2017).

11. Missiaglia, E. et al. Distal and proximal colon cancers differ in terms of molecular, pathological, and clinical features. Annals of oncology 25, 1995–2001 (2014).

12. Iacopetta, B. Are there two sides to colorectal cancer? International journal of cancer 101, 403–408 (2002).

13. Sottoriva, A. et al. A Big Bang model of human colorectal tumor growth. Nature genetics 47, 209 (2015).

14. Wei, Q. et al. Multiregion whole-exome sequencing of matched primary and metastatic tumors revealed genomic heterogeneity and suggested polyclonal seeding in colorectal cancer metastasis. Annals of Oncology 28, 2135–2141 (2017).

15. Mamlouk, S. et al. DNA copy number changes define spatial patterns of heterogeneity in colorectal cancer. Nature communications 8, 14093 (2017).

16. Roerink, S.F. et al. Intra-tumour diversification in colorectal cancer at the single-cell level. Nature 556, 457 (2018).

17. Uchi, R. et al. Integrated multiregional analysis proposing a new model of colorectal cancer evolution. PLoS genetics 12, e1005778 (2016).

18. Alves, J.M., Prado-López, S., Cameselle-Teijeiro, J.M. & Posada, D. Rapid evolution and biogeographic spread in a colorectal cancer. Nature communications 10, 5139 (2019).

19. Ishaque, N. et al. Whole genome sequencing puts forward hypotheses on metastasis evolution and therapy in colorectal cancer. Nature communications 9, 4782 (2018).

20. Sun, J. et al. Genomic signatures reveal DNA damage response deficiency in colorectal cancer brain metastases. Nature communications 10, 3190 (2019).

21. Dunne, P.D. et al. Cancer-cell intrinsic gene expression signatures overcome intratumoural heterogeneity bias in colorectal cancer patient classification. Nature communications 8, 15657 (2017).

22. Jamal-Hanjani, M. et al. Tracking the evolution of non–small-cell lung cancer. New England Journal of Medicine 376, 2109–2121 (2017).

23. Turajlic, S. et al. Tracking cancer evolution reveals constrained routes to metastases: TRACERx Renal. Cell 173, 581-594. e12 (2018).

24. Turajlic, S. et al. Deterministic evolutionary trajectories influence primary tumor growth: TRACERx renal. Cell 173, 595-610. e11 (2018).

25. Ling, S. et al. Extremely high genetic diversity in a single tumor points to prevalence of non-Darwinian cell evolution. Proceedings of the National Academy of Sciences 112, E6496–E6505 (2015).

26. Nikbakht, H. et al. Spatial and temporal homogeneity of driver mutations in diffuse intrinsic pontine glioma. Nature communications 7, 11185 (2016).

27. Kumar, A. et al. Substantial interindividual and limited intraindividual genomic diversity among tumors from men with metastatic prostate cancer. Nature medicine 22, 369 (2016).

28. Zhai, W. et al. The spatial organization of intra-tumour heterogeneity and evolutionary trajectories of metastases in hepatocellular carcinoma. Nature communications 8, 4565 (2017).

29. Network, C.G.A. Comprehensive molecular characterization of human colon and rectal cancer. Nature 487, 330 (2012).

30. Alexandrov, L.B. et al. Signatures of mutational processes in human cancer. Nature 500, 415–21 (2013).

31. Bielski, C.M. et al. Genome doubling shapes the evolution and prognosis of advanced cancers. Nature genetics 50, 1189 (2018).

32. Caravagna, G. et al. Detecting repeated cancer evolution from multi-region tumor sequencing data. Nature methods 15, 707 (2018).

33. Williams, M.J., Werner, B., Barnes, C.P., Graham, T.A. & Sottoriva, A. Identification of neutral tumor evolution across cancer types. Nature genetics 48, 238 (2016).

34. Loeb, L.A. et al. Extensive subclonal mutational diversity in human colorectal cancer and its significance. Proceedings of the National Academy of Sciences (2019).

35. Saito, T. et al. A temporal shift of the evolutionary principle shaping intratumor heterogeneity in colorectal cancer. Nature communications 9, 2884 (2018).

36. Lee, M.S., Menter, D.G. & Kopetz, S. Right versus left colon cancer biology: integrating the consensus molecular subtypes. Journal of the National Comprehensive Cancer Network 15, 411–419 (2017).

37. Hu, Z. et al. Quantitative evidence for early metastatic seeding in colorectal cancer. Nature genetics, 1 (2019).

38. Árnadóttir, S.S. et al. Characterization of genetic intratumor heterogeneity in colorectal cancer and matching patient-derived spheroid cultures. Molecular oncology 12, 132–147 (2018).

39. Dienstmann, R. et al. Consensus molecular subtypes and the evolution of precision medicine in colorectal cancer. Nature Reviews Cancer 17, 79 (2017).

40. Punt, C.J., Koopman, M. & Vermeulen, L. From tumour heterogeneity to advances in precision treatment of colorectal cancer. Nature reviews Clinical oncology 14, 235 (2017).

41. Chen, Y. et al. SOAPnuke: a MapReduce acceleration-supported software for integrated quality control and preprocessing of high-throughput sequencing data. Gigascience 7, 1–6 (2018).

42. Li, H. & Durbin, R. Fast and accurate short read alignment with Burrows-Wheeler transform. Bioinformatics 25, 1754–60 (2009).

43. McKenna, A. et al. The Genome Analysis Toolkit: a MapReduce framework for analyzing next-generation DNA sequencing data. Genome Res. 20, 1297–303 (2010).

44. Cibulskis, K. et al. ContEst: estimating cross-contamination of human samples in next-generation sequencing data. Bioinformatics 27, 2601–2602 (2011).

45. Wang, P.P., Parker, W.T., Branford, S. & Schreiber, A.W. BAM-matcher: a tool for rapid NGS sample matching. Bioinformatics 32, 2699–701 (2016).

46. Li, H. et al. The Sequence Alignment/Map format and SAMtools. Bioinformatics 25, 2078–9 (2009).

47. Koboldt, D.C. et al. VarScan 2: somatic mutation and copy number alteration discovery in cancer by exome sequencing. Genome Res. 22, 568–76 (2012).

48. Cibulskis, K. et al. Sensitive detection of somatic point mutations in impure and heterogeneous cancer samples. Nat. Biotechnol. 31, 213–9 (2013).

49. Wang, K., Li, M. & Hakonarson, H. ANNOVAR: functional annotation of genetic variants from high-throughput sequencing data. Nucleic Acids Res. 38, e164 (2010).

50. Forbes, S.A. et al. COSMIC: exploring the world’s knowledge of somatic mutations in human cancer. Nucleic Acids Res. 43, D805–11 (2015).

51. Kumar, P., Henikoff, S. & Ng, P.C. Predicting the effects of coding non-synonymous variants on protein function using the SIFT algorithm. Nature protocols 4, 1073 (2009).

52. Adzhubei, I., Jordan, D.M. & Sunyaev, S.R. Predicting functional effect of human missense mutations using PolyPhen-2. Current protocols in human genetics 76, 7.20. 1-7.20. 41 (2013).

53. Schwarz, J.M., Rödelsperger, C., Schuelke, M. & Seelow, D. MutationTaster evaluates disease-causing potential of sequence alterations. Nature methods 7, 575 (2010).

54. Roth, A. et al. PyClone: statistical inference of clonal population structure in cancer. Nat. Methods 11, 396–8 (2014).

55. Favero, F. et al. Sequenza: allele-specific copy number and mutation profiles from tumor sequencing data. Ann. Oncol. 26, 64–70 (2015).

56. Nilsen, G. et al. Copynumber: Efficient algorithms for single- and multi-track copy number segmentation. BMC Genomics 13, 591 (2012).

57. Rimmer, A. et al. Integrating mapping-, assembly- and haplotype-based approaches for calling variants in clinical sequencing applications. Nat. Genet. 46, 912–918 (2014).

58. Jakubek, Y.A., San Lucas, F.A. & Scheet, P. Directional allelic imbalance profiling and visualization from multi-sample data with RECUR. Bioinformatics 35, 2300–2302 (2019).

59. Zhang, H. et al. Sex difference of mutation clonality in diffuse glioma evolution. Neurooncology 21, 201–213 (2019).

60. Malikic, S., McPherson, A.W., Donmez, N. & Sahinalp, C.S. Clonality inference in multiple tumor samples using phylogeny. Bioinformatics 31, 1349–56 (2015).

61. Smith, M.A. et al. E-scape: interactive visualization of single-cell phylogenetics and cancer evolution. Nat. Methods 14, 549–550 (2017).

62. Falenstein, J. PHYLIP—Phylogeny inference packages (version 3.2). Cladistics 5, 164–166 (1989).

63. Rambaut, A. FigTree, a graphical viewer of phylogenetic trees. (2007).

64. Martincorena, I. et al. Universal Patterns of Selection in Cancer and Somatic Tissues. Cell 171, 1029-1041.e21 (2017).

65. Rosenthal, R., McGranahan, N., Herrero, J., Taylor, B.S. & Swanton, C. DeconstructSigs: delineating mutational processes in single tumors distinguishes DNA repair deficiencies and patterns of carcinoma evolution. Genome Biol. 17, 31 (2016).

