## Supplementary Figures for "Intratumor Heterogeneity and Evolution of Colorectal Cancer"

#### Figure S1

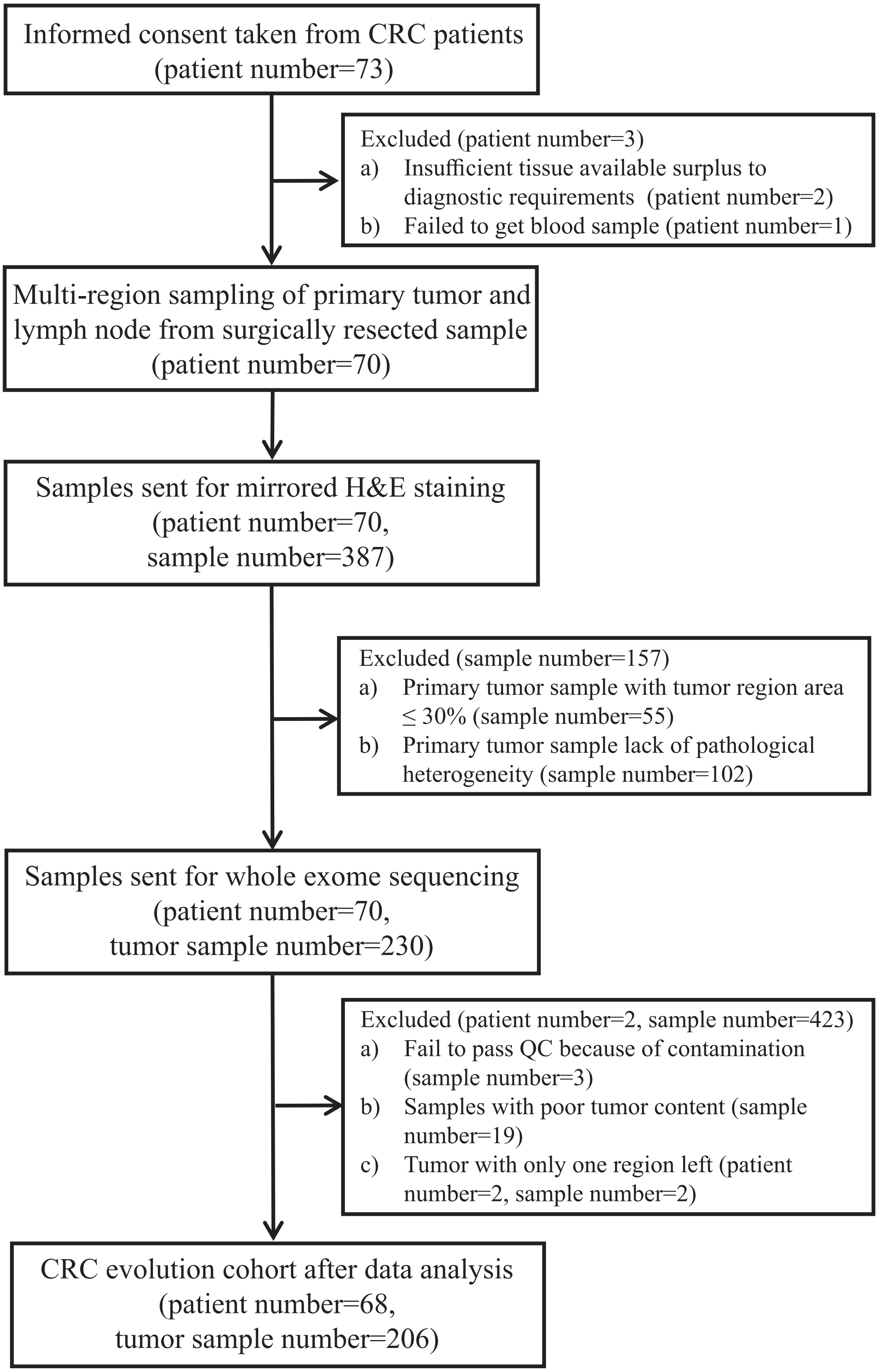

**Figure S1. CONSORT diagram for patient recruitment in this study and eventual selection of the 68 patients cohort.**

**Figure S2**

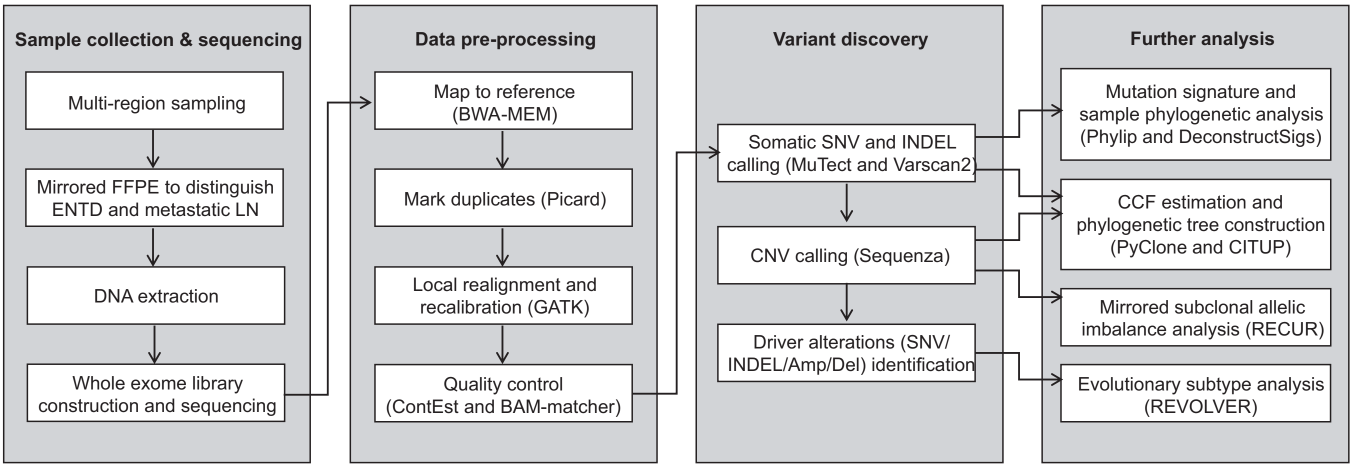

**Figure S2. Workflow summarizing experiments and data analysis.**
Overview of experiments and analysis workflow based on whole-exome sequencing of multi-region CRC tumors (primary tumors, lymph node metastasis and ENTDs) and paired normal controls. Analysis tools/methods are indicated by parentheses.

**Figure S3**

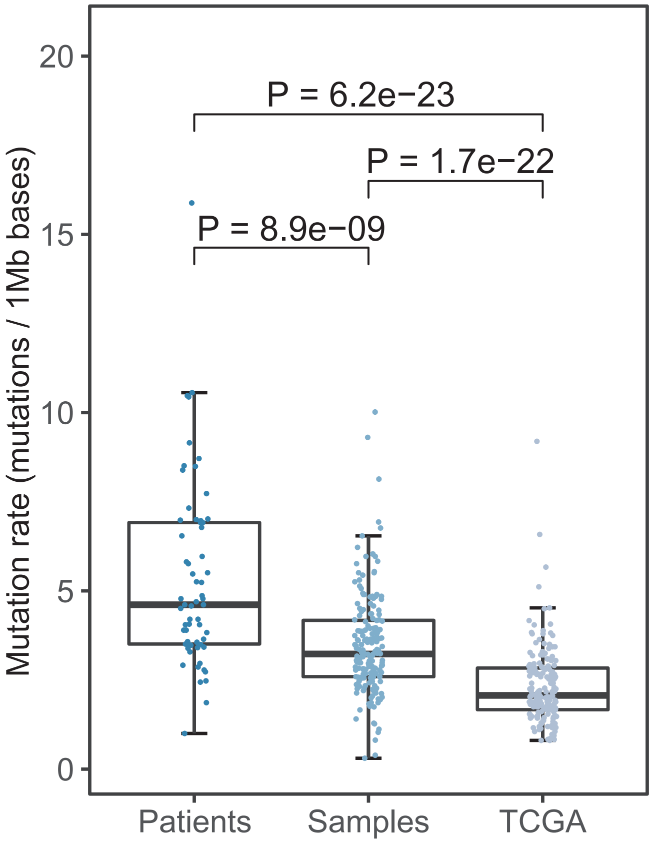

**Figure S3. Comparison of tumor mutation rate between TCGA and CRC tumors.** Box plots of mutation rate in non-hypermutated CRC patients analyzed as single samples, multi-region samples of non-hypermutated CRC patients and TCGA non-hypermutated CRC samples. The definition of hypermutated patients is all the samples in these patients have more than 10 mutations/1 Mb bases. In our CRC tumors, 6 patients are hypermutated patients and all other 62 patients are included into analysis.

#### Figure S4

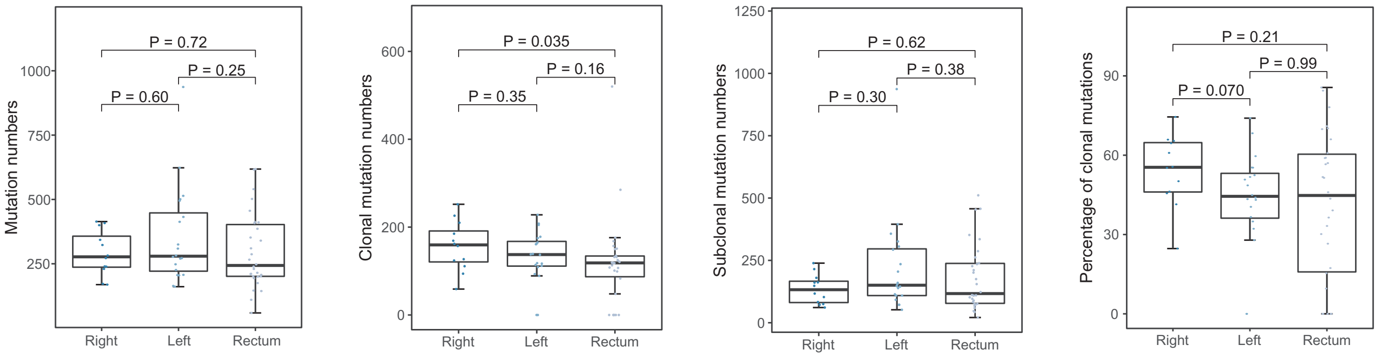

**Figure S4. Intratumor heterogeneity of mutations among right-sided colon, left-sided colon and rectal cancers.** Box plots of total, clonal, subclonal and percentage of clonal mutations by tumor position.

#### Figure S5

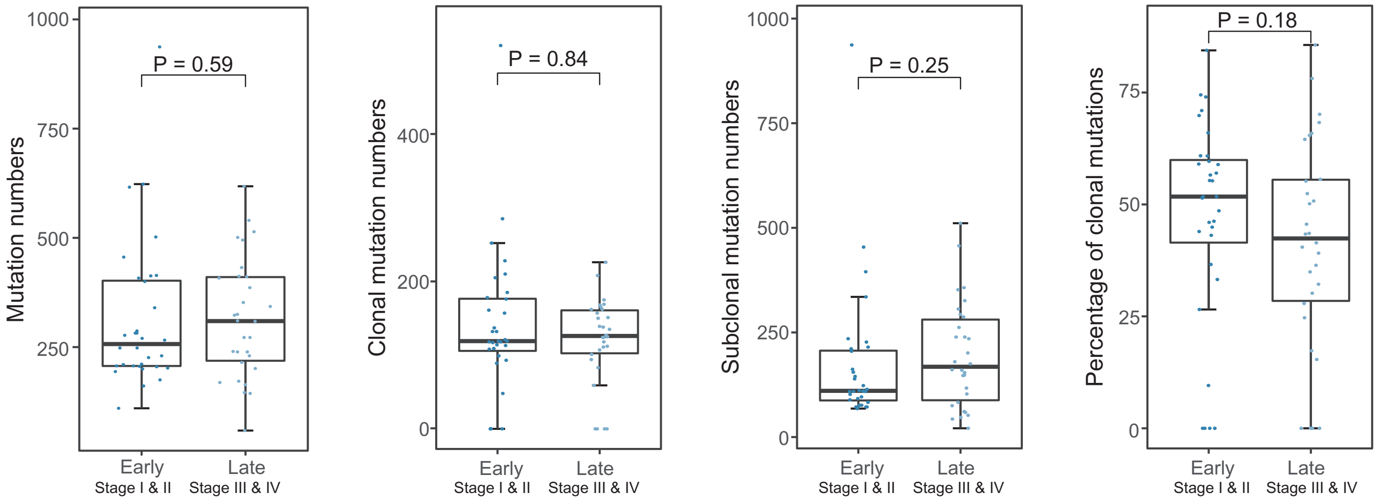

**Figure S5. Intratumor heterogeneity of mutations between early and late stage of CRC tumors.** Box plots of total, clonal, subclonal and percentage of clonal mutations by early (stage I and II) and late (stage III and IV) stage of CRC tumors.

**Figure S6**

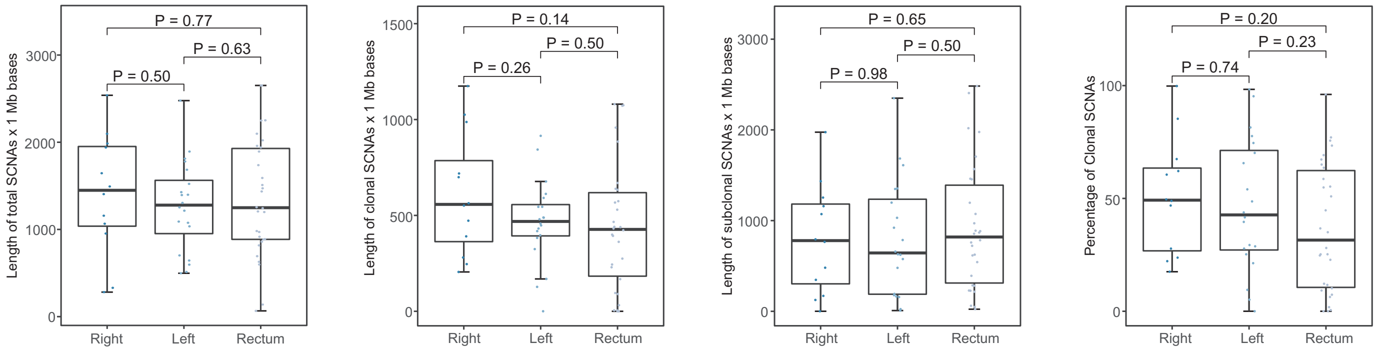

**Figure S6. Intratumor heterogeneity of somatic copy number alterations (SCNAs) between right-sided colon, left-sided colon and rectal cancers.** Box plots of total, clonal, subclonal and percentage of clonal SCNAs by tumor position.

#### Figure S7

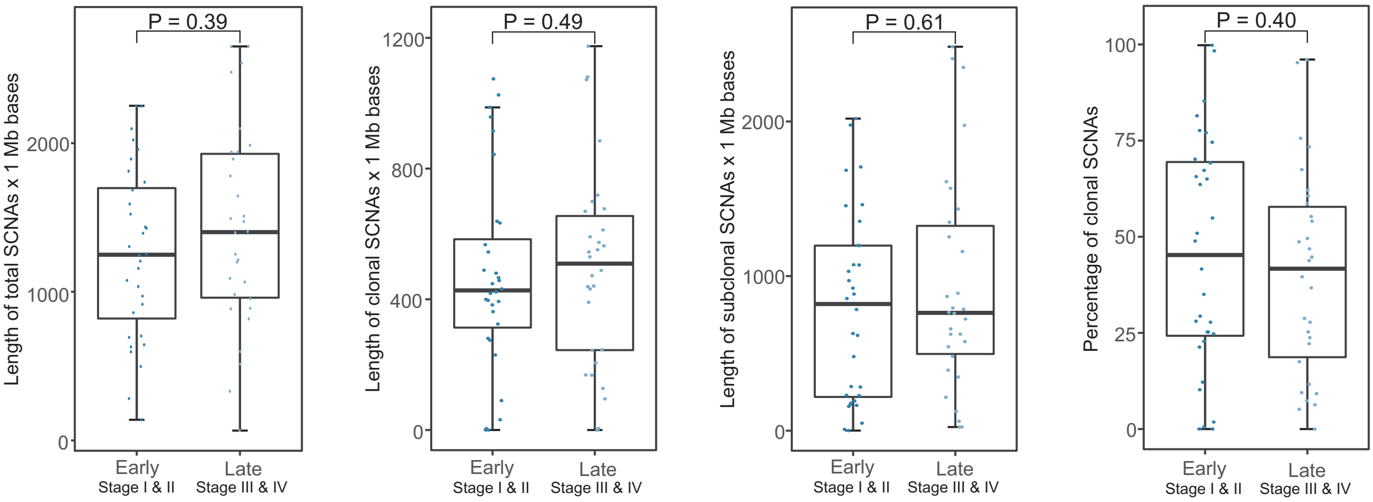

**Figure S7. Intratumor heterogeneity of SCNAs between early and late stage of CRC tumors.** Box plots of total, clonal, subclonal and percentage of clonal SCNAs by tumor stage.

### Figure S8

**Figure S8. Phylogenetic trees.**

Phylogenetic trees for 62 non-hypermutated CRC tumors. For each multi-panel figure, top-left panel showed cancer cell fractions (CCF) as a heatmap for all clustered mutations. CCF value for each mutation represents the mean of the mutation cluster CCF values, with darker color indicating higher CCF value. Regions were indicated below heatmap, "TR" represented primary tumor regions, "LN" represented lymph node metastasis regions and "LN_ENTD" represented extranodal tumour deposits regions. Middle top panel showed the complete phylogenetic tree as constructed based on the mutation clusters. Cancer driver genes found in the tumor, with cytoband, type [mutation (SNV and InDel)/copy number aberration (Amp and Del)] and mutation cluster were indicated on the right, with a colored bar representing individual clusters of driver genes. Below were phylogenetic trees drawn for each region. Grid of 100 representative cells were shown beneath the tree. The colors within each cell represented the mutational clusters.

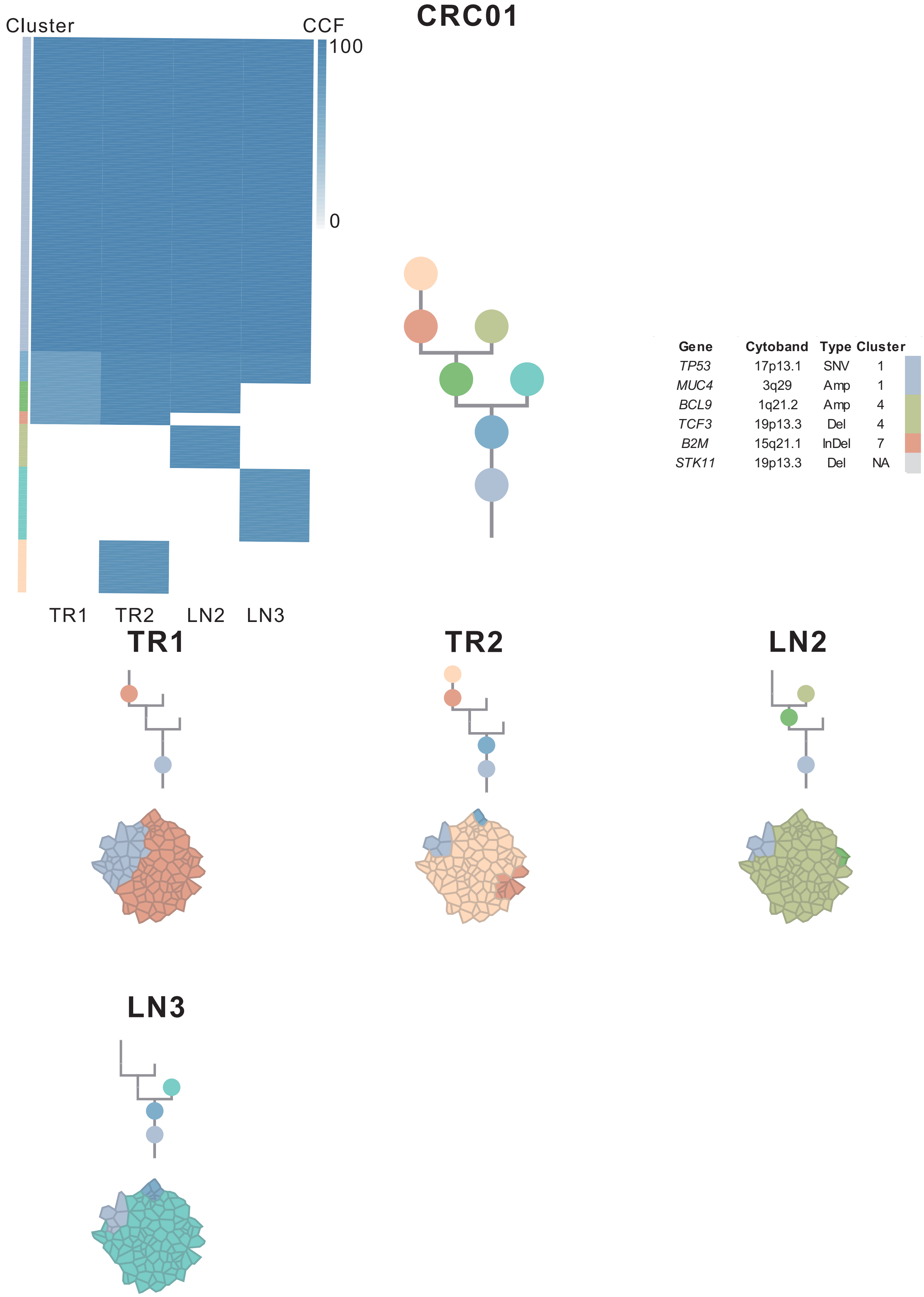
CRC01
Cluster
CCF
100
0
TR1
TR2
LN2
LN3
Gene Cytoband Type Cluster
TP53 17p13.1 SNV 1
MUC4 3q29 Amp 1
BCL9 1q21.2 Amp 4
TCF3 19p13.3 Del 4
B2M 15q21.1 InDel 7
STK11 19p13.3 Del NA
TR1
TR2
LN2
LN3

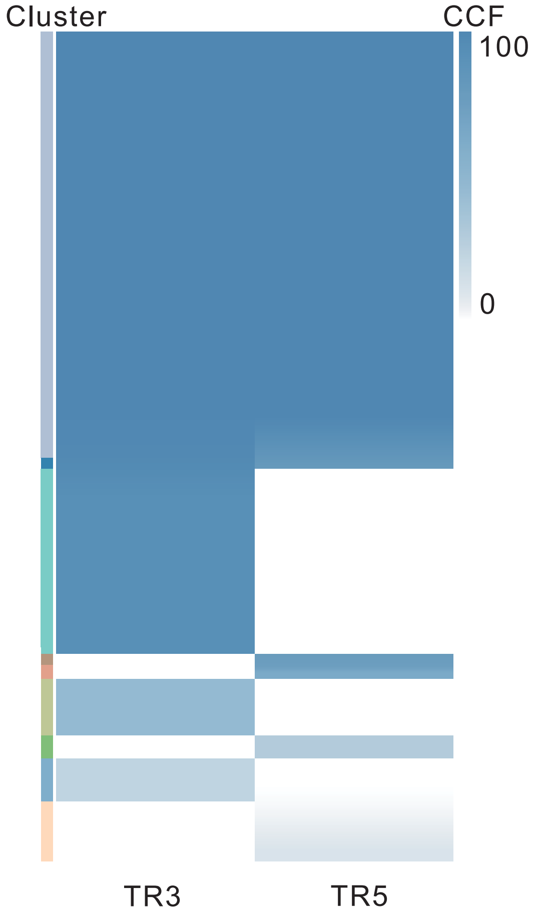

### CRC02

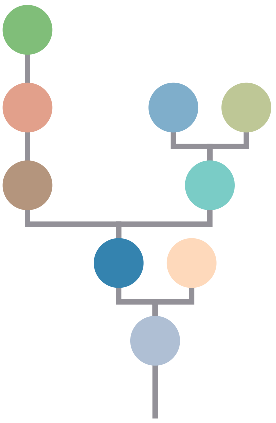

| Gene | Cytoband | Type | Cluster |
|---|---|---|---|
| * *APC* | 5q22.2 | SNV | 1 |
| *CUX1* | 7q22.1 | SNV | 1 |
| *TP53* | 17p13.1 | SNV | 1 |
| * *APC* | 5q22.2 | InDel | 1 |
| *FGFR1* | 8p11.23 | Amp | 1 |
| *CDKN2A* | 9p21.3 | Del | 1 |
| *AMER1* | Xq11.2 | SNV | NA |
| *ATRX* | Xq21.1 | SNV | NA |

## TR3

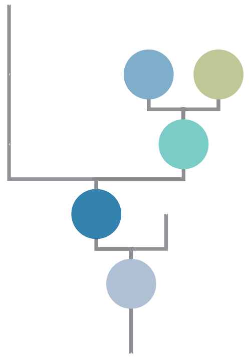

## TR5

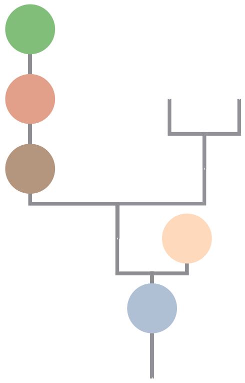

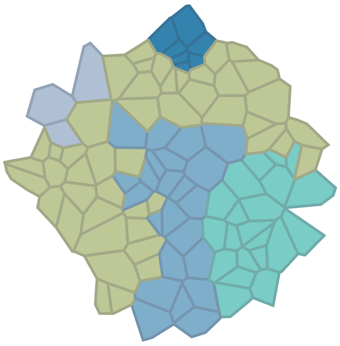

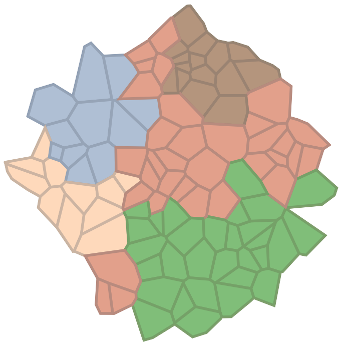

### CRC03

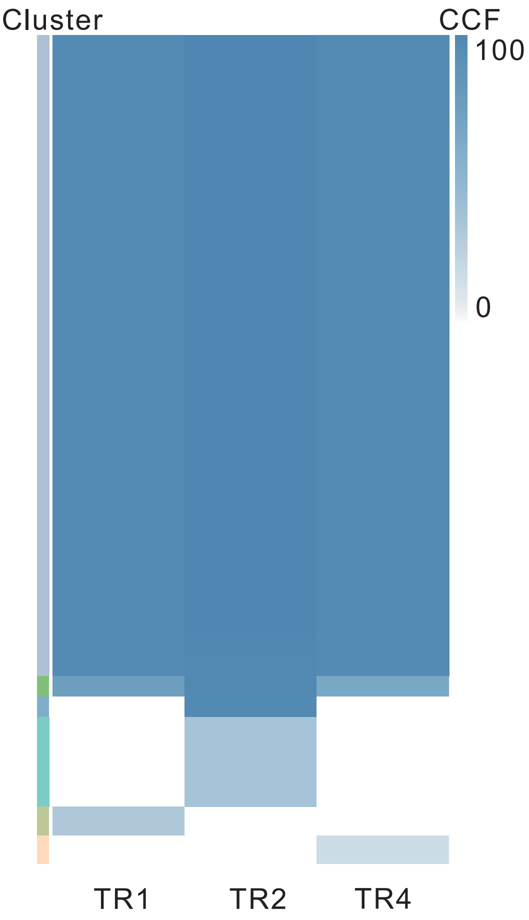

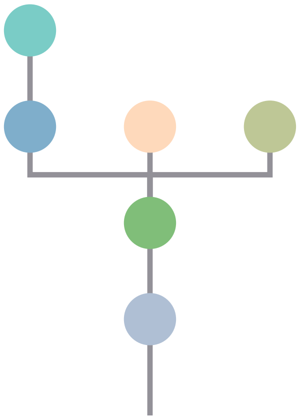

| Gene | Cytoband | Type | Cluster |
|---|---|---|---|
| *APC* | 5q22.2 | SNV | 1 |
| *KRAS* | 12p12.1 | SNV | 1 |
| *ETV6* | 12p13.2 | SNV | 1 |
| *CSF3R* | 1p34.3 | InDel | 6 |
| *AMER1* | Xq11.2 | SNV | NA |
| *PTEN* | 10q23.31 | Del | NA |

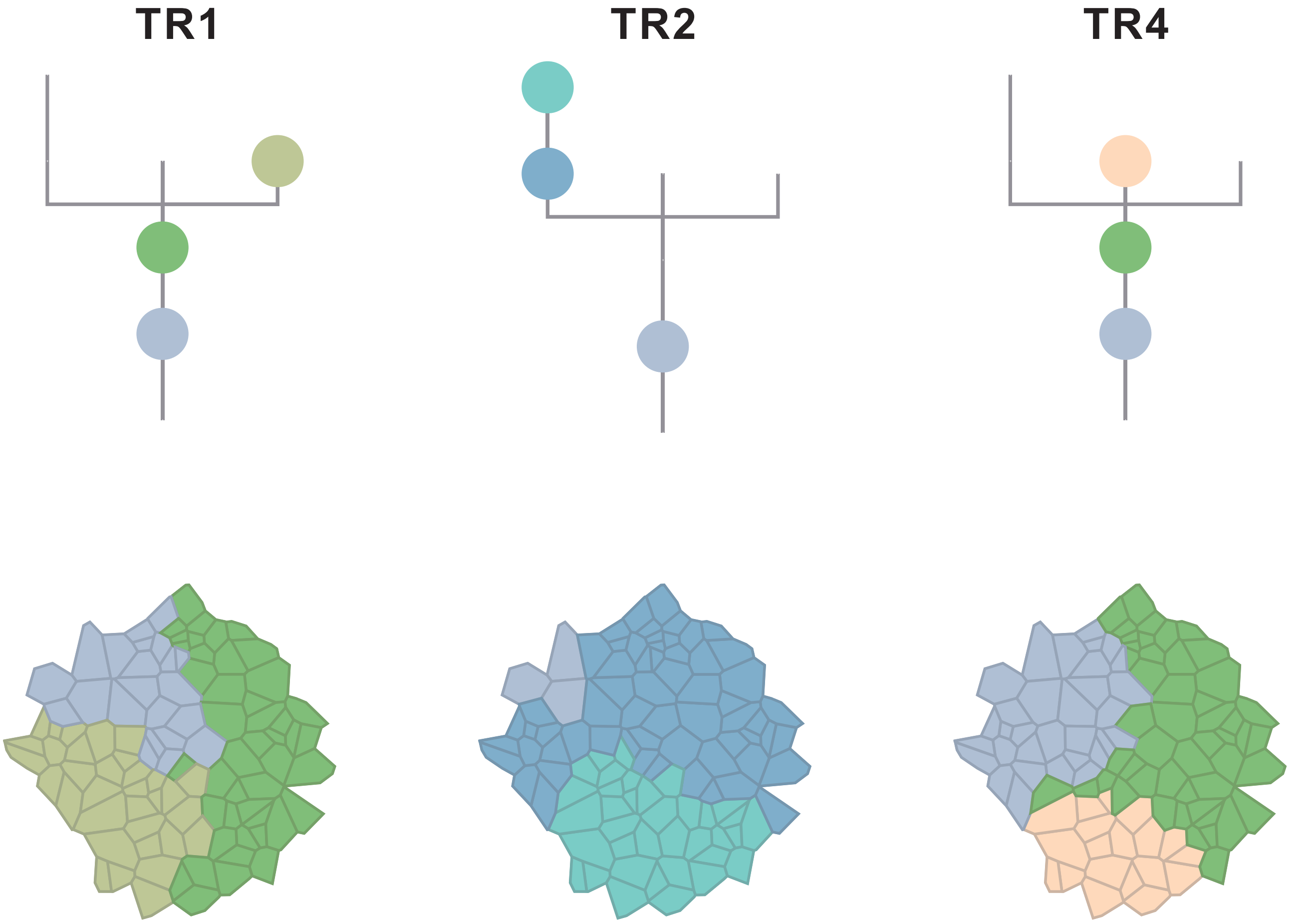

### CRC06

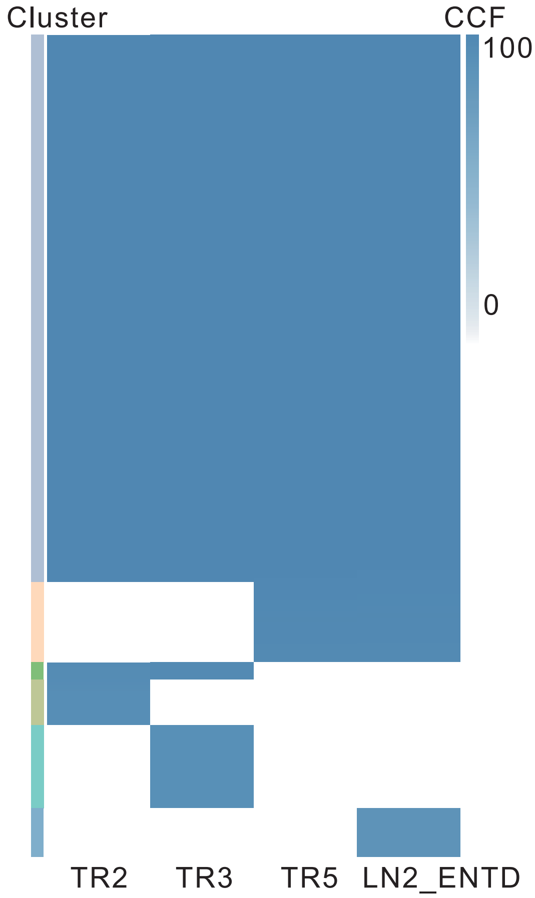

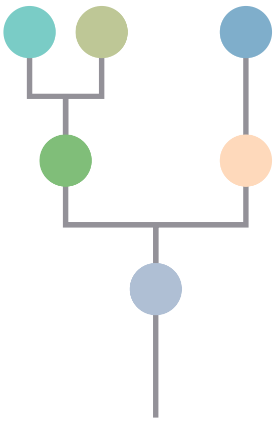

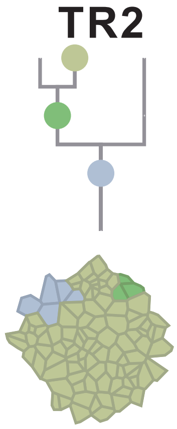

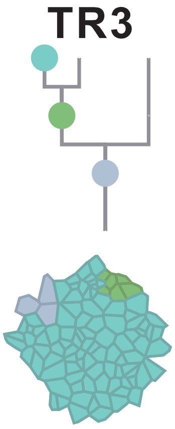

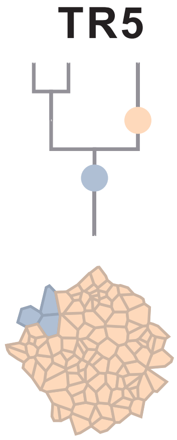

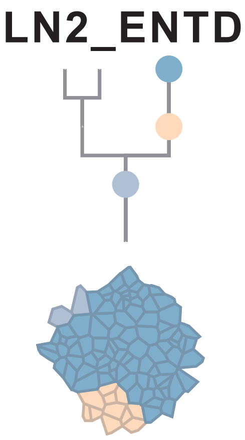

| Gene | Cytoband | Type | Cluster |
|---|---|---|---|
| * *PIK3CA* | 3q26.32 | SNV | 1 |
| * *PIK3CA* | 3q26.32 | SNV | 1 |
| *KRAS* | 12p12.1 | SNV | 1 |
| *ERCC2* | 19q13.32 | SNV | 1 |
| *APC* | 5q22.2 | InDel | 1 |
| *SSX2* | Xp11.22 | Amp | 1 |
| *ARAF* | Xp11.23 | Amp | 1 |
| *GATA1* | Xp11.23 | Amp | 1 |
| *SSX1* | Xp11.23 | Amp | 1 |
| *SSX4* | Xp11.23 | Amp | 1 |
| *TFE3* | Xp11.23 | Amp | 1 |
| *WAS* | Xp11.23 | Amp | 1 |
| *KDM6A* | Xp11.3 | Amp | 1 |
| *FOXO4* | Xq13.1 | Amp | 1 |
| *BTK* | Xq22.1 | Amp | 1 |
| *IRS4* | Xq22.3 | Amp | 1 |
| *BCORL1* | Xq26.1 | Amp | 1 |
| *ELF4* | Xq26.1 | Amp | 1 |
| *GPC3* | Xq26.2 | Amp | 1 |
| *CAMTA1* | 1p36.31 | Del | 1 |
| *RPL22* | 1p36.31 | Del | 1 |
| *TNFRSF14* | 1p36.32 | Del | 1 |
| *NOTCH1* | 9q34.3 | Del | 1 |
| *AXIN1* | 16p13.3 | Del | 1 |
| *CREBBP* | 16p13.3 | Del | 1 |
| *NTHL1* | 16p13.3 | Del | 1 |
| *TRAF7* | 16p13.3 | Del | 1 |
| *TSC2* | 16p13.3 | Del | 1 |
| *FLCN* | 17p11.2 | Del | 1 |
| *NCOR1* | 17p11.2 | Del | 1 |
| *MAP2K4* | 17p12 | Del | 1 |
| *PER1* | 17p13.1 | Del | 1 |
| *TP53* | 17p13.1 | Del | 1 |
| *YWHAE* | 17p13.3 | Del | 1 |
| *SMAD4* | 18q21.2 | InDel | 3 |
| *CARS* | 11p15.4 | Del | 3 |
| *NOTCH2* | 1p12 | Amp | 4 |
| *ARNT* | 1q21.3 | Amp | 4 |
| *SETDB1* | 1q21.3 | Amp | 4 |
| *STK11* | 19p13.3 | Del | 5 |
| *TCF3* | 19p13.3 | Del | 5 |
| *SDHA* | 5p15.33 | Del | 5 |
| *TERT* | 5p15.33 | Del | 5 |
| *TP53* | 17p13.1 | SNV | NA |

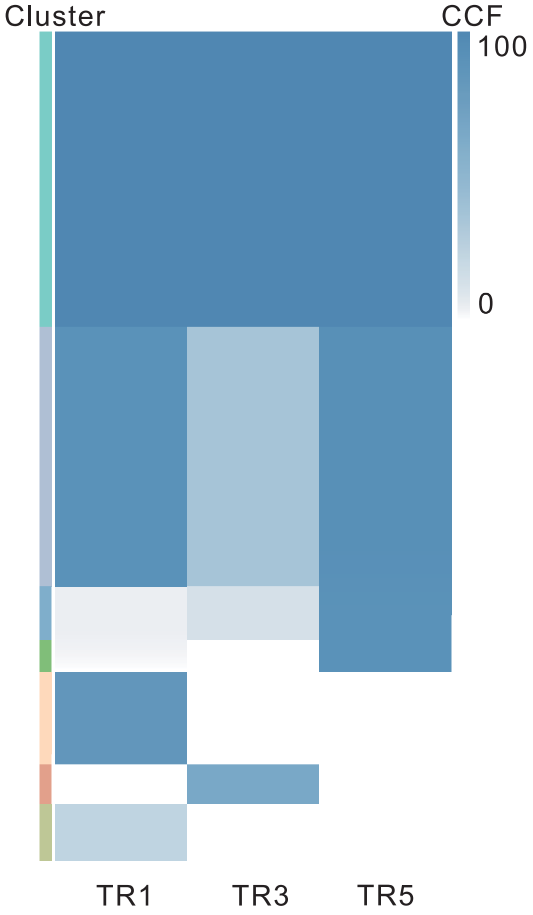

### CRC07

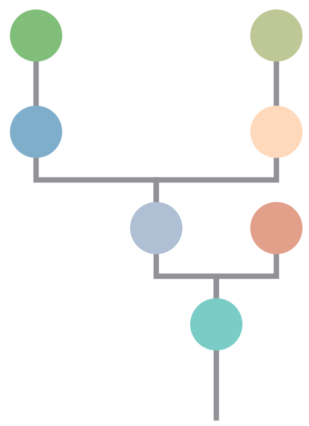

| Gene | Cytoband | Type | Cluster |
|---|---|---|---|
| * APC | 5q22.2 | SNV | 1 |
| TBL1XR1 | 3q26.32 | SNV | 2 |
| KRAS | 12p12.1 | SNV | 2 |
| TP53 | 17p13.1 | SNV | 2 |
| * APC | 5q22.2 | InDel | 2 |
| RUNX1 | 21q22.12 | Del | 2 |
| CLTCL1 | 22q11.21 | Del | 2 |
| LZTR1 | 22q11.21 | Del | 2 |
| SMARCB1 | 22q11.23 | Del | 2 |
| CHEK2 | 22q12.1 | Del | 2 |
| ZNRF3 | 22q12.1 | Del | 2 |
| NF2 | 22q12.2 | Del | 2 |
| MYH9 | 22q12.3 | Del | 2 |
| APOBEC3B | 22q13.1 | Del | 2 |
| EP300 | 22q13.2 | Del | 2 |
| MKL1 | 22q13.2 | Del | 2 |
| FOXO4 | Xq13.1 | Del | 2 |
| MED12 | Xq13.1 | Del | 2 |
| ZMYM3 | Xq13.1 | Del | 2 |
| ATRX | Xq21.1 | Del | 2 |
| BTK | Xq22.1 | Del | 2 |
| IRS4 | Xq22.3 | Del | 2 |
| STAG2 | Xq25 | Del | 2 |
| BCORL1 | Xq26.1 | Del | 2 |
| ELF4 | Xq26.1 | Del | 2 |
| GPC3 | Xq26.2 | Del | 2 |
| PHF6 | Xq26.2 | Del | 2 |
| ATP2B3 | Xq28 | Del | 2 |
| RPL10 | Xq28 | Del | 2 |
| ERBB4 | 2q34 | SNV | 3 |
| RNF43 | 17q22 | SNV | 4 |
| ATP1A1 | 1p13.1 | Amp | 5 |
| MUC4 | 3q29 | Amp | 5 |
| SIRPA | 20p13 | Del | 7 |
| ASXL1 | 20q11.21 | Del | 7 |
| PTPRT | 20q13.11 | Del | 7 |
| PTK6 | 20q13.33 | Del | 7 |

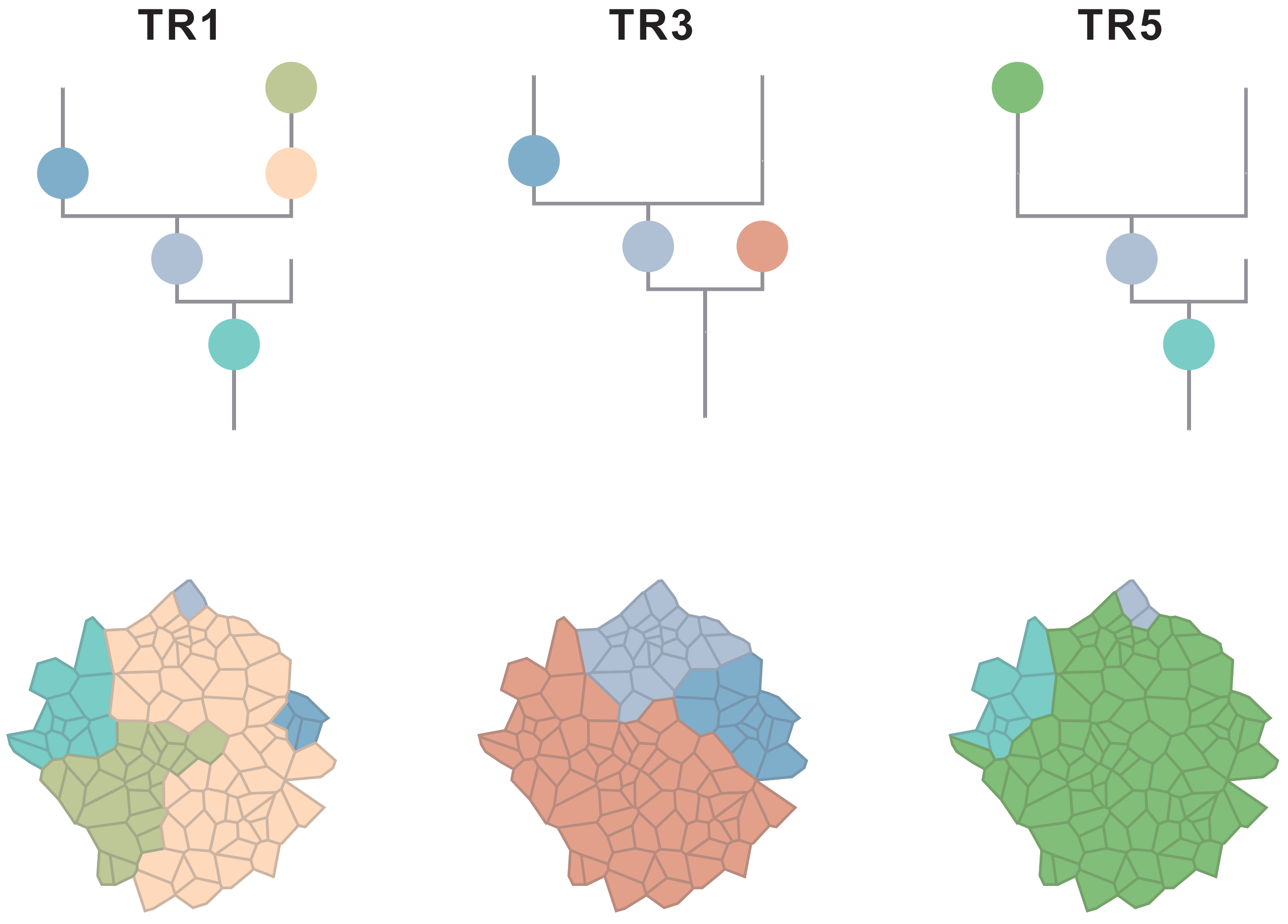

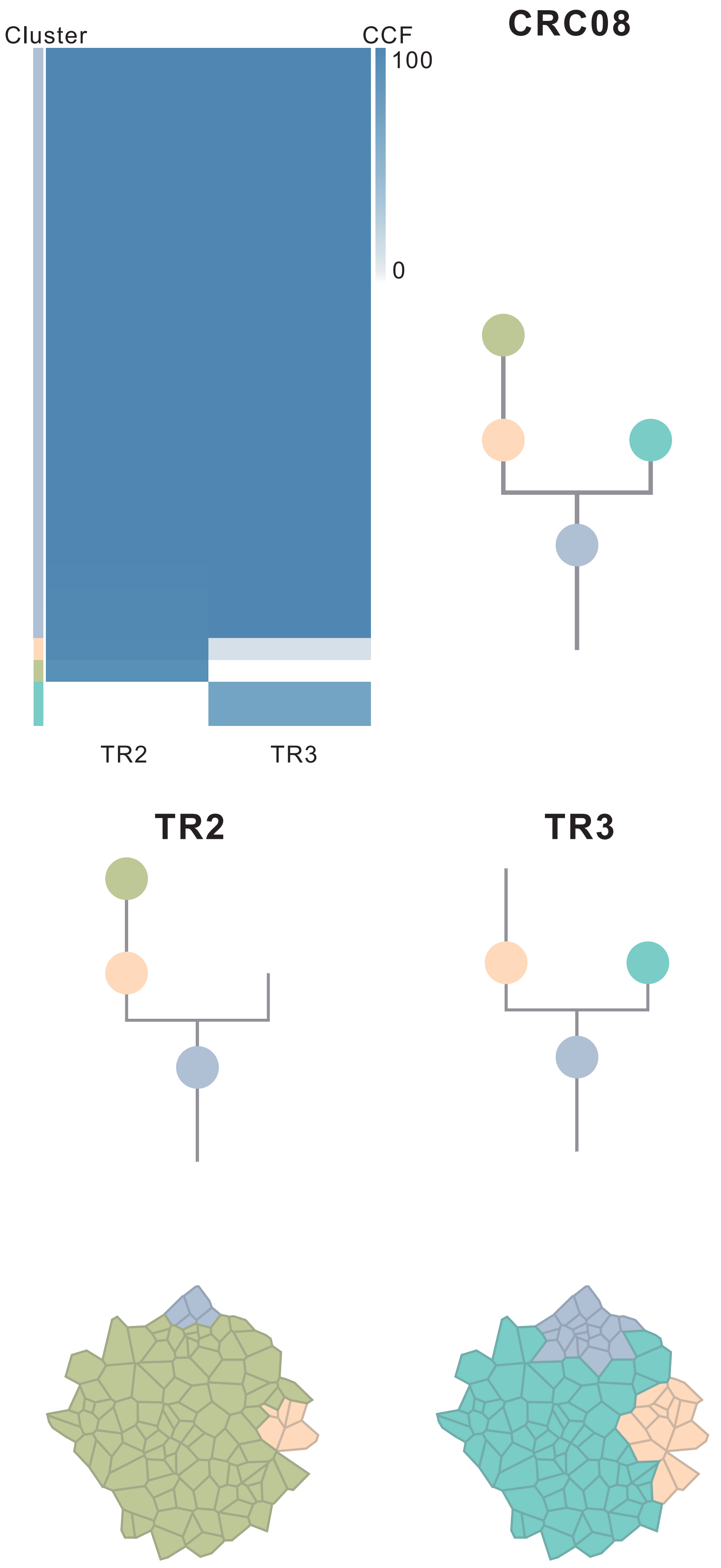

| Gene | Cytoband | Type | Cluster |
|---|---|---|---|
| *ESR1* | 6q25.1 | SNV | 1 |
| *ARHGEF10* | 8p23.3 | SNV | 1 |
| *PREX2* | 8q13.2 | SNV | 1 |
| *CSMD3* | 8q23.3 | SNV | 1 |
| *BCL9* | 1q21.2 | Amp | 1 |
| *ARNT* | 1q21.3 | Amp | 1 |
| *SETDB1* | 1q21.3 | Amp | 1 |
| *POU5F1* | 6p21.33 | Amp | 1 |
| *TRIM27* | 6p22.1 | Amp | 1 |
| *HIST1H3B* | 6p22.2 | Amp | 1 |
| *DEK* | 6p22.3 | Amp | 1 |
| *IRF4* | 6p25.3 | Amp | 1 |
| *CUX1* | 7q22.1 | Amp | 1 |
| *FLT3* | 13q12.2 | Amp | 1 |
| *FOXO1* | 13q14.11 | Amp | 1 |
| *CYSLTR2* | 13q14.2 | Amp | 1 |
| *SSX2* | Xp11.22 | Amp | 1 |
| *ARAF* | Xp11.23 | Amp | 1 |
| *GATA1* | Xp11.23 | Amp | 1 |
| *SSX1* | Xp11.23 | Amp | 1 |
| *SSX4* | Xp11.23 | Amp | 1 |
| *TFE3* | Xp11.23 | Amp | 1 |
| *WAS* | Xp11.23 | Amp | 1 |
| *KDM6A* | Xp11.3 | Amp | 1 |
| *BTK* | Xq22.1 | Amp | 1 |
| *IRS4* | Xq22.3 | Amp | 1 |
| *BCORL1* | Xq26.1 | Amp | 1 |
| *ELF4* | Xq26.1 | Amp | 1 |
| *GPC3* | Xq26.2 | Amp | 1 |
| *CASP9* | 1p36.21 | Del | 1 |
| *PRDM2* | 1p36.21 | Del | 1 |
| *SPEN* | 1p36.21 | Del | 1 |
| *CAMTA1* | 1p36.31 | Del | 1 |
| *RPL22* | 1p36.31 | Del | 1 |
| *TNFRSF14* | 1p36.32 | Del | 1 |
| *DAXX* | 6p21.32 | Del | 1 |
| *NDRG1* | 8q24.22 | Del | 1 |
| *RECQL4* | 8q24.3 | Del | 1 |
| *FEN1* | 11q12.2 | Del | 1 |
| *SDHAF2* | 11q12.2 | Del | 1 |
| *MEN1* | 11q13.1 | Del | 1 |
| *FLCN* | 17p11.2 | Del | 1 |
| *NCOR1* | 17p11.2 | Del | 1 |
| *MAP2K4* | 17p12 | Del | 1 |
| *PER1* | 17p13.1 | Del | 1 |
| *TP53* | 17p13.1 | Del | 1 |
| *YWHAE* | 17p13.3 | Del | 1 |
| *ELL* | 19p13.11 | Del | 1 |
| *DNM2* | 19p13.2 | Del | 1 |
| *KEAP1* | 19p13.2 | Del | 1 |
| *SMARCA4* | 19p13.2 | Del | 1 |
| *STK11* | 19p13.3 | Del | 1 |
| *TCF3* | 19p13.3 | Del | 1 |
| *CLTCL1* | 22q11.21 | Del | 1 |
| *LZTR1* | 22q11.21 | Del | 1 |
| *SMARCB1* | 22q11.23 | Del | 1 |
| *CHEK2* | 22q12.1 | Del | 1 |
| *ZNRF3* | 22q12.1 | Del | 1 |
| *NF2* | 22q12.2 | Del | 1 |
| *MYH9* | 22q12.3 | Del | 1 |
| *APOBEC3B* | 22q13.1 | Del | 1 |
| *EP300* | 22q13.2 | Del | 1 |
| *MKL1* | 22q13.2 | Del | 1 |
| *MUC4* | 3q29 | Amp | 2 |
| *NOTCH1* | 9q34.3 | Del | 2 |
| *TBX3* | 12q24.21 | Del | 2 |
| *HNF1A* | 12q24.31 | Del | 2 |
| *NCOR2* | 12q24.31 | Del | 2 |
| *SETD1B* | 12q24.31 | Del | 2 |
| *POLE* | 12q24.33 | Del | 2 |
| *TP53* | 17p13.1 | SNV | NA |

### CRC10

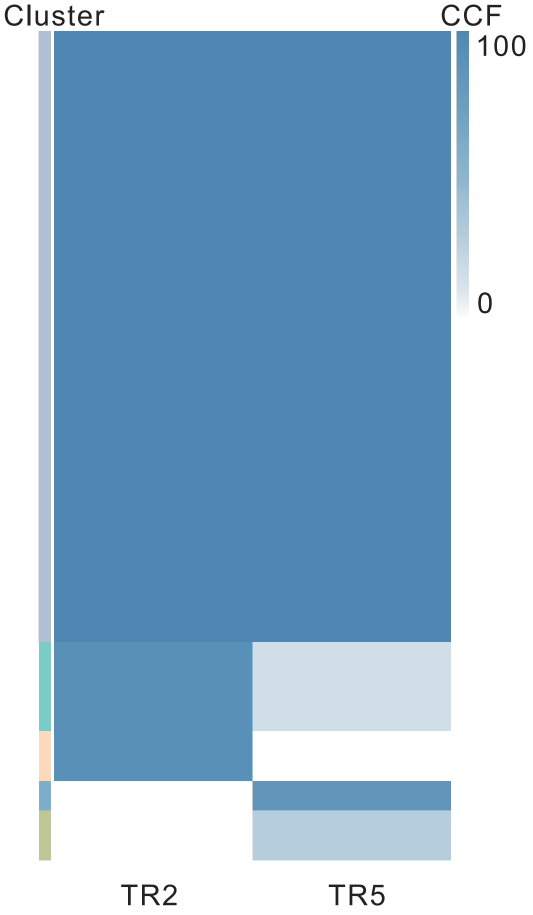

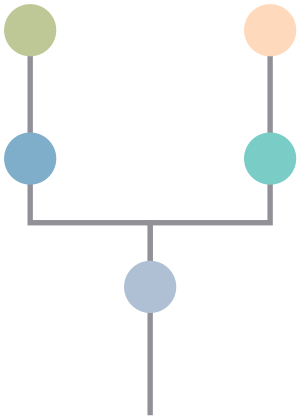

| Gene | Cytoband | Type | Cluster |
|---|---|---|---|
| *KRAS* | 12p12.1 | SNV | 1 |
| *APC* | 5q22.2 | InDel | 1 |
| *FLT3* | 13q12.2 | Amp | 1 |
| *FOXO1* | 13q14.11 | Amp | 1 |
| *CYSLTR2* | 13q14.2 | Amp | 1 |
| *CAMTA1* | 1p36.31 | Del | 1 |
| *RPL22* | 1p36.31 | Del | 1 |
| *TNFRSF14* | 1p36.32 | Del | 1 |
| *FLCN* | 17p11.2 | Del | 1 |
| *NCOR1* | 17p11.2 | Del | 1 |
| *MAP2K4* | 17p12 | Del | 1 |
| *PER1* | 17p13.1 | Del | 1 |
| *TP53* | 17p13.1 | Del | 1 |
| *YWHAE* | 17p13.3 | Del | 1 |
| *TSC1* | 9q34.13 | Del | 2 |
| *NOTCH1* | 9q34.3 | Del | 2 |
| *AXIN1* | 16p13.3 | Del | 2 |
| *STK11* | 19p13.3 | Del | 2 |
| *TCF3* | 19p13.3 | Del | 2 |

## TR2

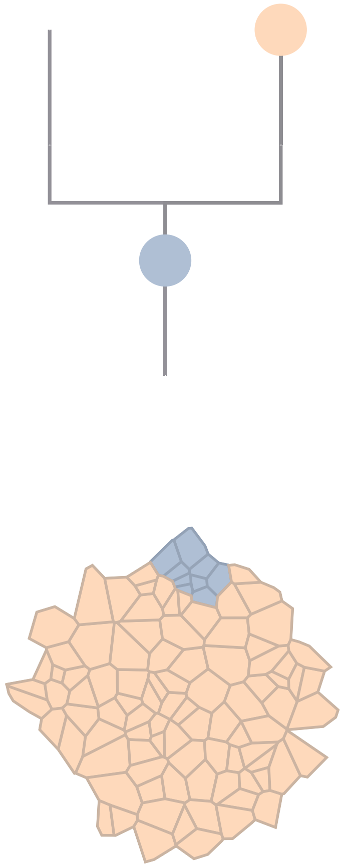

## TR5

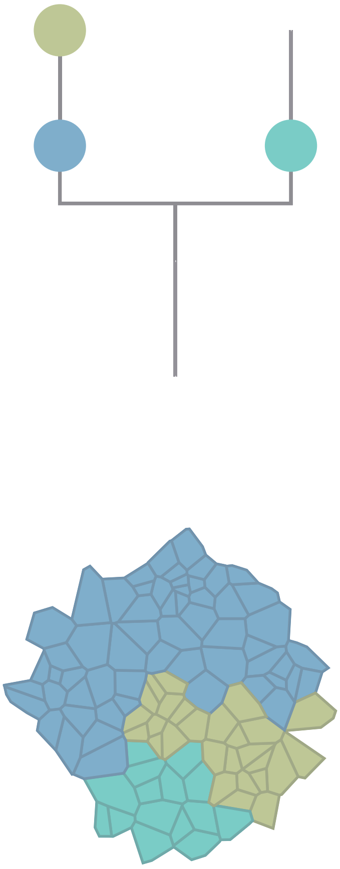

### CRC11

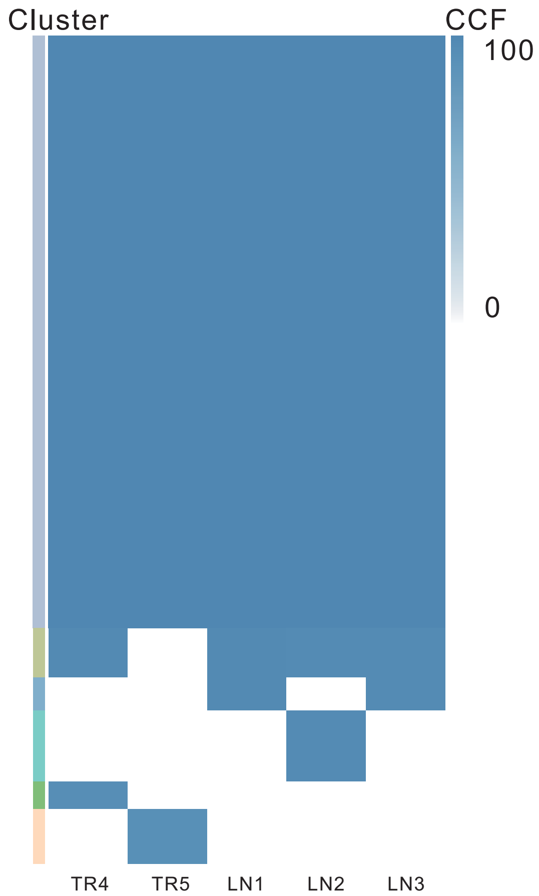

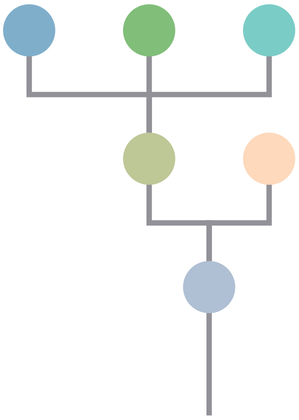

| Gene | Cytoband | Type | Cluster |
|---|---|---|---|
| * *APC* | 5q22.2 | SNV | 1 |
| *BRAF* | 7q34 | SNV | 1 |
| *ZFHX3* | 16q22.2 | SNV | 1 |
| * *SMAD4* | 18q21.2 | SNV | 1 |
| * *SMAD4* | 18q21.2 | SNV | 1 |
| * *APC* | 5q22.2 | InDel | 1 |
| *PAX5* | 9p13.2 | Amp | 1 |
| *PSIP1* | 9p22.3 | Amp | 1 |
| *JAK2* | 9p24.1 | Amp | 1 |
| *PDCD1LG2* | 9p24.1 | Amp | 1 |
| *GNAQ* | 9q21.2 | Amp | 1 |
| *SYK* | 9q22.2 | Amp | 1 |
| *NR4A3* | 9q31.1 | Amp | 1 |
| *KLF4* | 9q31.2 | Amp | 1 |
| *TAL2* | 9q31.2 | Amp | 1 |
| *TNC* | 9q33.1 | Amp | 1 |
| *SET* | 9q34.11 | Amp | 1 |
| *ABL1* | 9q34.12 | Amp | 1 |
| *BRD3* | 9q34.2 | Amp | 1 |
| *NOTCH1* | 9q34.3 | Amp | 1 |
| *FLT3* | 13q12.2 | Amp | 2 |
| *FOXO1* | 13q14.11 | Amp | 2 |
| *CYSLTR2* | 13q14.2 | Amp | 2 |
| *SDHA* | 5p15.33 | Del | 3 |
| *MUC4* | 3q29 | Amp | NA |
| *PLAG1* | 8q12.1 | Amp | NA |
| *PREX2* | 8q13.2 | Amp | NA |
| *NCOA2* | 8q13.3 | Amp | NA |
| *HEY1* | 8q21.13 | Amp | NA |
| *RUNX1T1* | 8q21.3 | Amp | NA |
| *CDH17* | 8q22.1 | Amp | NA |
| *PABPC1* | 8q22.3 | Amp | NA |
| *UBR5* | 8q22.3 | Amp | NA |
| *RAD21* | 8q24.11 | Amp | NA |
| *AXIN1* | 16p13.3 | Del | NA |

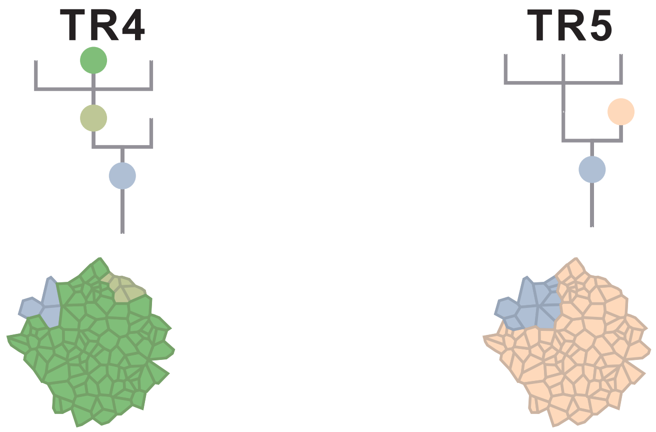

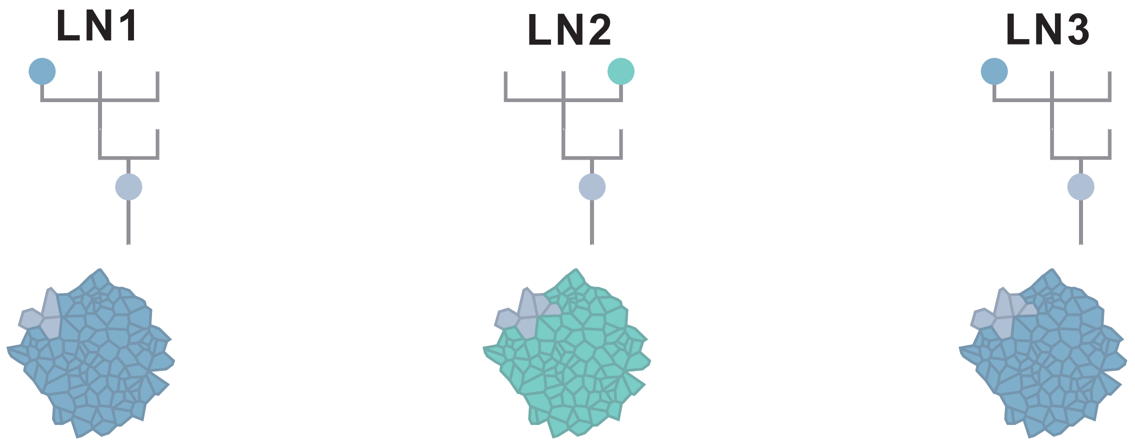

### CRC12

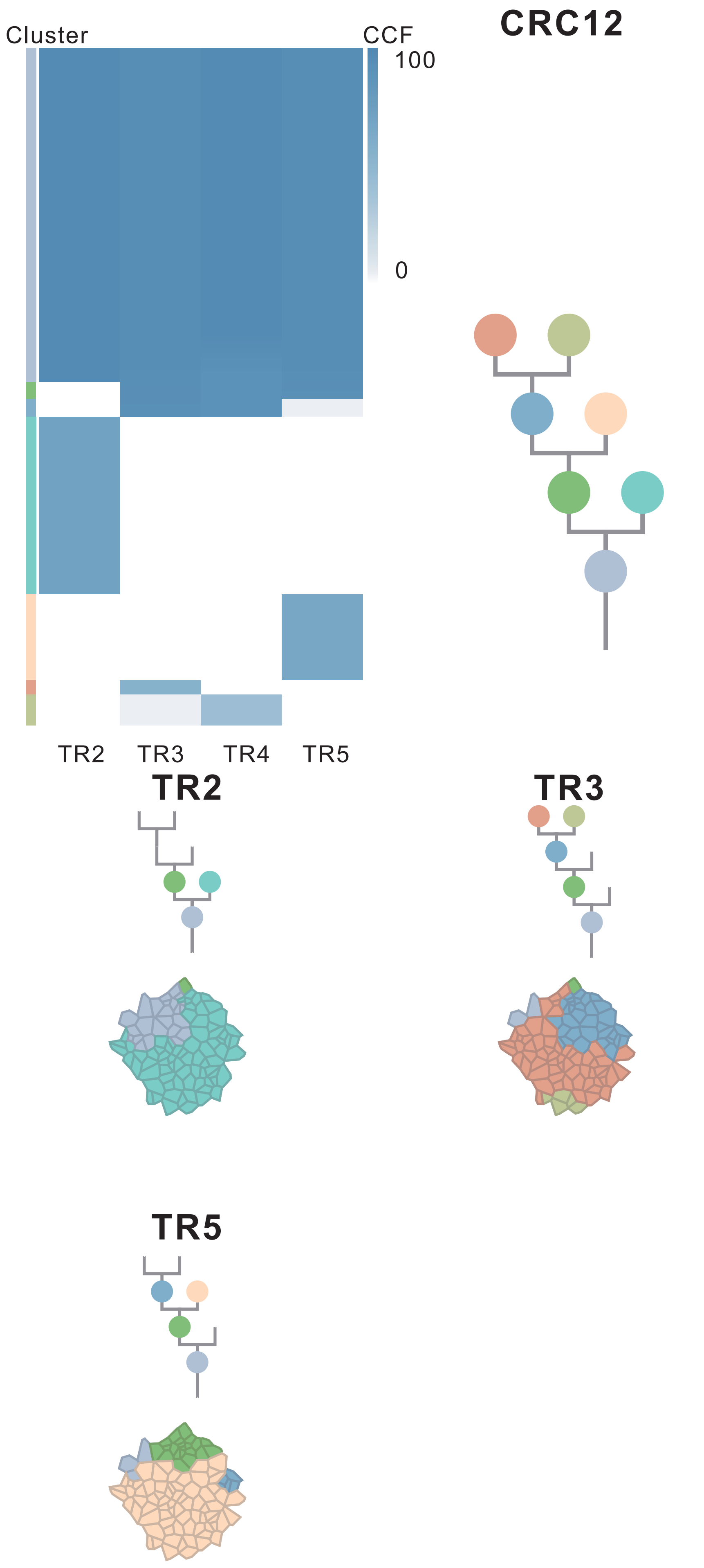

| Gene | Cytoband | Type | Cluster |
| --- | --- | --- | --- |
| *IDH1* | 2q34 | SNV | 1 |
| *BRAF* | 7q34 | SNV | 1 |
| *ARID2* | 12q12 | SNV | 1 |
| *RNF43* | 17q22 | SNV | 1 |
| *SMAD4* | 18q21.2 | SNV | 1 |
| *TP53* | 17p13.1 | InDel | 1 |
| *CREB3L2* | 7q33 | Amp | 1 |
| *TRIM24* | 7q33 | Amp | 1 |
| *BRAF* | 7q34 | Amp | 1 |
| *EZH2* | 7q36.1 | Amp | 1 |
| *RUNX1T1* | 8q21.3 | Amp | 1 |
| *CDH17* | 8q22.1 | Amp | 1 |
| *PABPC1* | 8q22.3 | Amp | 1 |
| *UBR5* | 8q22.3 | Amp | 1 |
| *RAD21* | 8q24.11 | Amp | 1 |
| *MYC* | 8q24.21 | Amp | 1 |
| *RECQL4* | 8q24.3 | Amp | 1 |
| *FLT3* | 13q12.2 | Amp | 1 |
| *FOXO1* | 13q14.11 | Amp | 1 |
| *CYSLTR2* | 13q14.2 | Amp | 1 |
| *ELK4* | 1q32.1 | Amp | 2 |
| *MDM4* | 1q32.1 | Amp | 2 |
| *ROBO2* | 3p12.3 | SNV | 3 |
| *FAT4* | 4q28.1 | SNV | 4 |
| *PRDM16* | 1p36.32 | Amp | NA |
| *ARNT* | 1q21.3 | Amp | NA |
| *SETDB1* | 1q21.3 | Amp | NA |
| *FCRL4* | 1q23.1 | Amp | NA |
| *NTRK1* | 1q23.1 | Amp | NA |
| *DDR2* | 1q23.3 | Amp | NA |
| *FCGR2B* | 1q23.3 | Amp | NA |
| *PBX1* | 1q23.3 | Amp | NA |

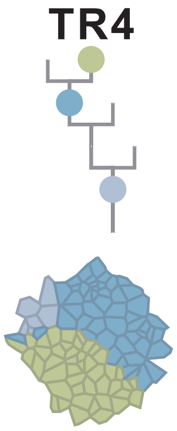

### CRC14

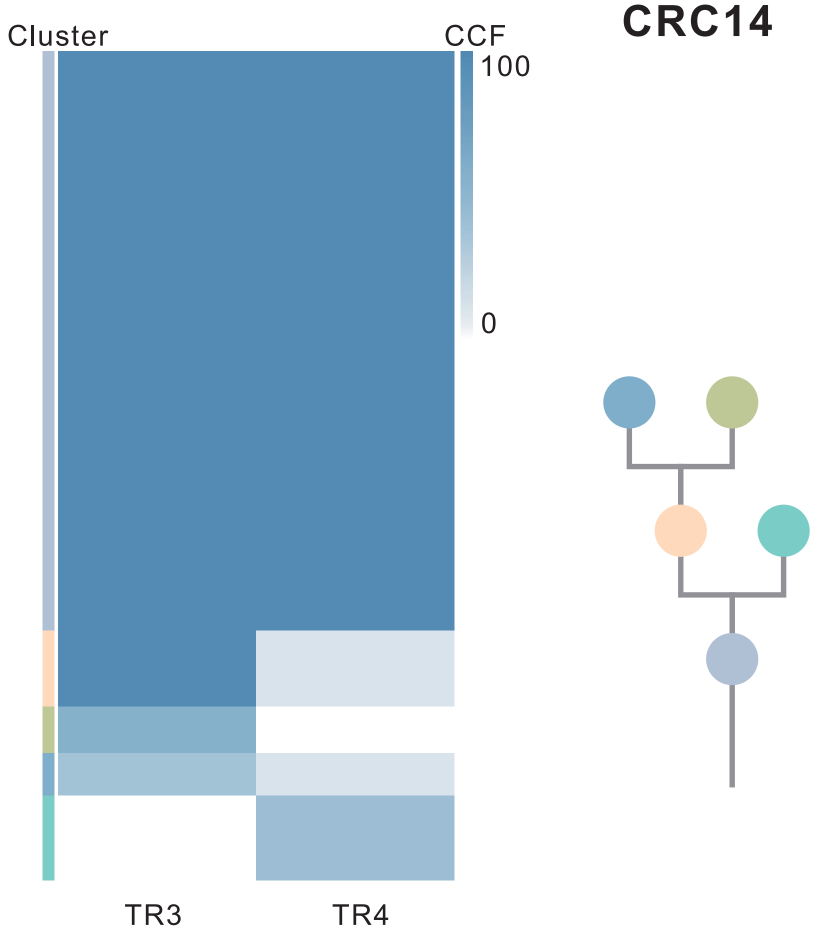

| Gene | Cytoband | Type | Cluster |
|---|---|---|---|
| *SDHB* | 1p36.13 | SNV | 1 |
| *TP63* | 3q28 | SNV | 1 |
| *APC* | 5q22.2 | SNV | 1 |
| *ESR1* | 6q25.1 | SNV | 1 |
| *DICER1* | 14q32.13 | SNV | 1 |
| *TP53* | 17p13.1 | SNV | 1 |
| *IKBKB* | 8p11.21 | Amp | 1 |
| *KAT6A* | 8p11.21 | Amp | 1 |
| *FGFR1* | 8p11.23 | Amp | 1 |
| *SRC* | 20q11.23 | Amp | 1 |
| *MAFB* | 20q12 | Amp | 1 |
| *PLCG1* | 20q12 | Amp | 1 |
| *NFATC2* | 20q13.2 | Amp | 1 |
| *SALL4* | 20q13.2 | Amp | 1 |
| *GNAS* | 20q13.32 | Amp | 1 |
| *PTK6* | 20q13.33 | Amp | 1 |

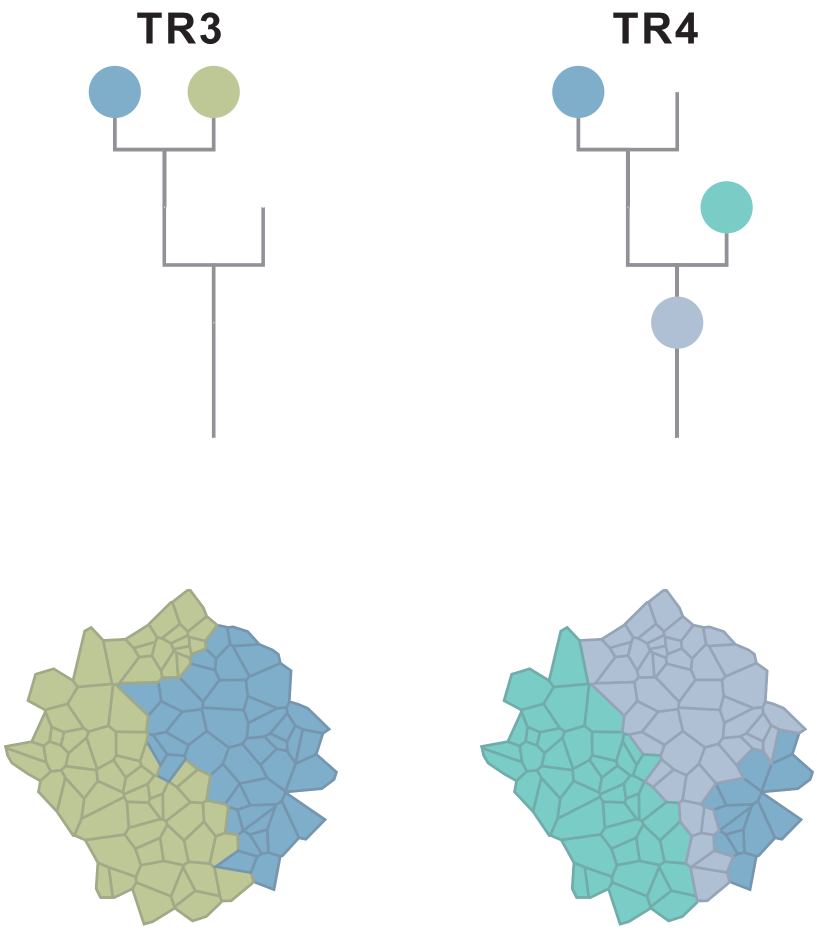

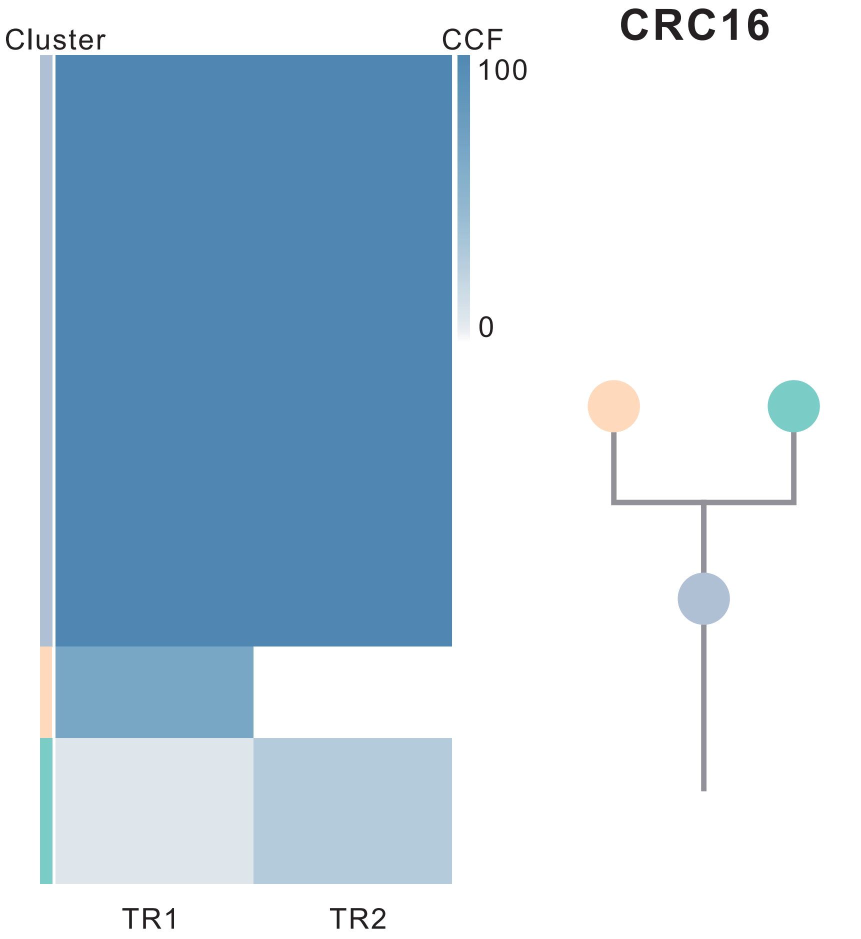

| Gene | Cytoband | Type | Cluster |
|---|---|---|---|
| *MITF* | 3p13 | SNV | 1 |
| *CTNNB1* | 3p22.1 | SNV | 1 |
| *PIK3CA* | 3q26.32 | SNV | 1 |
| *APC* | 5q22.2 | SNV | 1 |
| *KRAS* | 12p12.1 | SNV | 1 |
| *RSPO2* | 8q23.1 | SNV | 2 |

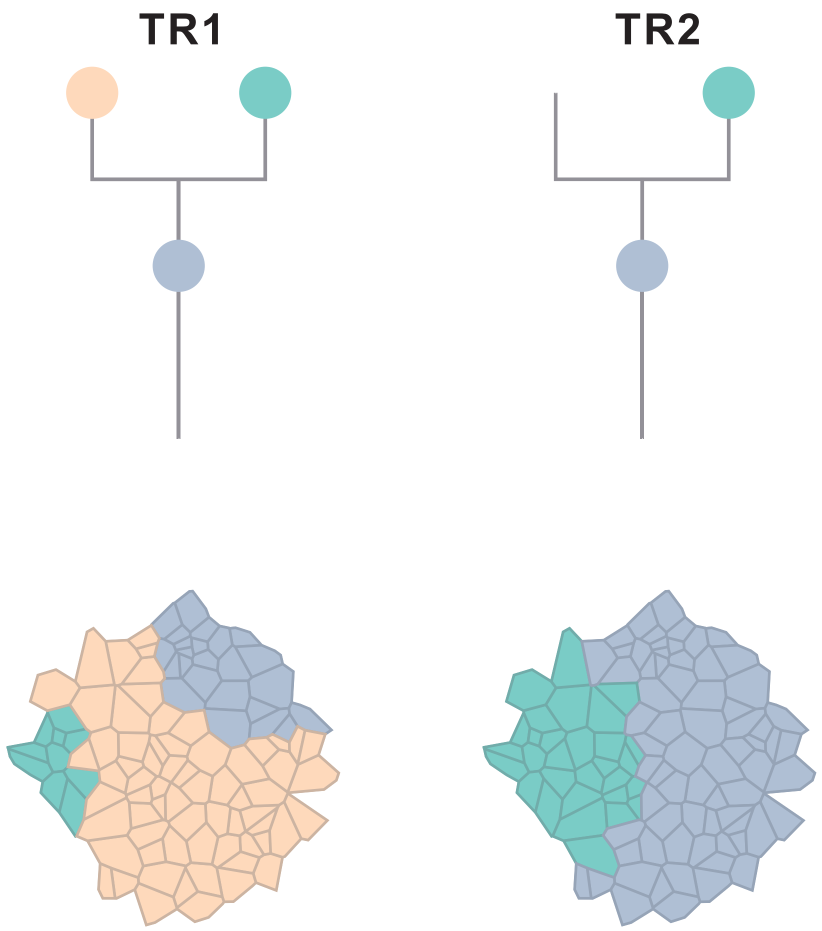

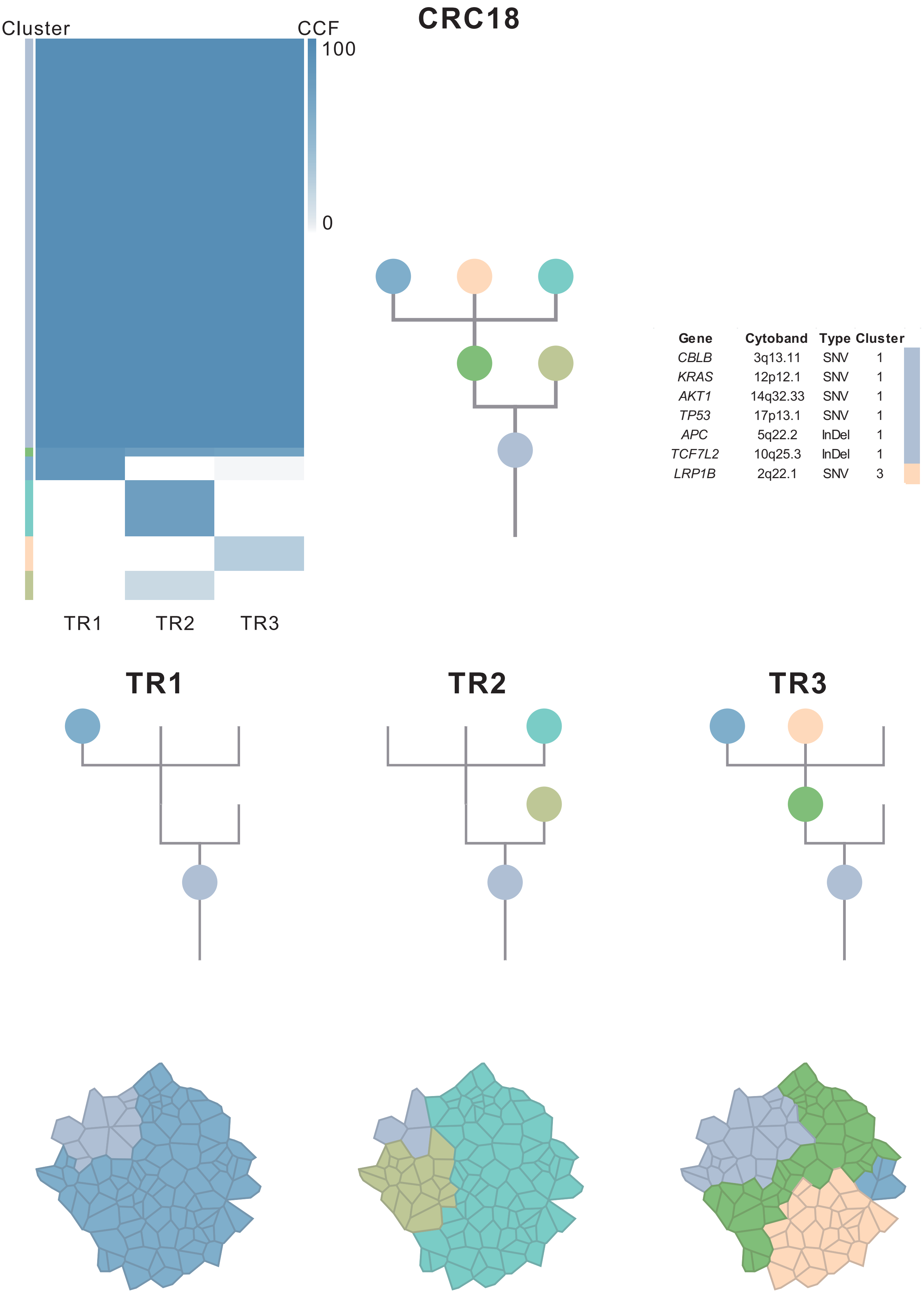
CRC18
Cluster
CCF
100
0
TR1
TR2
TR3
Gene
Cytoband
Type
Cluster
CBLB 3q13.11 SNV 1
KRAS 12p12.1 SNV 1
AKT1 14q32.33 SNV 1
TP53 17p13.1 SNV 1
APC 5q22.2 InDel 1
TCF7L2 10q25.3 InDel 1
LRP1B 2q22.1 SNV 3
TR1
TR2
TR3

### CRC19

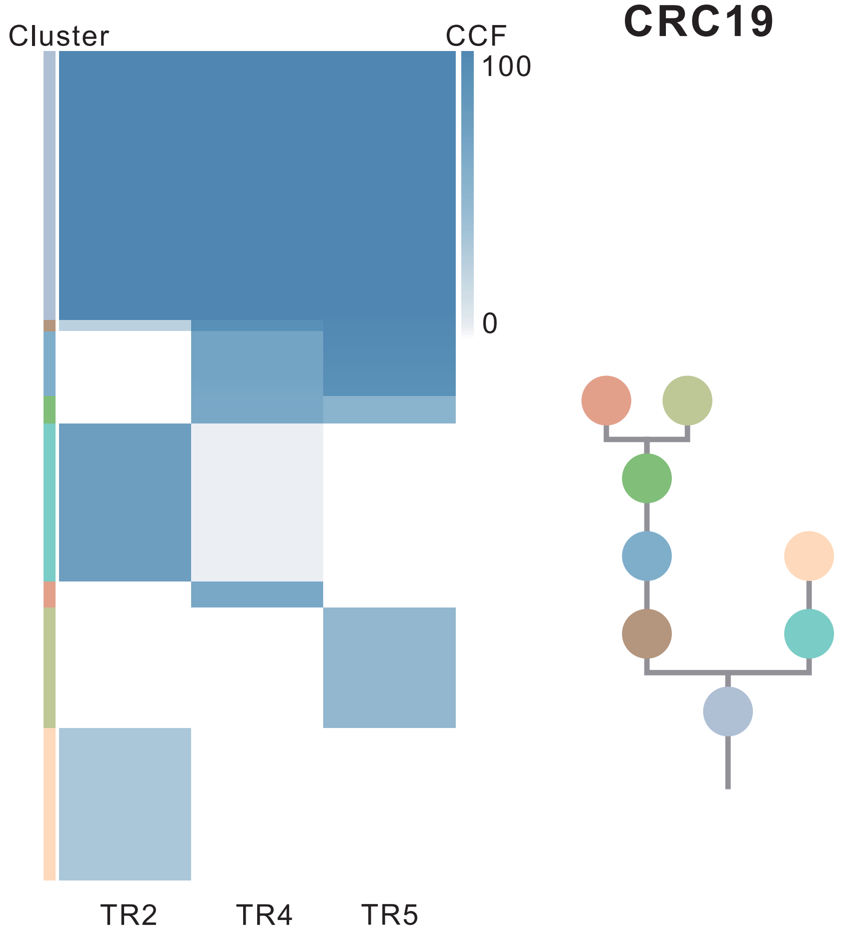

| Gene | Cytoband | Type | Cluster |
|---|---|---|---|
| *APC* | 5q22.2 | SNV | 1 |
| *KRAS* | 12p12.1 | SNV | 1 |
| *ARID2* | 12q12 | SNV | 1 |
| *FLCN* | 17p11.2 | SNV | 1 |
| *TP53* | 17p13.1 | SNV | 1 |
| *SRC* | 20q11.23 | Amp | 1 |
| *MAFB* | 20q12 | Amp | 1 |
| *PLCG1* | 20q12 | Amp | 1 |
| *NFATC2* | 20q13.2 | Amp | 1 |
| *SALL4* | 20q13.2 | Amp | 1 |
| *GNAS* | 20q13.32 | Amp | 1 |
| *PTK6* | 20q13.33 | Amp | 1 |
| *PER1* | 17p13.1 | SNV | 3 |
| *AMER1* | Xq11.2 | SNV | NA |

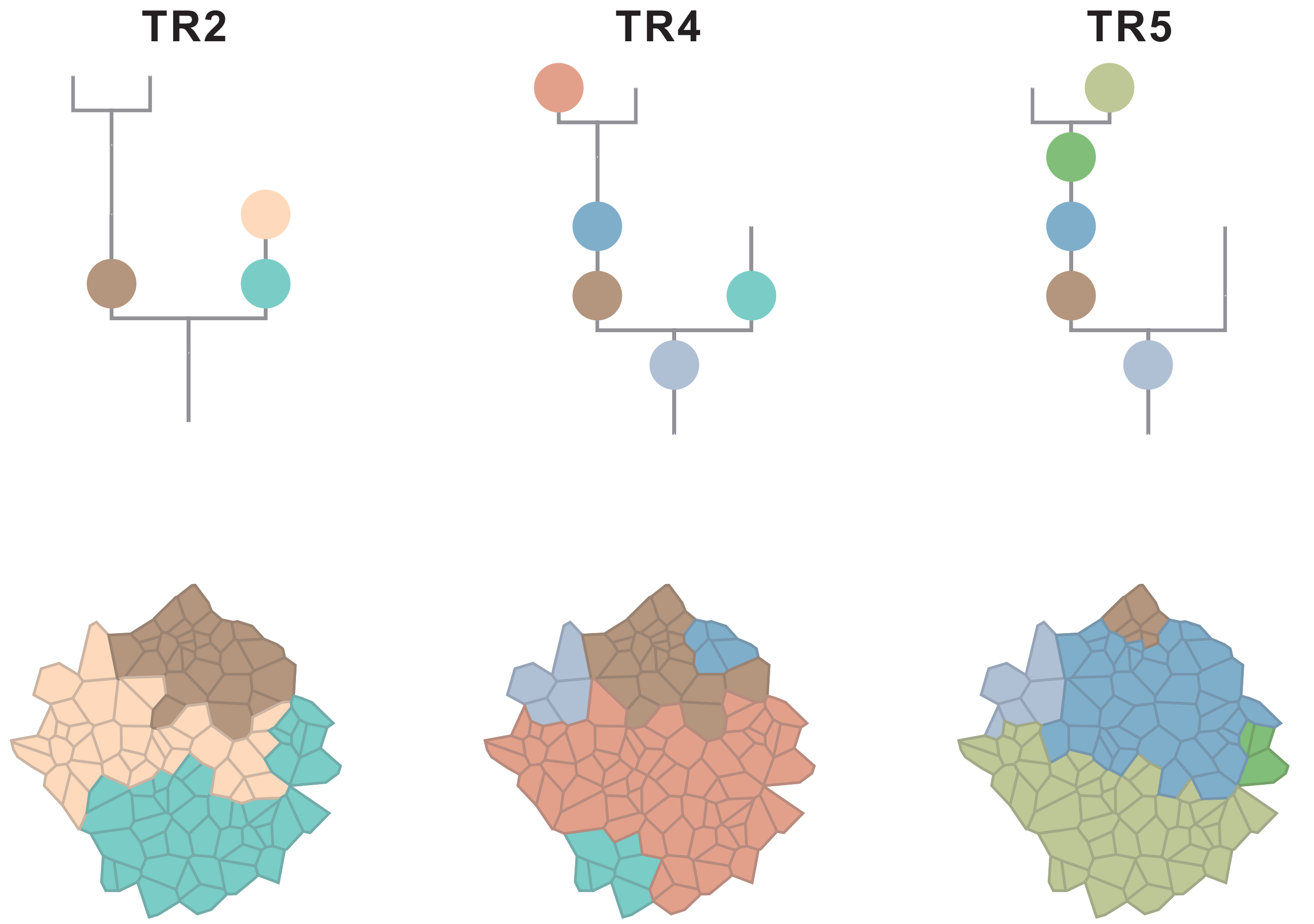

### CRC20

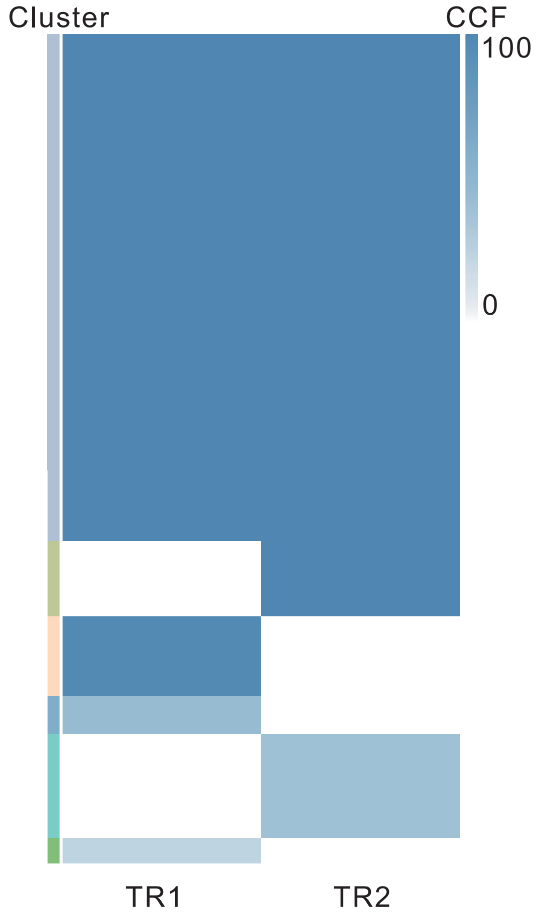

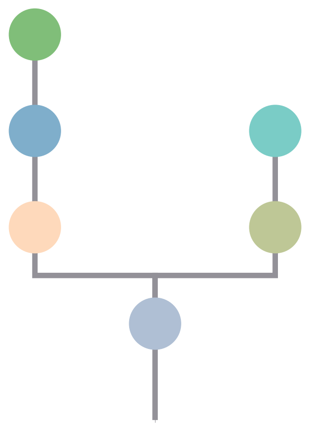

| Gene | Cytoband | Type | Cluster |
|---|---|---|---|
| *FBXW7* | 4q31.3 | SNV | 1 |
| *APC* | 5q22.2 | SNV | 1 |
| *TP53* | 17p13.1 | SNV | 1 |
| *SRC* | 20q11.23 | Amp | 1 |
| *MAFB* | 20q12 | Amp | 1 |
| *PLCG1* | 20q12 | Amp | 1 |
| *NFATC2* | 20q13.2 | Amp | 1 |
| *SALL4* | 20q13.2 | Amp | 1 |
| *GNAS* | 20q13.32 | Amp | 1 |
| *PTK6* | 20q13.33 | Amp | 1 |

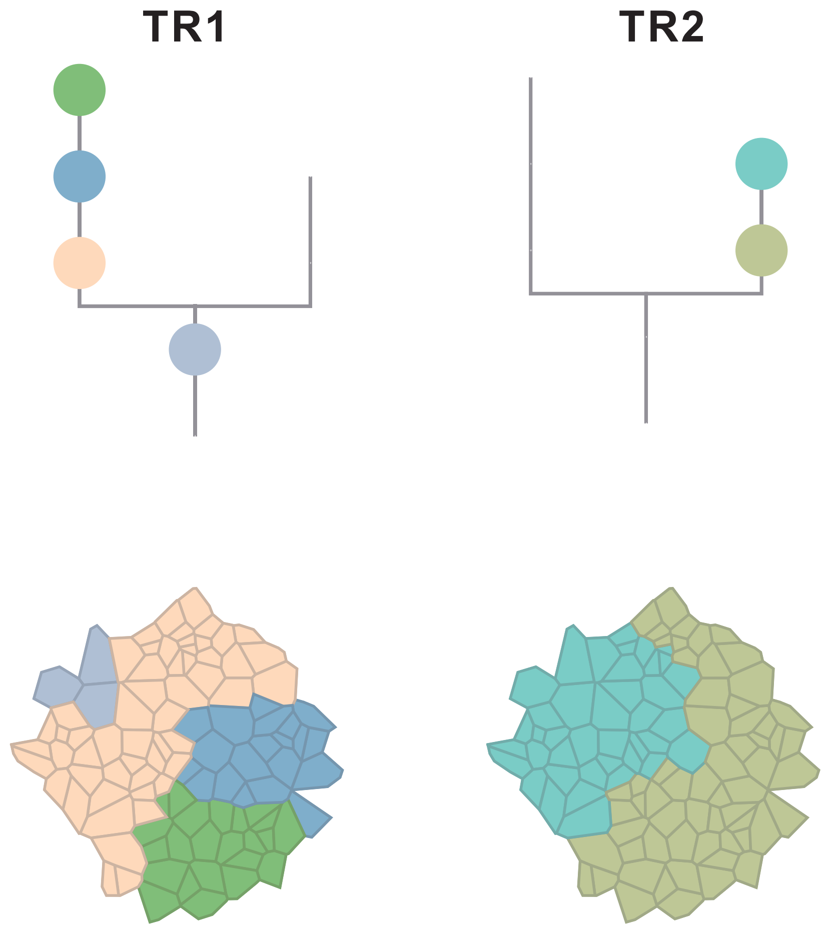

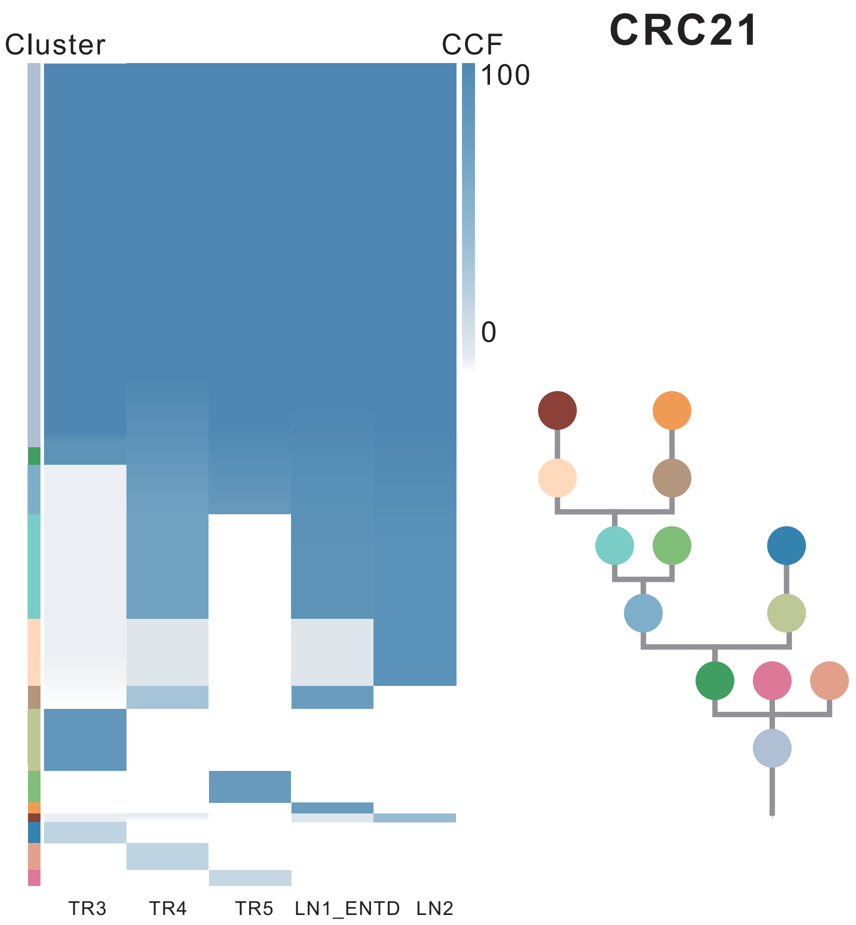

| Gene | Cytoband | Type | Cluster |
|---|---|---|---|
| * *ASXL2* | 2p23.3 | SNV | 1 |
| *BRAF* | 7q34 | SNV | 1 |
| * *ASXL2* | 2p23.3 | InDel | 1 |
| *TP53* | 17p13.1 | InDel | 1 |
| *RNF43* | 17q22 | InDel | 1 |
| *NCOA2* | 8q13.3 | Amp | 1 |
| *HEY1* | 8q21.13 | Amp | 1 |
| *RUNX1T1* | 8q21.3 | Amp | 1 |
| *CDH17* | 8q22.1 | Amp | 1 |
| *PABPC1* | 8q22.3 | Amp | 1 |
| *UBR5* | 8q22.3 | Amp | 1 |
| *RAD21* | 8q24.11 | Amp | 1 |
| *MYC* | 8q24.21 | Amp | 1 |
| *RECQL4* | 8q24.3 | Amp | 1 |
| *CCND2* | 12p13.32 | Amp | 2 |
| *KDM5A* | 12p13.33 | Amp | 2 |
| *ERBB4* | 2q34 | SNV | 3 |
| *TSC1* | 9q34.13 | SNV | 3 |

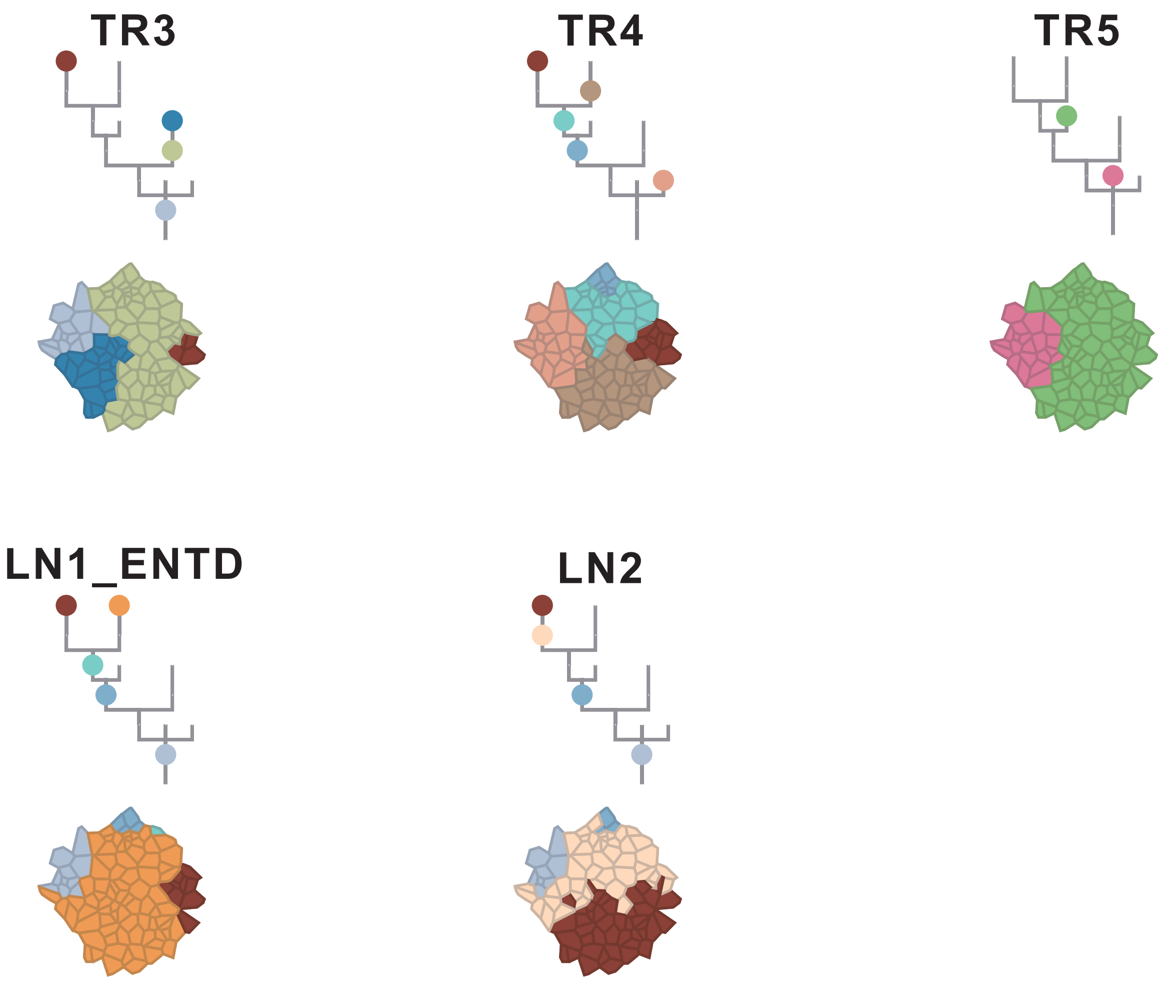

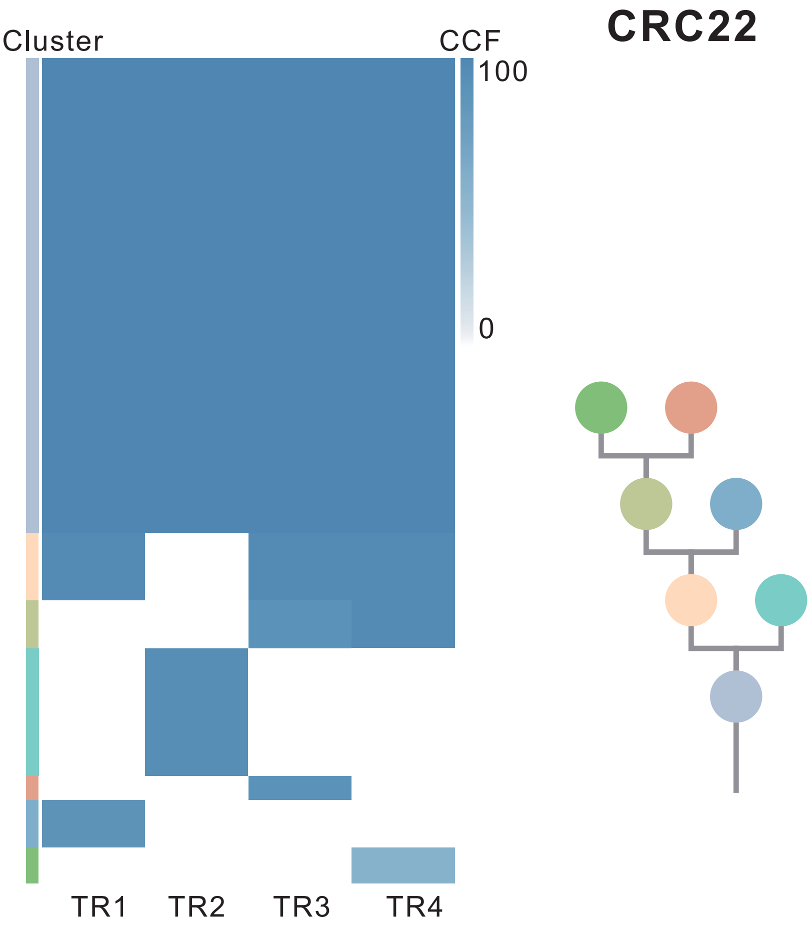

| Gene | Cytoband | Type | Cluster |
|---|---|---|---|
| *APC* | 5q22.2 | SNV | 1 |
| *KRAS* | 12p12.1 | SNV | 1 |
| *TP53* | 17p13.1 | SNV | 1 |
| *FLT3* | 13q12.2 | Amp | 1 |
| *FOXO1* | 13q14.11 | Amp | 1 |
| *CYSLTR2* | 13q14.2 | Amp | 1 |
| *SRC* | 20q11.23 | Amp | 1 |
| *MAFB* | 20q12 | Amp | 1 |
| *PLCG1* | 20q12 | Amp | 1 |
| *NFATC2* | 20q13.2 | Amp | 1 |
| *SALL4* | 20q13.2 | Amp | 1 |
| *GNAS* | 20q13.32 | Amp | 1 |
| *PTK6* | 20q13.33 | Amp | 1 |
| *KMT2D* | 12q13.12 | SNV | 2 |
| *AR* | Xq12 | SNV | NA |

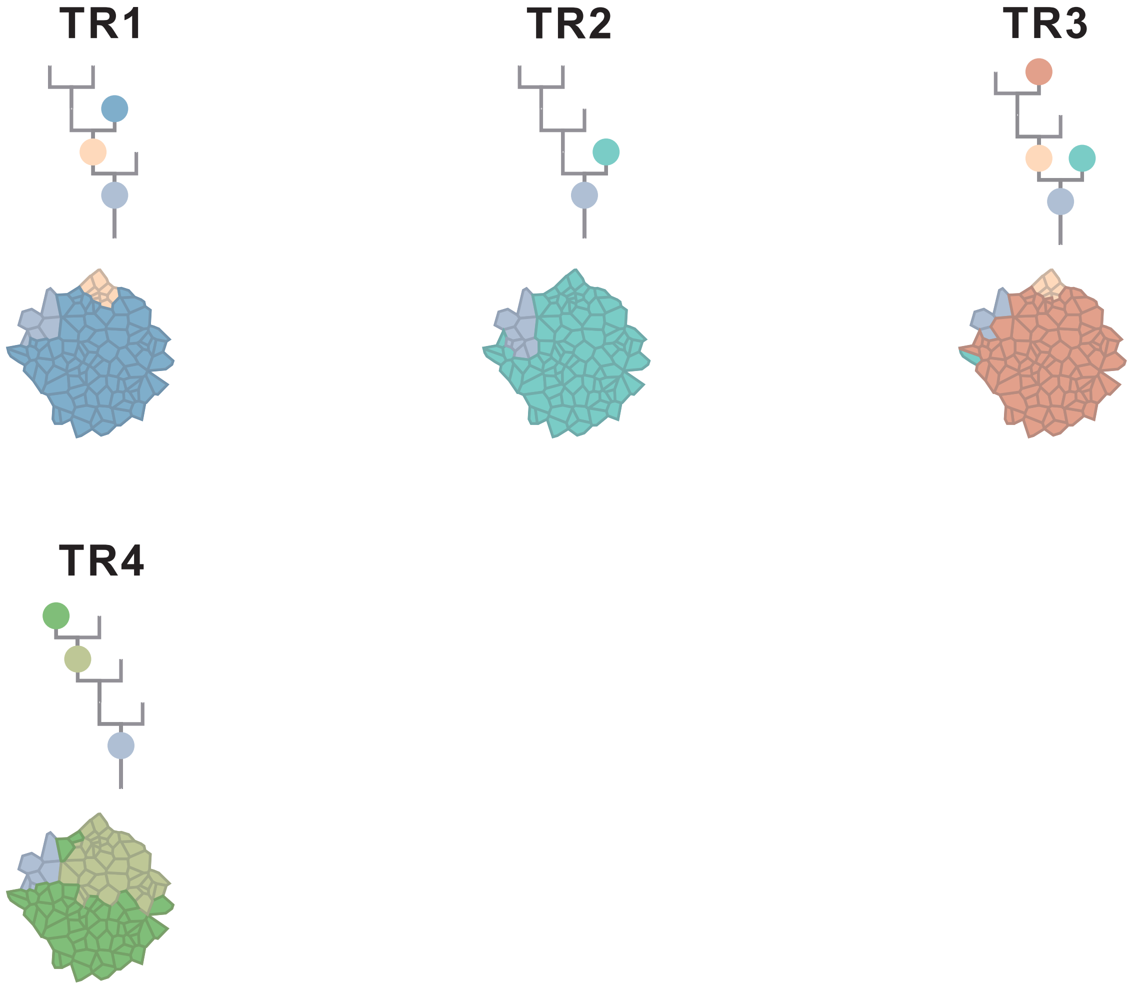

### CRC23

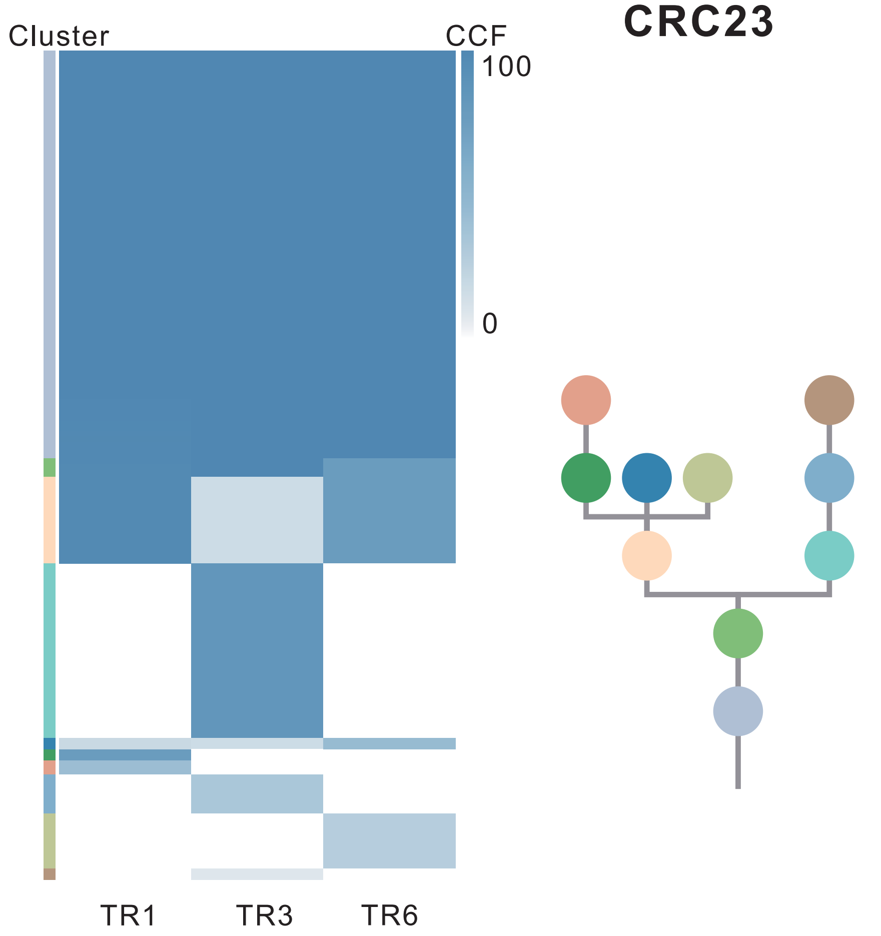

| Gene | Cytoband | Type | Cluster |
|---|---|---|---|
| *ROBO2* | 3p12.3 | SNV | 1 |
| *FAT4* | 4q28.1 | SNV | 1 |
| *APC* | 5q22.2 | SNV | 1 |
| *STAT5B* | 17q21.2 | SNV | 1 |
| *BCL9L* | 11q23.3 | InDel | 1 |
| *LRP1B* | 2q22.1 | SNV | 2 |
| *MSH6* | 2p16.3 | SNV | 4 |
| *BCORL1* | Xq26.1 | SNV | NA |

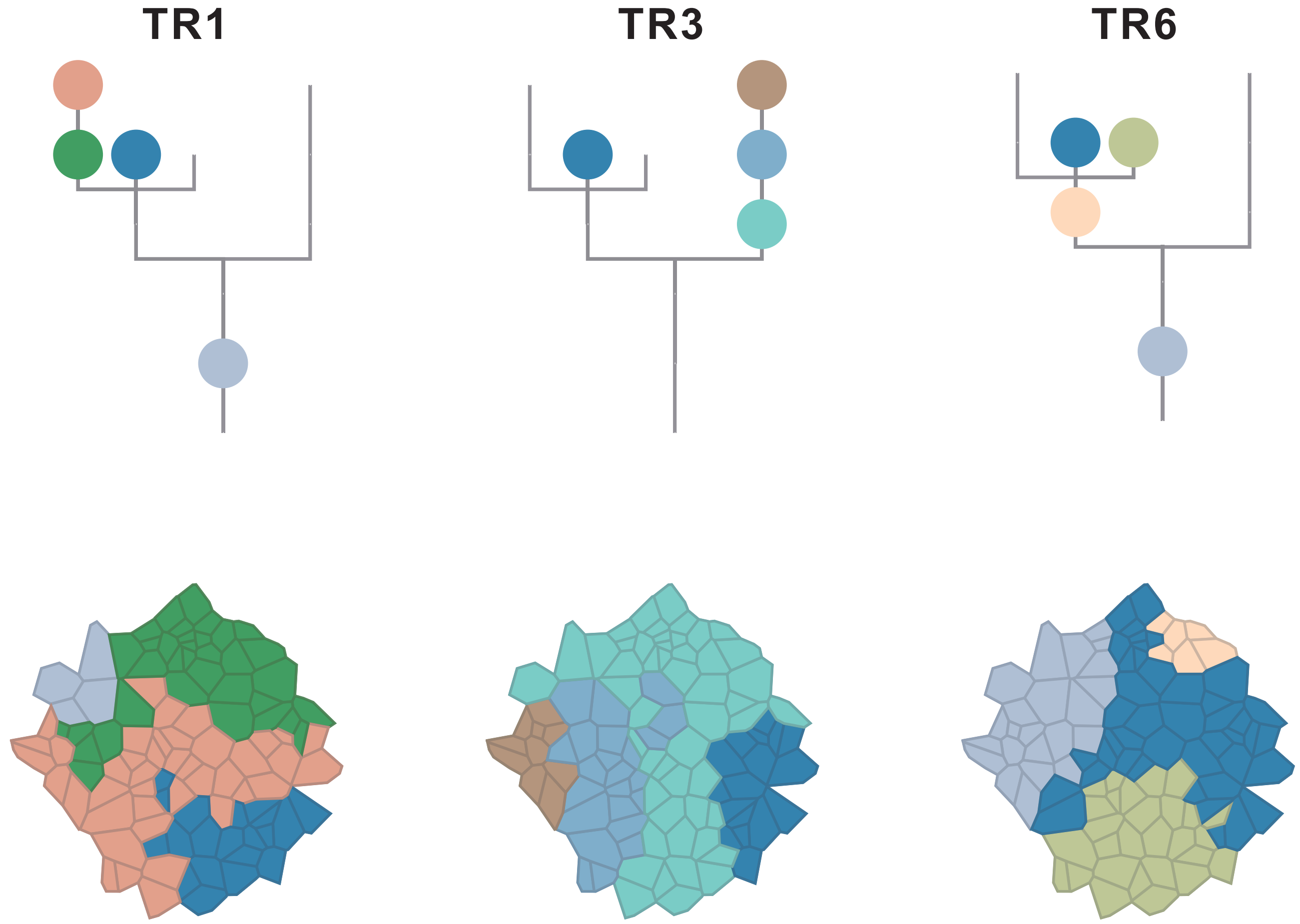

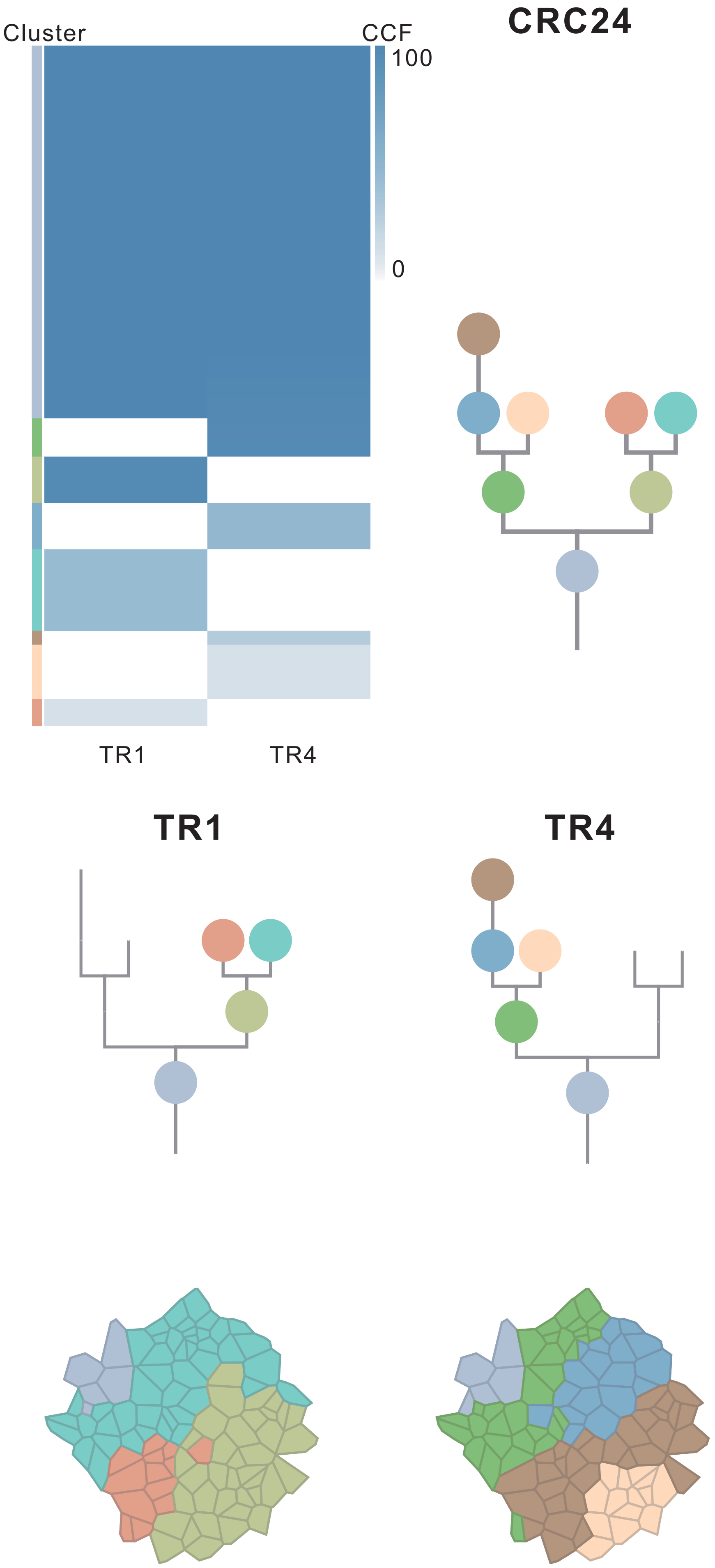

| Gene | Cytoband | Type | Cluster |
|---|---|---|---|
| *BMPR1A* | 10q23.2 | SNV | 1 |
| *KRAS* | 12p12.1 | SNV | 1 |
| *TP53* | 17p13.1 | SNV | 1 |
| *GRIN2A* | 16p13.2 | SNV | 3 |

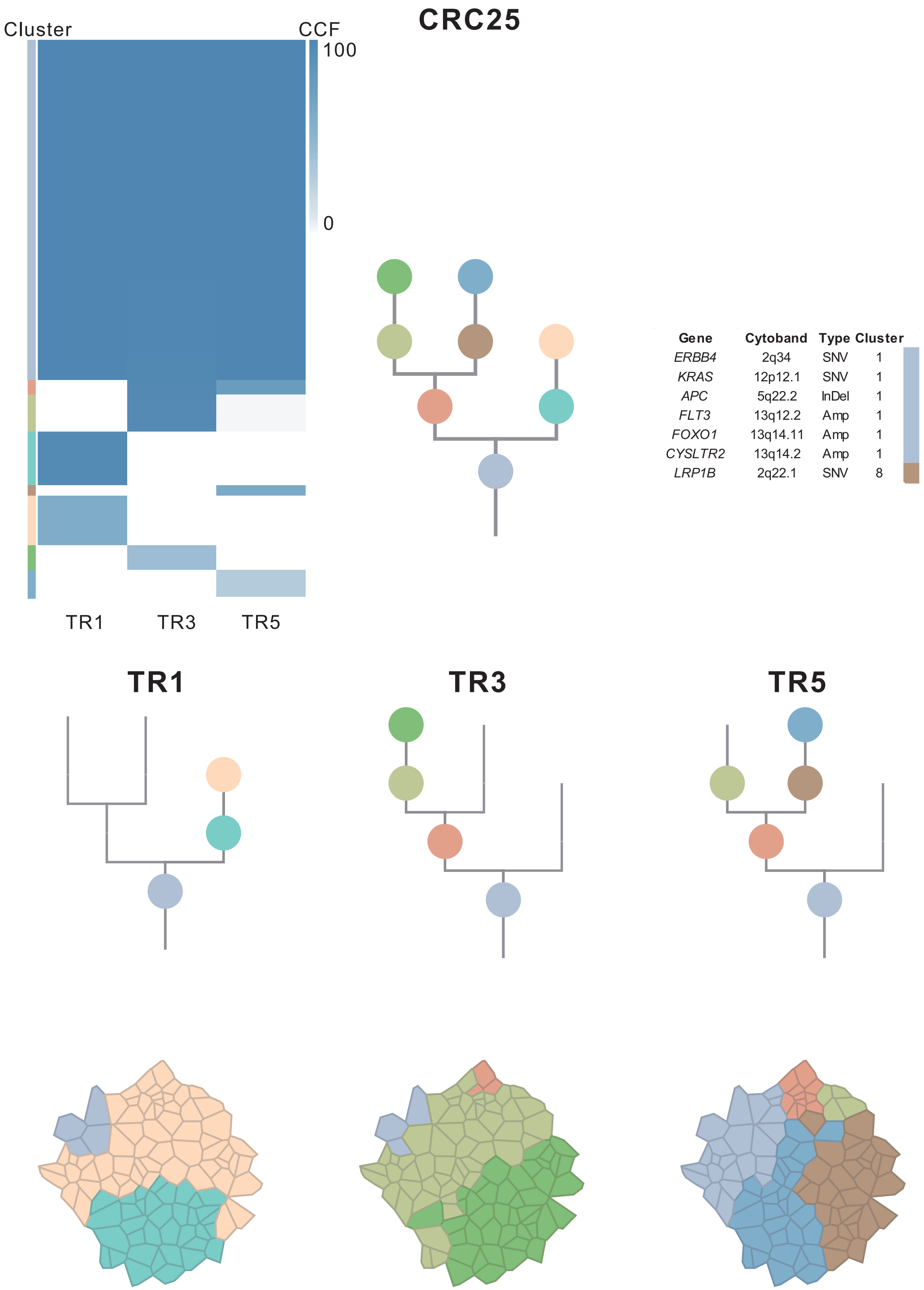

| Gene | Cytoband | Type | Cluster |
| --- | --- | --- | --- |
| *ERBB4* | 2q34 | SNV | 1 |
| *KRAS* | 12p12.1 | SNV | 1 |
| *APC* | 5q22.2 | InDel | 1 |
| *FLT3* | 13q12.2 | Amp | 1 |
| *FOXO1* | 13q14.11 | Amp | 1 |
| *CYSLTR2* | 13q14.2 | Amp | 1 |
| *LRP1B* | 2q22.1 | SNV | 8 |

### CRC26

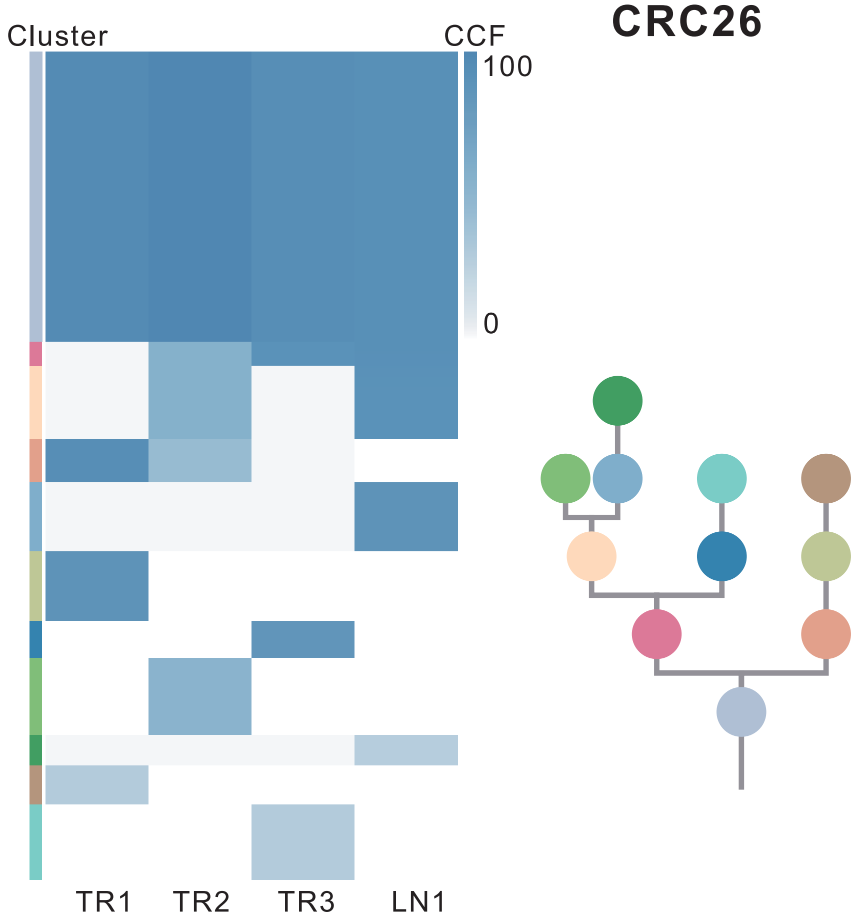

| Gene | Cytoband | Type | Cluster |
|---|---|---|---|
| *LEF1* | 4q25 | SNV | 1 |
| *FBXW7* | 4q31.3 | SNV | 1 |
| * *APC* | 5q22.2 | SNV | 1 |
| *KRAS* | 12p12.1 | SNV | 1 |
| * *APC* | 5q22.2 | InDel | 1 |
| *TP53* | 17p13.1 | InDel | 1 |
| *FLT3* | 13q12.2 | Amp | 1 |
| *CYSLTR2* | 13q14.2 | Amp | 1 |
| *FOXO1* | 13q14.11 | Amp | 1 |
| *CSMD3* | 8q23.3 | SNV | 3 |
| *EIF3E* | 8q23.1 | SNV | 5 |
| *SMAD3* | 15q22.33 | InDel | 6 |
| *MUC4* | 3q29 | Amp | 6 |
| *CREB3L2* | 7q33 | Amp | 6 |
| *TRIM24* | 7q33 | Amp | 6 |
| *BRAF* | 7q34 | Amp | 6 |
| *EZH2* | 7q36.1 | Amp | 6 |
| *SRC* | 20q11.23 | Amp | 6 |
| *MAFB* | 20q12 | Amp | 6 |
| *PLCG1* | 20q12 | Amp | 6 |
| *NFATC2* | 20q13.2 | Amp | 6 |
| *SALL4* | 20q13.2 | Amp | 6 |
| *GNAS* | 20q13.32 | Amp | 6 |
| *PTK6* | 20q13.33 | Amp | 6 |
| *AXIN1* | 16p13.3 | Del | 7 |
| *FGFR1* | 8p11.23 | Amp | NA |

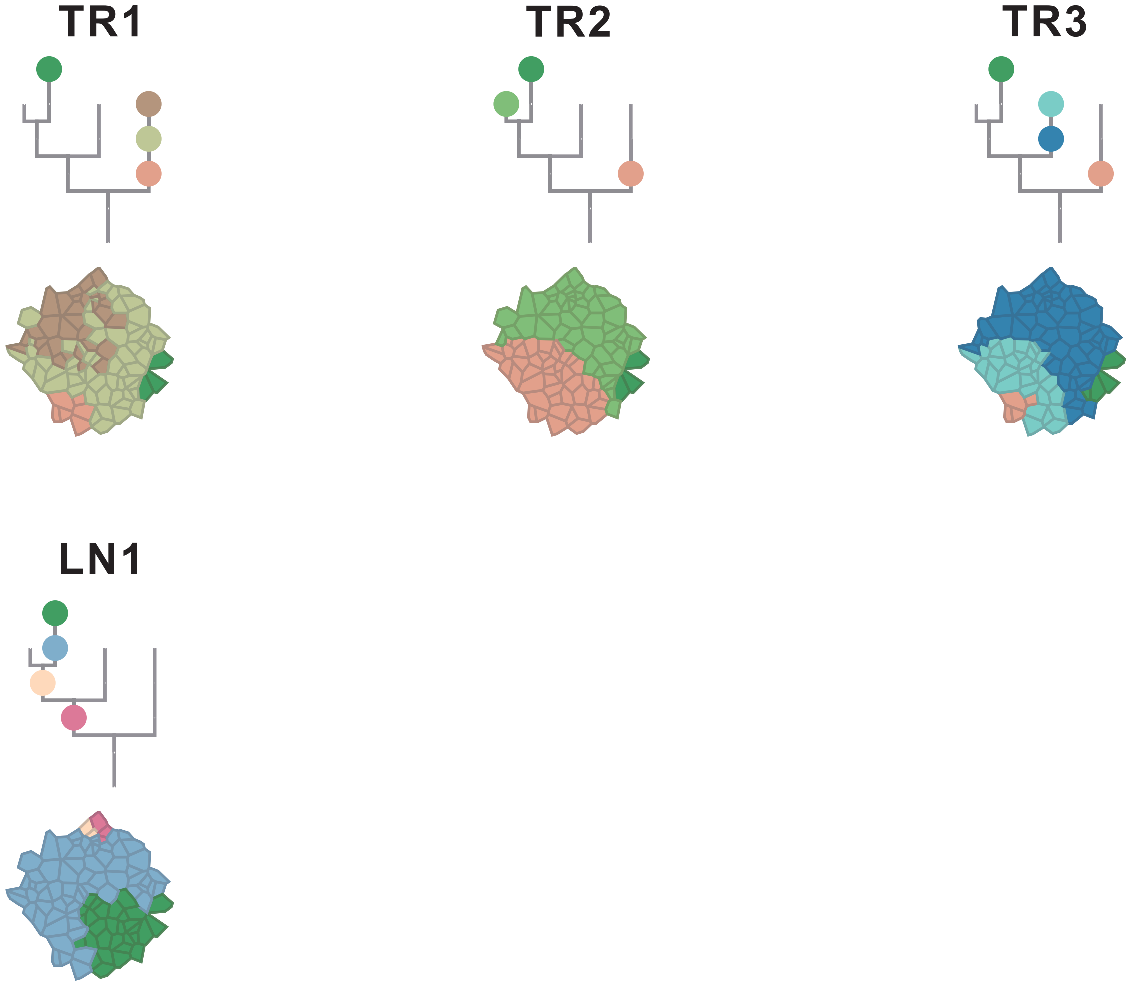

### CRC27

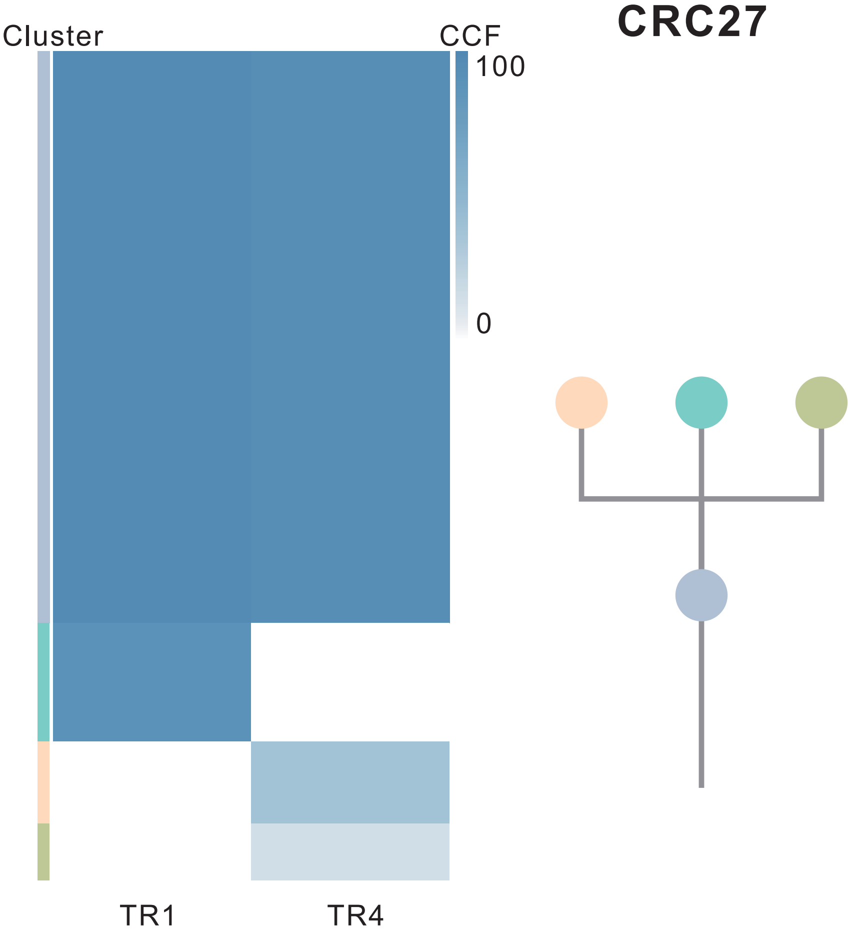

| Gene | Cytoband | Type | Cluster |
|---|---|---|---|
| * *APC* | 5q22.2 | SNV | 1 |
| * *TP53* | 17p13.1 | SNV | 1 |
| * *TP53* | 17p13.1 | SNV | 1 |
| * *APC* | 5q22.2 | InDel | 1 |
| *LZTR1* | 22q11.21 | InDel | 1 |
| *MUC4* | 3q29 | Amp | 1 |
| *SRC* | 20q11.23 | Amp | 1 |
| *MAFB* | 20q12 | Amp | 1 |
| *PLCG1* | 20q12 | Amp | 1 |
| *NFATC2* | 20q13.2 | Amp | 1 |
| *SALL4* | 20q13.2 | Amp | 1 |
| *SSX2* | Xp11.22 | Amp | 1 |
| *ARAF* | Xp11.23 | Amp | 1 |
| *GATA1* | Xp11.23 | Amp | 1 |
| *SSX1* | Xp11.23 | Amp | 1 |
| *SSX4* | Xp11.23 | Amp | 1 |
| *TFE3* | Xp11.23 | Amp | 1 |
| *WAS* | Xp11.23 | Amp | 1 |
| *KDM6A* | Xp11.3 | Amp | 1 |
| *IRS4* | Xq22.3 | Del | 1 |
| *STAG2* | Xq25 | Del | 1 |
| *BCORL1* | Xq26.1 | Del | 1 |
| *ELF4* | Xq26.1 | Del | 1 |
| *GPC3* | Xq26.2 | Del | 1 |
| *PHF6* | Xq26.2 | Del | 1 |
| *ATP2B3* | Xq28 | Del | 1 |
| *RPL10* | Xq28 | Del | 1 |
| *NOTCH2* | 1p12 | Amp | NA |
| *BCL9* | 1q21.2 | Amp | NA |

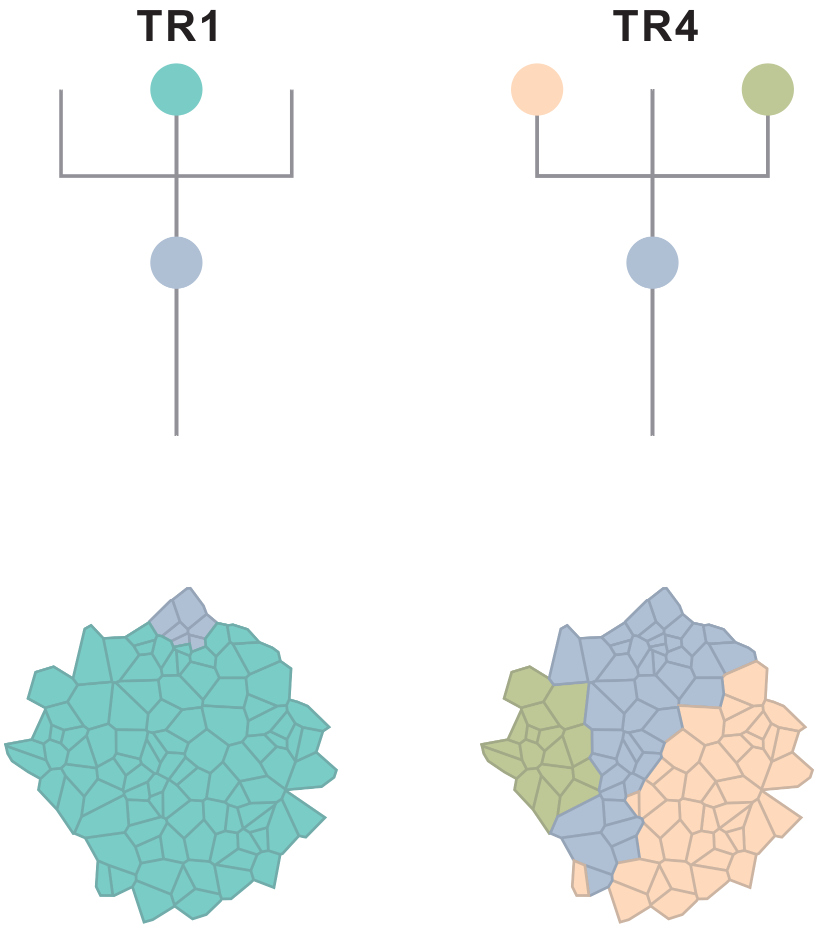

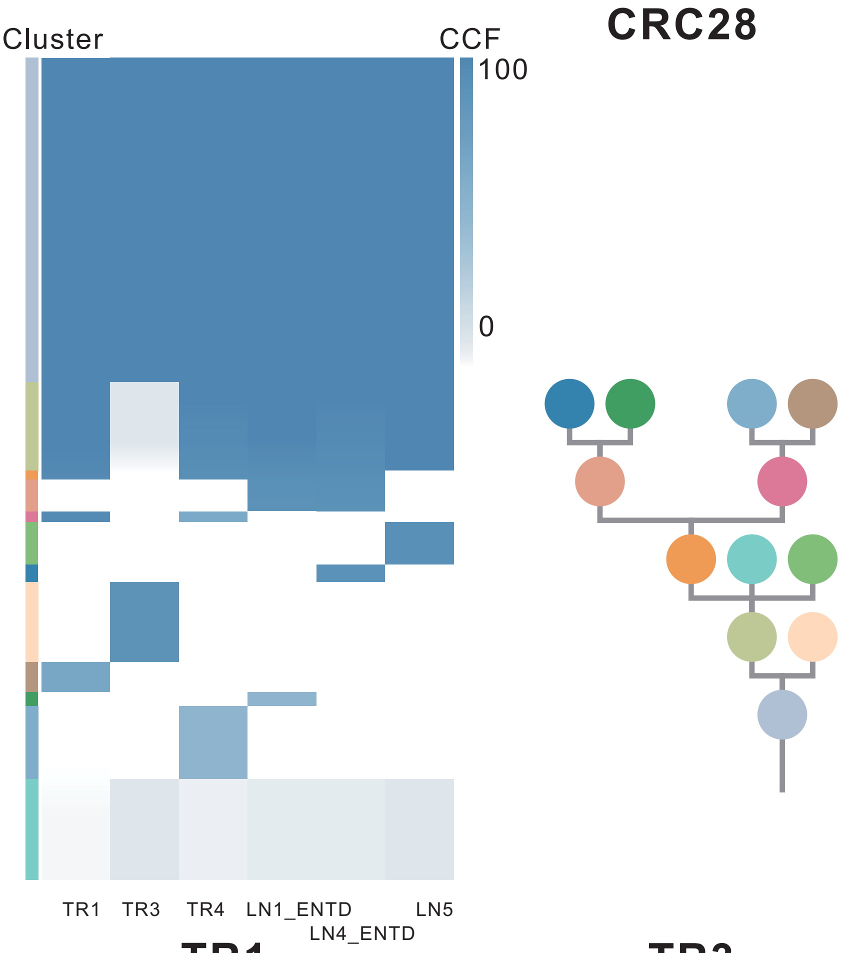

| Gene | Cytoband | Type | Cluster |
|---|---|---|---|
| * *APC* | 5q22.2 | SNV | 1 |
| * *APC* | 5q22.2 | SNV | 1 |
| *KRAS* | 12p12.1 | SNV | 1 |
| *CLTCL1* | 22q11.21 | SNV | 1 |
| *KRAS* | 12p12.1 | Amp | 1 |
| *CHD4* | 12p13.31 | Amp | 1 |
| *CCND2* | 12p13.32 | Amp | 1 |
| *KDM5A* | 12p13.33 | Amp | 1 |
| *IRS4* | Xq22.3 | Amp | 1 |
| *BCORL1* | Xq26.1 | Amp | 1 |
| *ELF4* | Xq26.1 | Amp | 1 |
| *GPC3* | Xq26.2 | Amp | 1 |
| *FAT4* | 4q28.1 | SNV | 2 |
| *ARID1A* | 1p36.11 | SNV | 3 |
| *ID3* | 1p36.12 | Del | 4 |
| *ARHGEF10L* | 1p36.13 | Del | 4 |
| *SDHB* | 1p36.13 | Del | 4 |
| *CAMTA1* | 1p36.31 | Del | 4 |
| *RPL22* | 1p36.31 | Del | 4 |
| *TNFRSF14* | 1p36.32 | Del | 4 |
| *CASP8* | 2q33.1 | SNV | 6 |
| *JAK2* | 9p24.1 | Amp | 6 |
| *PDCD1LG2* | 9p24.1 | Amp | 6 |
| *KDM5C* | Xp11.22 | SNV | NA |

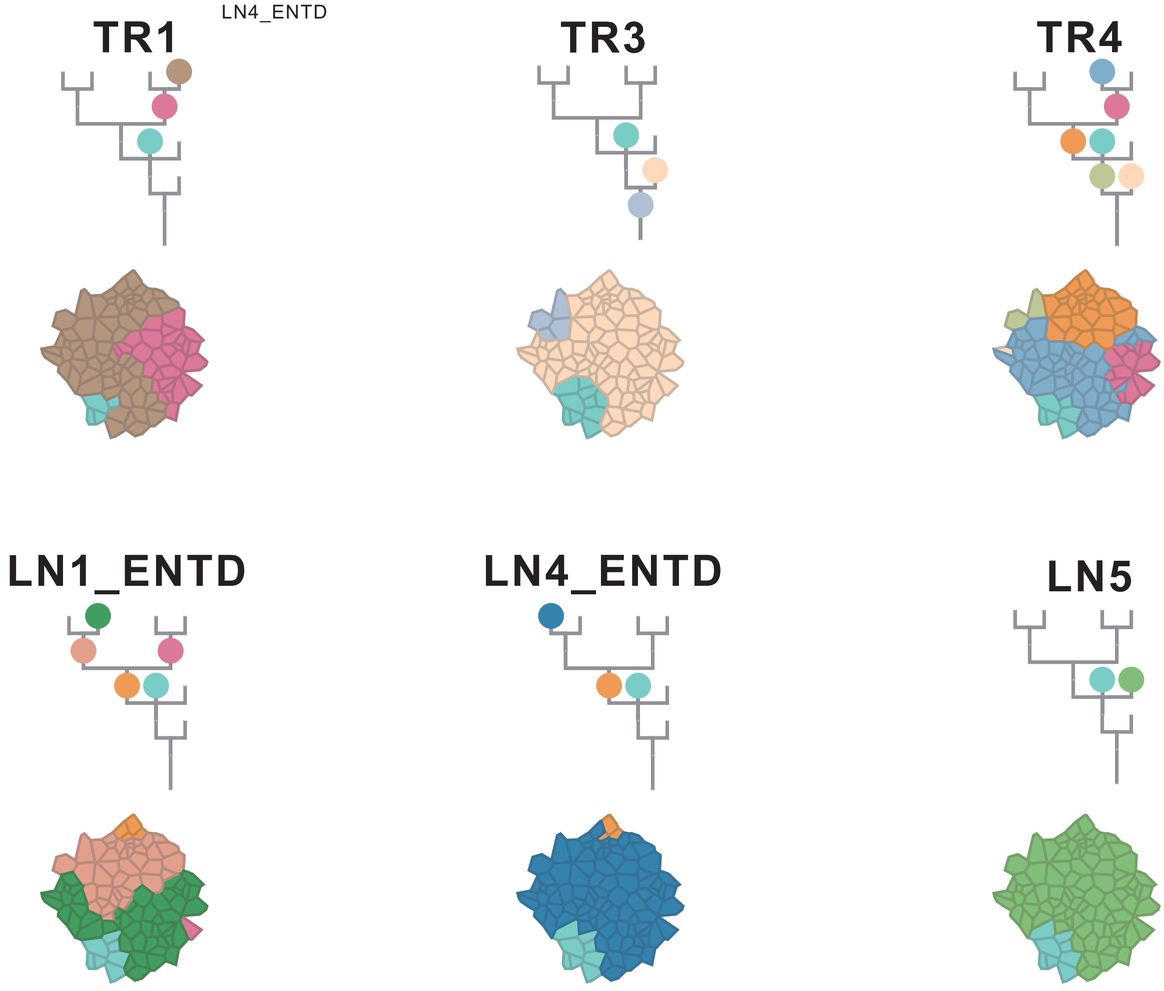

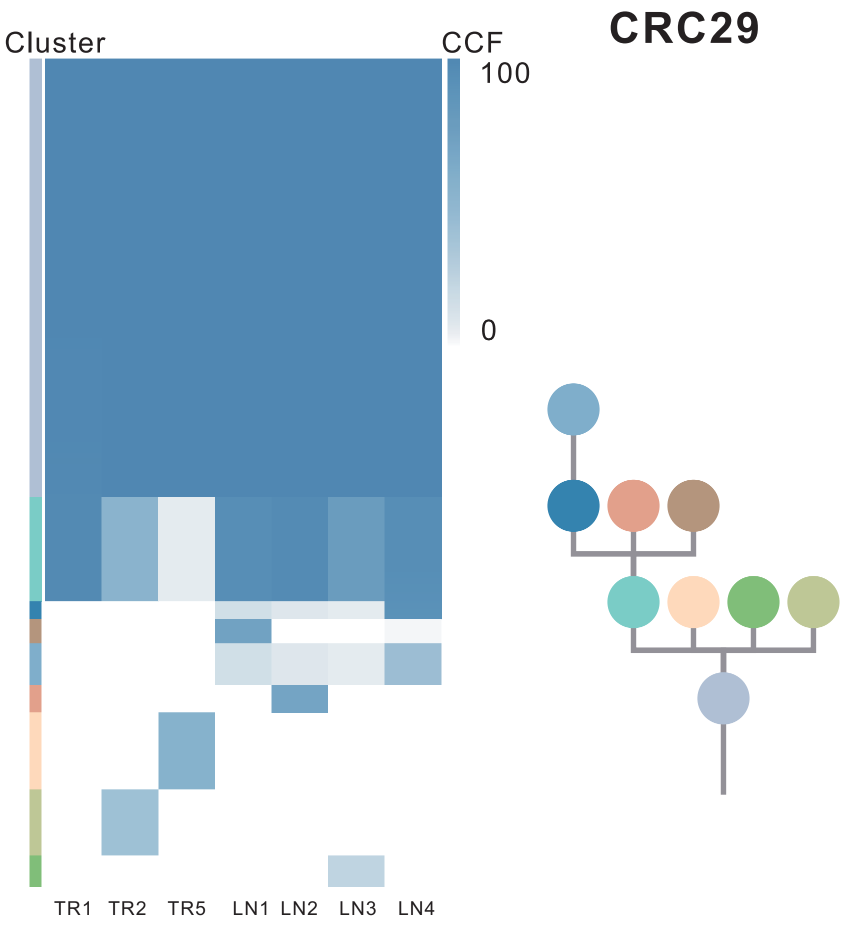

| Gene | Cytoband | Type | Cluster |
|---|---|---|---|
| *APC* | 5q22.2 | SNV | 1 |
| *BCL9L* | 11q23.3 | SNV | 1 |
| *NCOR2* | 12q24.31 | SNV | 1 |
| *TP53* | 17p13.1 | SNV | 1 |
| *FLT3* | 13q12.2 | Amp | 1 |
| *SRC* | 20q11.23 | Amp | 1 |
| *MAFB* | 20q12 | Amp | 1 |
| *PLCG1* | 20q12 | Amp | 1 |
| *NFATC2* | 20q13.2 | Amp | 1 |
| *SALL4* | 20q13.2 | Amp | 1 |
| *GNAS* | 20q13.32 | Amp | 1 |
| *PTK6* | 20q13.33 | Amp | 1 |
| *DDX5* | 17q23.3 | Amp | 2 |
| *SMAD3* | 15q22.33 | SNV | 4 |
| *LZTR1* | 22q11.21 | InDel | NA |
| *MUC4* | 3q29 | Amp | NA |
| *EGFR* | 7p11.2 | Amp | NA |
| *PLAG1* | 8q12.1 | Amp | NA |
| *PREX2* | 8q13.2 | Amp | NA |
| *NCOA2* | 8q13.3 | Amp | NA |
| *HEY1* | 8q21.13 | Amp | NA |
| *RUNX1T1* | 8q21.3 | Amp | NA |
| *CDH17* | 8q22.1 | Amp | NA |
| *PABPC1* | 8q22.3 | Amp | NA |
| *UBR5* | 8q22.3 | Amp | NA |
| *RAD21* | 8q24.11 | Amp | NA |
| *MYC* | 8q24.21 | Amp | NA |
| *RECQL4* | 8q24.3 | Amp | NA |
| *PPM1D* | 17q23.2 | Amp | NA |
| *CD79B* | 17q23.3 | Amp | NA |
| *AXIN1* | 16p13.3 | Del | NA |
| *PRKAR1A* | 17q24.2 | Del | NA |

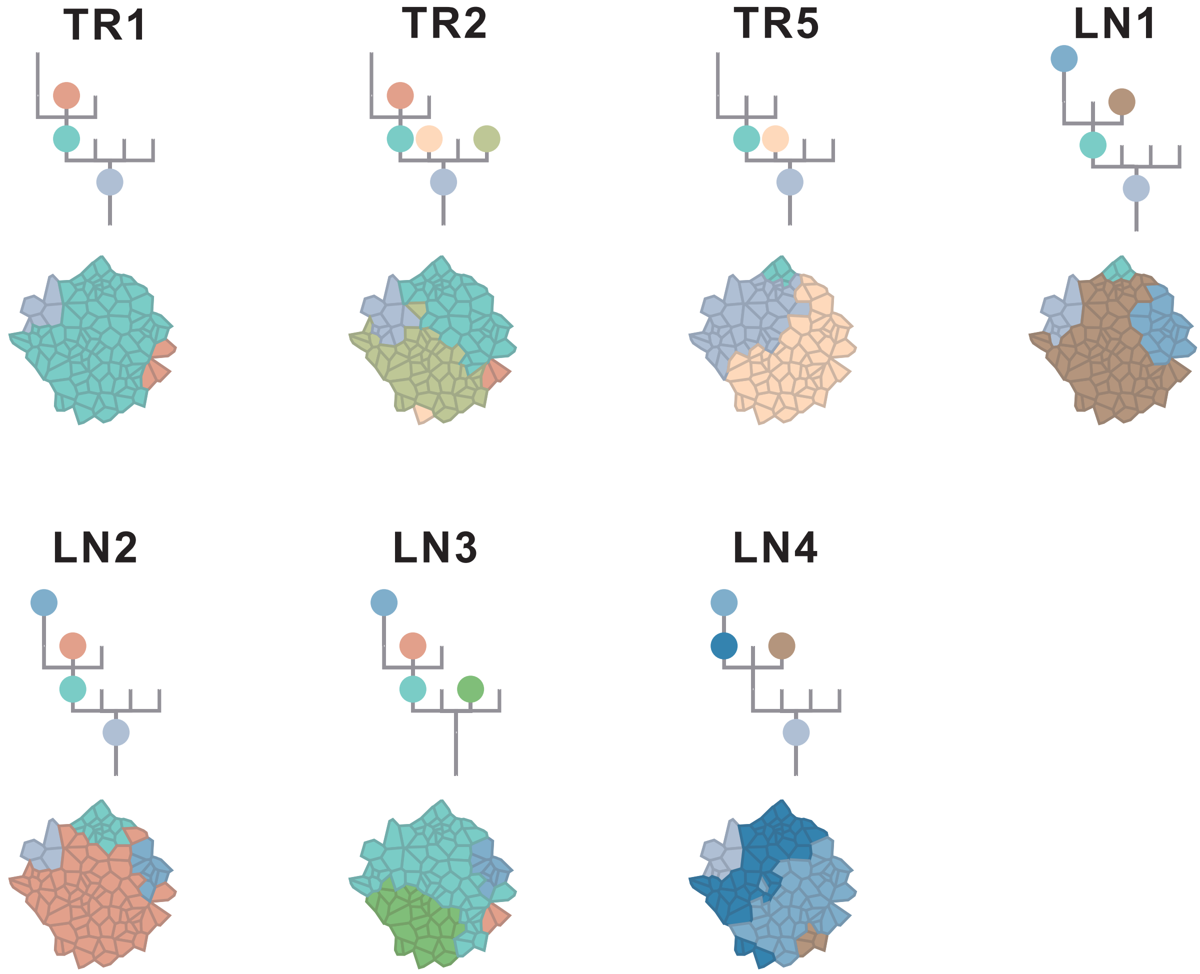

### CRC30

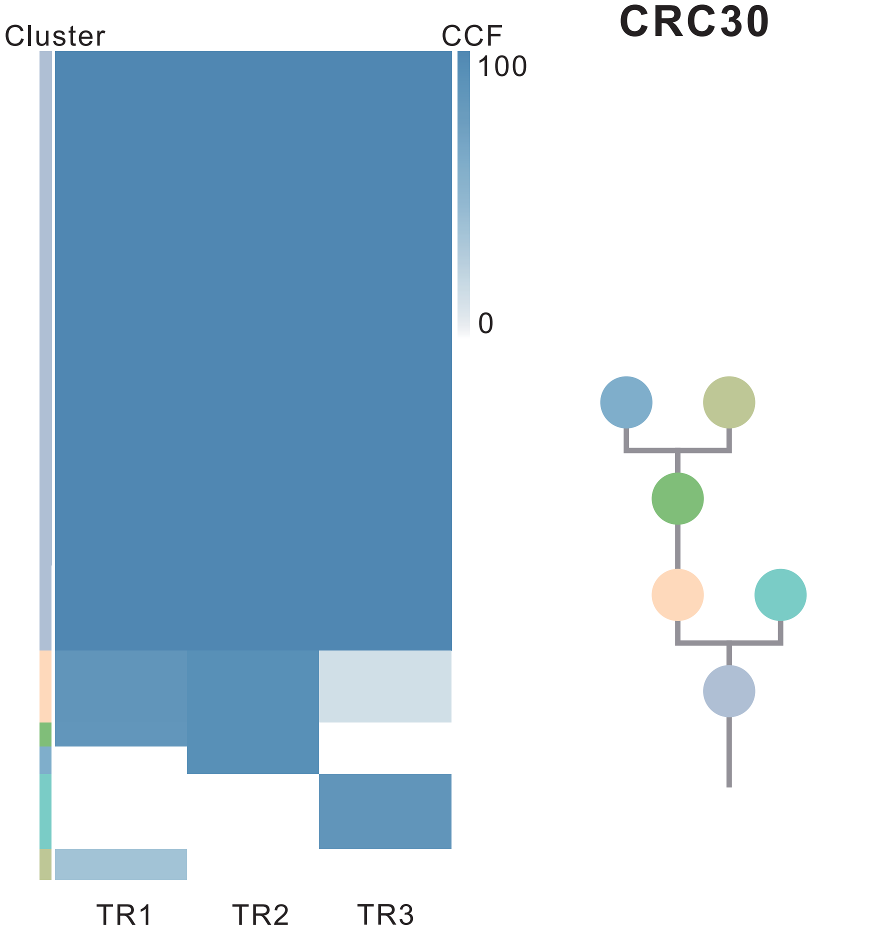

| Gene | Cytoband | Type | Cluster |
|---|---|---|---|
| *APC* | 5q22.2 | SNV | 1 |
| *PTPRB* | 12q15 | SNV | 1 |
| *TP53* | 17p13.1 | SNV | 1 |
| *IL7R* | 5p13.2 | Amp | 1 |
| *IKBKB* | 8p11.21 | Amp | 1 |
| *KAT6A* | 8p11.21 | Amp | 1 |
| *PLAG1* | 8q12.1 | Amp | 1 |
| *PREX2* | 8q13.2 | Amp | 1 |
| *NCOA2* | 8q13.3 | Amp | 1 |
| *HEY1* | 8q21.13 | Amp | 1 |
| *RUNX1T1* | 8q21.3 | Amp | 1 |
| *CDH17* | 8q22.1 | Amp | 1 |
| *PABPC1* | 8q22.3 | Amp | 1 |
| *UBR5* | 8q22.3 | Amp | 1 |
| *RAD21* | 8q24.11 | Amp | 1 |
| *MYC* | 8q24.21 | Amp | 1 |
| *RECQL4* | 8q24.3 | Amp | 1 |

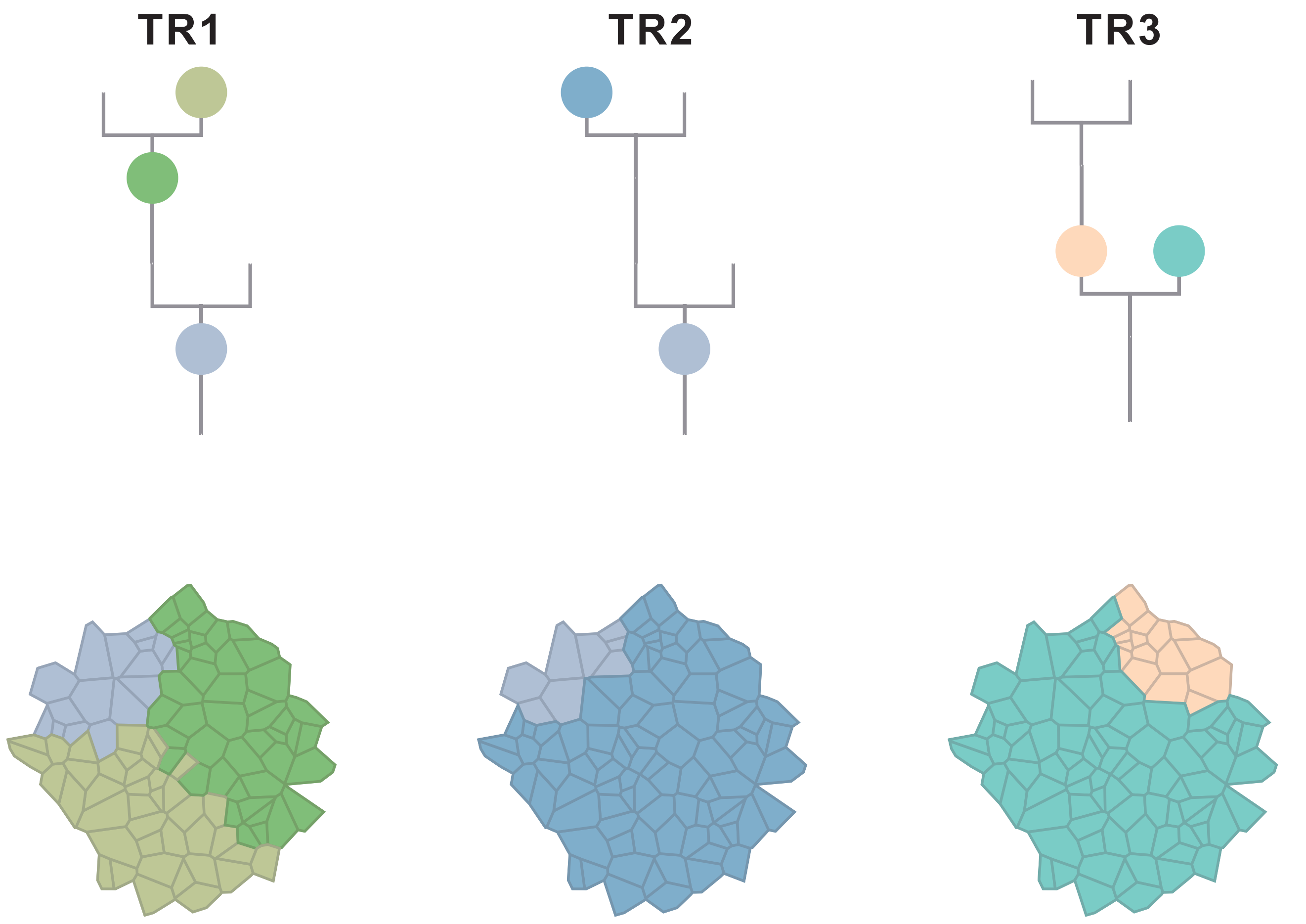

### CRC31

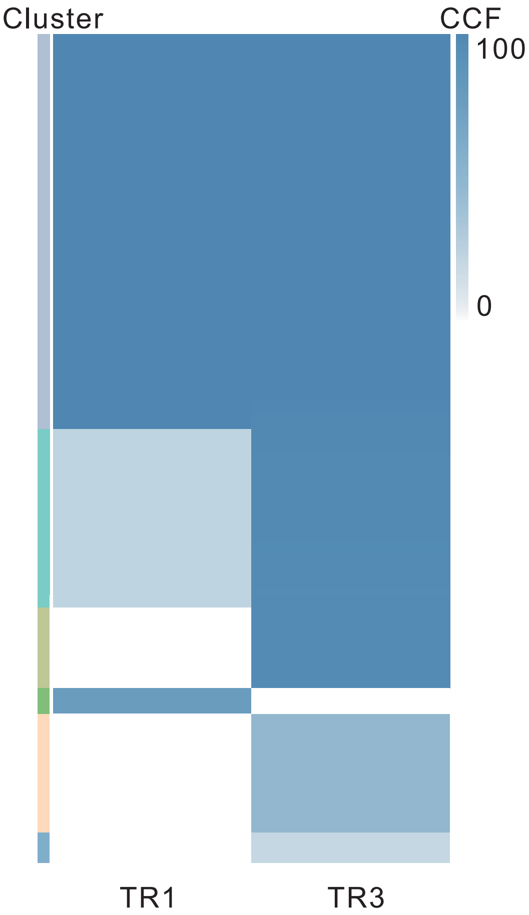

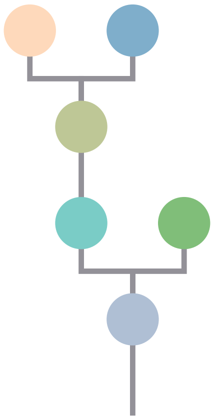

| Gene | Cytoband | Type | Cluster |
|---|---|---|---|
| *APC* | 5q22.2 | SNV | 1 |
| *KRAS* | 12p12.1 | SNV | 1 |
| *TP53* | 17p13.1 | SNV | 1 |
| *ARID1A* | 1p36.11 | SNV | 2 |
| *CLTCL1* | 22q11.21 | SNV | 3 |

## TR1

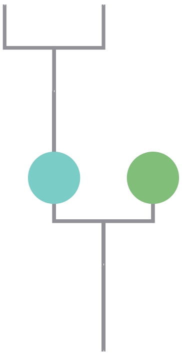

## TR3

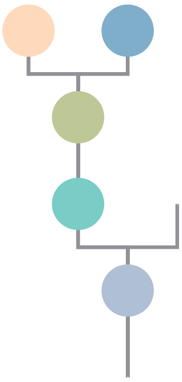

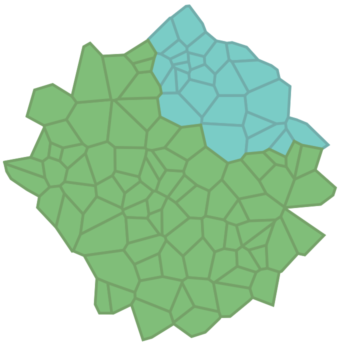

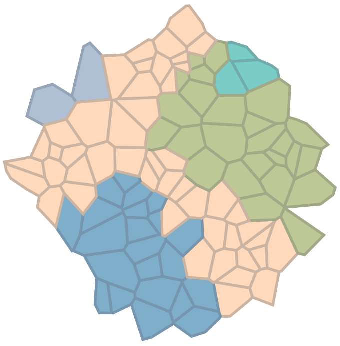

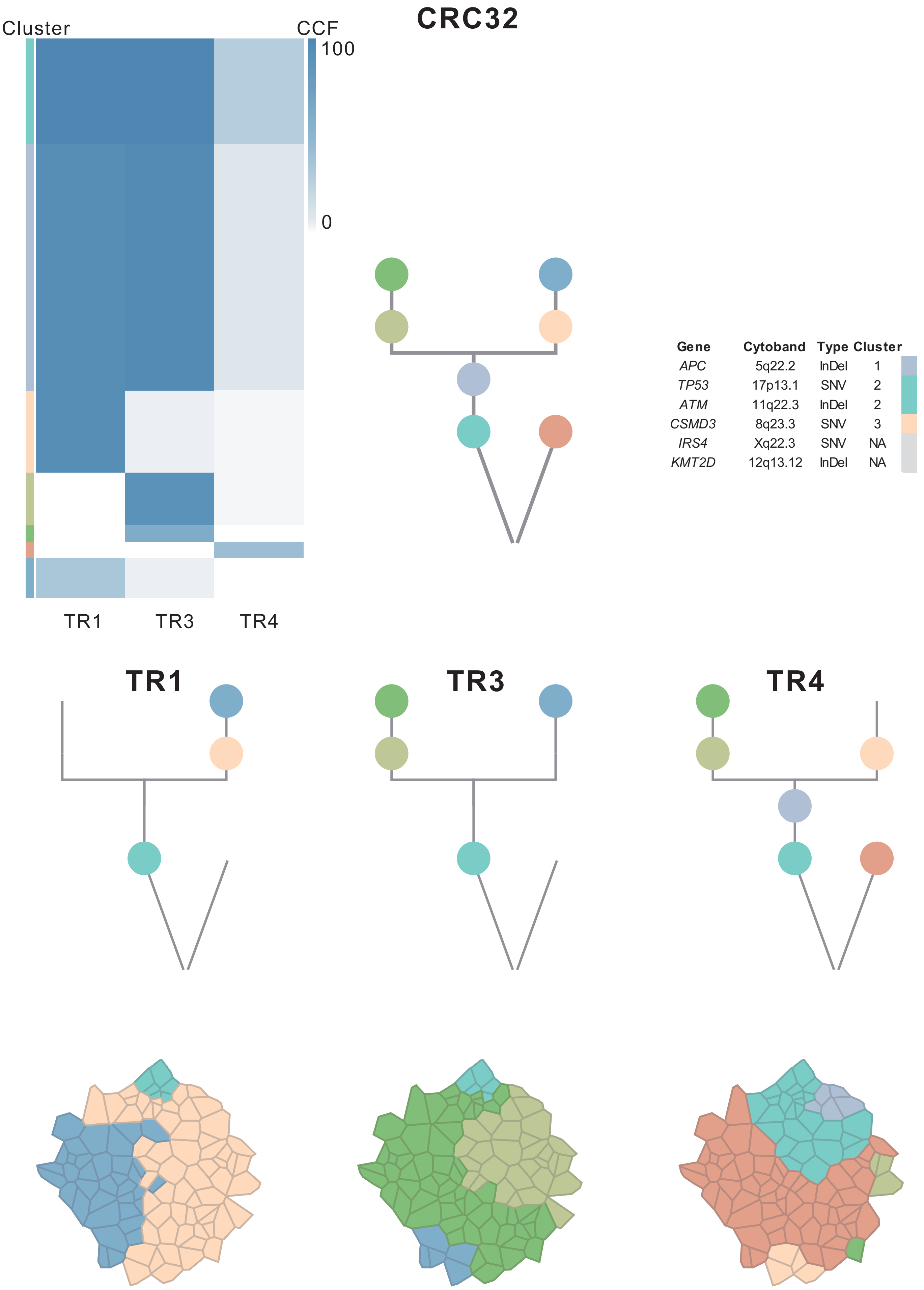
CRC32
Cluster
CCF
100
0
TR1
TR3
TR4
Gene Cytoband Type Cluster
APC 5q22.2 InDel 1
TP53 17p13.1 SNV 2
ATM 11q22.3 InDel 2
CSMD3 8q23.3 SNV 3
IRS4 Xq22.3 SNV NA
KMT2D 12q13.12 InDel NA
TR1
TR3
TR4

#### CRC33

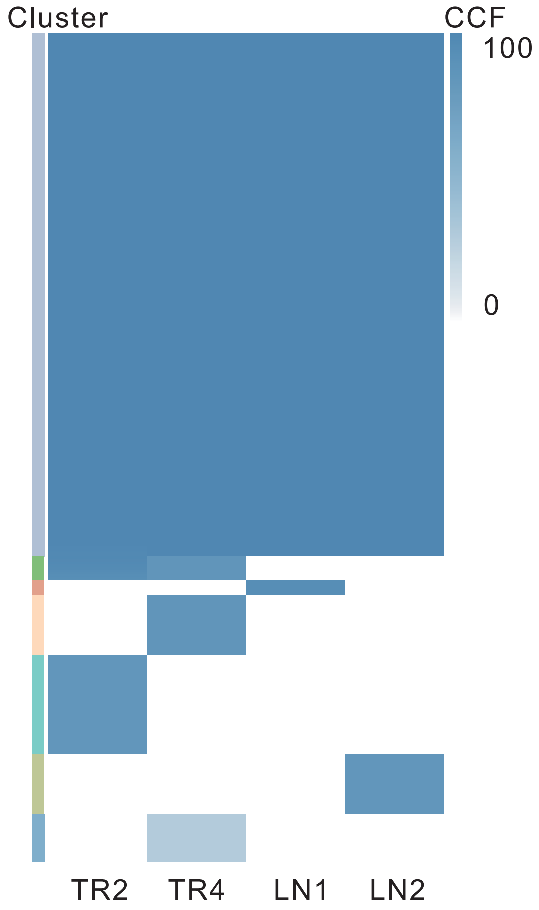

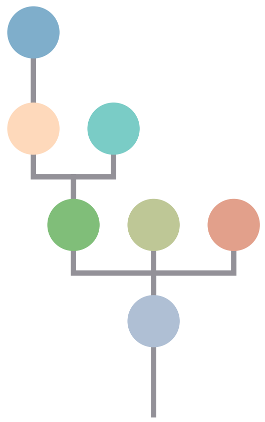

| Gene | Cytoband | Type | Cluster |
|---|---|---|---|
| *TP53* | 17p13.1 | SNV | 1 |
| *JAK3* | 19p13.11 | SNV | 1 |
| *APC* | 5q22.2 | InDel | 1 |
| *FLT3* | 13q12.2 | Amp | 1 |
| *FOXO1* | 13q14.11 | Amp | 1 |
| *CYSLTR2* | 13q14.2 | Amp | 1 |
| *SRC* | 20q11.23 | Amp | 1 |
| *MAFB* | 20q12 | Amp | 1 |
| *PLCG1* | 20q12 | Amp | 1 |
| *NFATC2* | 20q13.2 | Amp | 1 |
| *SALL4* | 20q13.2 | Amp | 1 |
| *GNAS* | 20q13.32 | Amp | 1 |
| *PTK6* | 20q13.33 | Amp | 1 |
| *NRG1* | 8p12 | SNV | 4 |
| *EGFR* | 7p11.2 | Amp | 4 |
| *IRS4* | Xq22.3 | InDel | NA |

**TR2**

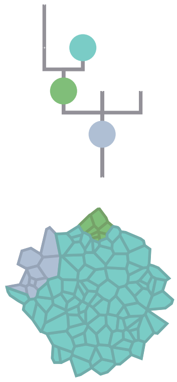

**TR4**

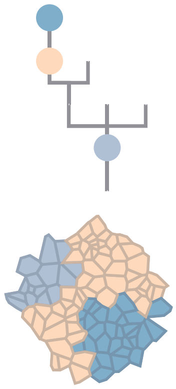

**LN1**

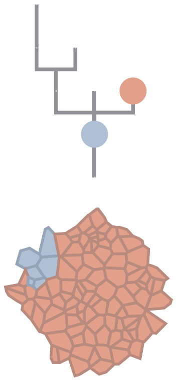

**LN2**

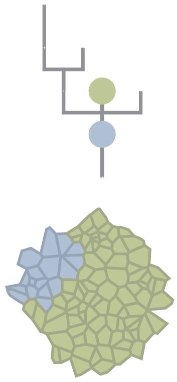

### CRC34

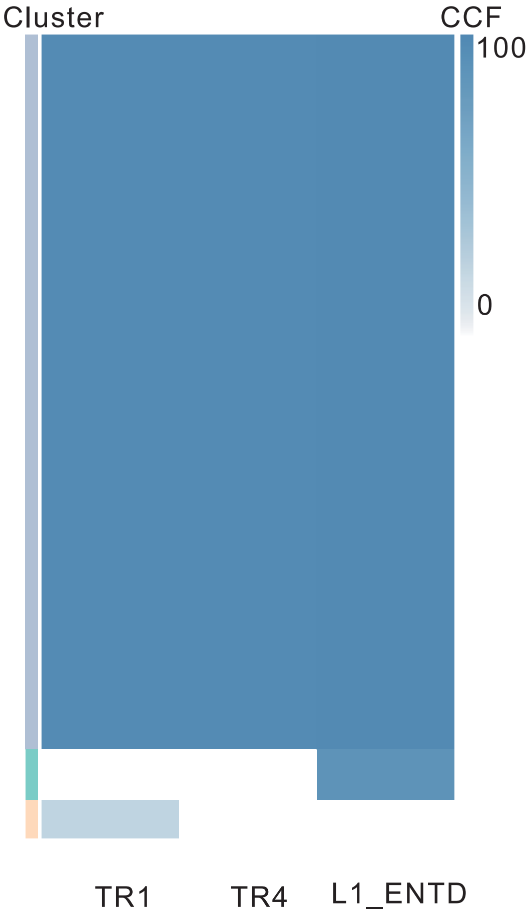

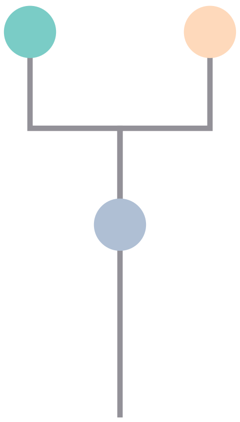

| Gene | Cytoband | Type | Cluster |
|---|---|---|---|
| * *APC* | 5q22.2 | SNV | 1 |
| * *APC* | 5q22.2 | SNV | 1 |
| *KRAS* | 12p12.1 | SNV | 1 |
| *TP53* | 17p13.1 | SNV | 1 |
| *CSMD3* | 8q23.3 | InDel | 1 |
| *AR* | Xq12 | SNV | NA |

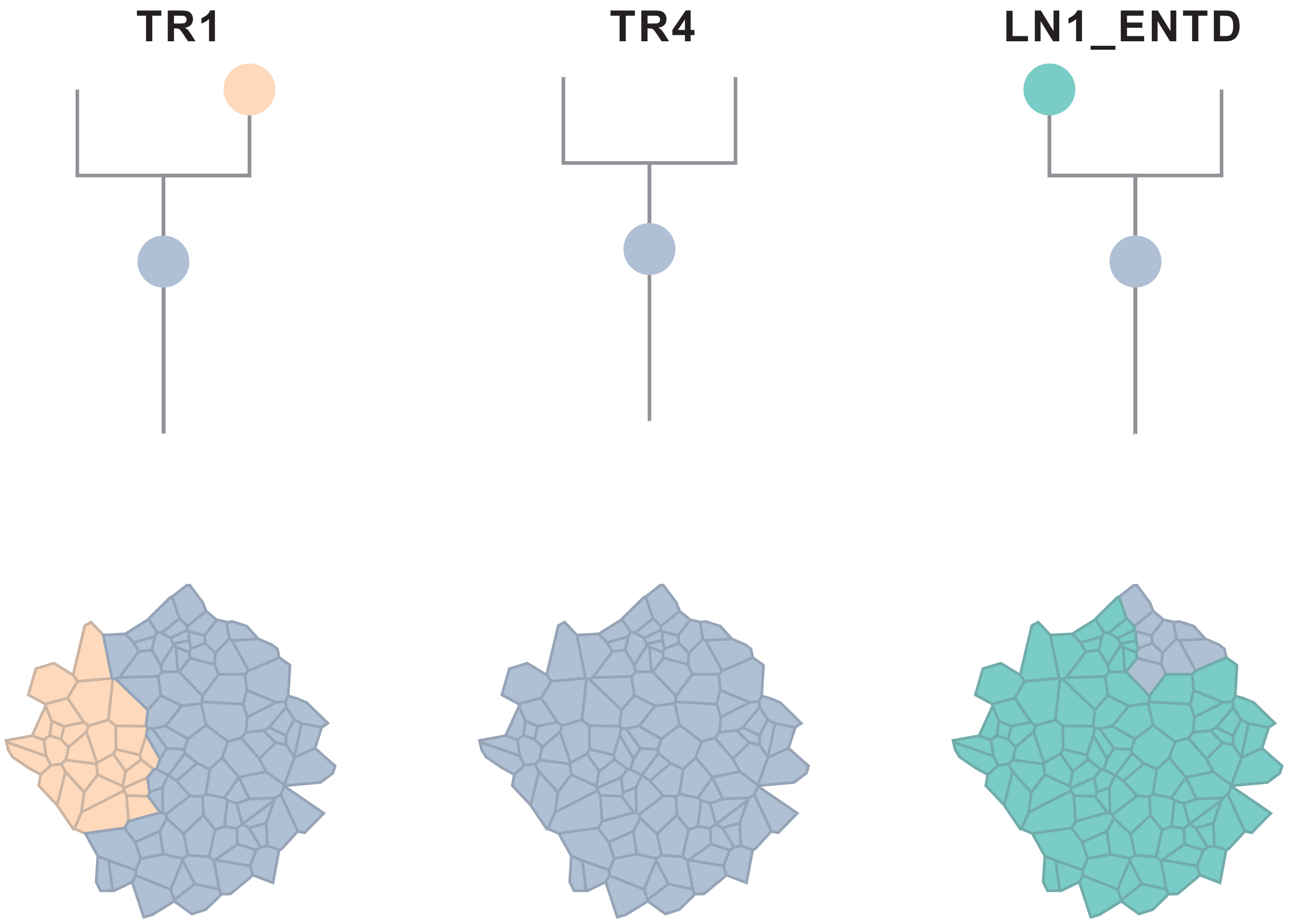

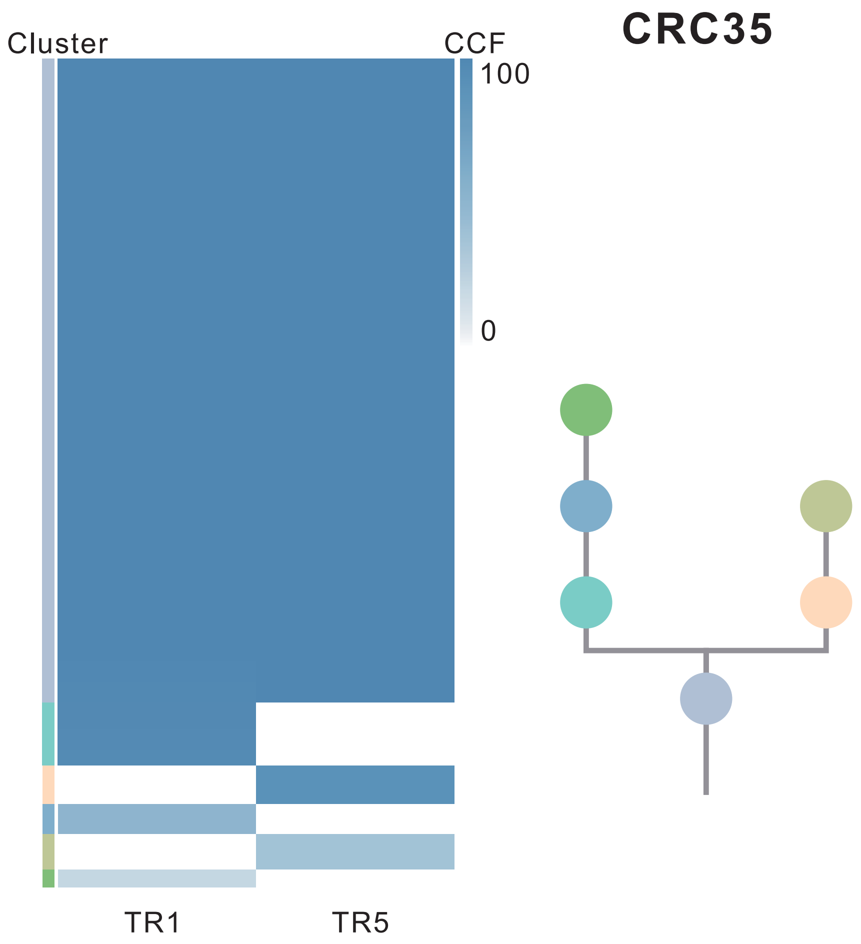

| Gene | Cytoband | Type | Cluster |
|---|---|---|---|
| *APC* | 5q22.2 | SNV | 1 |
| *ATM* | 11q22.3 | SNV | 1 |
| *KRAS* | 12p12.1 | SNV | 1 |
| *POLG* | 15q26.1 | SNV | 1 |
| *TP53* | 17p13.1 | SNV | 1 |
| *IL7R* | 5p13.2 | Amp | 1 |
| *CTNND2* | 5p15.2 | Amp | 1 |
| *TERT* | 5p15.33 | Amp | 1 |
| *SSX2* | Xp11.22 | Amp | 1 |
| *ARAF* | Xp11.23 | Amp | 1 |
| *GATA1* | Xp11.23 | Amp | 1 |
| *SSX1* | Xp11.23 | Amp | 1 |
| *SSX4* | Xp11.23 | Amp | 1 |
| *TFE3* | Xp11.23 | Amp | 1 |
| *WAS* | Xp11.23 | Amp | 1 |
| *KDM6A* | Xp11.3 | Amp | 1 |

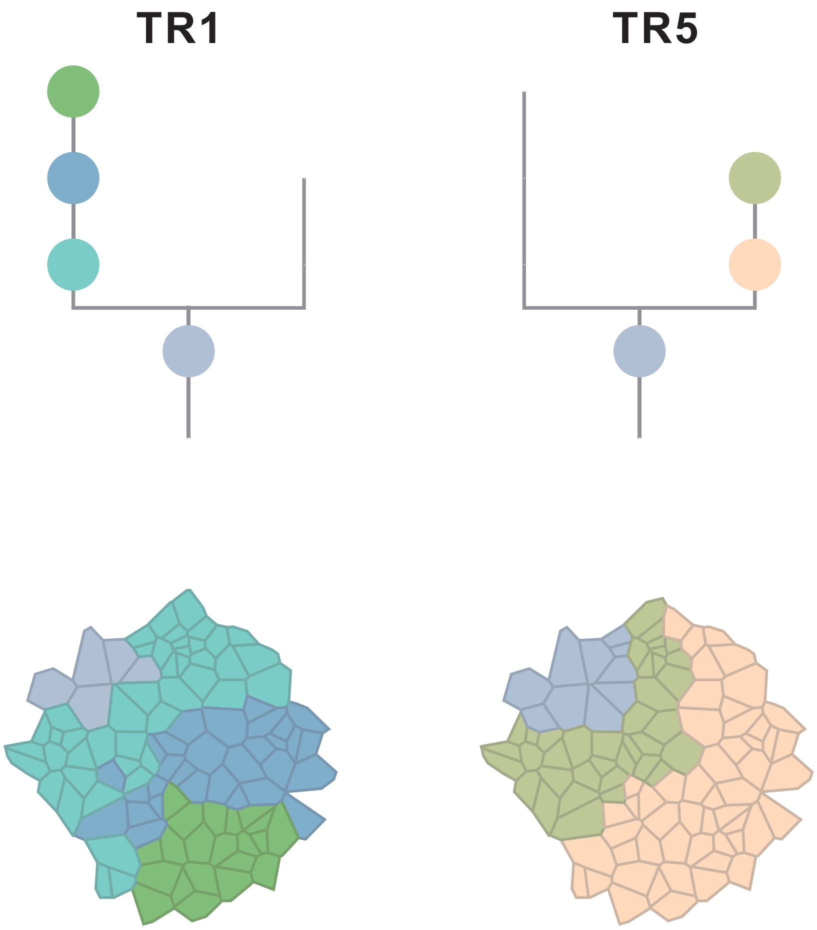

### CRC36

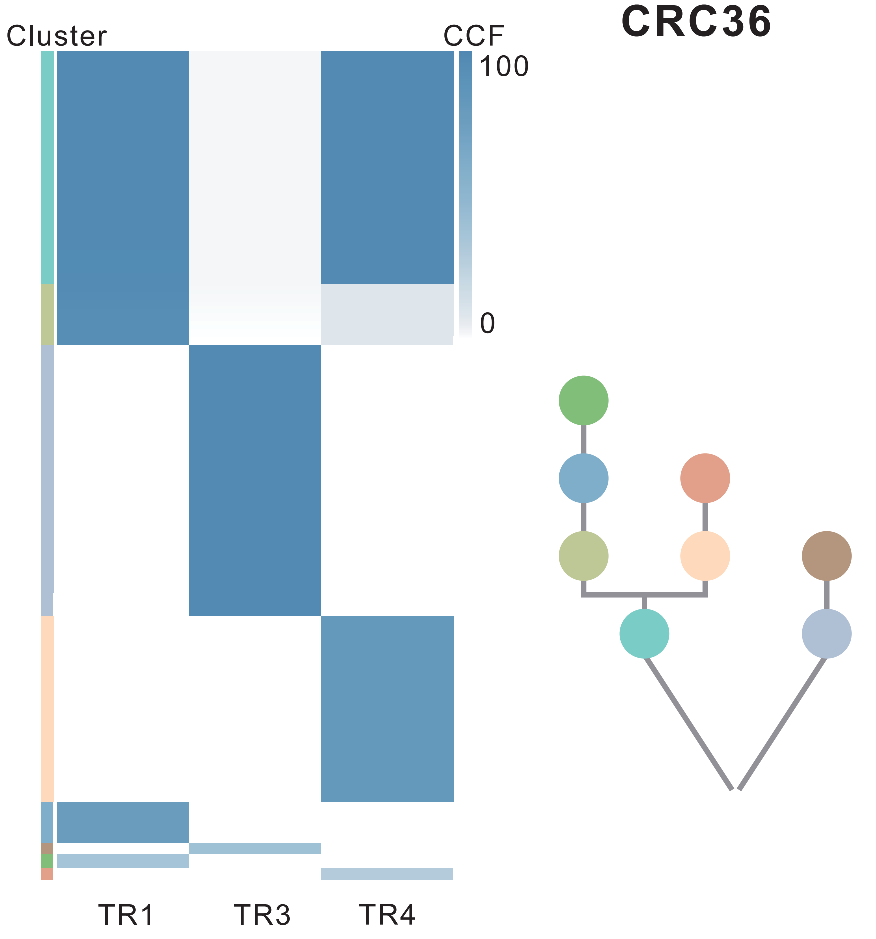

| Gene | Cytoband | Type | Cluster |
|---|---|---|---|
| *FBXO11* | 2p16.3 | SNV | 1 |
| *FBXW7* | 4q31.3 | SNV | 1 |
| * *APC* | 5q22.2 | SNV | 1 |
| *WRN* | 8p12 | SNV | 1 |
| *ATM* | 11q22.3 | SNV | 1 |
| *KRAS* | 12p12.1 | SNV | 1 |
| * *TP53* | 17p13.1 | SNV | 1 |
| * *TP53* | 17p13.1 | SNV | 1 |
| *PTPRT* | 20q12 | SNV | 1 |
| *ERBB4* | 2q34 | SNV | 2 |
| * *APC* | 5q22.2 | SNV | 2 |
| *ABI1* | 10p12.1 | SNV | 2 |
| * *CREBBP* | 16p13.3 | SNV | 2 |
| * *CREBBP* | 16p13.3 | SNV | 2 |
| * *TP53* | 17p13.1 | SNV | 2 |
| *LZTR1* | 22q11.21 | InDel | 2 |
| *NFATC2* | 20q13.2 | Amp | 2 |
| *SALL4* | 20q13.2 | Amp | 2 |
| *GNAS* | 20q13.32 | Amp | 2 |
| *RAC1* | 7p22.1 | Amp | 3 |
| *CARD11* | 7p22.2 | Amp | 3 |
| *SRC* | 20q11.23 | Amp | 3 |
| *MAFB* | 20q12 | Amp | 3 |
| *PLCG1* | 20q12 | Amp | 3 |
| *KMT2C* | 7q36.1 | SNV | 4 |
| *PTK6* | 20q13.33 | Amp | 4 |
| *ZFHX3* | 16q22.3 | SNV | 5 |
| *LRIG3* | 12q14.1 | SNV | 7 |
| *LARP4B* | 10p15.3 | InDel | 8 |
| *BCOR* | Xp11.4 | SNV | NA |

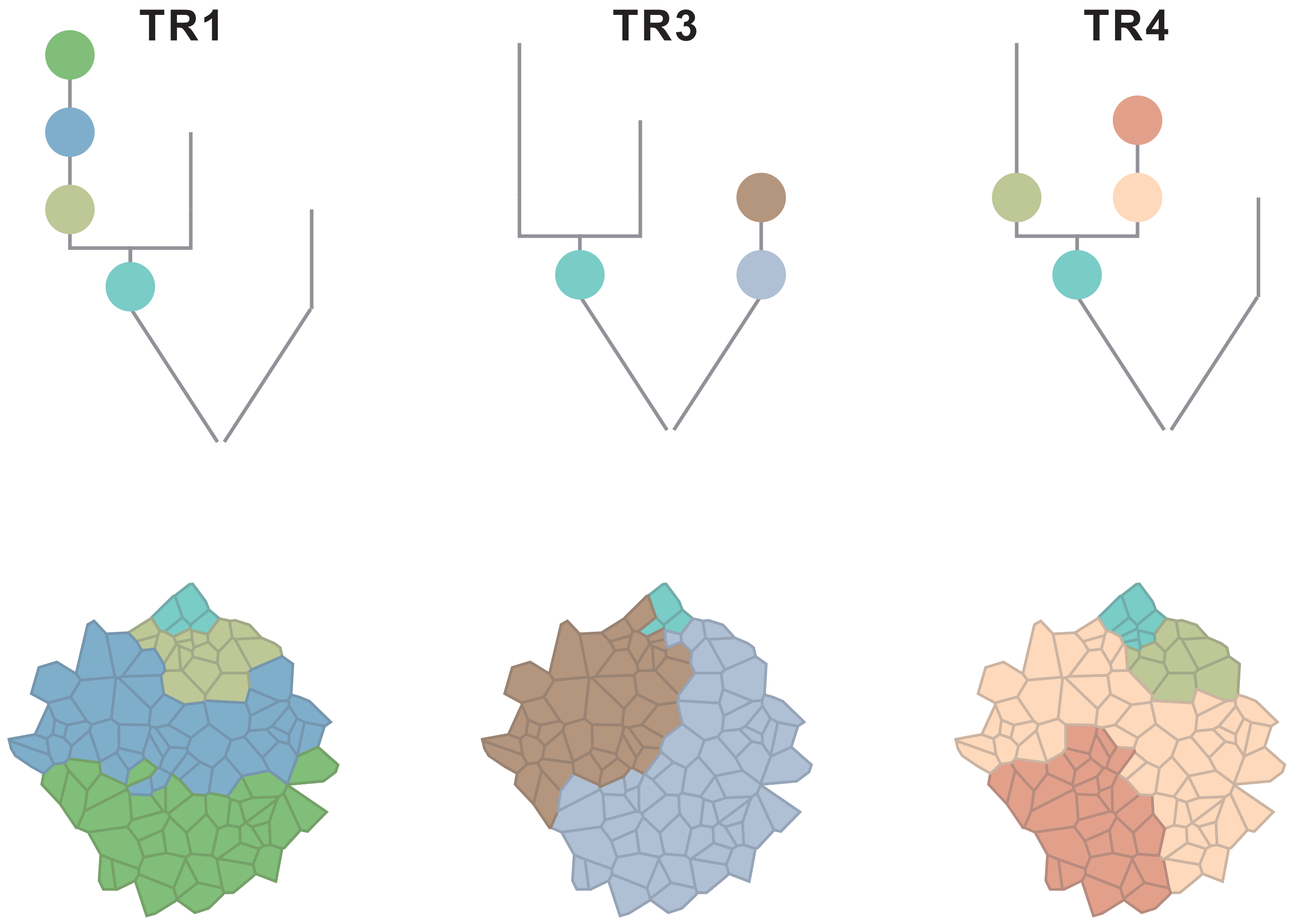

### CRC37

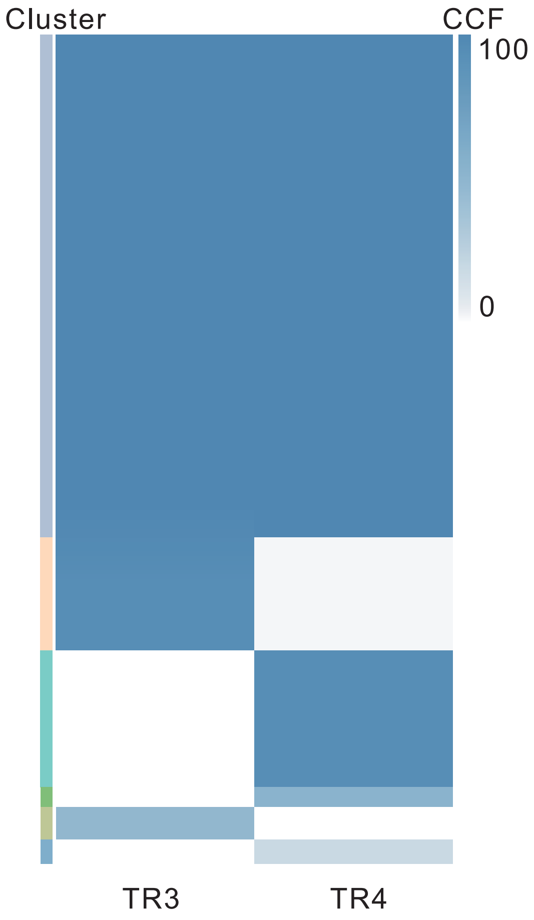

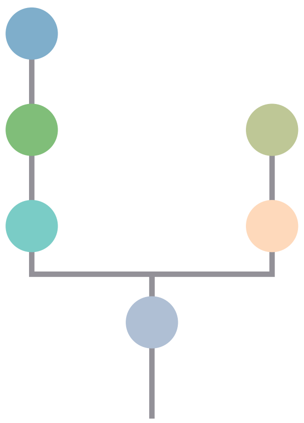

| Gene | Cytoband | Type | Cluster |
|---|---|---|---|
| * *TP53* | 17p13.1 | SNV | 1 |
| * *TP53* | 17p13.1 | SNV | 1 |
| * *APC* | 5q22.2 | InDel | 1 |
| *FLT3* | 13q12.2 | Amp | 1 |
| *SRC* | 20q11.23 | Amp | 1 |
| *MAFB* | 20q12 | Amp | 1 |
| *PLCG1* | 20q12 | Amp | 1 |
| *NFATC2* | 20q13.2 | Amp | 1 |
| *SALL4* | 20q13.2 | Amp | 1 |
| *GNAS* | 20q13.32 | Amp | 1 |
| *PTK6* | 20q13.33 | Amp | 1 |
| * *APC* | 5q22.2 | InDel | NA |

## TR3 TR4

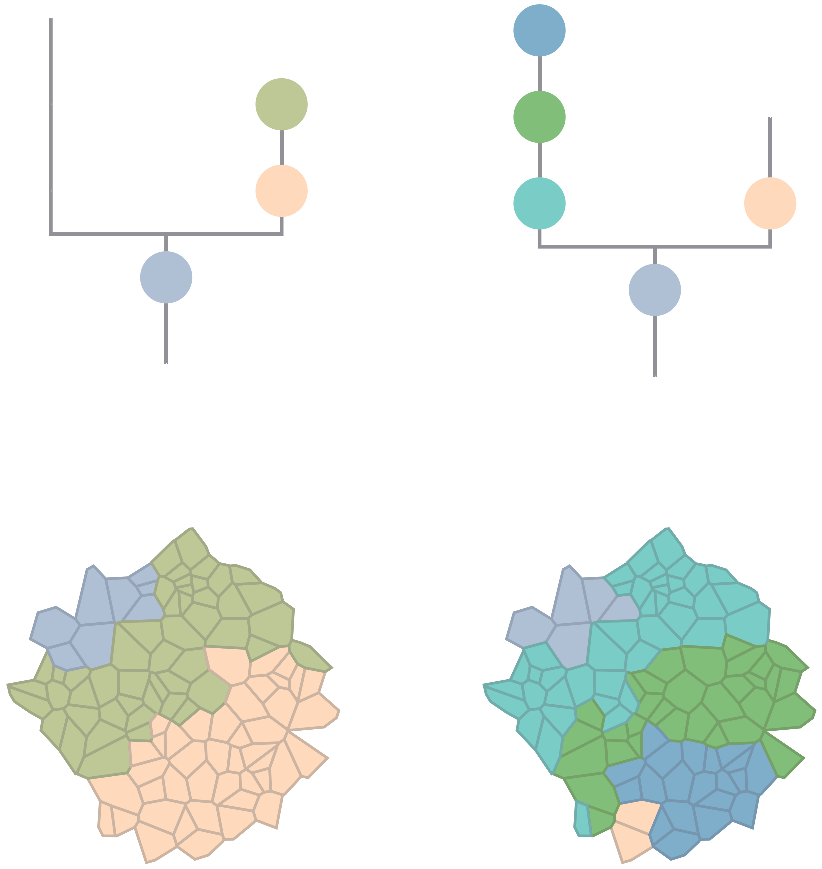

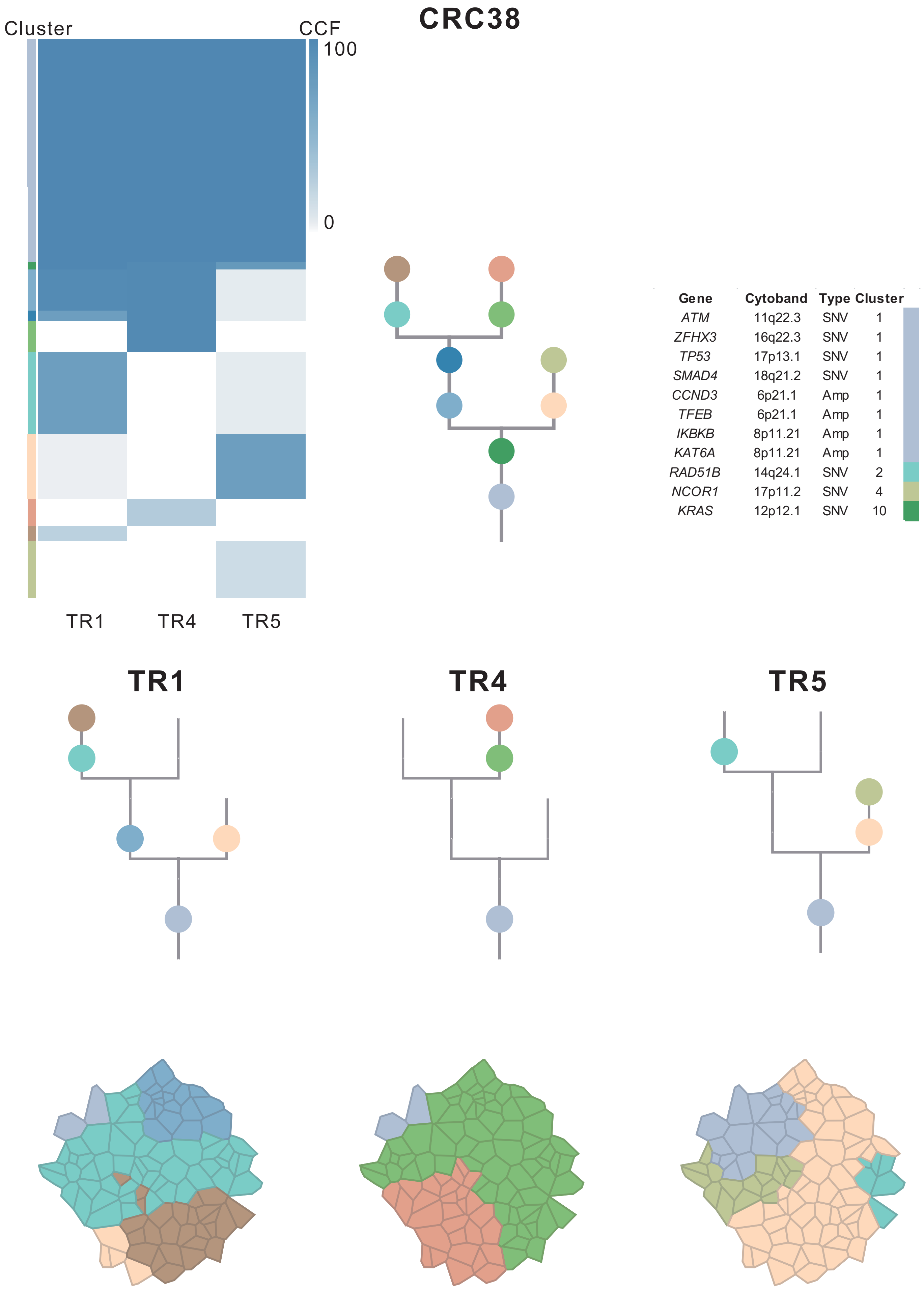

| Gene | Cytoband | Type | Cluster |
| --- | --- | --- | --- |
| *ATM* | 11q22.3 | SNV | 1 |
| *ZFHX3* | 16q22.3 | SNV | 1 |
| *TP53* | 17p13.1 | SNV | 1 |
| *SMAD4* | 18q21.2 | SNV | 1 |
| *CCND3* | 6p21.1 | Amp | 1 |
| *TFEB* | 6p21.1 | Amp | 1 |
| *IKBKB* | 8p11.21 | Amp | 1 |
| *KAT6A* | 8p11.21 | Amp | 1 |
| *RAD51B* | 14q24.1 | SNV | 2 |
| *NCOR1* | 17p11.2 | SNV | 4 |
| *KRAS* | 12p12.1 | SNV | 10 |

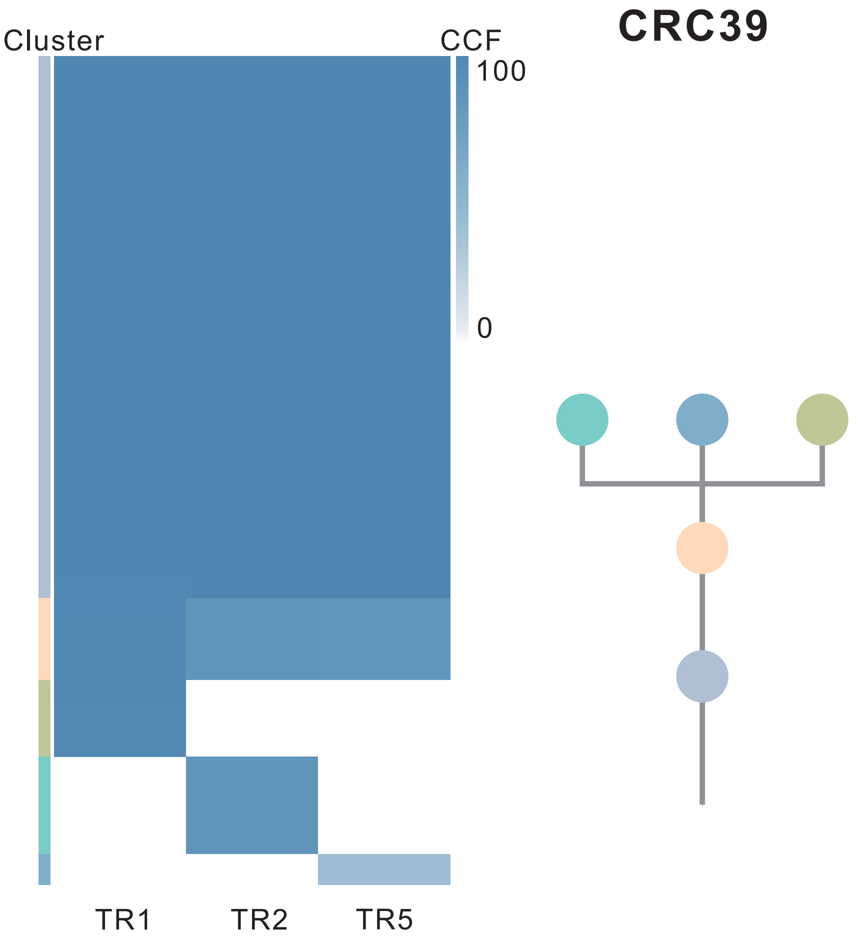

| Gene | Cytoband | Type | Cluster |
| --- | --- | --- | --- |
| *TP53* | 17p13.1 | SNV | 1 |
| *MAP3K1* | 5q11.2 | InDel | 1 |
| *ATM* | 11q22.3 | InDel | 1 |
| *FLT3* | 13q12.2 | Amp | 1 |
| *FOXO1* | 13q14.11 | Amp | 1 |
| *CYSLTR2* | 13q14.2 | Amp | 1 |
| *KRAS* | 12p12.1 | SNV | 3 |
| *FBXW7* | 4q31.3 | SNV | NA |
| *IRS4* | Xq22.3 | Amp | NA |
| *BCORL1* | Xq26.1 | Amp | NA |
| *ELF4* | Xq26.1 | Amp | NA |
| *GPC3* | Xq26.2 | Amp | NA |

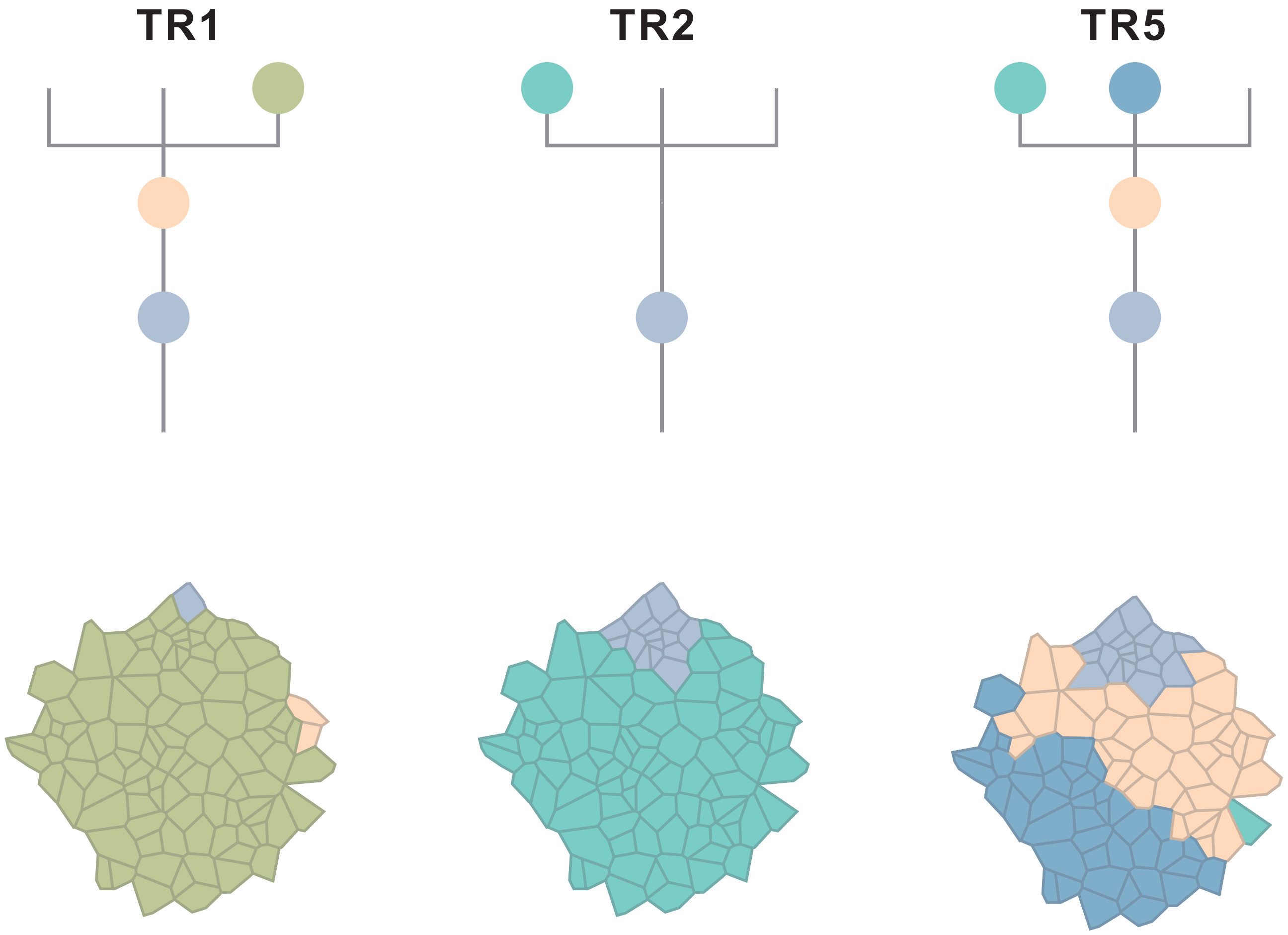

### CRC40

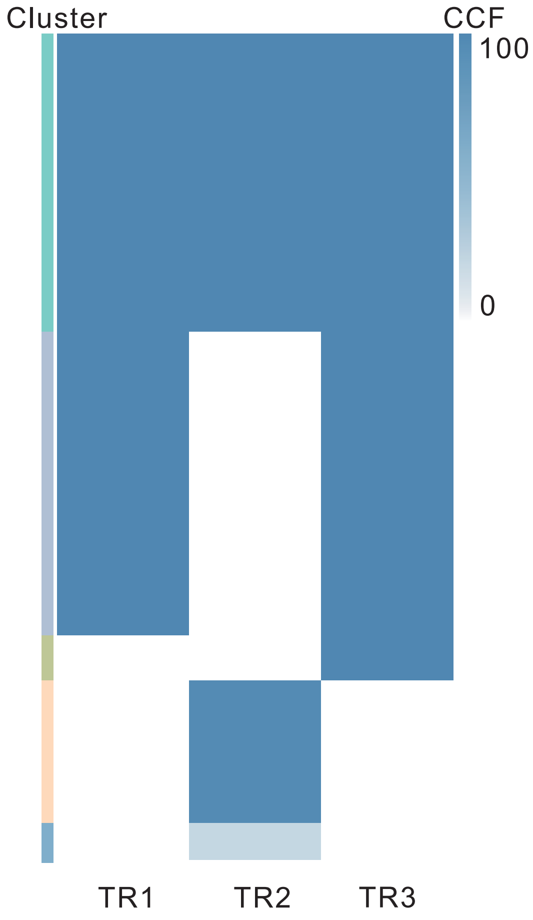

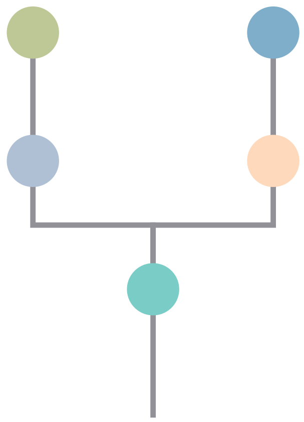

| Gene | Cytoband | Type | Cluster |
|---|---|---|---|
| *PIK3CA* | 3q26.32 | SNV | 1 |
| *TCF7L2* | 10q25.3 | SNV | 1 |
| *LZTR1* | 22q11.21 | InDel | 1 |
| * *APC* | 5q22.2 | SNV | 2 |
| * *APC* | 5q22.2 | SNV | 2 |
| *KRAS* | 12p12.1 | SNV | 2 |
| *ARID1A* | 1p36.11 | InDel | 2 |
| *ELF3* | 1q32.1 | InDel | 2 |
| *KMT2C* | 7q36.1 | SNV | 3 |
| *NDRG1* | 8q24.22 | SNV | 3 |
| *GNAS* | 20q13.32 | SNV | 3 |
| *MED12* | Xq13.1 | SNV | NA |
| *RECQL4* | 8q24.3 | Amp | NA |
| *RUNX1* | 21q22.12 | Del | NA |
| *SMARCB1* | 22q11.23 | Del | NA |
| *CHEK2* | 22q12.1 | Del | NA |
| *ZNRF3* | 22q12.1 | Del | NA |
| *NF2* | 22q12.2 | Del | NA |
| *MYH9* | 22q12.3 | Del | NA |
| *APOBEC3B* | 22q13.1 | Del | NA |
| *EP300* | 22q13.2 | Del | NA |
| *MKL1* | 22q13.2 | Del | NA |

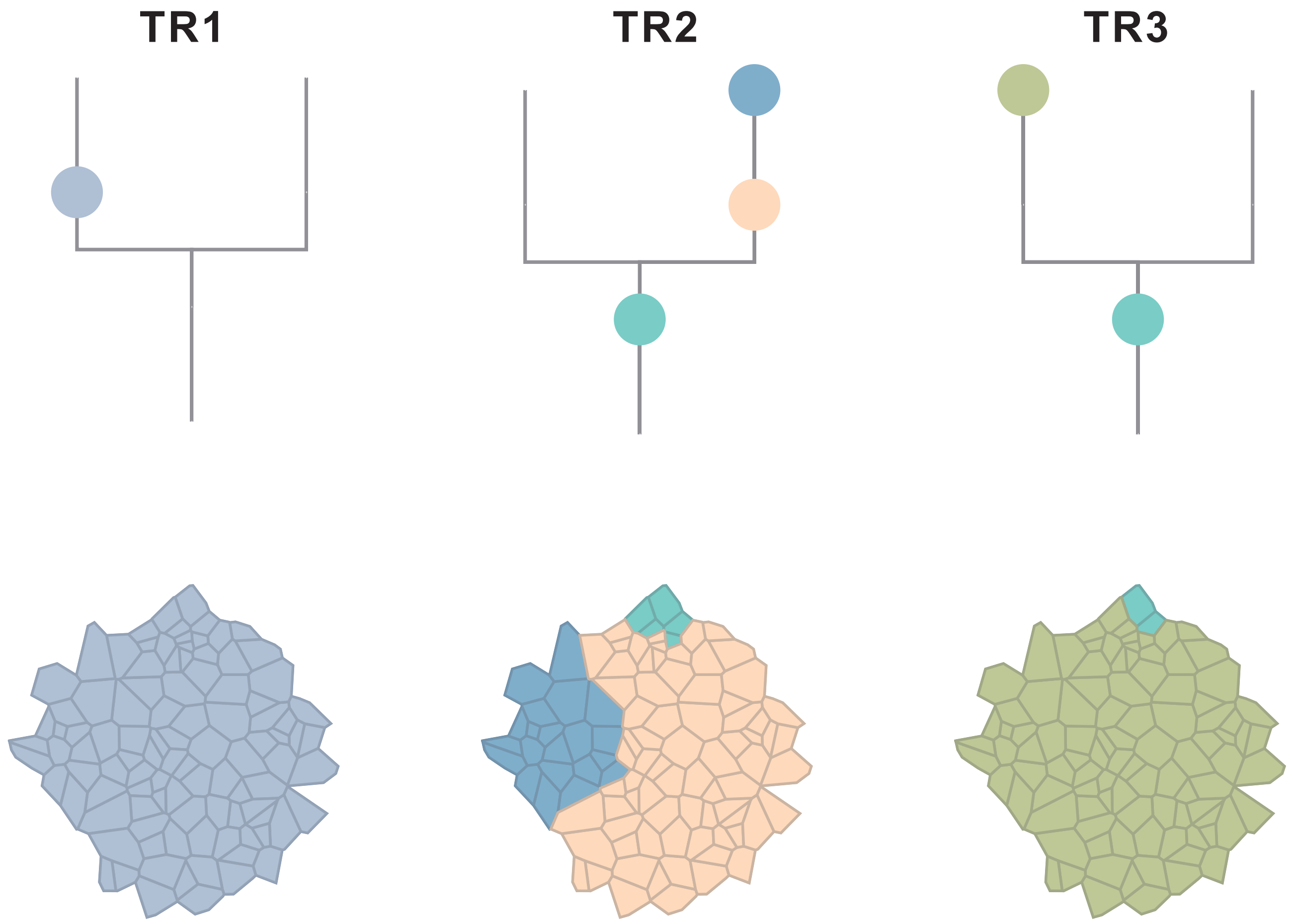

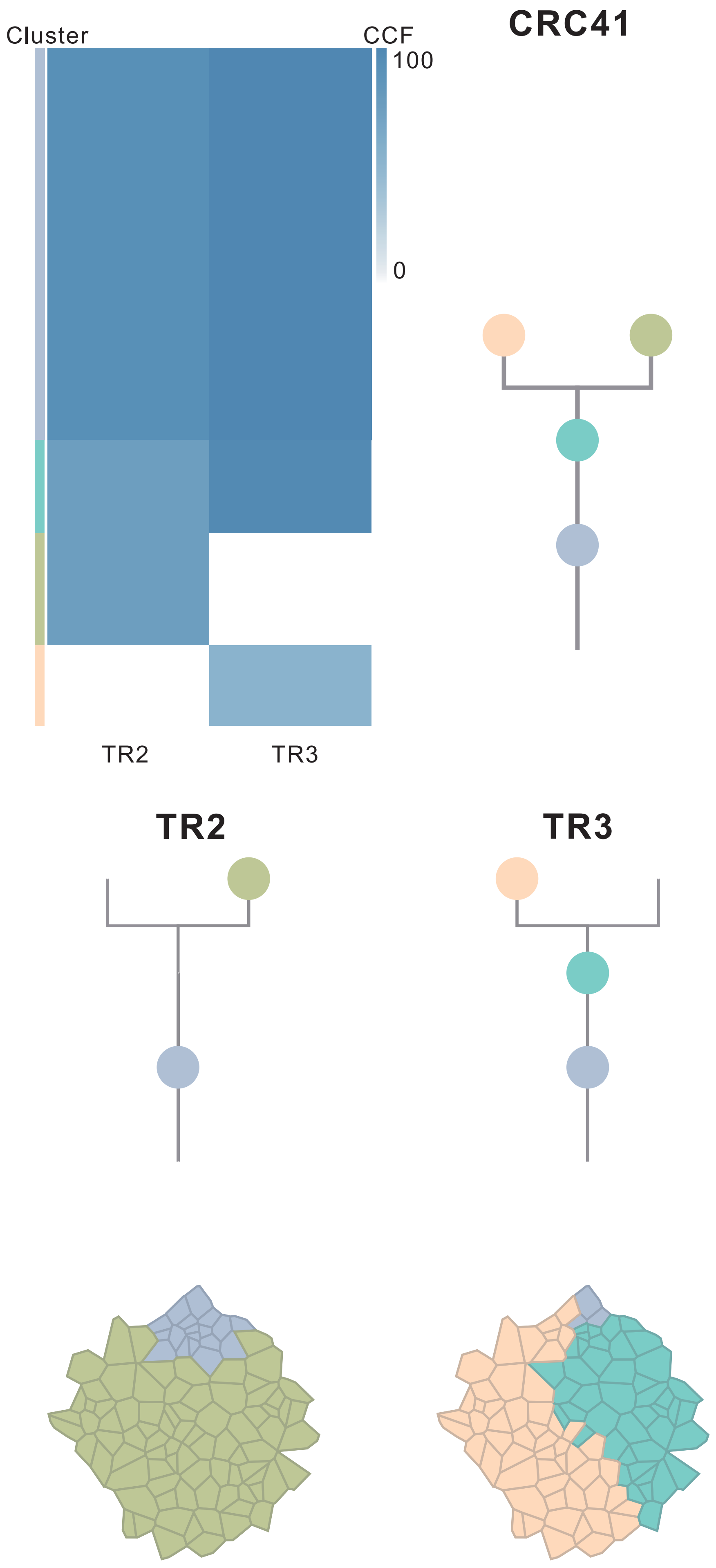

| Gene | Cytoband | Type | Cluster |
|---|---|---|---|
| *TCF7L2* | 10q25.3 | SNV | 1 |
| *TP53* | 17p13.1 | SNV | 1 |
| *FBXW7* | 4q31.3 | InDel | 1 |
| *LZTR1* | 22q11.21 | InDel | 1 |
| *IKBKB* | 8p11.21 | Amp | 1 |
| *KAT6A* | 8p11.21 | Amp | 1 |
| *FGFR1* | 8p11.23 | Amp | 1 |
| *ARID1A* | 1p36.11 | SNV | 2 |
| *SRC* | 20q11.23 | Amp | 3 |
| *MAFB* | 20q12 | Amp | 3 |
| *PLCG1* | 20q12 | Amp | 3 |
| *NFATC2* | 20q13.2 | Amp | 3 |
| *SALL4* | 20q13.2 | Amp | 3 |
| *GNAS* | 20q13.32 | Amp | 3 |
| *PTK6* | 20q13.33 | Amp | 3 |
| *RUNX1* | 21q22.12 | Del | 4 |
| *SMARCB1* | 22q11.23 | Del | 4 |
| *CHEK2* | 22q12.1 | Del | 4 |
| *ZNRF3* | 22q12.1 | Del | 4 |
| *NF2* | 22q12.2 | Del | 4 |
| *MYH9* | 22q12.3 | Del | 4 |
| *APOBEC3B* | 22q13.1 | Del | 4 |
| *EP300* | 22q13.2 | Del | 4 |
| *MKL1* | 22q13.2 | Del | 4 |
| *KDM5C* | Xp11.22 | Del | 4 |
| *SMC1A* | Xp11.22 | Del | 4 |
| *GATA1* | Xp11.23 | Del | 4 |
| *RBM10* | Xp11.23 | Del | 4 |
| *KDM6A* | Xp11.3 | Del | 4 |
| *BCOR* | Xp11.4 | Del | 4 |
| *DDX3X* | Xp11.4 | Del | 4 |
| *ZRSR2* | Xp22.2 | Del | 4 |
| *FOXO4* | Xq13.1 | Del | 4 |
| *MED12* | Xq13.1 | Del | 4 |
| *ZMYM3* | Xq13.1 | Del | 4 |
| *ATRX* | Xq21.1 | Del | 4 |
| *BTK* | Xq22.1 | Del | 4 |
| *IRS4* | Xq22.3 | Del | 4 |
| *STAG2* | Xq25 | Del | 4 |
| *BCORL1* | Xq26.1 | Del | 4 |
| *ELF4* | Xq26.1 | Del | 4 |
| *GPC3* | Xq26.2 | Del | 4 |
| *PHF6* | Xq26.2 | Del | 4 |
| *ATP2B3* | Xq28 | Del | 4 |
| *RPL10* | Xq28 | Del | 4 |
| *AR* | Xq12 | SNV | NA |
| *FBXW7* | 4q31.3 | InDel | NA |

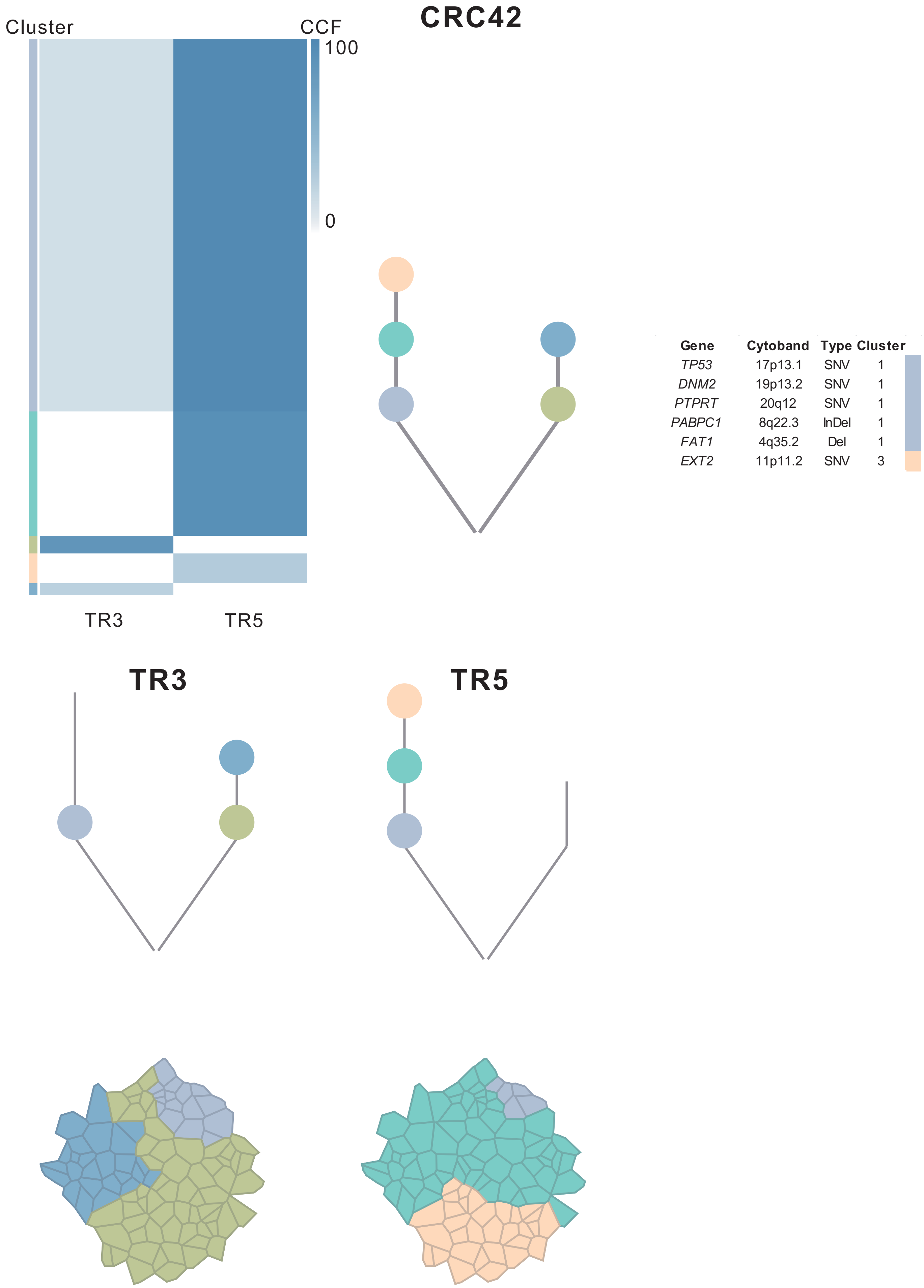
CRC42
Cluster
CCF
100
0
TR3
TR5
Gene Cytoband Type Cluster
TP53 17p13.1 SNV 1
DNM2 19p13.2 SNV 1
PTPRT 20q12 SNV 1
PABPC1 8q22.3 InDel 1
FAT1 4q35.2 Del 1
EXT2 11p11.2 SNV 3
TR3
TR5

### CRC43

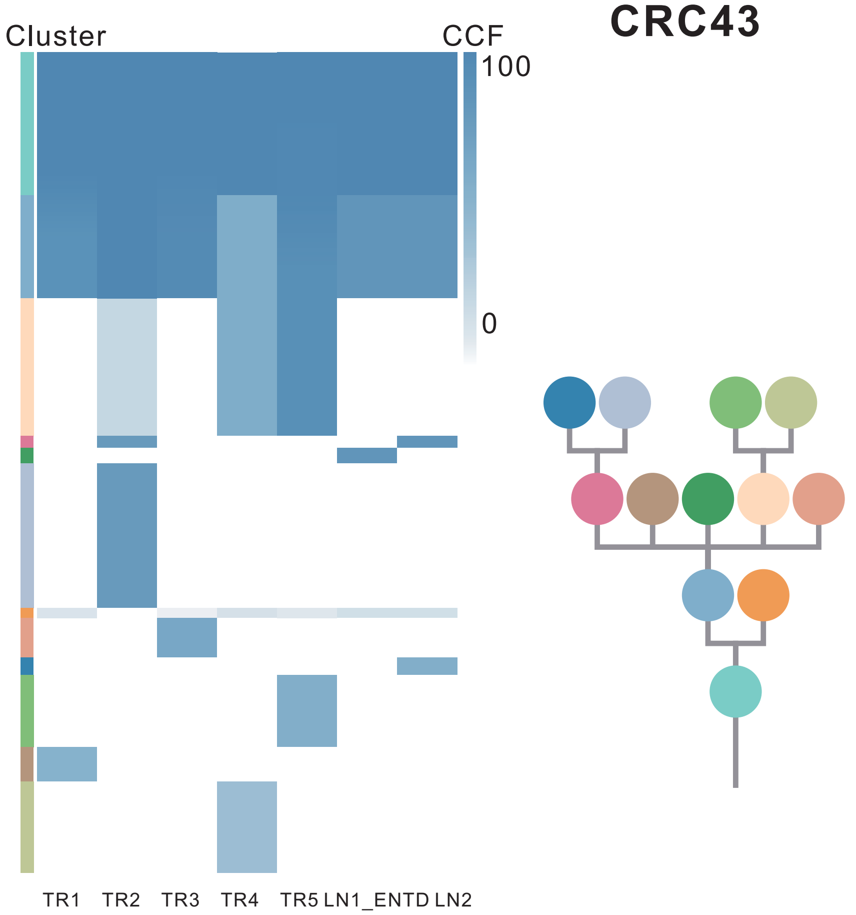

| Gene | Cytoband | Type | Cluster |
|---|---|---|---|
| *LRP1B* | 2q22.1 | SNV | 1 |
| *IRF4* | 6p25.3 | SNV | 1 |
| *FBXW7* | 4q31.3 | SNV | 2 |
| *APC* | 5q22.2 | SNV | 2 |
| *BRAF* | 7q34 | SNV | 2 |
| *CNTNAP2* | 7q35 | SNV | 2 |
| *BCL9L* | 11q23.3 | SNV | 2 |
| *KRAS* | 12p12.1 | SNV | 2 |
| *ZNF521* | 18q11.2 | SNV | 2 |
| *ELF3* | 1q32.1 | SNV | 5 |
| *TP53* | 17p13.1 | SNV | 5 |
| *SMAD4* | 18q21.2 | SNV | 5 |
| *WNK2* | 9q22.31 | SNV | 7 |
| *FOXP1* | 3p13 | Amp | 8 |
| *MITF* | 3p13 | Amp | 8 |

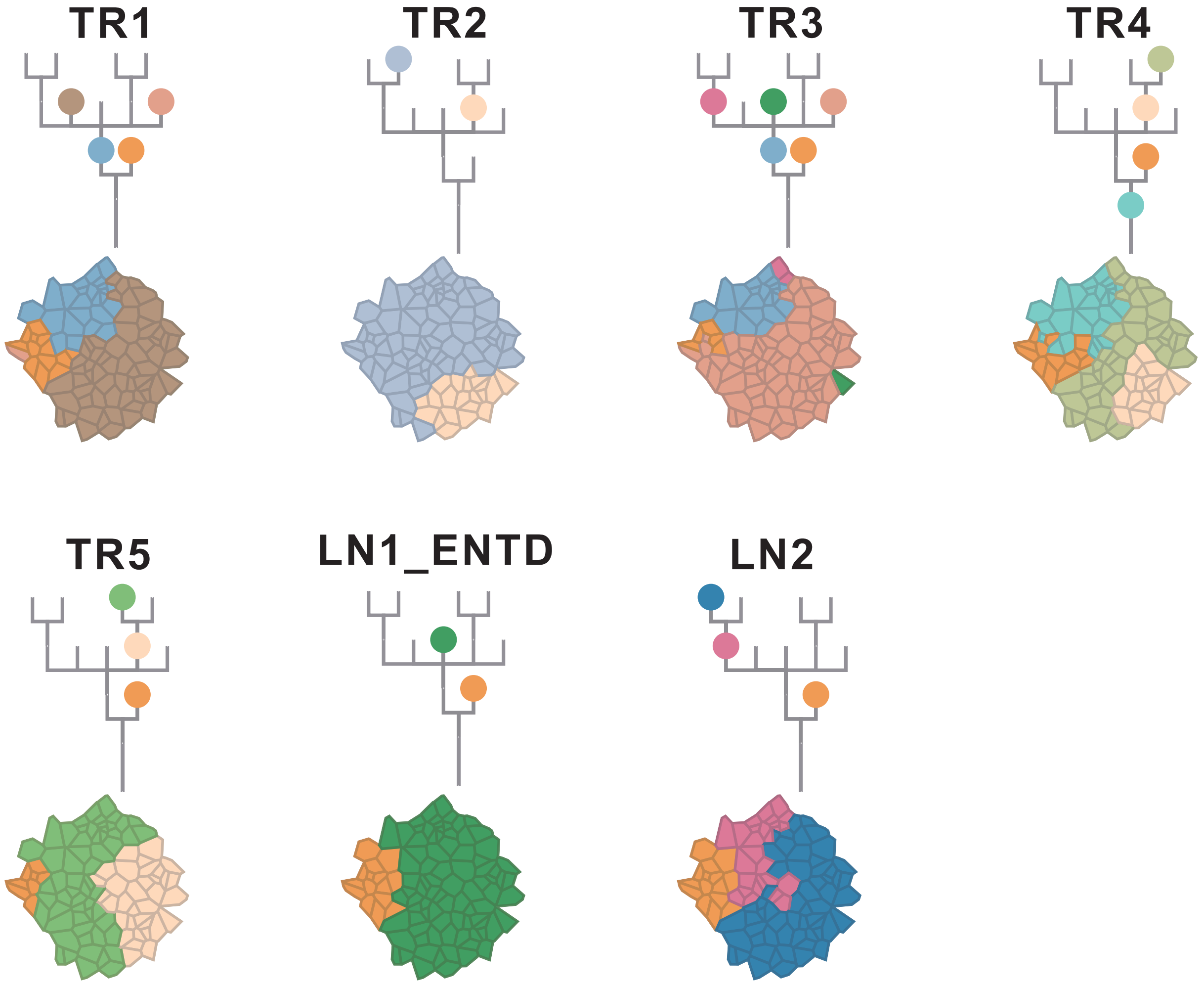

### CRC44

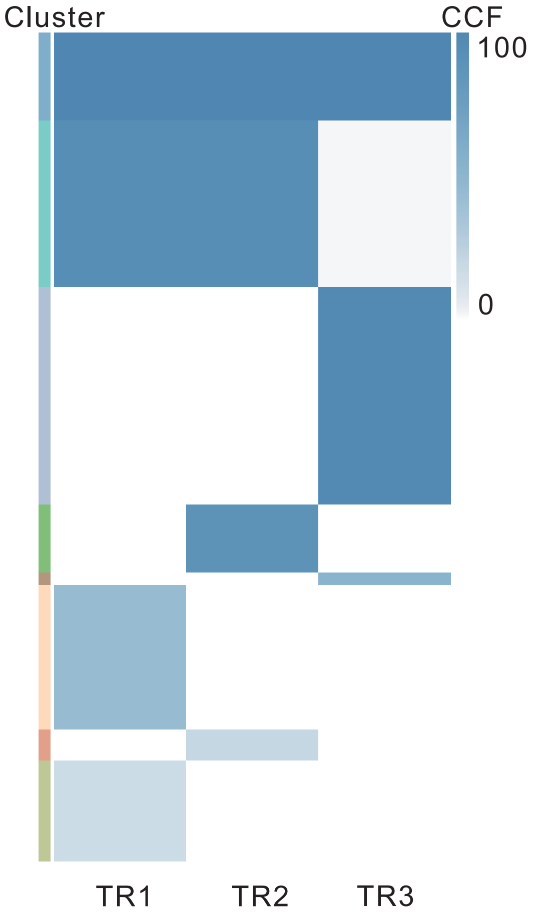

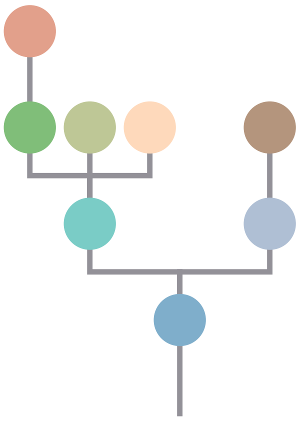

| Gene | Cytoband | Type | Cluster |
|---|---|---|---|
| * *FBXW7* | 4q31.3 | SNV | 1 |
| *TP53* | 17p13.1 | SNV | 1 |
| *FLT3* | 13q12.2 | Amp | 1 |
| *FOXO1* | 13q14.11 | Amp | 1 |
| *CYSLTR2* | 13q14.2 | Amp | 1 |
| *ARID1A* | 1p36.11 | SNV | 2 |
| * *FBXW7* | 4q31.3 | SNV | 2 |
| *KRAS* | 12p12.1 | SNV | 2 |
| *GNAS* | 20q13.32 | SNV | 4 |
| *MUTYH* | 1p34.1 | SNV | 5 |
| *APC* | 5q22.2 | SNV | 5 |
| *ACVR2A* | 2q22.3 | SNV | 7 |
| *LZTR1* | 22q11.21 | InDel | NA |
| *AMER1* | Xq11.2 | SNV | NA |

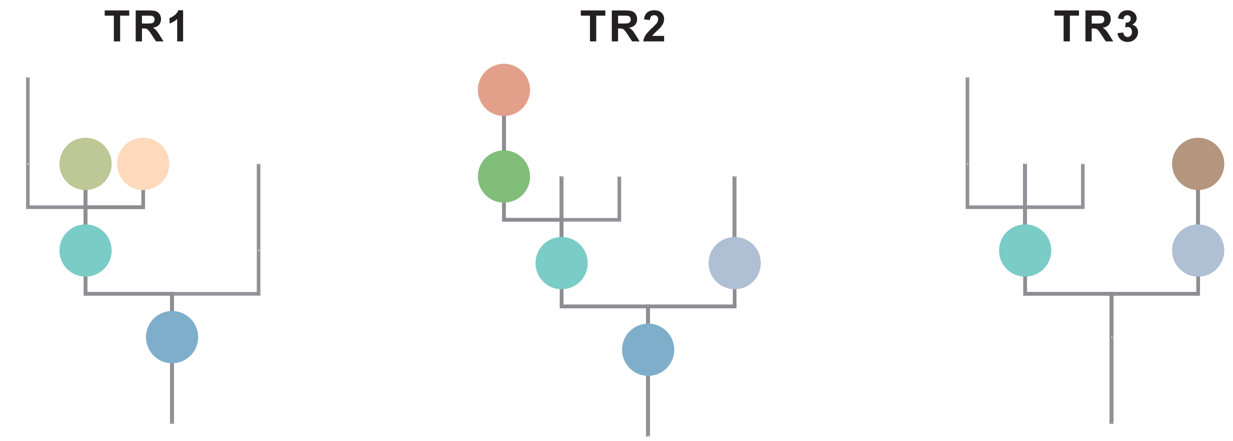

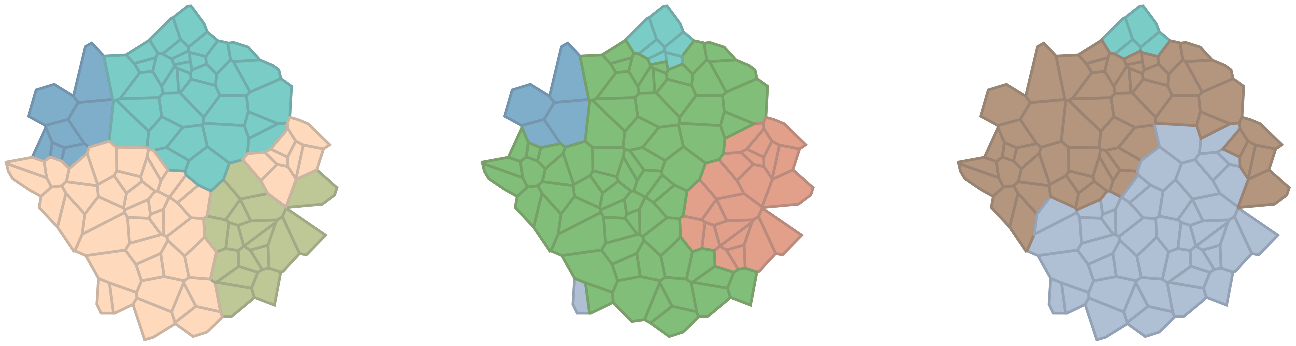

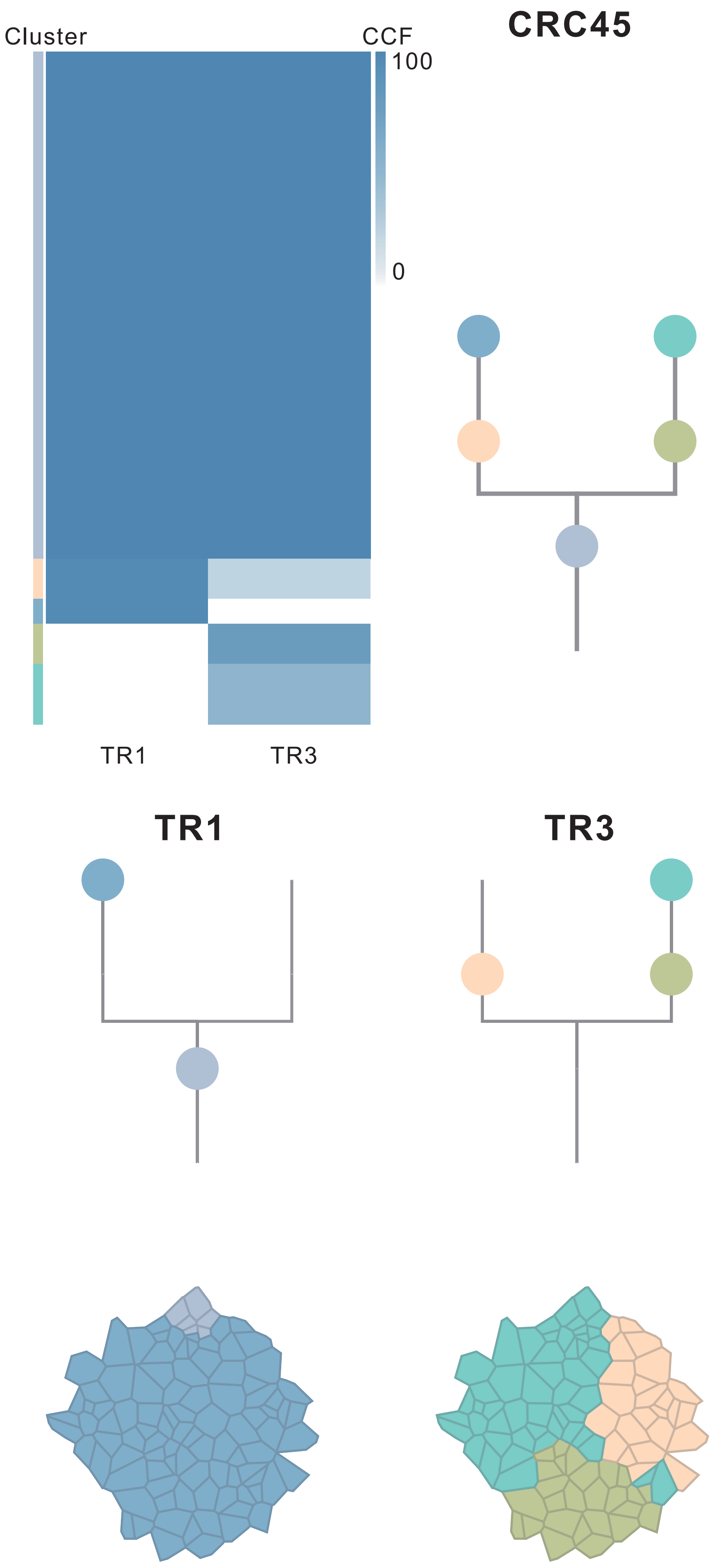

| Gene | Cytoband | Type | Cluster |
| --- | --- | --- | --- |
| *APC* | 5q22.2 | SNV | 1 |
| *POT1* | 7q31.33 | SNV | 1 |
| *TP53* | 17p13.1 | SNV | 1 |
| *FGFR1* | 8p11.23 | Amp | 4 |

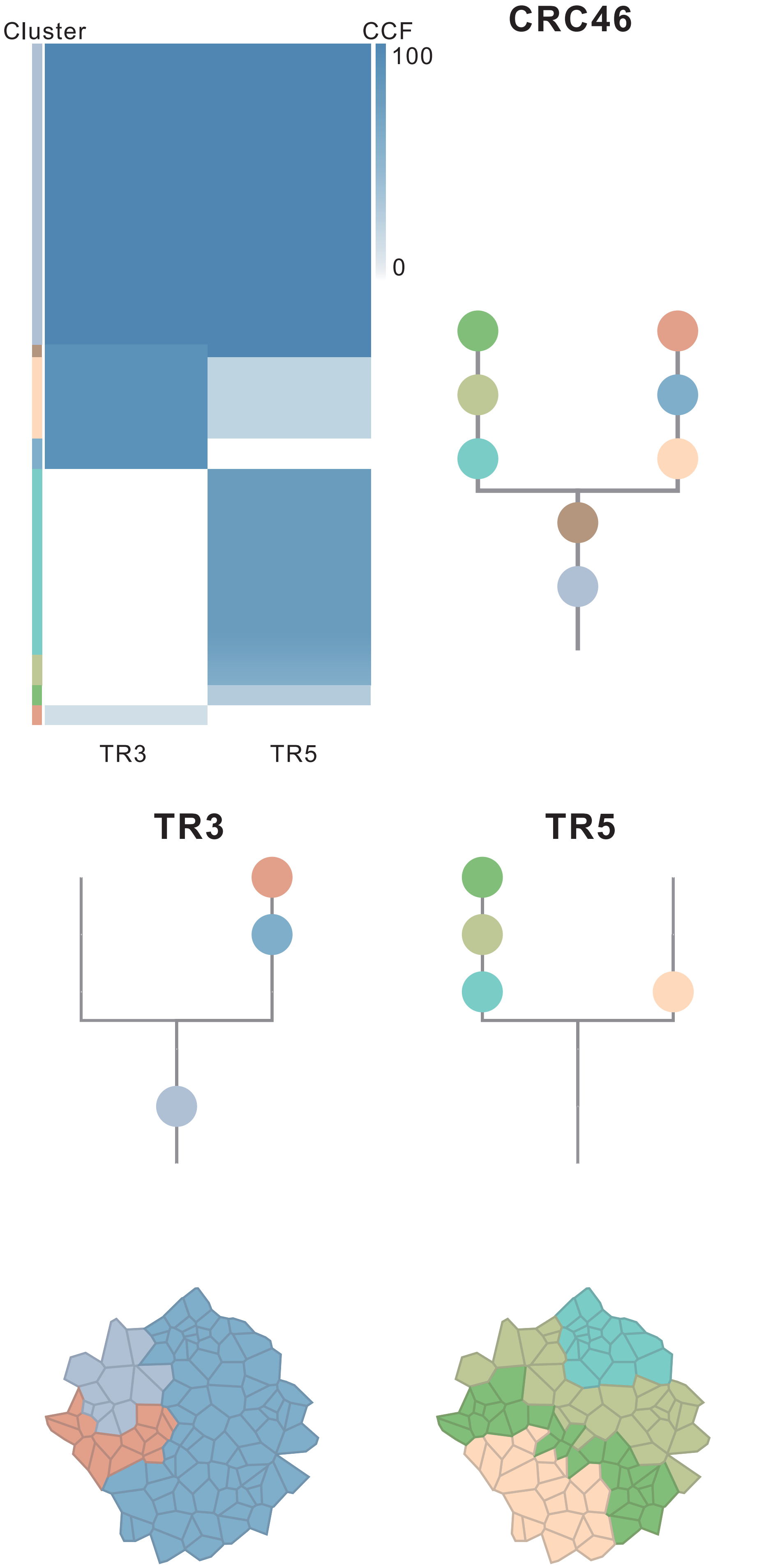

| Gene | Cytoband | Type | Cluster |
|---|---|---|---|
| *APC* | 5q22.2 | SNV | 1 |
| * *ATM* | 11q22.3 | SNV | 1 |
| *TBX3* | 12q24.21 | SNV | 1 |
| *ARID1A* | 1p36.11 | InDel | 1 |
| *NFE2L2* | 2q31.2 | InDel | 1 |
| *DDB2* | 11p11.2 | InDel | 1 |
| * *ATM* | 11q22.3 | InDel | 1 |
| *PLAG1* | 8q12.1 | Amp | 1 |
| *PREX2* | 8q13.2 | Amp | 1 |
| *NCOA2* | 8q13.3 | Amp | 1 |
| *HEY1* | 8q21.13 | Amp | 1 |
| *RUNX1T1* | 8q21.3 | Amp | 1 |
| *CDH17* | 8q22.1 | Amp | 1 |
| *PABPC1* | 8q22.3 | Amp | 1 |
| *UBR5* | 8q22.3 | Amp | 1 |
| *RAD21* | 8q24.11 | Amp | 1 |
| *MYC* | 8q24.21 | Amp | 1 |
| *RECQL4* | 8q24.3 | Amp | 1 |
| *FLT3* | 13q12.2 | Amp | 1 |
| *IRS4* | Xq22.3 | Amp | 1 |
| *BCORL1* | Xq26.1 | Amp | 1 |
| *ELF4* | Xq26.1 | Amp | 1 |
| *GPC3* | Xq26.2 | Amp | 1 |
| *EGFR* | 7p11.2 | Amp | 2 |
| *HNRNPA2B1* | 7p15.2 | Amp | 2 |
| *HOXA11* | 7p15.2 | Amp | 2 |
| *HOXA13* | 7p15.2 | Amp | 2 |
| *HOXA9* | 7p15.2 | Amp | 2 |
| *MACC1* | 7p21.1 | Amp | 2 |
| *ETV1* | 7p21.2 | Amp | 2 |
| *RAC1* | 7p22.1 | Amp | 2 |
| *CARD11* | 7p22.2 | Amp | 2 |
| *SSX2* | Xp11.22 | Amp | 2 |
| *ARAF* | Xp11.23 | Amp | 2 |
| *GATA1* | Xp11.23 | Amp | 2 |
| *SSX1* | Xp11.23 | Amp | 2 |
| *SSX4* | Xp11.23 | Amp | 2 |
| *TFE3* | Xp11.23 | Amp | 2 |
| *WAS* | Xp11.23 | Amp | 2 |
| *KDM6A* | Xp11.3 | Amp | 2 |
| *LRP1B* | 2q22.1 | SNV | 3 |
| *FOXO1* | 13q14.11 | Amp | 3 |
| *NTRK3* | 15q25.3 | SNV | 8 |
| *CYSLTR2* | 13q14.2 | Amp | NA |

### CRC47

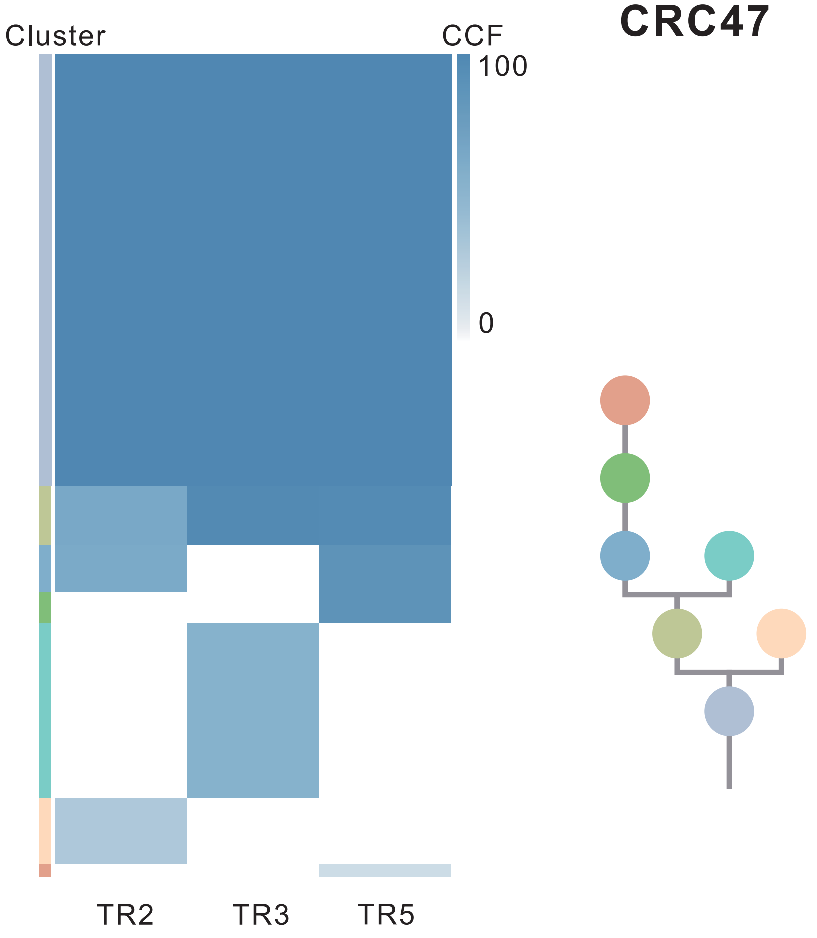

| Gene | Cytoband | Type | Cluster |
|---|---|---|---|
| *APC* | 5q22.2 | SNV | 1 |
| *TP53* | 17p13.1 | SNV | 1 |
| *MAFB* | 20q12 | Amp | 1 |
| *PLCG1* | 20q12 | Amp | 1 |
| *NFATC2* | 20q13.2 | Amp | 1 |
| *SALL4* | 20q13.2 | Amp | 1 |
| *GNAS* | 20q13.32 | Amp | 1 |
| *PTK6* | 20q13.33 | Amp | 1 |
| *SRC* | 20q11.23 | Amp | 2 |
| *PIK3CA* | 3q26.32 | SNV | 5 |
| *RBM10* | Xp11.23 | InDel | NA |

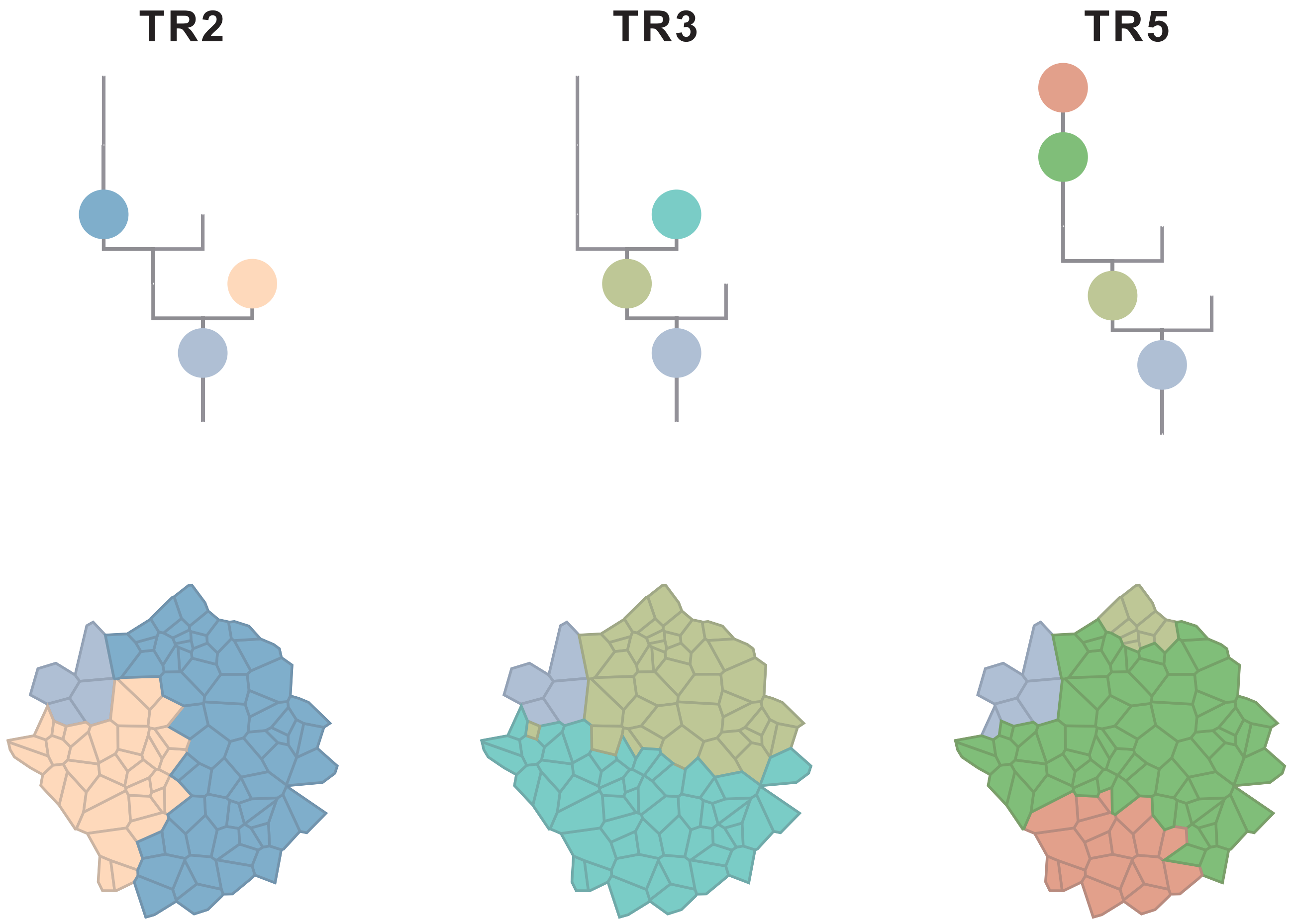

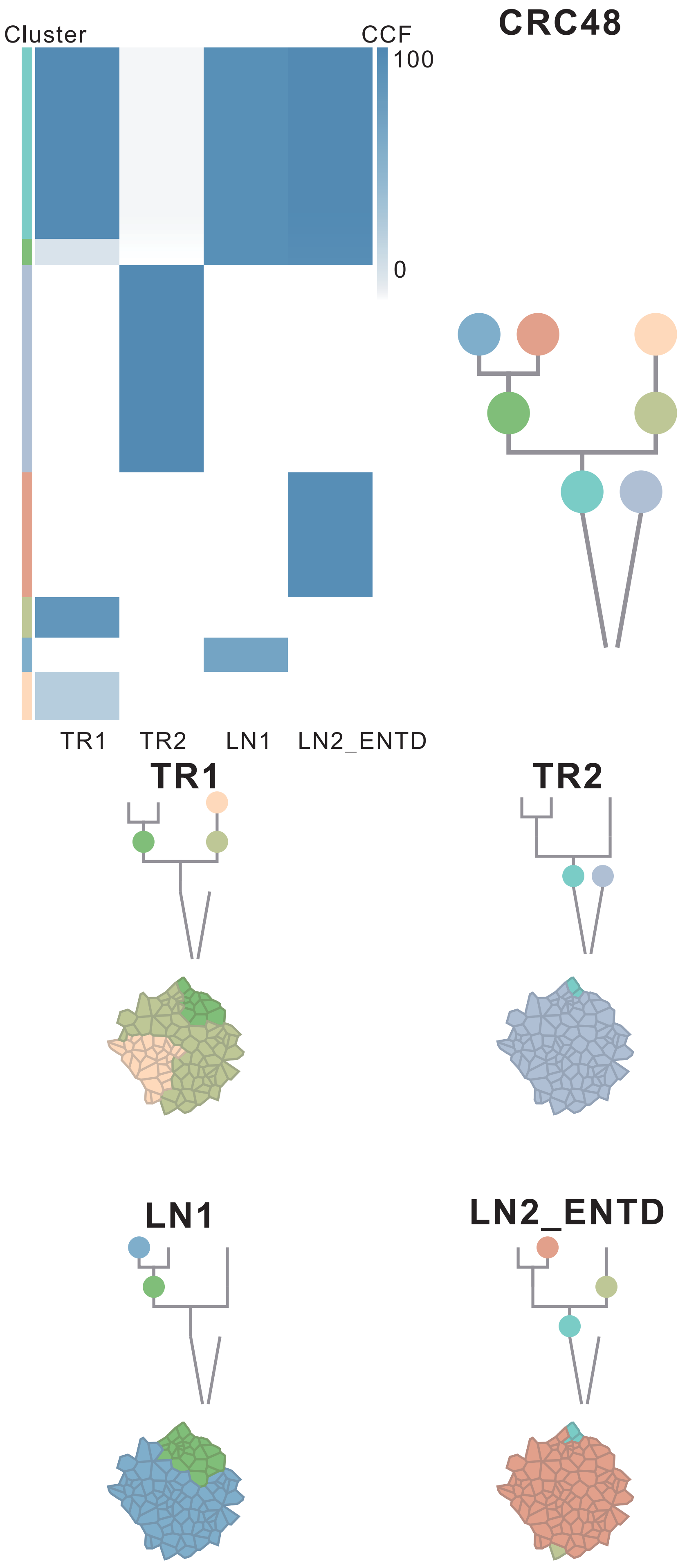

| Gene | Cytoband | Type | Cluster |
|---|---|---|---|
| * *APC* | 5q22.2 | InDel | 2 |
| * *APC* | 5q22.2 | InDel | 2 |
| *FLT3* | 13q12.2 | Amp | 2 |
| *FOXO1* | 13q14.11 | Amp | 2 |
| *CYSLTR2* | 13q14.2 | Amp | 2 |
| *PTPRD* | 9p24.1 | SNV | 6 |
| *RAD21* | 8q24.11 | Amp | 6 |
| *MYC* | 8q24.21 | Amp | 6 |
| *MUC4* | 3q29 | Amp | 7 |
| *PLAG1* | 8q12.1 | Amp | 7 |
| *PREX2* | 8q13.2 | Amp | 7 |
| *NCOA2* | 8q13.3 | Amp | 7 |
| *RUNX1T1* | 8q21.3 | Amp | 7 |
| *HEY1* | 8q21.13 | Amp | 7 |
| *CDH17* | 8q22.1 | Amp | 7 |
| *PABPC1* | 8q22.3 | Amp | 7 |
| *UBR5* | 8q22.3 | Amp | 7 |
| *CASP9* | 1p36.21 | Del | 7 |
| *PRDM2* | 1p36.21 | Del | 7 |
| *SPEN* | 1p36.21 | Del | 7 |
| *CAMTA1* | 1p36.31 | Del | 7 |
| *RPL22* | 1p36.31 | Del | 7 |
| *TNFRSF14* | 1p36.32 | Del | 7 |
| *SDHA* | 5p15.33 | Del | 7 |
| *FEN1* | 11q12.2 | Del | 7 |
| *SDHAF2* | 11q12.2 | Del | 7 |
| *MEN1* | 11q13.1 | Del | 7 |
| *EED* | 11q14.2 | Del | 7 |
| *BIRC3* | 11q22.2 | Del | 7 |
| *ATM* | 11q22.3 | Del | 7 |
| *DDX10* | 11q22.3 | Del | 7 |
| *SDHD* | 11q23.1 | Del | 7 |
| *ZBTB16* | 11q23.2 | Del | 7 |
| *ARHGEF12* | 11q23.3 | Del | 7 |
| *BCL9L* | 11q23.3 | Del | 7 |
| *CBL* | 11q23.3 | Del | 7 |
| *CCNB1IP1* | 14q11.2 | Del | 7 |
| *BAZ1A* | 14q13.2 | Del | 7 |
| *NKX2-1* | 14q13.3 | Del | 7 |
| *MAX* | 14q23.3 | Del | 7 |
| *RAD51B* | 14q24.1 | Del | 7 |
| *BCL11B* | 14q32.2 | Del | 7 |
| *DICER1* | 14q32.13 | Del | 7 |
| *AXIN1* | 16p13.3 | Del | 7 |
| *NTHL1* | 16p13.3 | Del | 7 |
| *TRAF7* | 16p13.3 | Del | 7 |
| *TSC2* | 16p13.3 | Del | 7 |
| *FLCN* | 17p11.2 | Del | 7 |
| *NCOR1* | 17p11.2 | Del | 7 |
| *MAP2K4* | 17p12 | Del | 7 |
| *PER1* | 17p13.1 | Del | 7 |
| *TP53* | 17p13.1 | Del | 7 |
| *YWHAE* | 17p13.3 | Del | 7 |
| *DNM2* | 19p13.2 | Del | 7 |
| *KEAP1* | 19p13.2 | Del | 7 |
| *SMARCA4* | 19p13.2 | Del | 7 |
| *STK11* | 19p13.3 | Del | 7 |
| *TCF3* | 19p13.3 | Del | 7 |
| *ELL* | 19p13.11 | Del | 7 |
| *SIRPA* | 20p13 | Del | 7 |
| *KRAS* | 12p12.1 | SNV | NA |
| *TP53* | 17p13.1 | SNV | NA |
| *SMAD4* | 18q21.2 | SNV | NA |
| *AMER1* | Xq11.2 | SNV | NA |
| *ATP2B3* | Xq28 | SNV | NA |
| *LZTR1* | 22q11.21 | InDel | NA |
| *BCORL1* | Xq26.1 | InDel | NA |

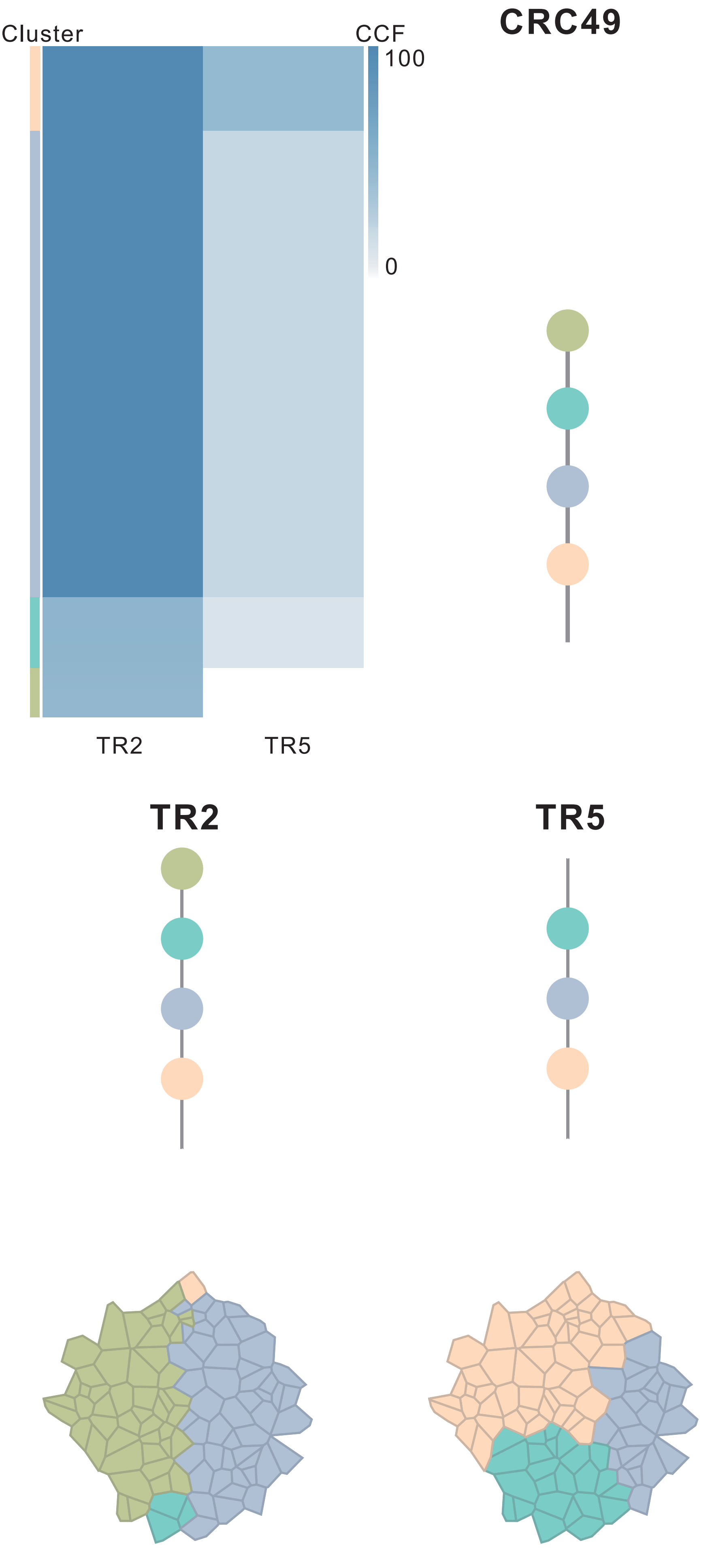

| Gene | Cytoband | Type | Cluster |
|---|---|---|---|
| * *APC* | 5q22.2 | SNV | 1 |
| * *APC* | 5q22.2 | SNV | 1 |
| *TP53* | 17p13.1 | SNV | 1 |
| *MUC4* | 3q29 | Amp | 3 |
| *RECQL4* | 8q24.3 | Amp | 3 |

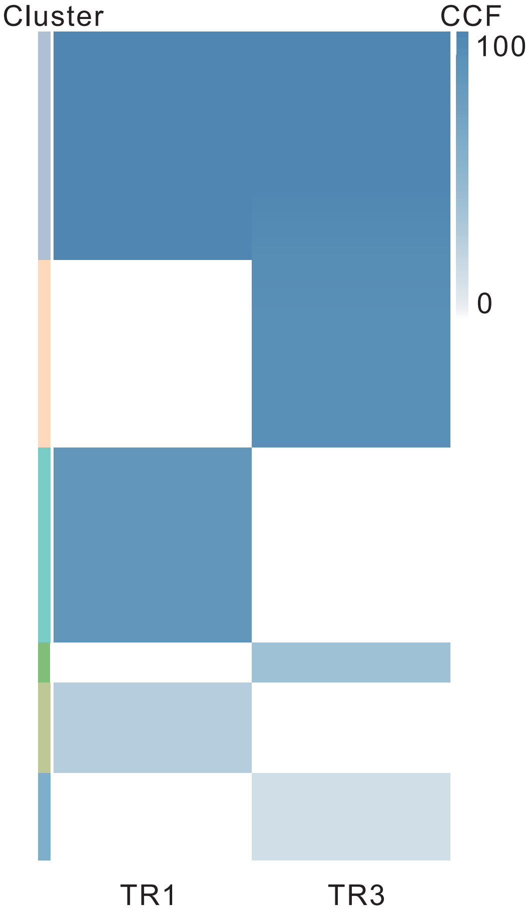

#### CRC50

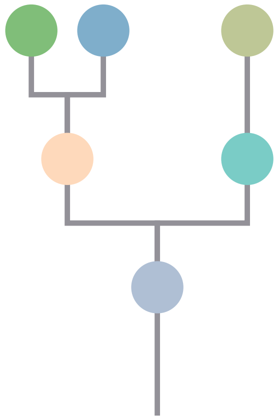

| Gene | Cytoband | Type | Cluster |
|---|---|---|---|
| *ARID1A* | 1p36.11 | SNV | 1 |
| *PIK3CA* | 3q26.32 | SNV | 1 |
| *KRAS* | 12p12.1 | SNV | 1 |
| *BAP1* | 3p21.1 | InDel | 1 |
| *APC* | 5q22.2 | InDel | 1 |
| *FBXW7* | 4q31.3 | SNV | 2 |
| *FAT4* | 4q28.1 | SNV | 3 |
| *DICER1* | 14q32.13 | InDel | 3 |
| *TSC1* | 9q34.13 | SNV | 5 |

## TR1

## TR3

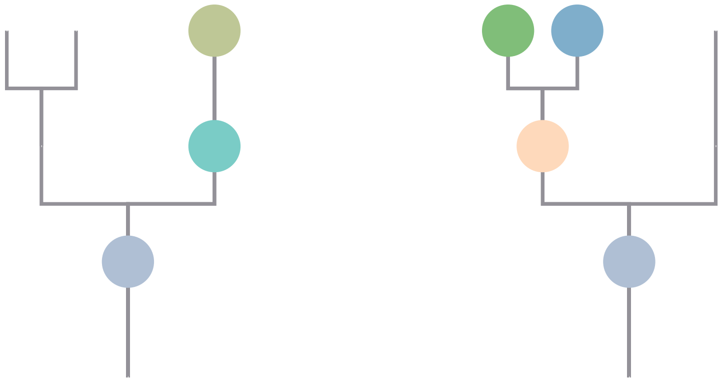

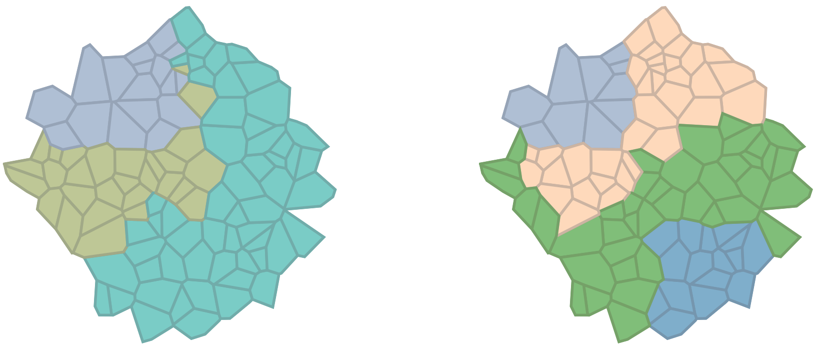

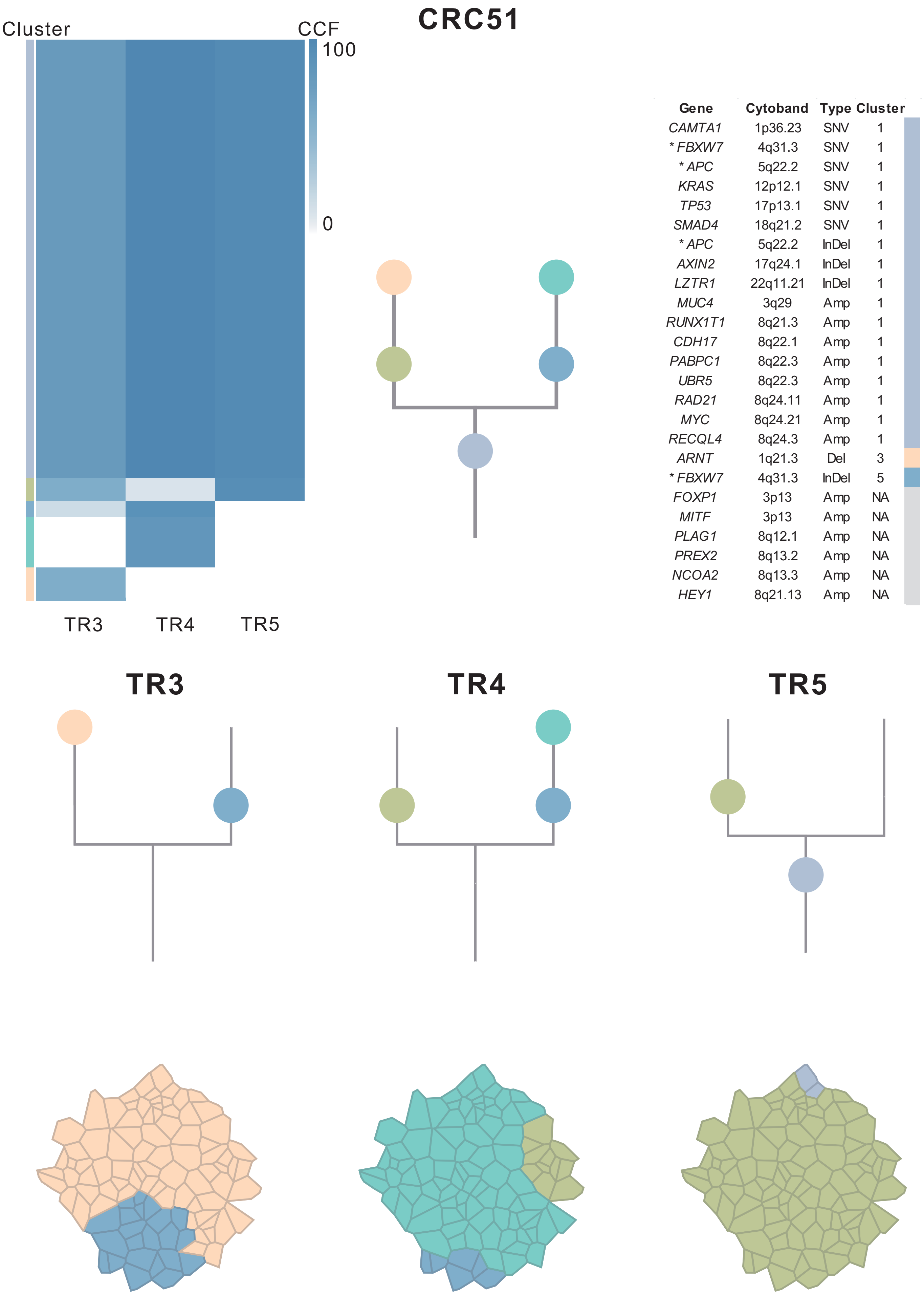

| Gene | Cytoband | Type | Cluster |
|---|---|---|---|
| *CAMTA1* | 1p36.23 | SNV | 1 |
| * *FBXW7* | 4q31.3 | SNV | 1 |
| * *APC* | 5q22.2 | SNV | 1 |
| *KRAS* | 12p12.1 | SNV | 1 |
| *TP53* | 17p13.1 | SNV | 1 |
| *SMAD4* | 18q21.2 | SNV | 1 |
| * *APC* | 5q22.2 | InDel | 1 |
| *AXIN2* | 17q24.1 | InDel | 1 |
| *LZTR1* | 22q11.21 | InDel | 1 |
| *MUC4* | 3q29 | Amp | 1 |
| *RUNX1T1* | 8q21.3 | Amp | 1 |
| *CDH17* | 8q22.1 | Amp | 1 |
| *PABPC1* | 8q22.3 | Amp | 1 |
| *UBR5* | 8q22.3 | Amp | 1 |
| *RAD21* | 8q24.11 | Amp | 1 |
| *MYC* | 8q24.21 | Amp | 1 |
| *RECQL4* | 8q24.3 | Amp | 1 |
| *ARNT* | 1q21.3 | Del | 3 |
| * *FBXW7* | 4q31.3 | InDel | 5 |
| *FOXP1* | 3p13 | Amp | NA |
| *MITF* | 3p13 | Amp | NA |
| *PLAG1* | 8q12.1 | Amp | NA |
| *PREX2* | 8q13.2 | Amp | NA |
| *NCOA2* | 8q13.3 | Amp | NA |
| *HEY1* | 8q21.13 | Amp | NA |

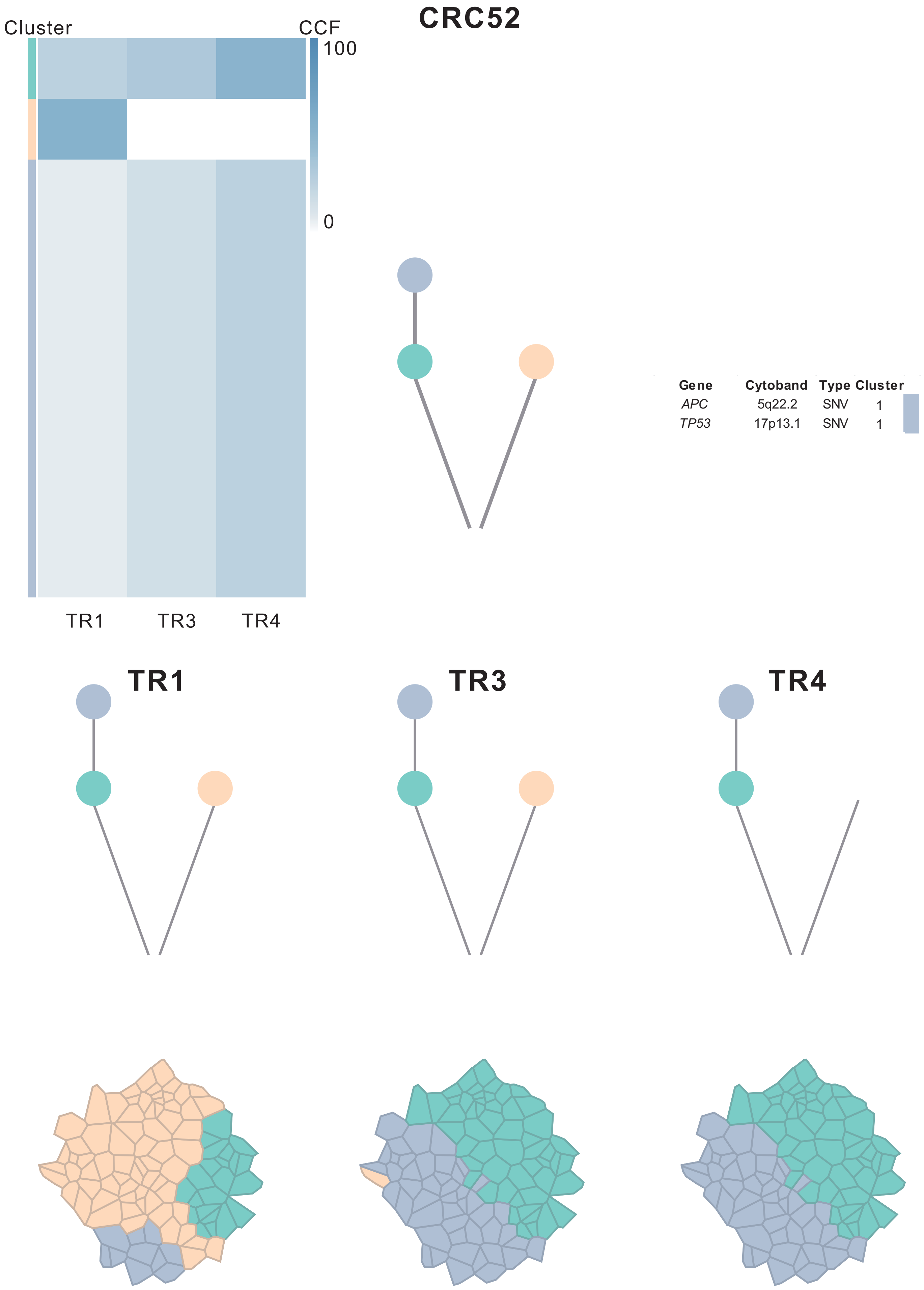
CRC52
Cluster
CCF
100
0
TR1
TR3
TR4
Gene Cytoband Type Cluster
APC 5q22.2 SNV 1
TP53 17p13.1 SNV 1
TR1
TR3
TR4

#### CRC53

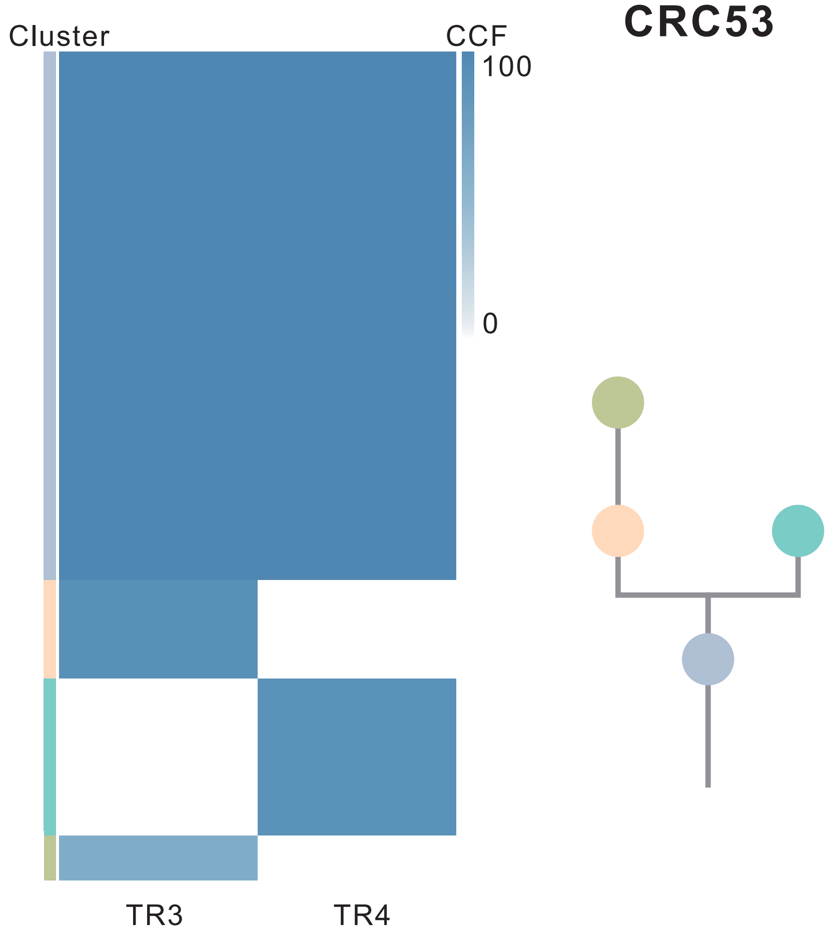

| Gene | Cytoband | Type | Cluster |
|---|---|---|---|
| *FBXW7* | 4q31.3 | SNV | 1 |
| *KRAS* | 12p12.1 | SNV | 1 |
| *TP53* | 17p13.1 | SNV | 1 |
| *SMARCA4* | 19p13.2 | SNV | 1 |
| *APC* | 5q22.2 | InDel | 1 |
| *LZTR1* | 22q11.21 | InDel | 1 |
| *KRAS* | 12p12.1 | Amp | 2 |
| *AMER1* | Xq11.2 | SNV | NA |

**TR3** **TR4**

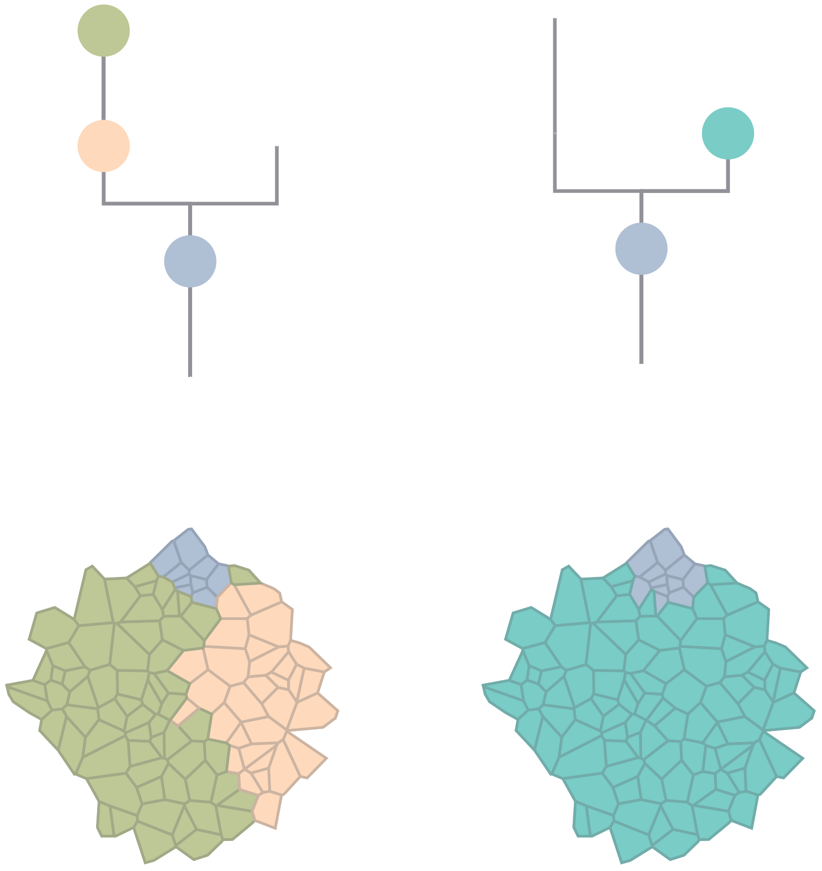

### CRC54

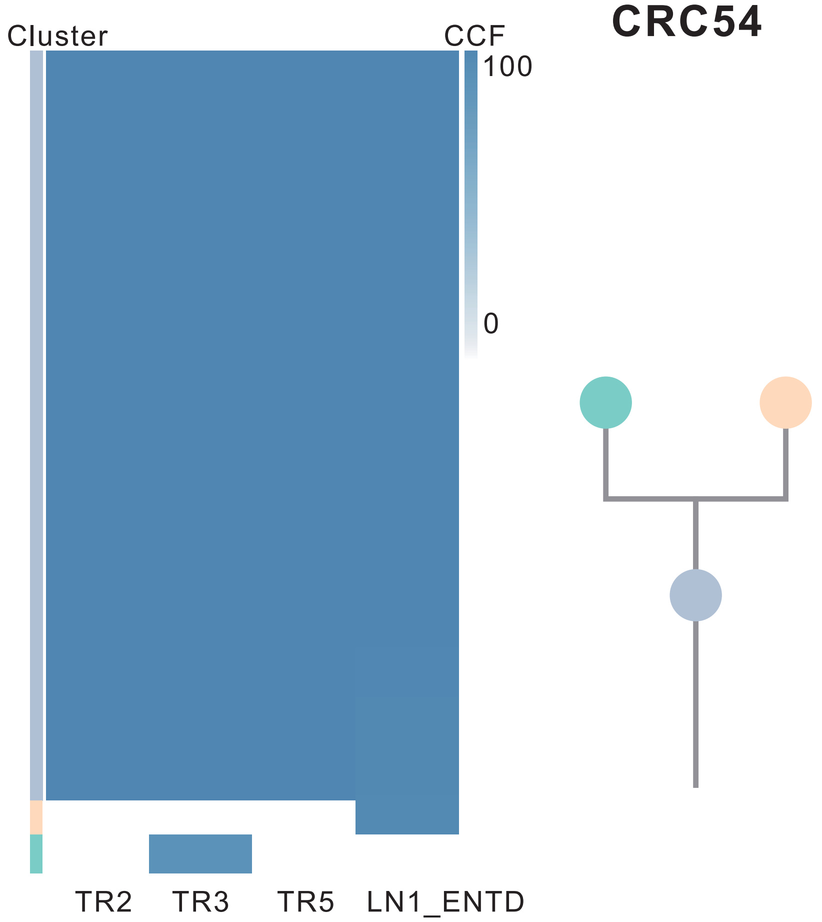

| Gene | Cytoband | Type | Cluster |
|---|---|---|---|
| * *APC* | 5q22.2 | SNV | 1 |
| *KRAS* | 12p12.1 | SNV | 1 |
| *MAX* | 14q23.3 | SNV | 1 |
| *TP53* | 17p13.1 | SNV | 1 |
| *SMAD4* | 18q21.2 | SNV | 1 |
| *LZTR1* | 22q11.21 | InDel | 1 |
| *SSX2* | Xp11.22 | Amp | 1 |
| *ARAF* | Xp11.23 | Amp | 1 |
| *GATA1* | Xp11.23 | Amp | 1 |
| *SSX1* | Xp11.23 | Amp | 1 |
| *SSX4* | Xp11.23 | Amp | 1 |
| *TFE3* | Xp11.23 | Amp | 1 |
| *WAS* | Xp11.23 | Amp | 1 |
| *KDM6A* | Xp11.3 | Amp | 1 |
| * *APC* | 5q22.2 | SNV | NA |
| *AMER1* | Xq11.2 | SNV | NA |
| *HRAS* | 11p15.5 | Amp | NA |

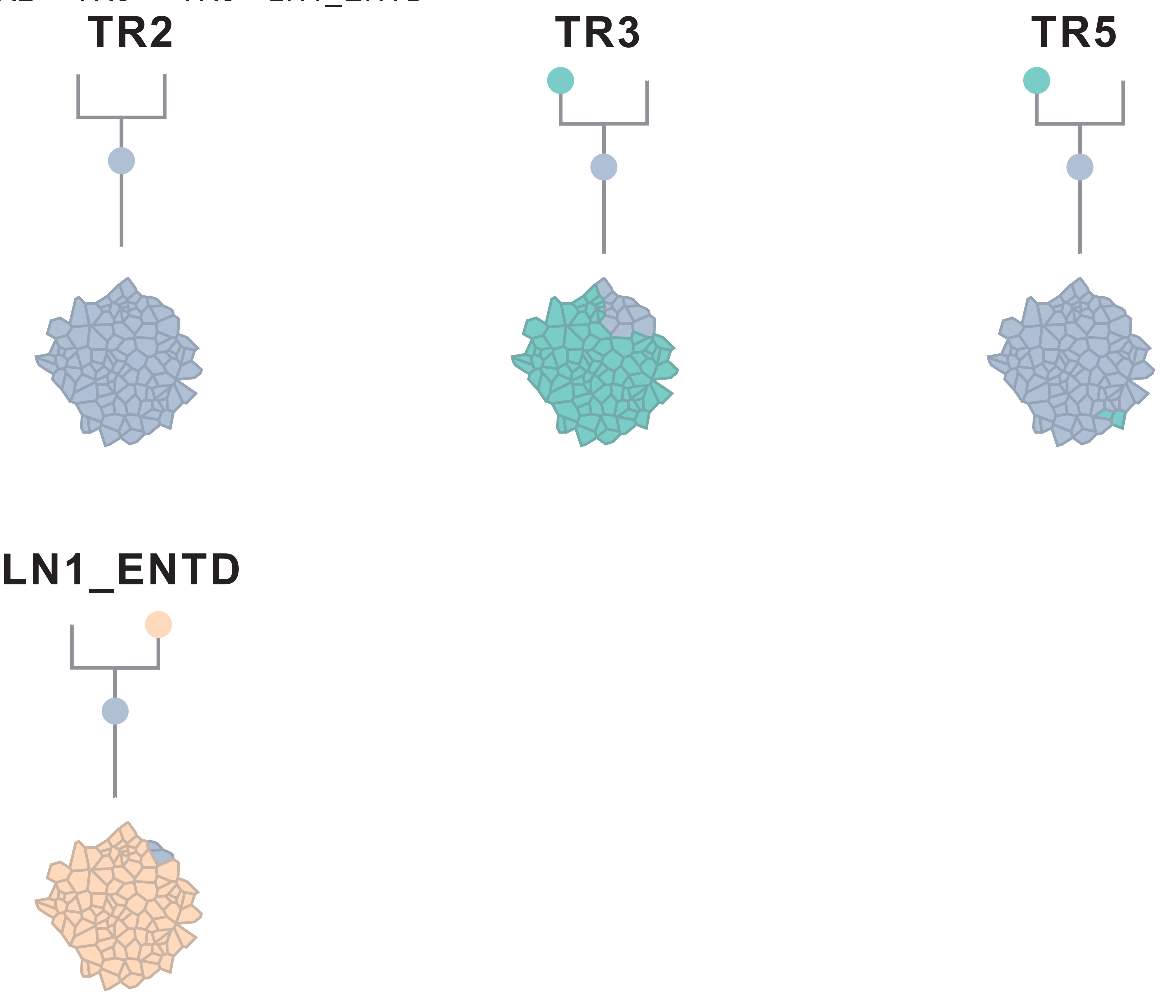

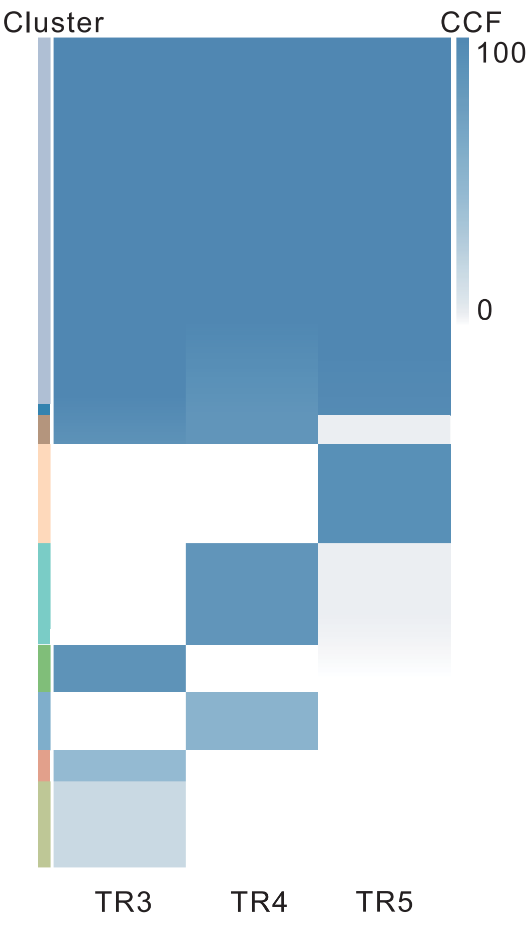

### CRC55

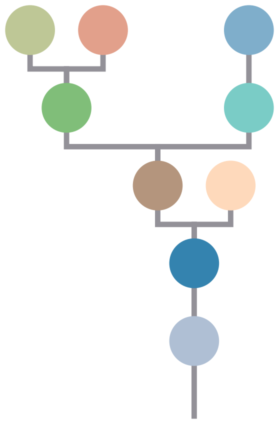

| Gene | Cytoband | Type | Cluster |
|---|---|---|---|
| *APC* | 5q22.2 | SNV | 1 |
| *CDH1* | 16q22.1 | SNV | 1 |
| *TP53* | 17p13.1 | SNV | 1 |
| *ARID1A* | 1p36.11 | InDel | 1 |
| *EGFR* | 7p11.2 | Amp | 1 |
| *HNRNPA2B1* | 7p15.2 | Amp | 1 |
| *HOXA11* | 7p15.2 | Amp | 1 |
| *HOXA13* | 7p15.2 | Amp | 1 |
| *HOXA9* | 7p15.2 | Amp | 1 |
| *MACC1* | 7p21.1 | Amp | 1 |
| *ETV1* | 7p21.2 | Amp | 1 |
| *RAC1* | 7p22.1 | Amp | 1 |
| *CARD11* | 7p22.2 | Amp | 1 |
| *FLT3* | 13q12.2 | Amp | 1 |
| *FOXO1* | 13q14.11 | Amp | 1 |
| *CYSLTR2* | 13q14.2 | Amp | 1 |
| *SRC* | 20q11.23 | Amp | 1 |
| *MUC4* | 3q29 | Amp | 2 |
| *NCOR1* | 17p12 | SNV | 3 |
| *MAFB* | 20q12 | Amp | 3 |
| *PLCG1* | 20q12 | Amp | 3 |
| *NFATC2* | 20q13.2 | Amp | 3 |
| *SALL4* | 20q13.2 | Amp | 3 |
| *GNAS* | 20q13.32 | Amp | 3 |
| *PTK6* | 20q13.33 | Amp | 3 |
| *AR* | Xq12 | SNV | NA |

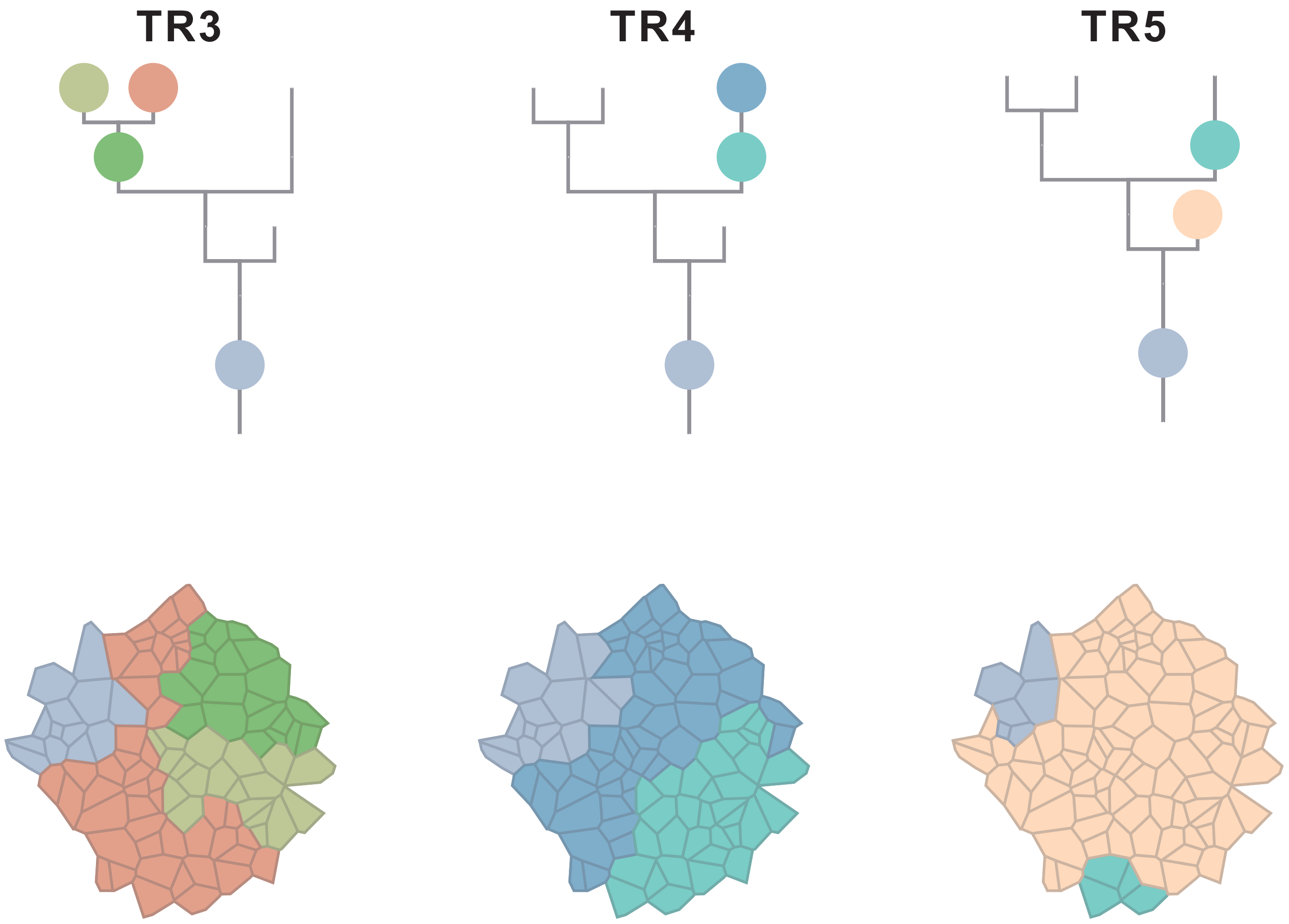

### CRC56

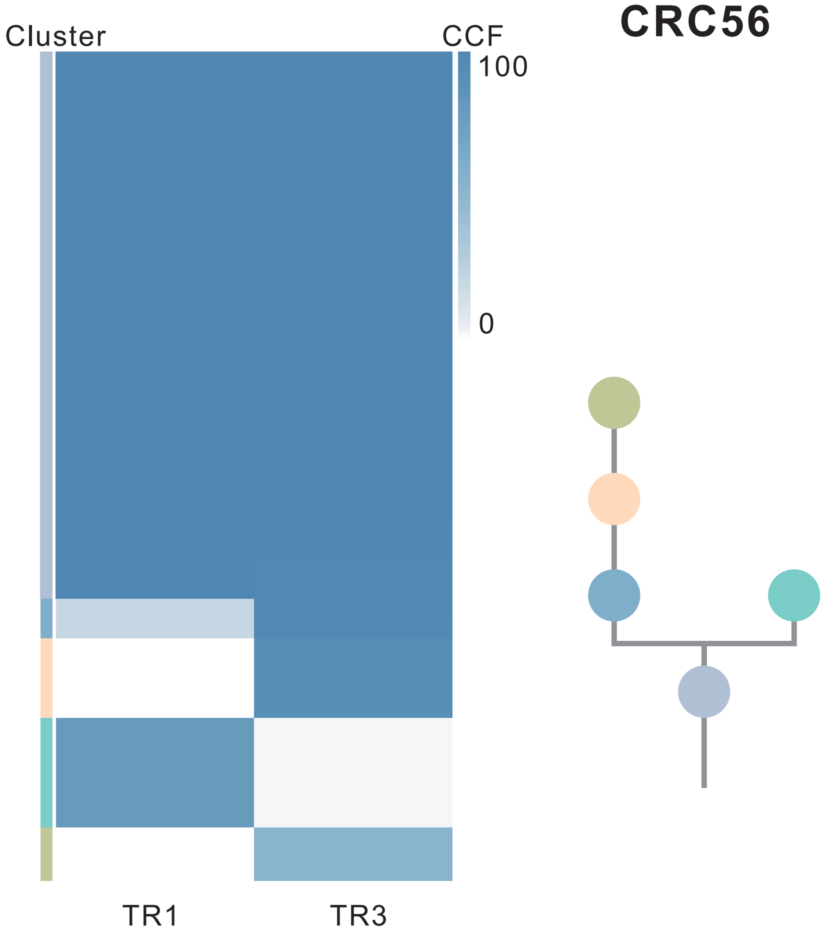

| Gene | Cytoband | Type | Cluster |
|---|---|---|---|
| *PIK3CA* | 3q26.32 | SNV | 1 |
| *KDR* | 4q12 | SNV | 1 |
| * *FBXW7* | 4q31.3 | SNV | 1 |
| * *APC* | 5q22.2 | SNV | 1 |
| *TP53* | 17p13.1 | SNV | 1 |
| *BCL9L* | 11q23.3 | InDel | 1 |
| *SRC* | 20q11.23 | Amp | 1 |
| *MAFB* | 20q12 | Amp | 1 |
| *PLCG1* | 20q12 | Amp | 1 |
| *NFATC2* | 20q13.2 | Amp | 1 |
| *SALL4* | 20q13.2 | Amp | 1 |
| *GNAS* | 20q13.32 | Amp | 1 |
| *PTK6* | 20q13.33 | Amp | 1 |
| *ARHGEF10* | 8p23.3 | Del | 1 |
| * *FBXW7* | 4q31.3 | SNV | 5 |
| * *APC* | 5q22.2 | SNV | 5 |
| *RBM10* | Xp11.23 | SNV | NA |

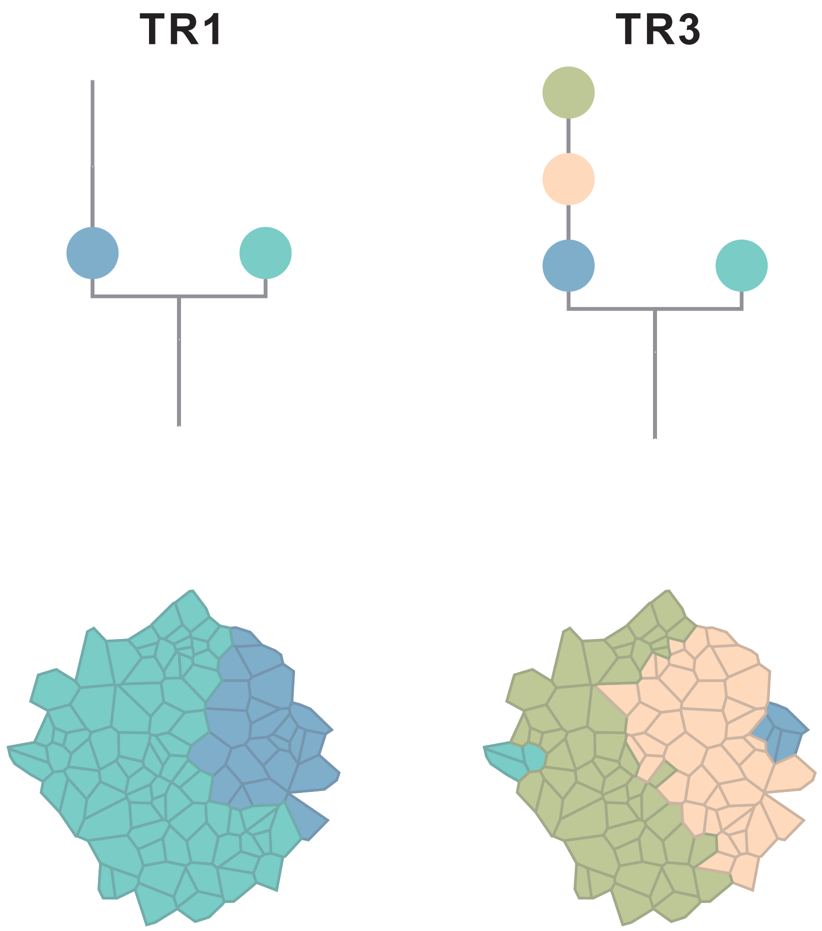

### CRC57

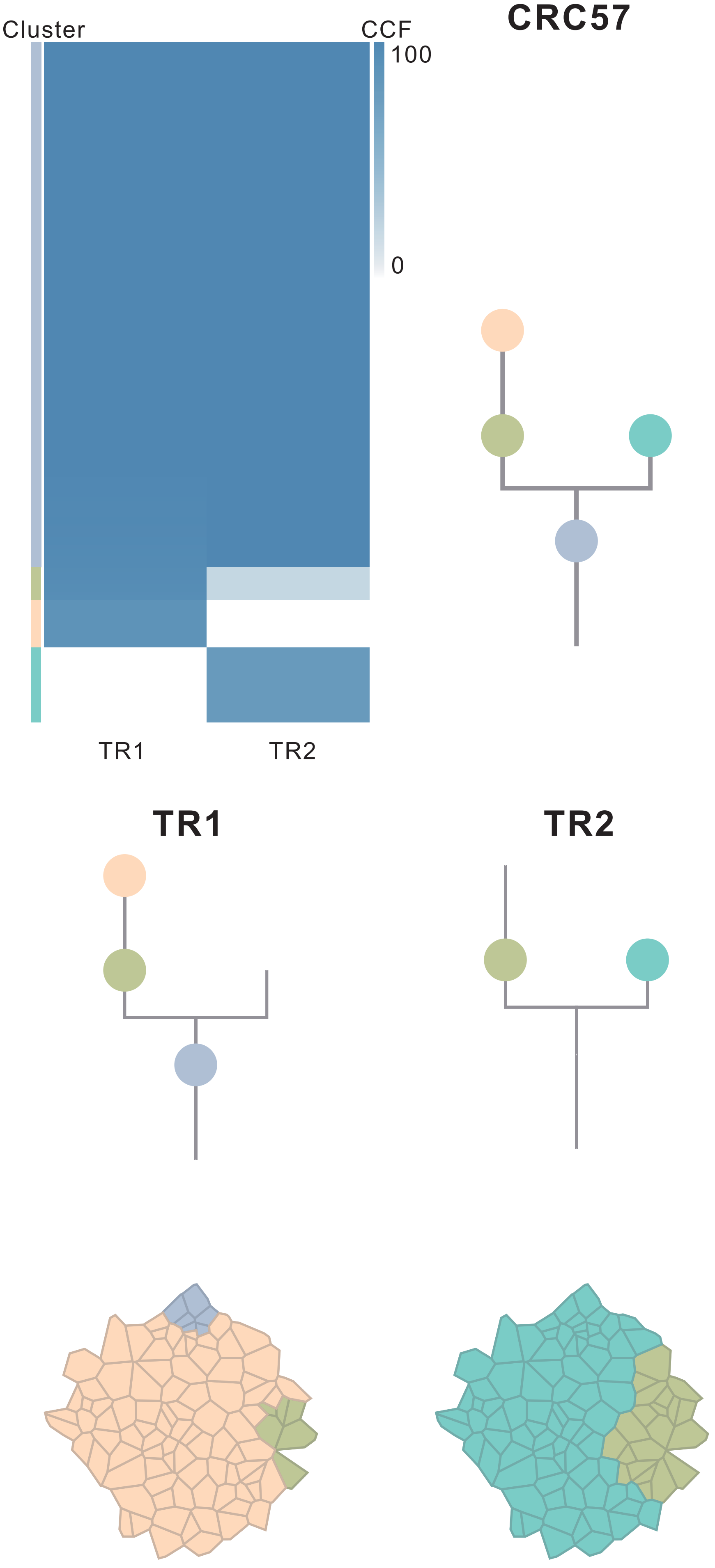

| Gene | Cytoband | Type | Cluster |
| --- | --- | --- | --- |
| *SETD2* | 3p21.31 | SNV | 1 |
| *KRAS* | 12p12.1 | SNV | 1 |
| *AXIN2* | 17q24.1 | InDel | NA |

### CRC58

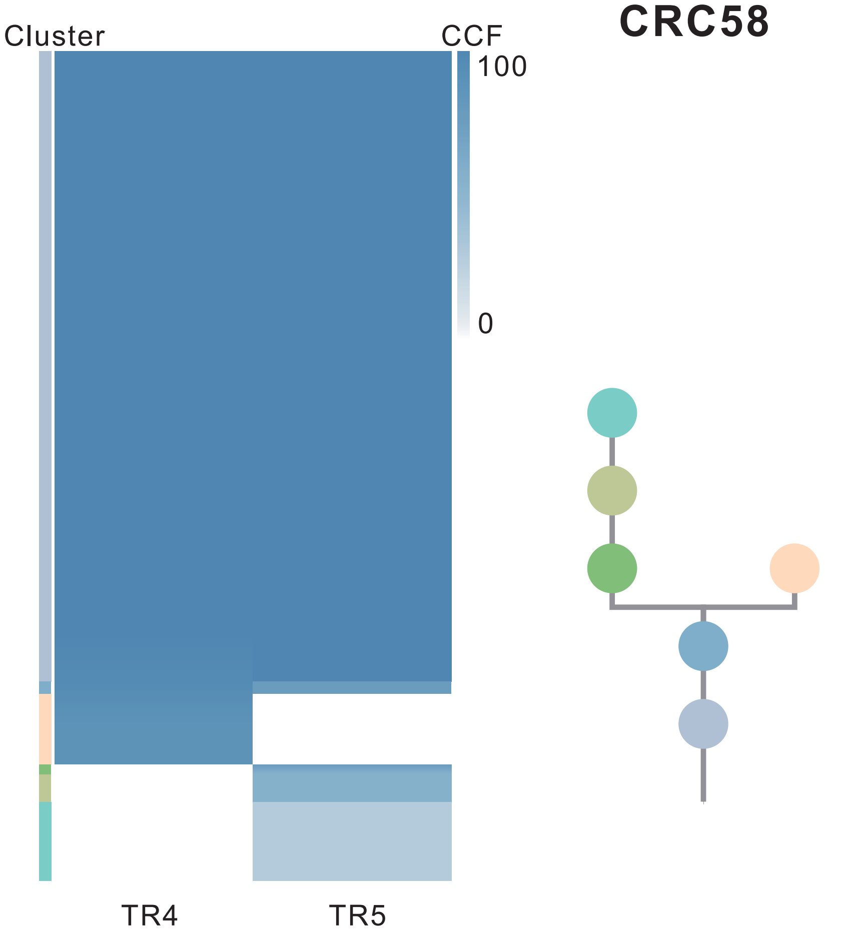

| Gene | Cytoband | Type | Cluster |
|---|---|---|---|
| *CTNNB1* | 3p22.1 | SNV | 1 |
| *ATR* | 3q23 | SNV | 1 |
| *TP53* | 17p13.1 | SNV | 1 |
| *EGFR* | 7p11.2 | Amp | 1 |
| *PLAG1* | 8q12.1 | Amp | 1 |
| *PREX2* | 8q13.2 | Amp | 1 |
| *NCOA2* | 8q13.3 | Amp | 1 |
| *RUNX1T1* | 8q21.3 | Amp | 1 |
| *HEY1* | 8q21.13 | Amp | 1 |
| *CDH17* | 8q22.1 | Amp | 1 |
| *PABPC1* | 8q22.3 | Amp | 1 |
| *UBR5* | 8q22.3 | Amp | 1 |
| *RECQL4* | 8q24.3 | Amp | 1 |
| *RAD21* | 8q24.11 | Amp | 1 |
| *MYC* | 8q24.21 | Amp | 1 |
| *FLT3* | 13q12.2 | Amp | 1 |
| *SRC* | 20q11.23 | Amp | 1 |
| *MAFB* | 20q12 | Amp | 1 |
| *PLCG1* | 20q12 | Amp | 1 |
| *NFATC2* | 20q13.2 | Amp | 1 |
| *SALL4* | 20q13.2 | Amp | 1 |
| *LZTR1* | 22q11.21 | InDel | 3 |
| *GNAS* | 20q13.32 | Amp | NA |

**TR4** **TR5**

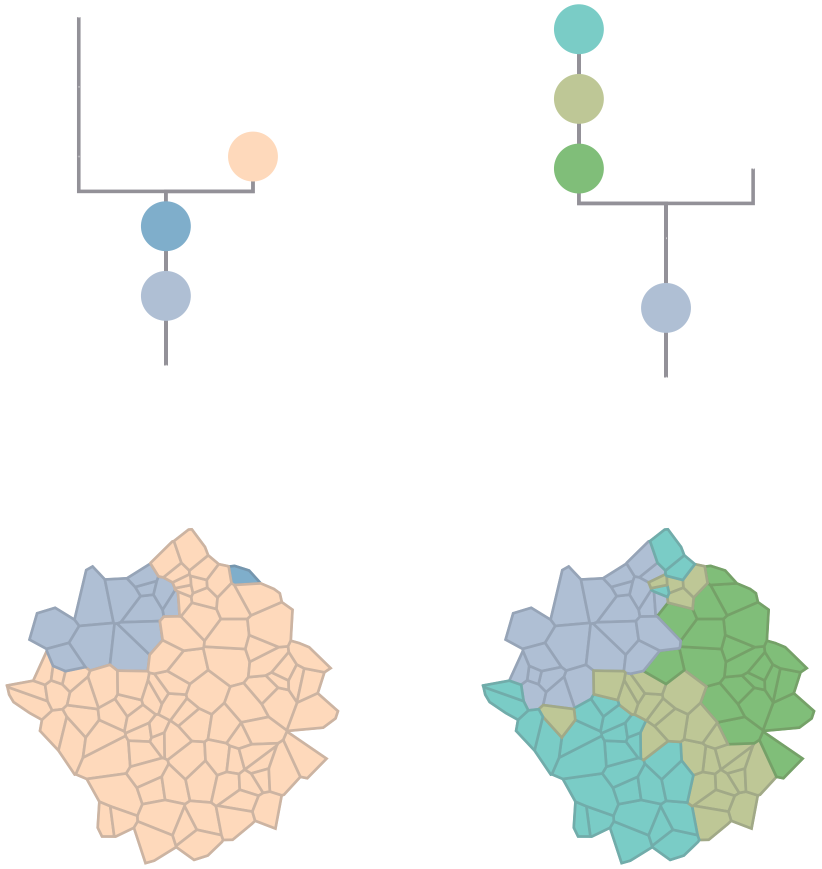

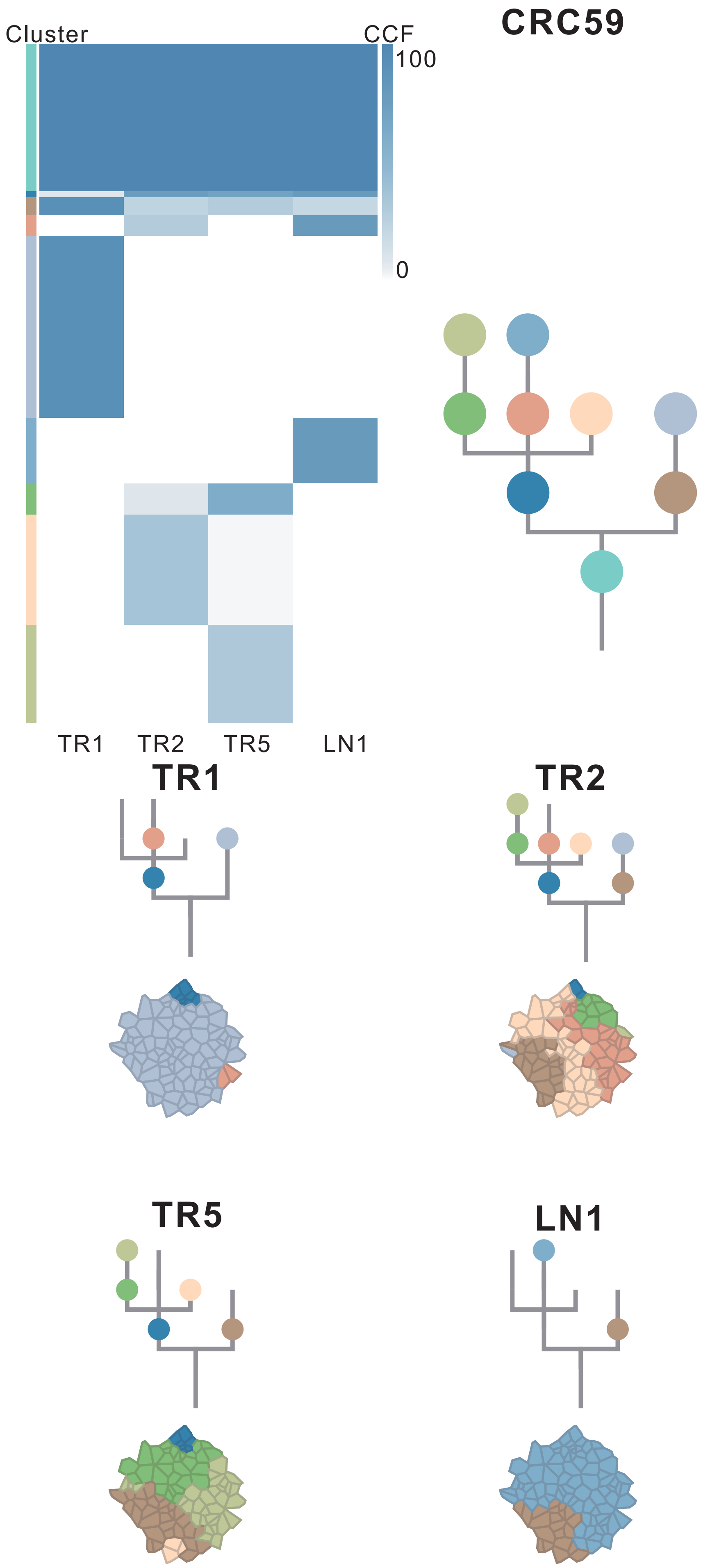

| Gene | Cytoband | Type | Cluster |
|---|---|---|---|
| *LRP1B* | 2q22.1 | SNV | 2 |
| * *APC* | 5q22.2 | SNV | 2 |
| *TCF7L2* | 10q25.3 | SNV | 2 |
| *TP53* | 17p13.1 | SNV | 2 |
| * *APC* | 5q22.2 | InDel | 2 |
| *FLT3* | 13q12.2 | Amp | 2 |
| *SRC* | 20q11.23 | Amp | 2 |
| *MAFB* | 20q12 | Amp | 2 |
| *PLCG1* | 20q12 | Amp | 2 |
| *NFATC2* | 20q13.2 | Amp | 2 |
| *SALL4* | 20q13.2 | Amp | 2 |
| *GNAS* | 20q13.32 | Amp | 2 |
| *PTK6* | 20q13.33 | Amp | 2 |
| *LEPROTL1* | 8p12 | Del | 2 |
| *WRN* | 8p12 | Del | 2 |
| *ARHGEF10* | 8p23.3 | Del | 2 |
| *SMAD2* | 18q21.1 | Del | 2 |
| *SMAD4* | 18q21.2 | Del | 2 |
| *RUNX1* | 21q22.12 | Del | 2 |
| *PIK3CA* | 3q26.32 | SNV | 3 |
| *CHD2* | 15q26.1 | SNV | 3 |
| *CREBBP* | 16p13.3 | SNV | 4 |
| *PBRM1* | 3p21.1 | InDel | 4 |
| *IKBKB* | 8p11.21 | Amp | 8 |
| *KAT6A* | 8p11.21 | Amp | 8 |
| *PLAG1* | 8q12.1 | Amp | 8 |
| *RUNX1T1* | 8q21.3 | Amp | 8 |
| *CDH17* | 8q22.1 | Amp | 8 |
| *PABPC1* | 8q22.3 | Amp | 8 |
| *UBR5* | 8q22.3 | Amp | 8 |
| *RAD21* | 8q24.11 | Amp | 8 |
| *MYC* | 8q24.21 | Amp | 8 |
| *NRG1* | 8p12 | Del | 8 |
| *HEY1* | 8q21.13 | Amp | NA |
| *OLIG2* | 21q22.11 | Amp | NA |

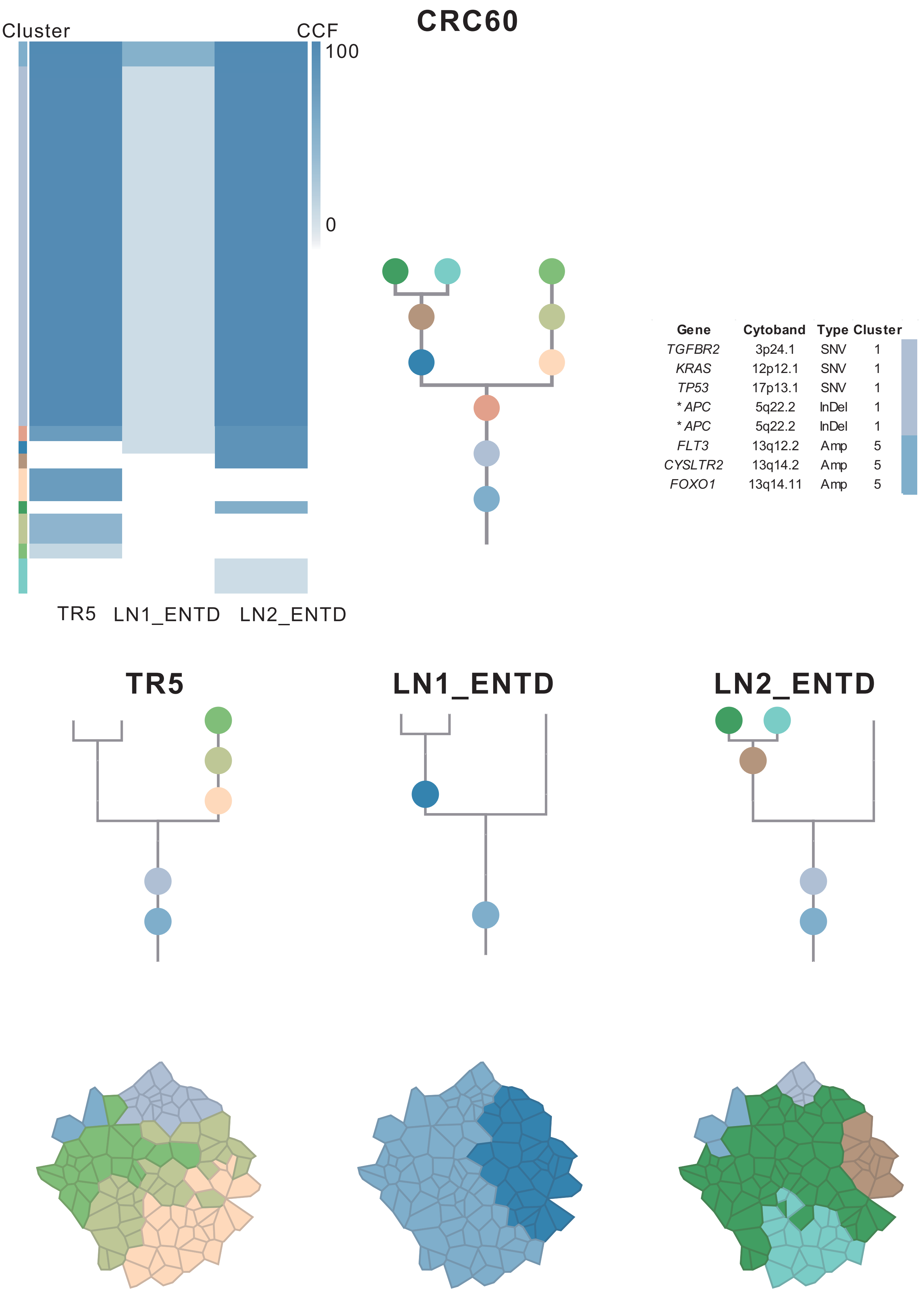

| Gene | Cytoband | Type | Cluster |
|---|---|---|---|
| *TGFBR2* | 3p24.1 | SNV | 1 |
| *KRAS* | 12p12.1 | SNV | 1 |
| *TP53* | 17p13.1 | SNV | 1 |
| * *APC* | 5q22.2 | InDel | 1 |
| * *APC* | 5q22.2 | InDel | 1 |
| *FLT3* | 13q12.2 | Amp | 5 |
| *CYSLTR2* | 13q14.2 | Amp | 5 |
| *FOXO1* | 13q14.11 | Amp | 5 |

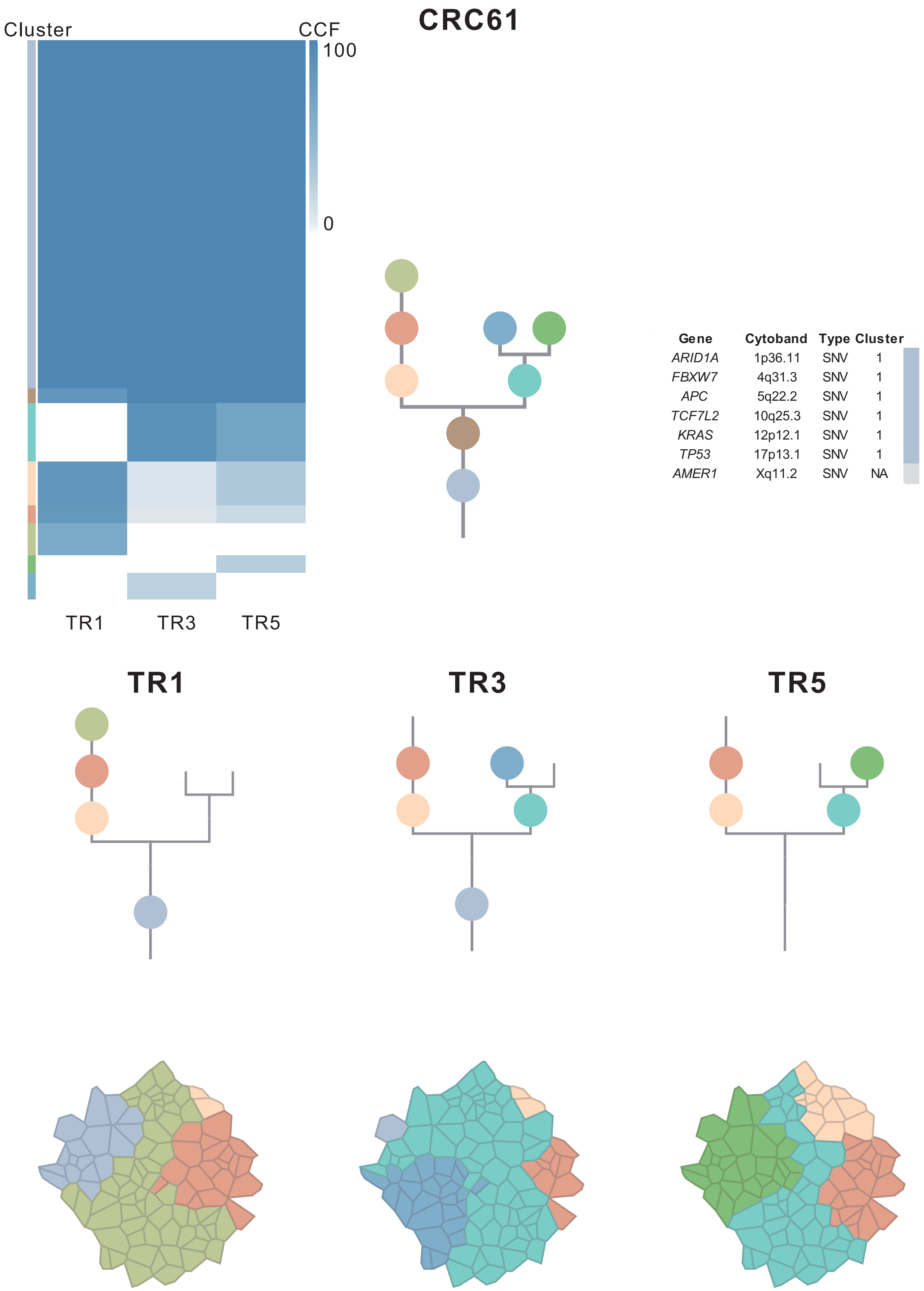

| Gene | Cytoband | Type | Cluster |
|---|---|---|---|
| *ARID1A* | 1p36.11 | SNV | 1 |
| *FBXW7* | 4q31.3 | SNV | 1 |
| *APC* | 5q22.2 | SNV | 1 |
| *TCF7L2* | 10q25.3 | SNV | 1 |
| *KRAS* | 12p12.1 | SNV | 1 |
| *TP53* | 17p13.1 | SNV | 1 |
| *AMER1* | Xq11.2 | SNV | NA |

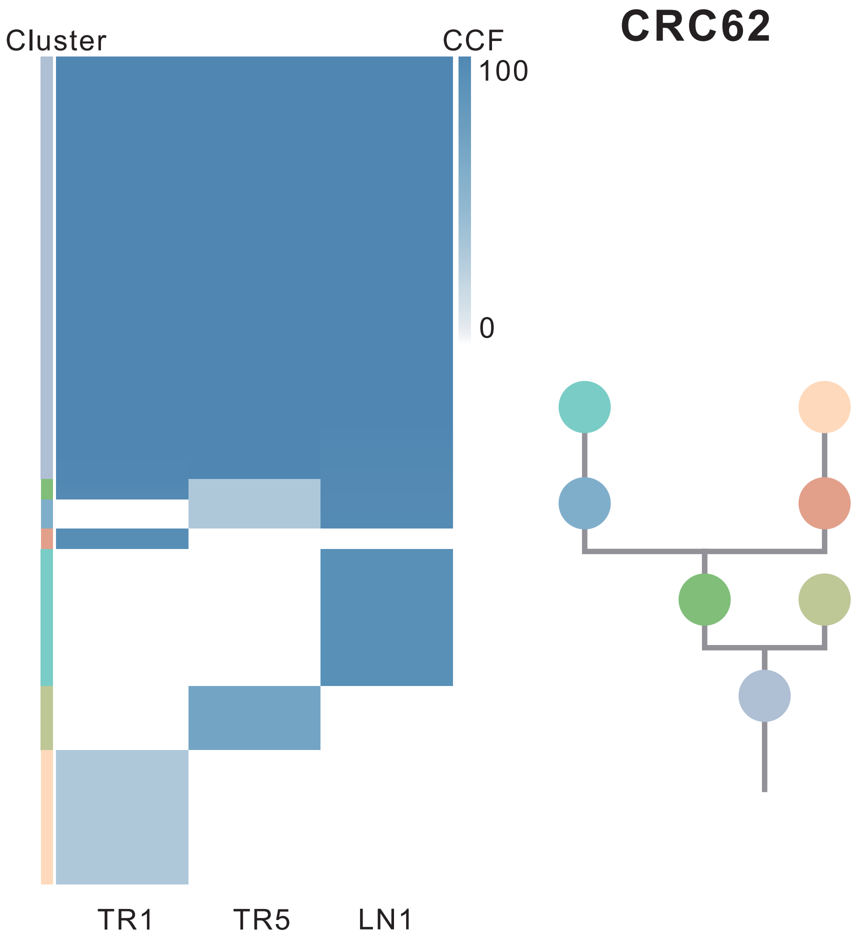

| Gene | Cytoband | Type | Cluster |
|---|---|---|---|
| *TP53* | 17p13.1 | SNV | 1 |
| *ARID1A* | 1p36.11 | InDel | 1 |
| *IL7R* | 5p13.2 | Amp | 1 |
| *CTNND2* | 5p15.2 | Amp | 1 |
| *TERT* | 5p15.33 | Amp | 1 |
| *SRC* | 20q11.23 | Amp | 1 |
| *MAFB* | 20q12 | Amp | 1 |
| *PLCG1* | 20q12 | Amp | 1 |
| *NFATC2* | 20q13.2 | Amp | 1 |
| *SALL4* | 20q13.2 | Amp | 1 |
| *GNAS* | 20q13.32 | Amp | 1 |
| *PTK6* | 20q13.33 | Amp | 1 |
| *GPC5* | 13q31.3 | SNV | NA |

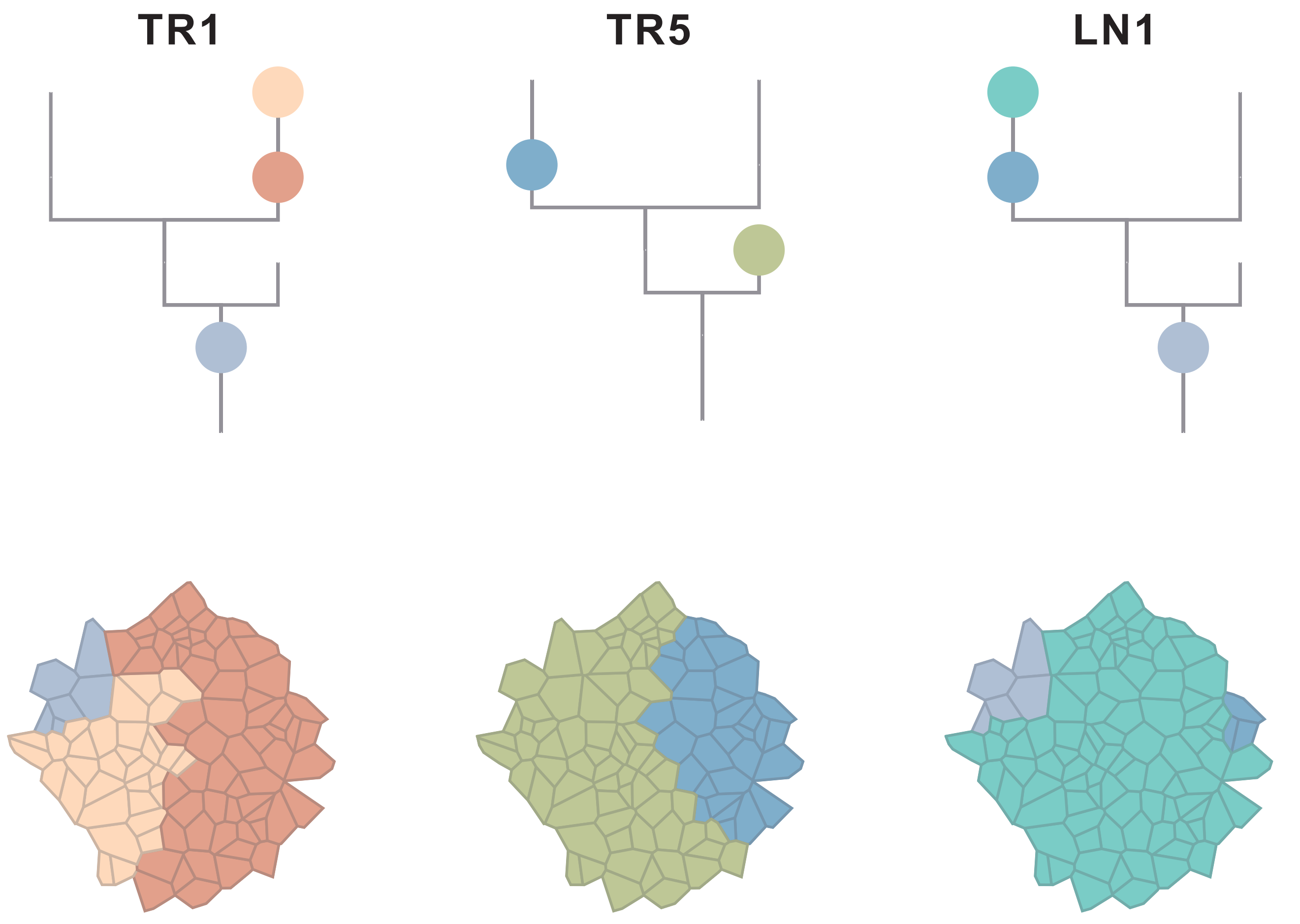

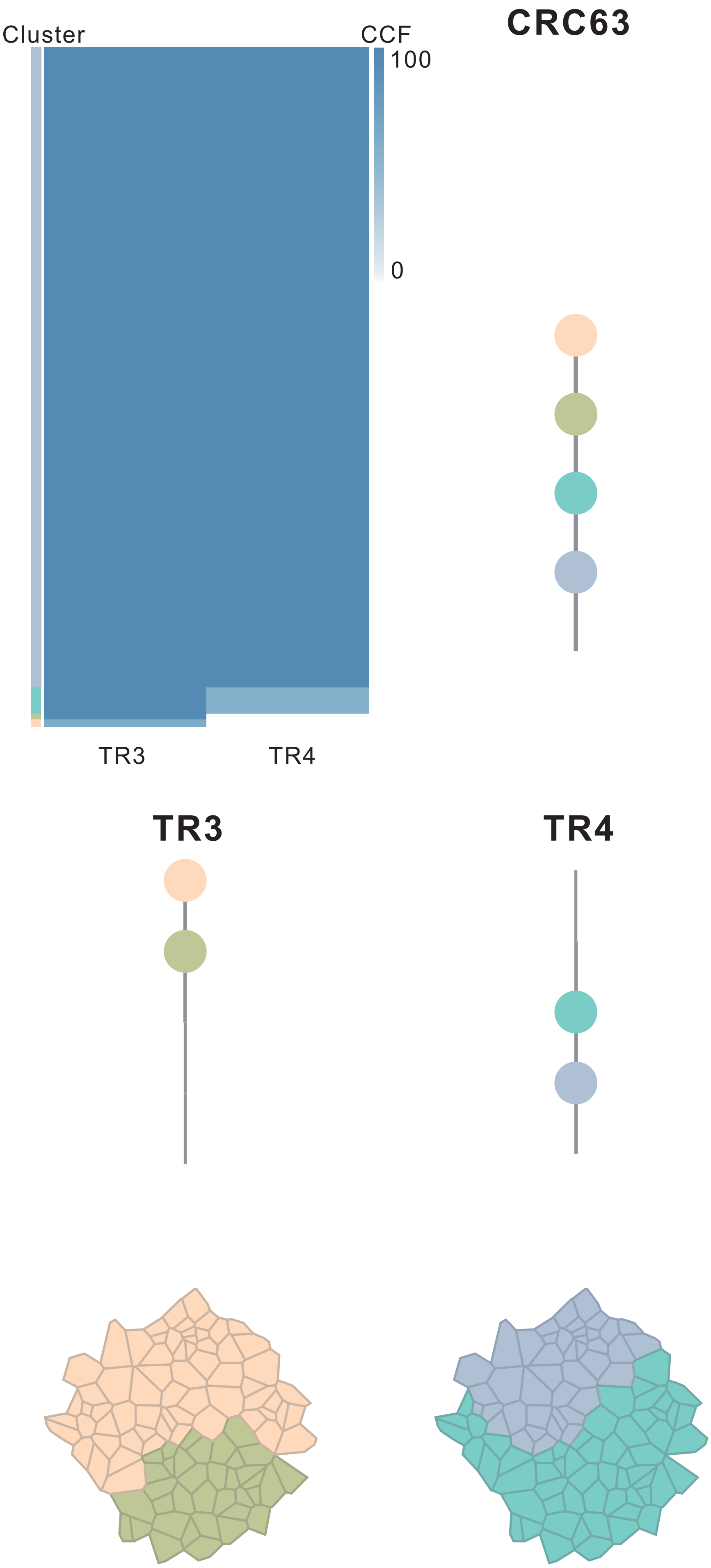

| Gene | Cytoband | Type | Cluster |
| --- | --- | --- | --- |
| *ARNT* | 1q21.3 | SNV | 1 |
| *ASXL2* | 2p23.3 | SNV | 1 |
| *FBXW7* | 4q31.3 | SNV | 1 |
| * *APC* | 5q22.2 | SNV | 1 |
| *TSC1* | 9q34.13 | SNV | 1 |
| * *TSC2* | 16p13.3 | SNV | 1 |
| * *TSC2* | 16p13.3 | SNV | 1 |
| *ERBB2* | 17q12 | SNV | 1 |
| *ARHGAP26* | 5q31.3 | SNV | 2 |
| * *APC* | 5q22.2 | InDel | 2 |
| *TP53* | 17p13.1 | SNV | NA |

### CRC64

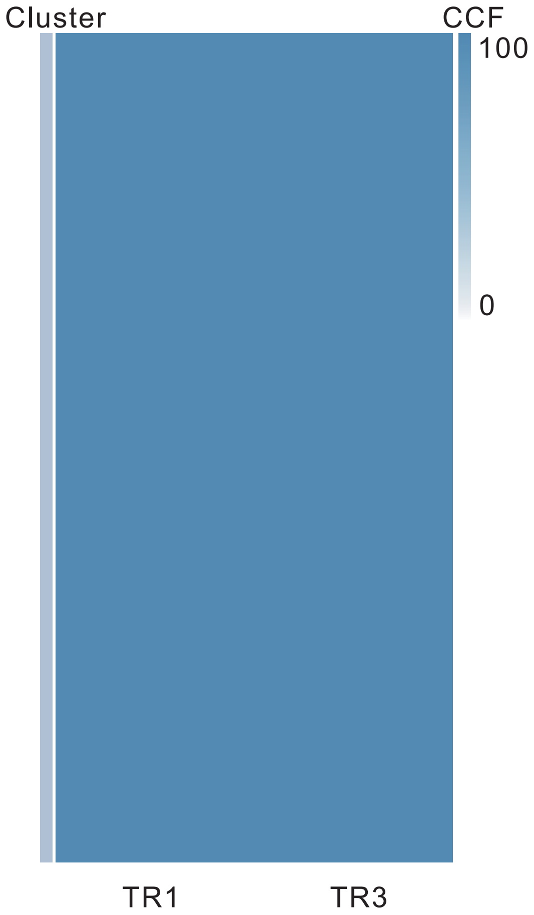

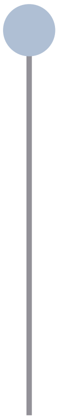

| Gene | Cytoband | Type | Cluster |
|---|---|---|---|
| *PIK3CA* | 3q26.32 | SNV | NA |
| *FBXW7* | 4q31.3 | SNV | NA |
| *KRAS* | 12p12.1 | SNV | NA |
| *SMAD2* | 18q21.1 | SNV | NA |
| *APC* | 5q22.2 | InDel | NA |
| *CBFA2T3* | 16q24.3 | Del | NA |

**TR1**

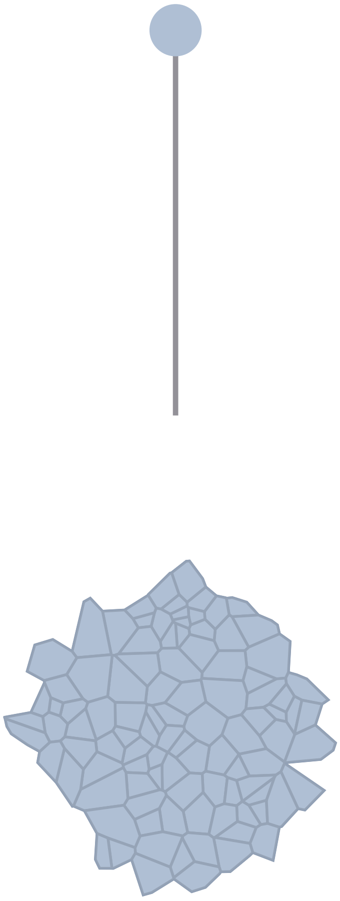

**TR3**

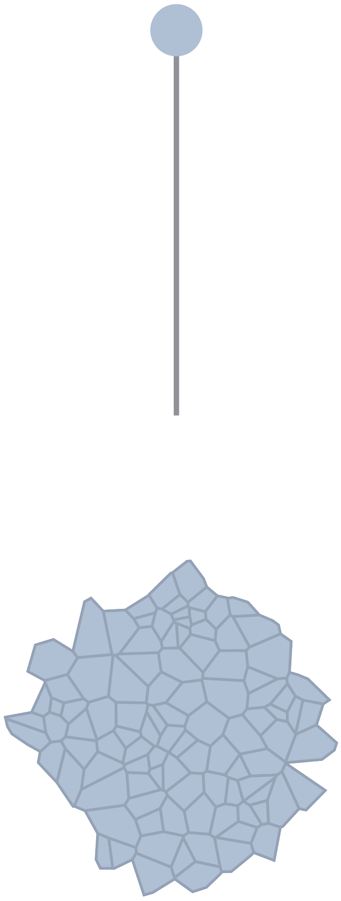

### CRC65

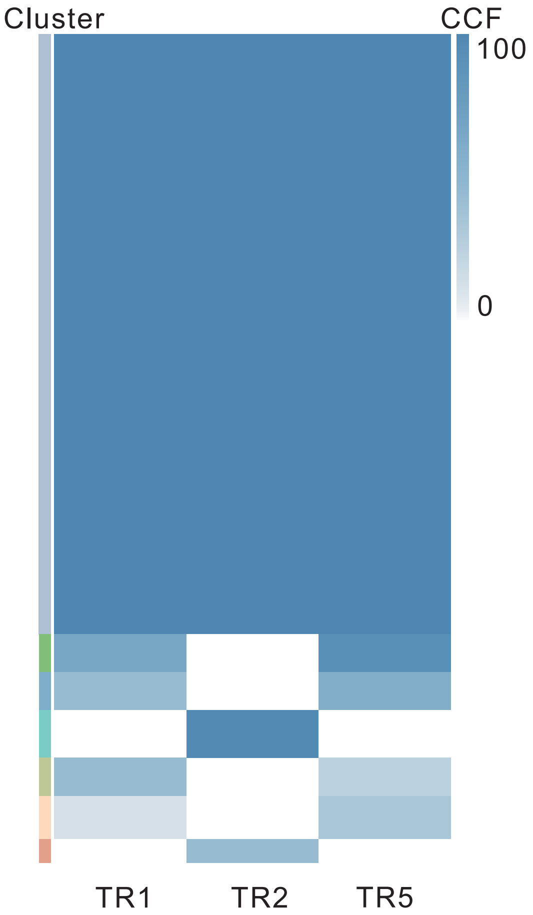

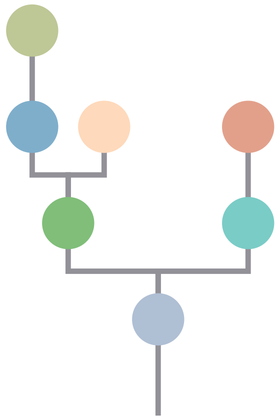

| Gene | Cytoband | Type | Cluster |
|---|---|---|---|
| *FAT4* | 4q28.1 | SNV | 1 |
| *CSMD3* | 8q23.3 | SNV | 1 |
| *KRAS* | 12p12.1 | SNV | 1 |
| *TP53* | 17p13.1 | SNV | 1 |
| *APC* | 5q22.2 | InDel | 1 |
| *SRC* | 20q11.23 | Amp | 1 |
| *MAFB* | 20q12 | Amp | 1 |
| *PLCG1* | 20q12 | Amp | 1 |
| *NFATC2* | 20q13.2 | Amp | 1 |
| *SALL4* | 20q13.2 | Amp | 1 |
| *PTK6* | 20q13.33 | Amp | 1 |
| *SIRPA* | 20p13 | Del | 1 |
| *CLTCL1* | 22q11.21 | Del | 1 |
| *LZTR1* | 22q11.21 | Del | 1 |
| *SMARCB1* | 22q11.23 | Del | 1 |
| *CHEK2* | 22q12.1 | Del | 1 |
| *ZNRF3* | 22q12.1 | Del | 1 |
| *NF2* | 22q12.2 | Del | 1 |
| *MYH9* | 22q12.3 | Del | 1 |
| *APOBEC3B* | 22q13.1 | Del | 1 |
| *EP300* | 22q13.2 | Del | 1 |
| *MKL1* | 22q13.2 | Del | 1 |
| *TBX3* | 12q24.21 | InDel | NA |
| *LZTR1* | 22q11.21 | InDel | NA |
| *GNAS* | 20q13.32 | Amp | NA |

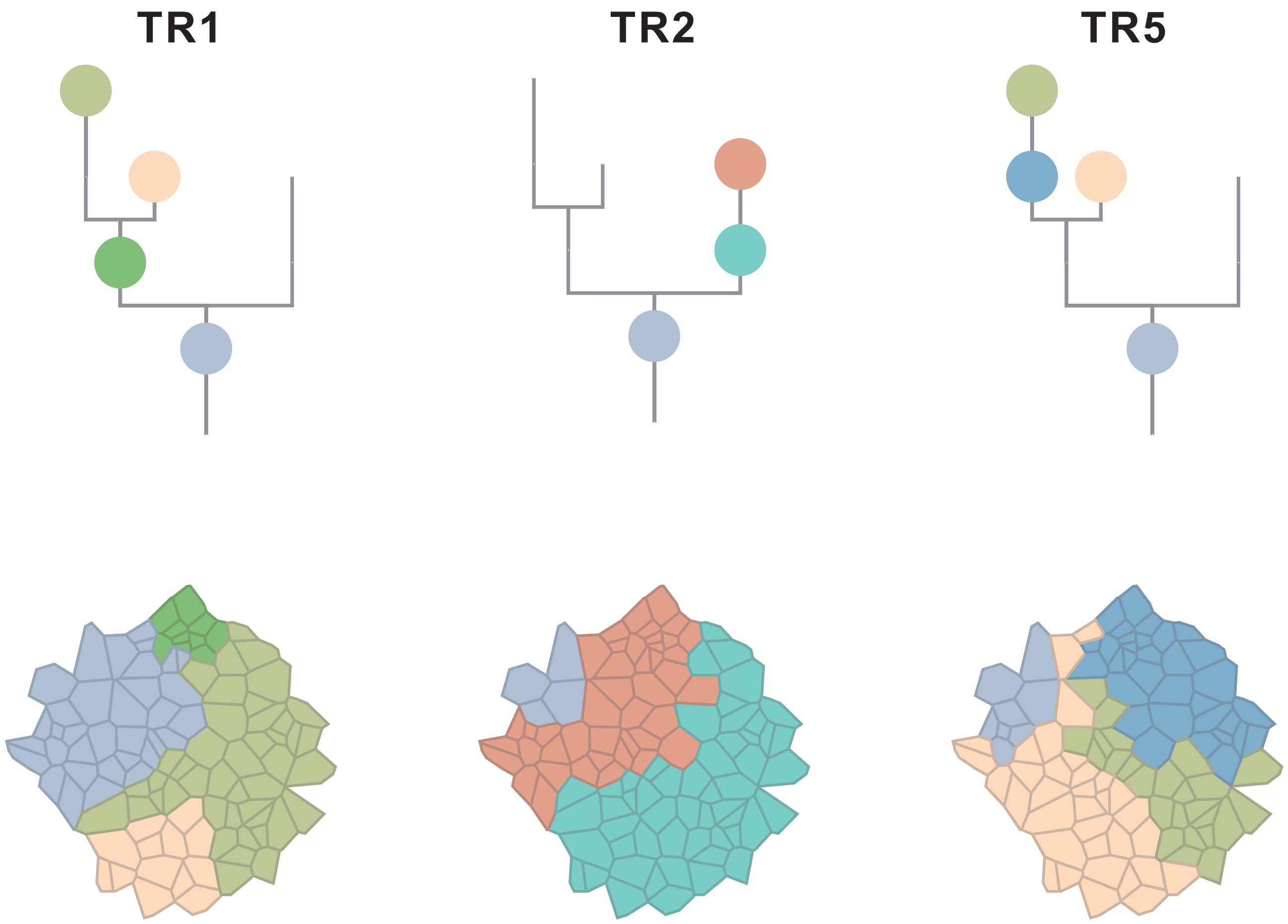

### CRC66

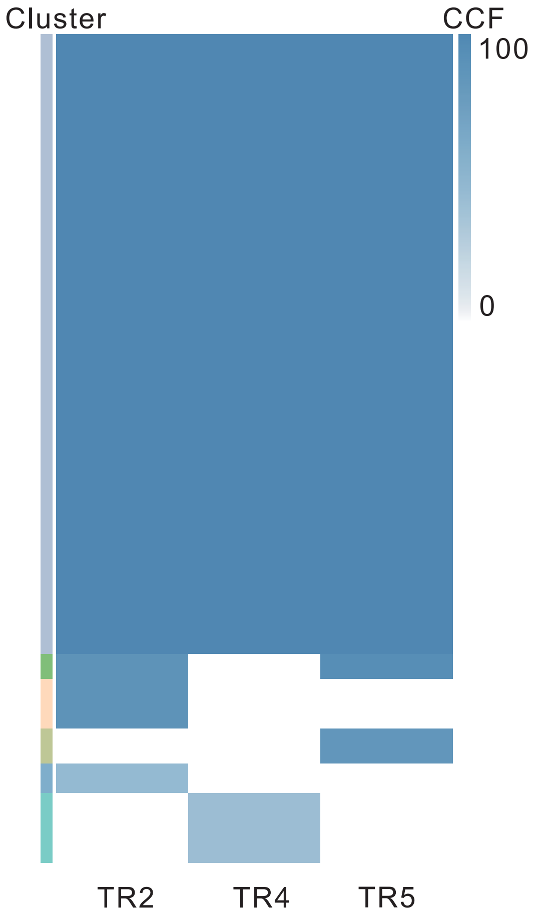

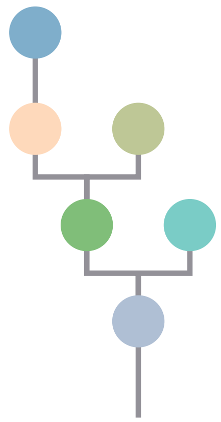

| Gene | Cytoband | Type | Cluster |
|---|---|---|---|
| * *FBXW7* | 4q31.3 | SNV | 1 |
| * *FBXW7* | 4q31.3 | SNV | 1 |
| * *FBXW7* | 4q31.3 | SNV | 1 |
| *APC* | 5q22.2 | SNV | 1 |
| *CDH11* | 16q21 | SNV | 1 |
| *TP53* | 17p13.1 | SNV | 1 |
| *SRC* | 20q11.23 | Amp | 1 |
| *MAFB* | 20q12 | Amp | 1 |
| *PLCG1* | 20q12 | Amp | 1 |
| *NFATC2* | 20q13.2 | Amp | 1 |
| *SALL4* | 20q13.2 | Amp | 1 |
| *GNAS* | 20q13.32 | Amp | 1 |
| *PTK6* | 20q13.33 | Amp | 1 |

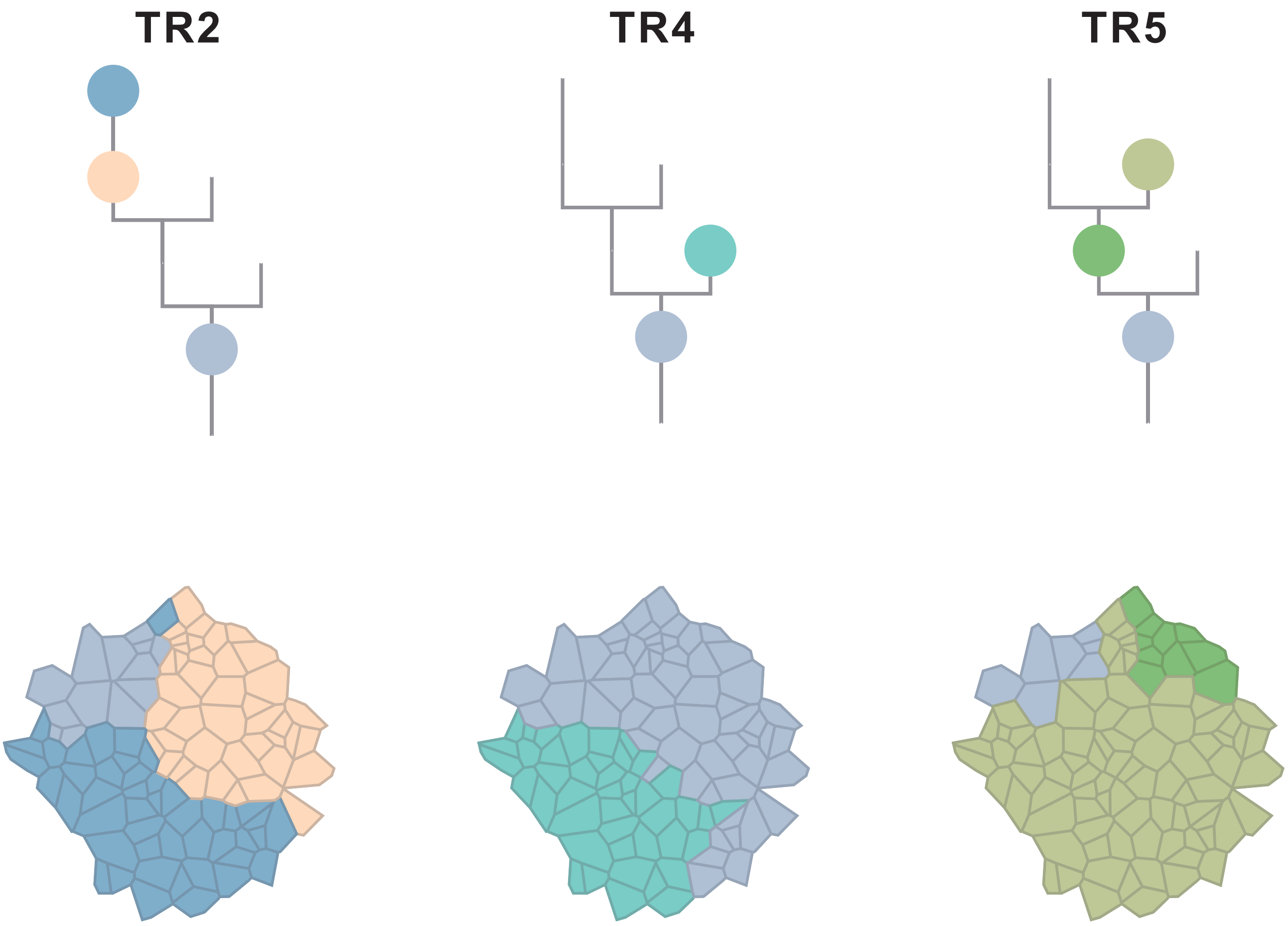

### CRC67

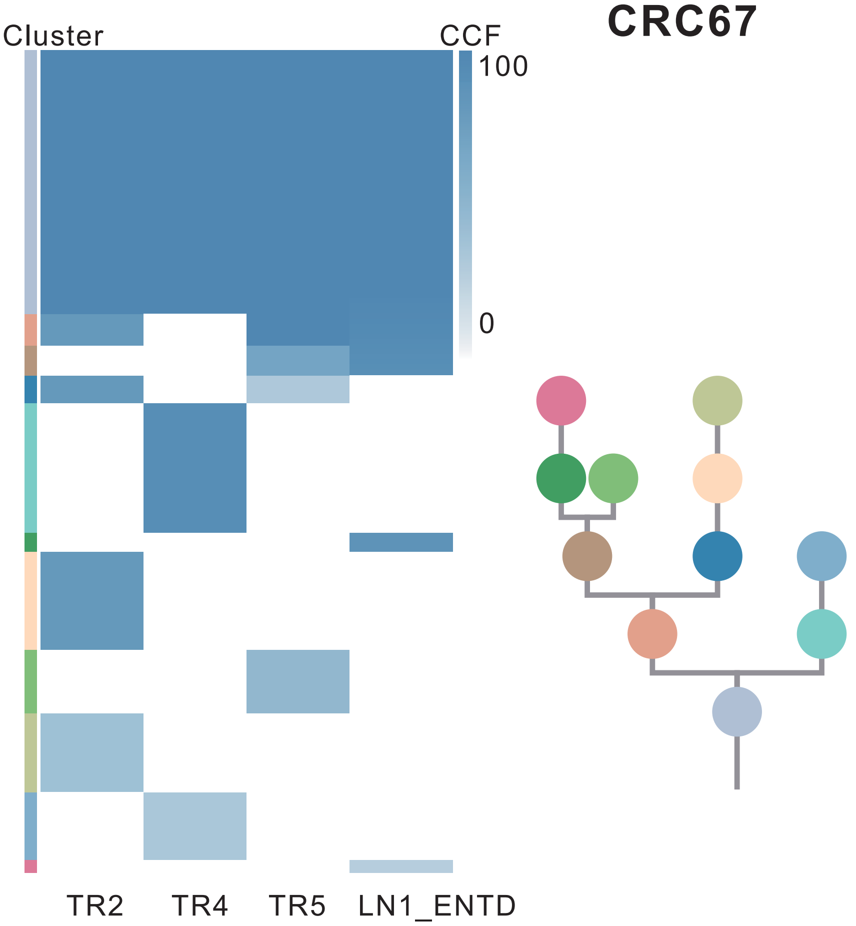

| Gene | Cytoband | Type | Cluster |
|---|---|---|---|
| * *LRP1B* | 2q22.1 | SNV | 1 |
| *PIK3CA* | 3q26.32 | SNV | 1 |
| *APC* | 5q22.2 | SNV | 1 |
| *EZH2* | 7q36.1 | SNV | 1 |
| *TP53* | 17p13.1 | SNV | 1 |
| *SMAD4* | 18q21.2 | SNV | 1 |
| *ARHGAP26* | 5q31.3 | SNV | 3 |
| * *LRP1B* | 2q22.1 | SNV | 6 |
| * *LRP1B* | 2q22.1 | SNV | 9 |

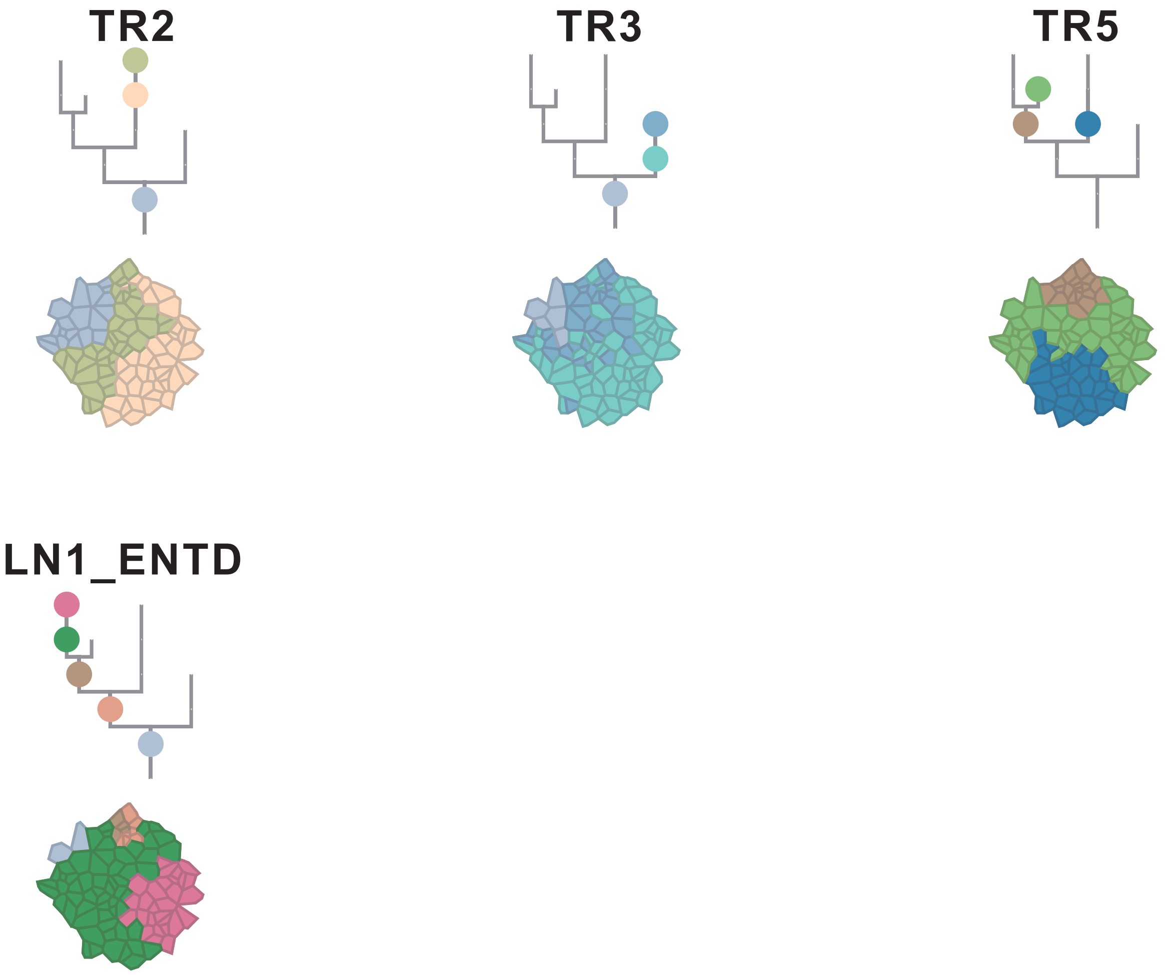

### CRC68

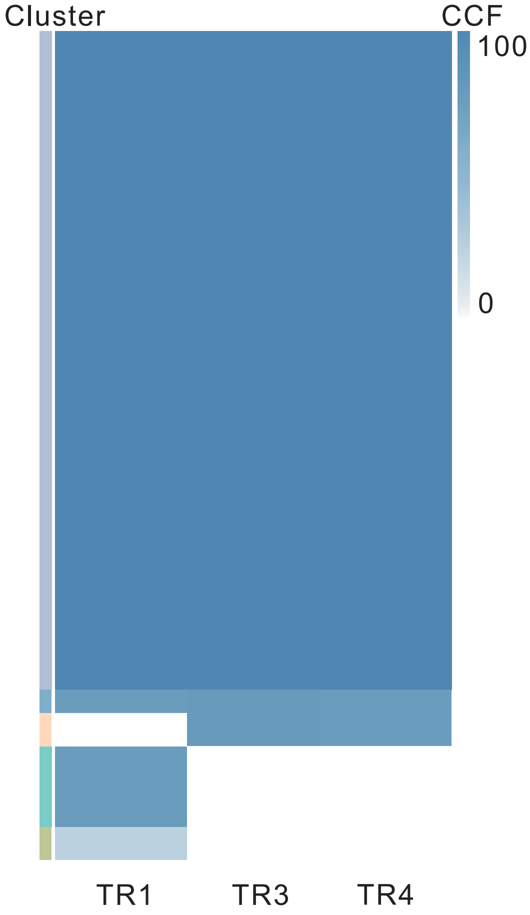

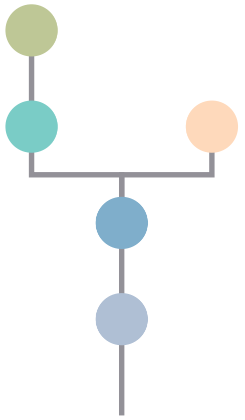

| Gene | Cytoband | Type | Cluster |
|---|---|---|---|
| *NRAS* | 1p13.2 | SNV | 1 |
| *PIK3CA* | 3q26.32 | SNV | 1 |
| * *APC* | 5q22.2 | SNV | 1 |
| *KRAS* | 12p12.1 | SNV | 1 |
| *CDH11* | 16q21 | SNV | 1 |
| *TP53* | 17p13.1 | SNV | 1 |
| * *APC* | 5q22.2 | InDel | 1 |
| *SRC* | 20q11.23 | Amp | 1 |
| *MAFB* | 20q12 | Amp | 1 |
| *PLCG1* | 20q12 | Amp | 1 |
| *NFATC2* | 20q13.2 | Amp | 1 |
| *SALL4* | 20q13.2 | Amp | 1 |
| *GNAS* | 20q13.32 | Amp | 1 |
| *PTK6* | 20q13.33 | Amp | 1 |
| *PRKAR1A* | 17q24.2 | SNV | NA |

## TR1

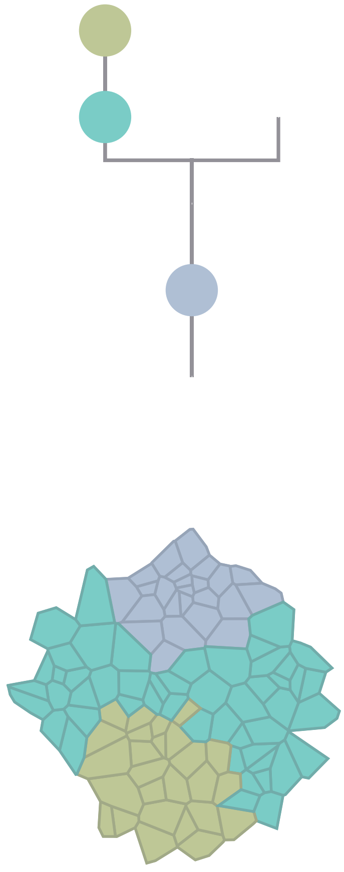

## TR3

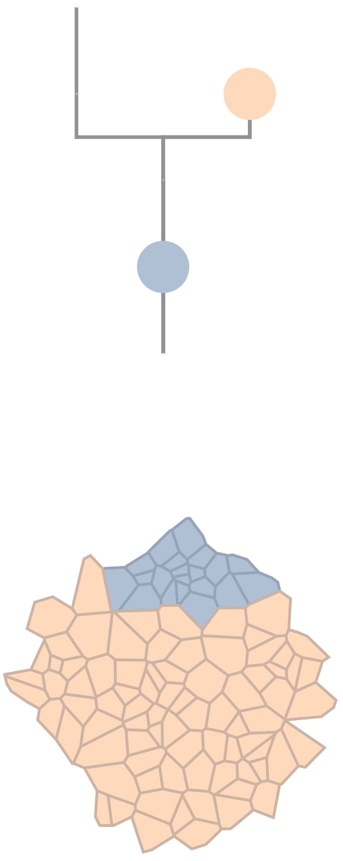

## TR4

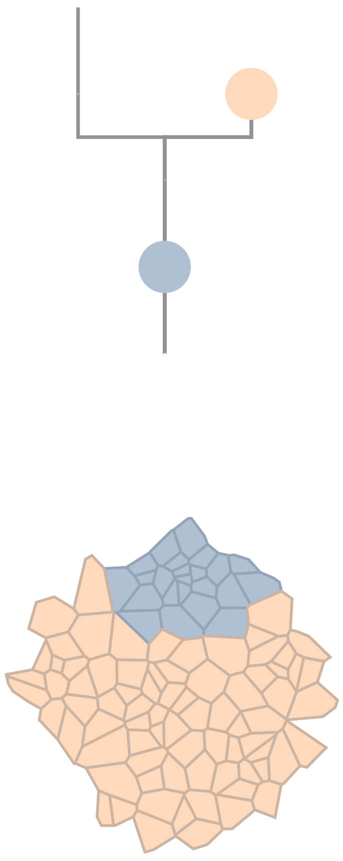

### Figure S9

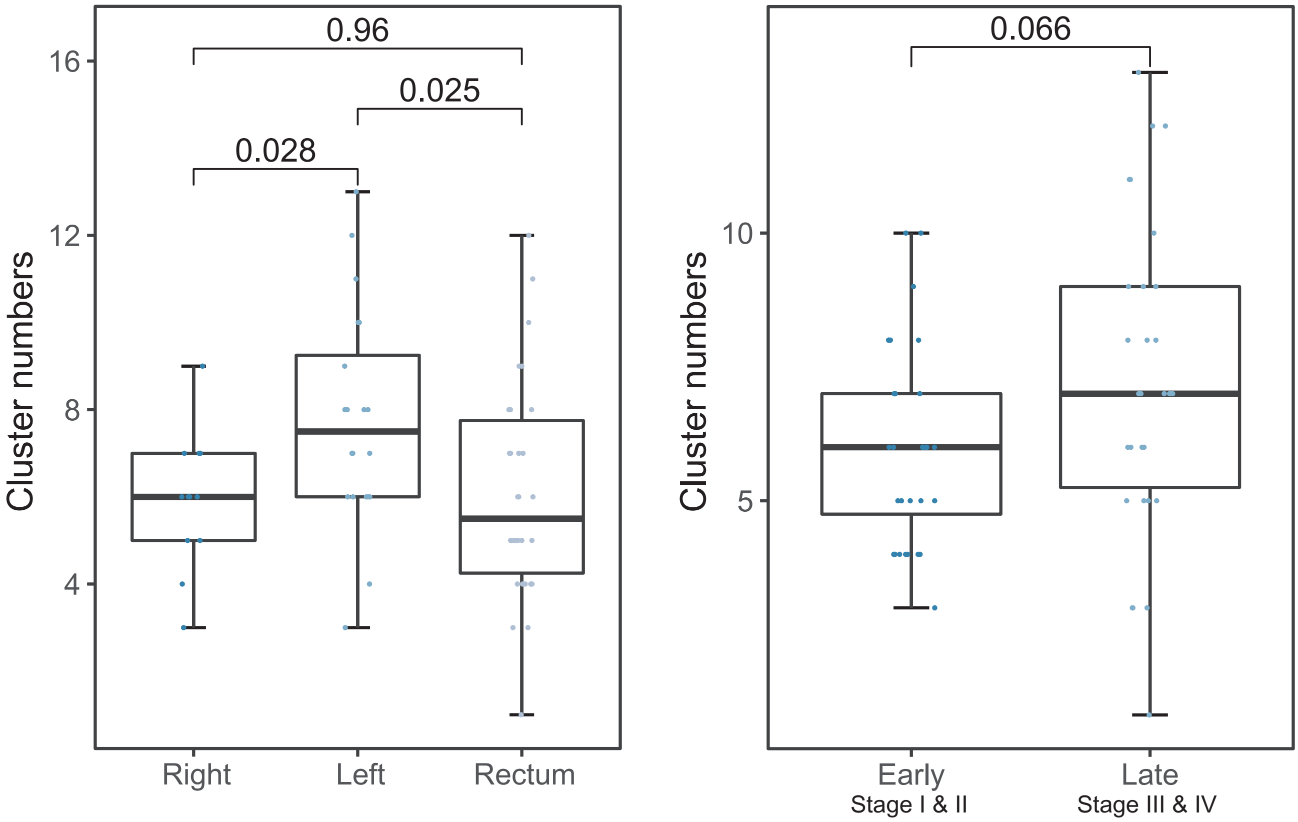

**Figure S9. Difference of cluster numbers in CRC tumors by right / left / rectum and early / late stage of cancers.**
Box plots of cluster numbers in CRC tumors by position and stage.

**Figure S10**

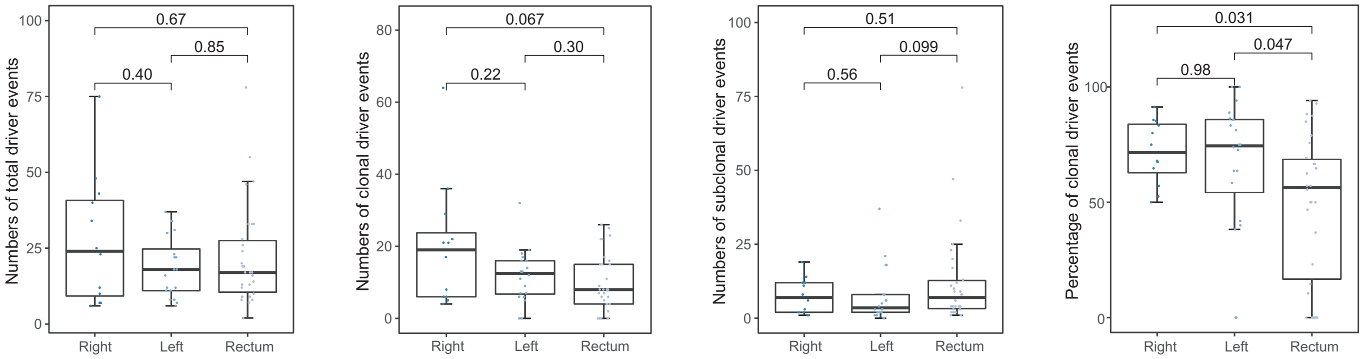

**Figure S10. Difference in numbers of driver events in CRC tumors among right-sided colon, left-sided colon and rectal cancers.** Box plots of total, clonal, subclonal and percentage of clonal numbers of driver events by tumor position.

#### Figure S11

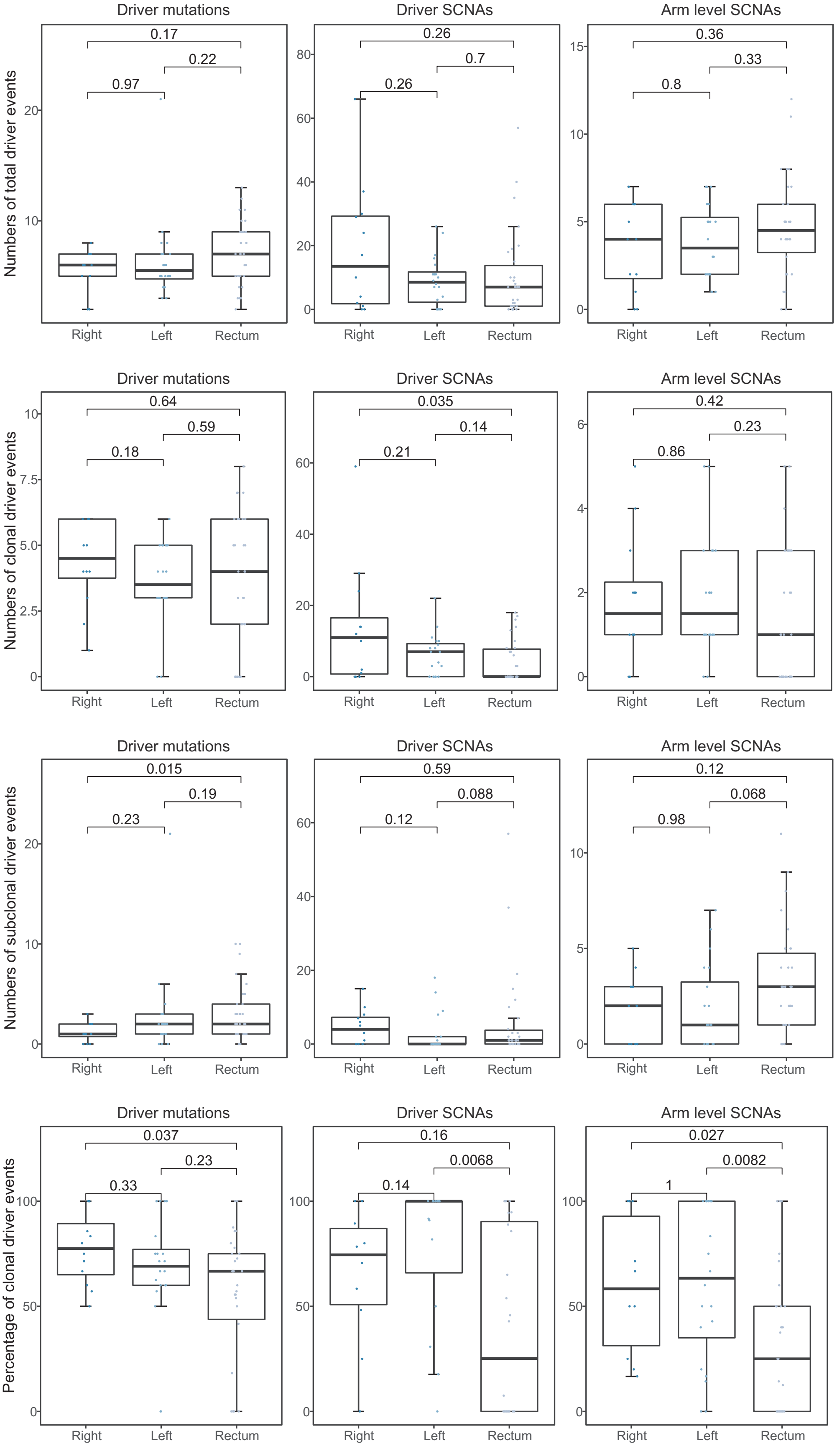

**Figure S11. Difference in types of driver events in CRC tumors among right-sided colon, left-sided colon and rectal cancers.**
Box plots of total, clonal, subclonal and percentage of clonal numbers of different driver events types by tumor position.

**Figure S12**

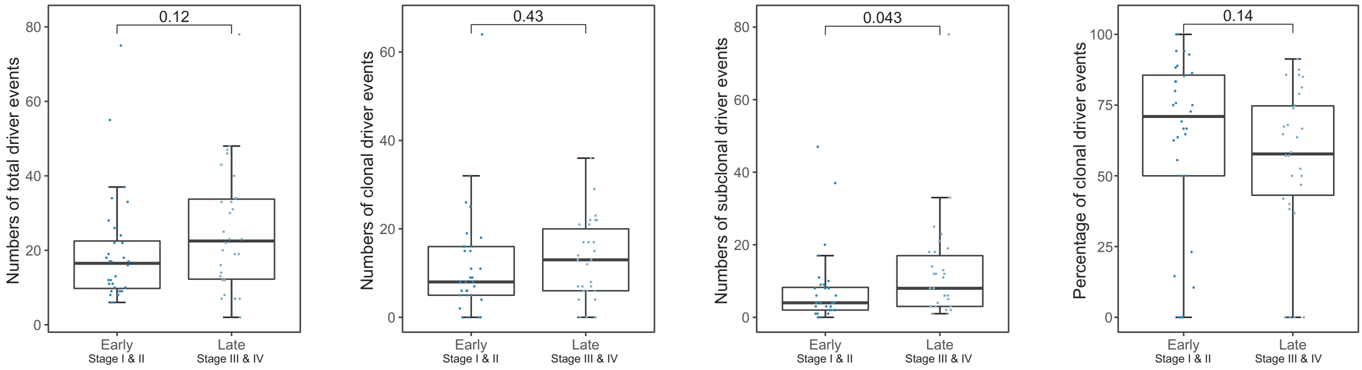

**Figure S12. Difference in numbers of driver events in CRC tumors between early and late stage of cancers.**

Box plots of total, clonal, subclonal and percentage of clonal numbers of driver events by tumor stage.

#### Figure S13

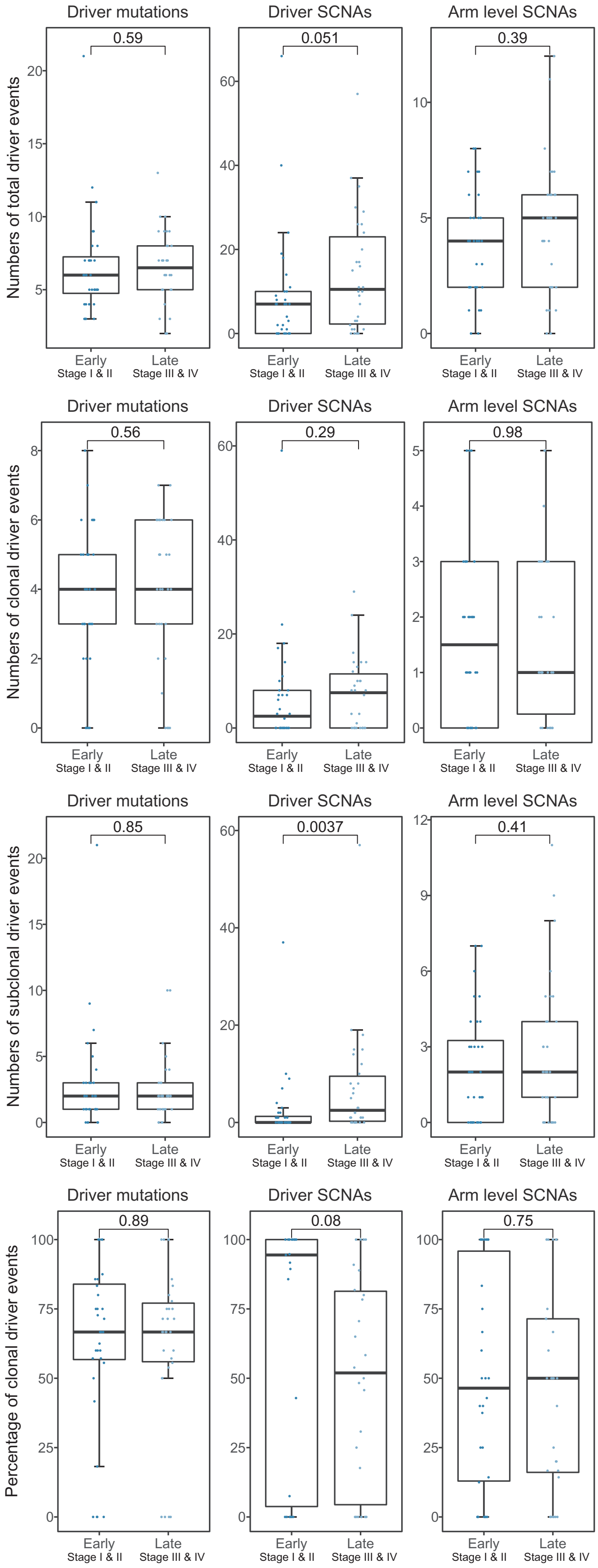

**Figure S13. Difference in types of driver events in CRC tumors between early and late stage of cancers.**

Box plots of total, clonal, subclonal and percentage of clonal numbers of different driver events types by tumor stage.

#### Figure S14

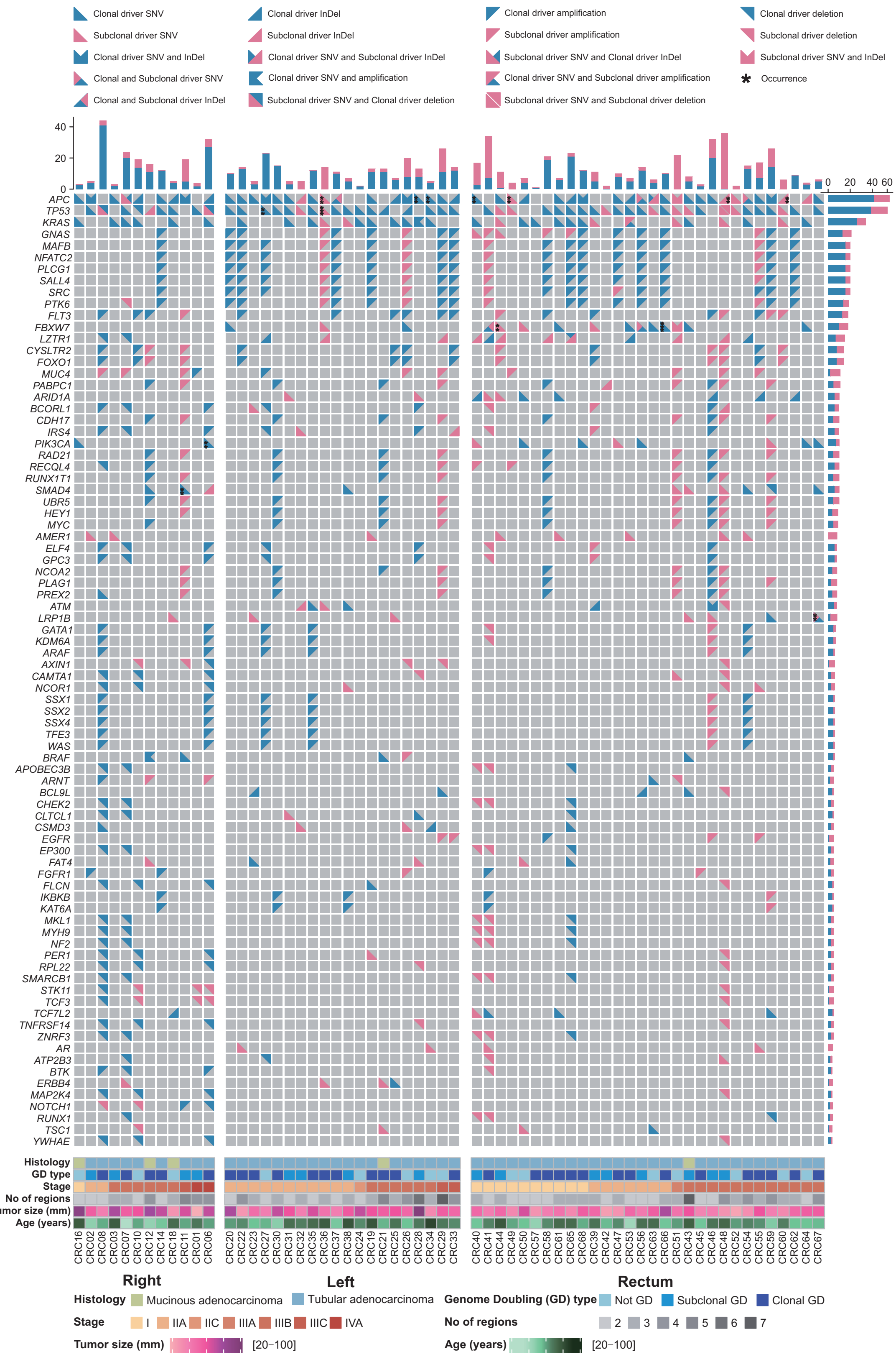

**Figure S14. Heterogeneity of driver alterations in CRC tumors.**

Driver alterations detected in each tumor. Only genes containing ≥ 4 driver alterations across the CRC patients are included. The shape and color represented the alteration type and clonal status as shown in the top panel. The second top panel displayed the number of driver alterations identified across individual CRC tumors and the bar plots to the right showed the number of the variants in right-sided colon, left-sided colon and rectal cancers for each gene. The lower panel showed the demographic and clinical characteristics of the 62 CRC patients in this study (divided by histology; whole genome doubling status; stage; number of regions; tumor size; age and tumor location).

### Figure S15

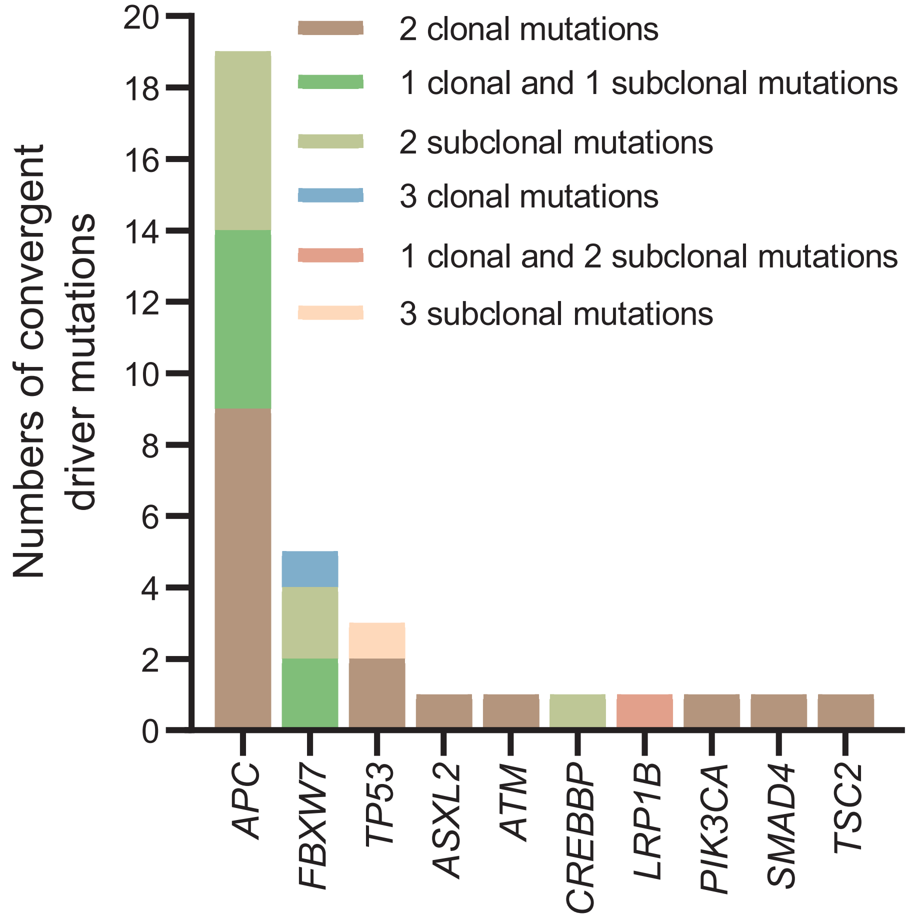

**Figure S15. Convergent driver mutations in CRC tumors.** All convergent driver mutation events were summarized in 62 CRC tumors.

### Figure S16

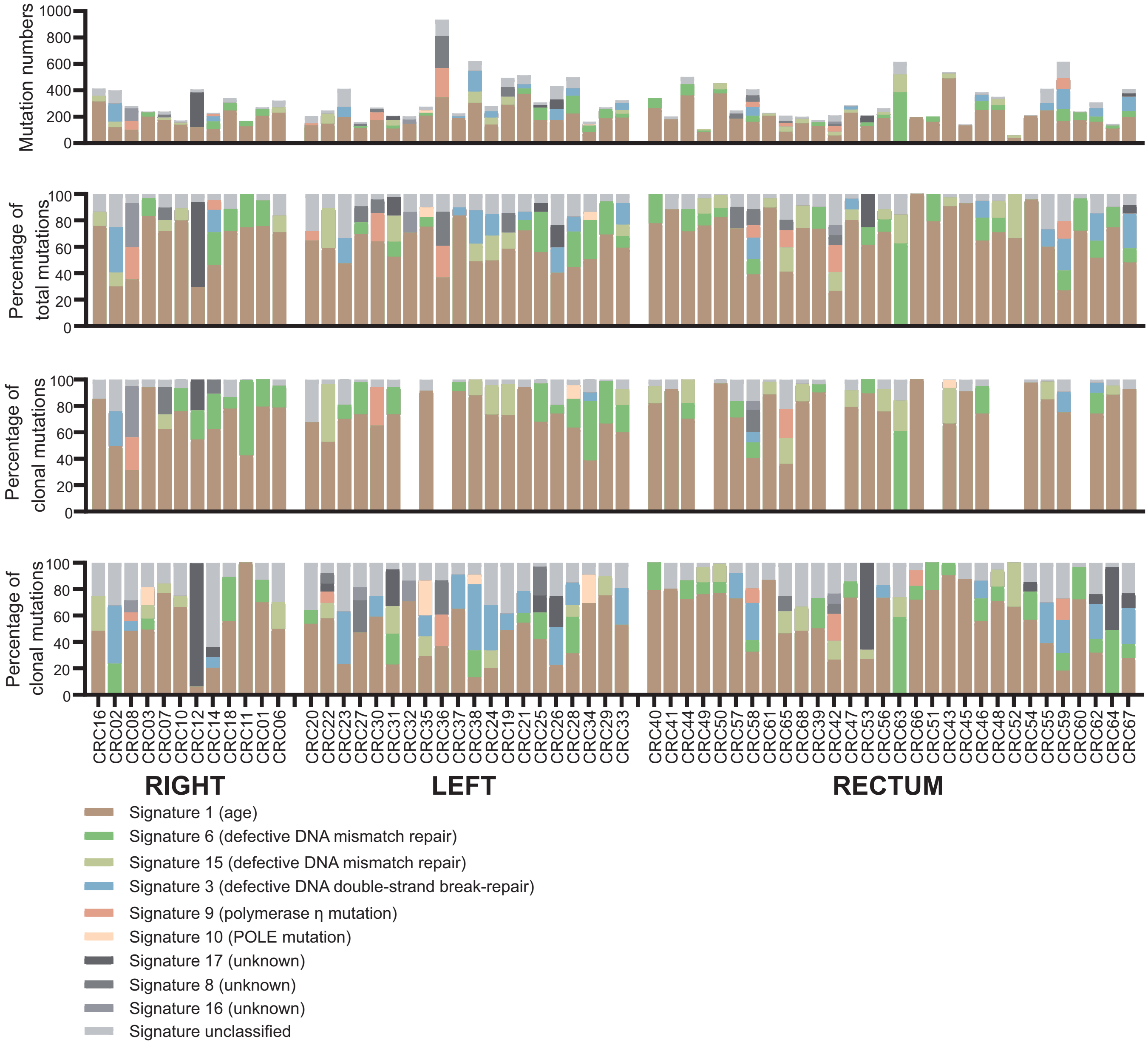

**Figure S16. Mutation signature in CRC tumors.**
Mutation signatures identified in CRC tumors, split according to tumor location and clonality of mutations.

### Figure S17

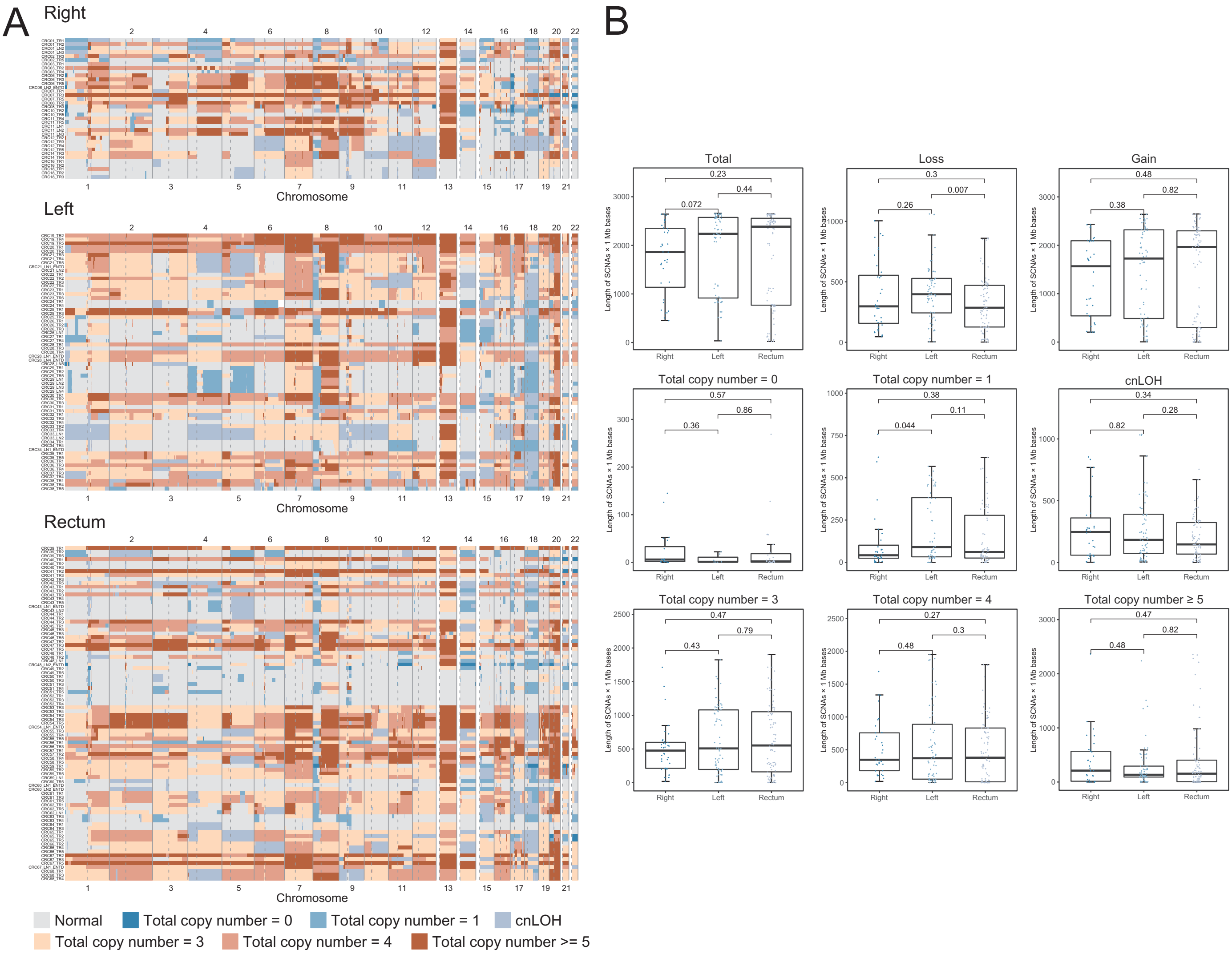

**Figure S17. Heatmap of SCNAs among right-sided colon, left-sided colon and rectal cancers.**
(A) Heatmap of genome-wide view SCNAs were shown for all the regions of CRC tumors by position.
(B) Box plots of different SCNAs types of CRC tumors by position. Loss included total copy number 0, 1 and copy neutral loss of heterozygosity (cnLOH). Gain included total copy number 3, 4 and ≥5.

### Figure S18

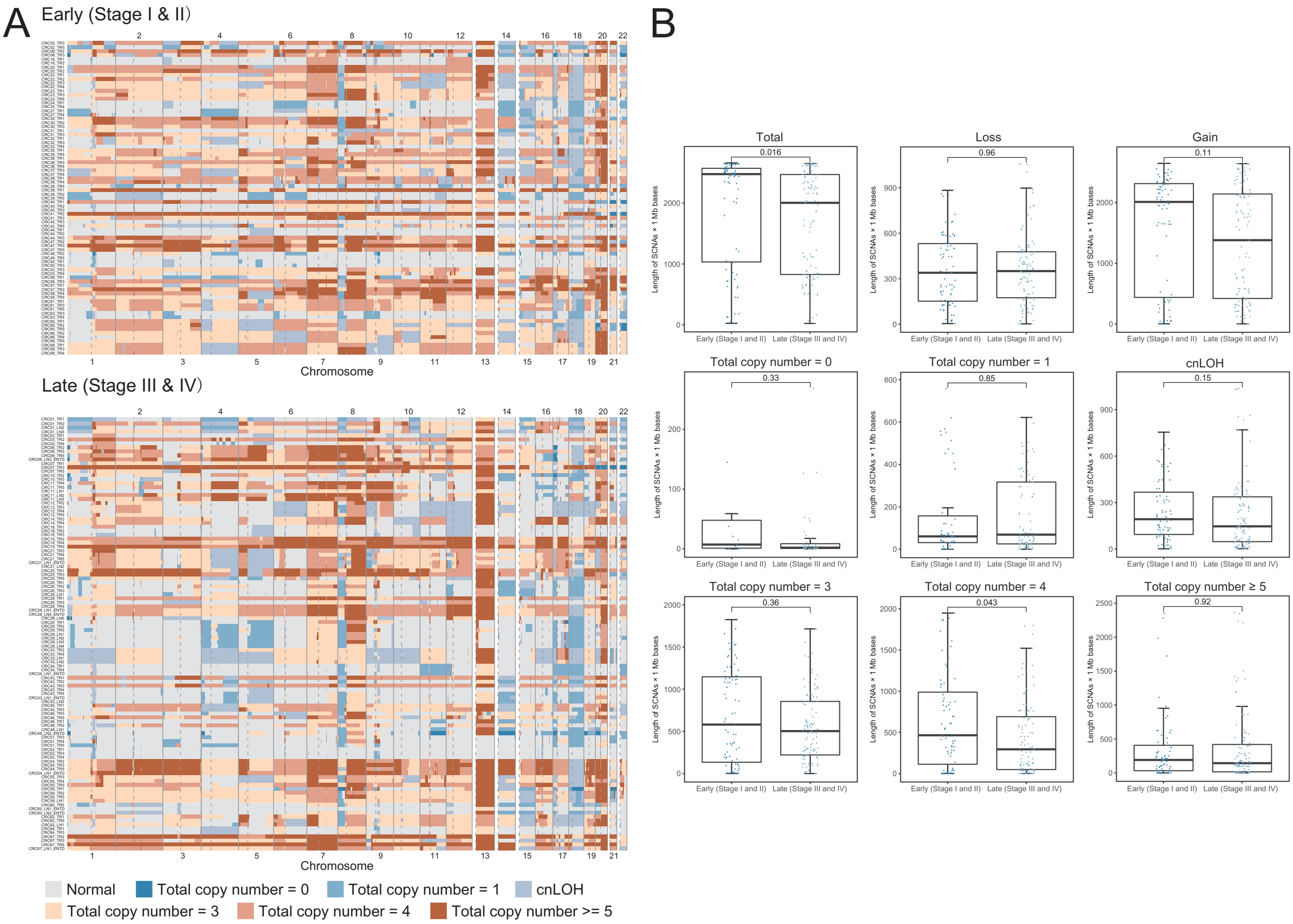

**Figure S18. Heatmap of SCNAs between early and late stage of CRC tumors.**
(A) Heatmap of genome-wide view SCNAs were shown for all the regions of early and late stage of CRC tumors.
(B) Box plots of different SCNAs types by early and late stage of CRC tumors. Loss included total copy number 0, 1 and cnLOH. Gain included total copy number 3, 4 and ≥5.

### Figure S19

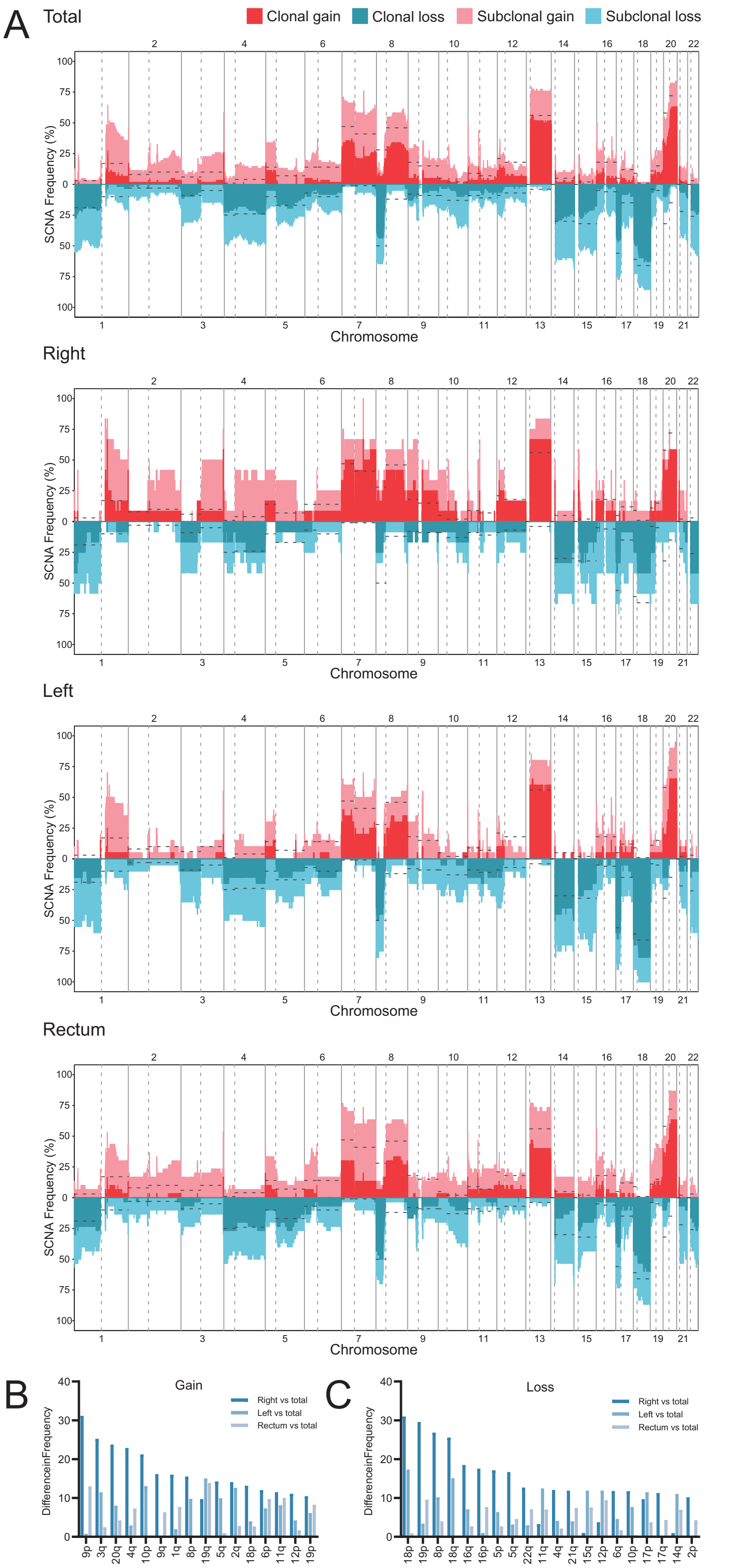

**Figure S19. Differences in SCNA frequencies among total, right-sided colon, left-sided colon and rectal cancers.**

(A) SCNA frequency of CRC tumors based on position. The dotted lines were frequency of SCNAs in TCGA CRC samples.

(B) and (C) The mean frequency of each chromosome arm by tumor position were calculated and only the chromosome arm with absolute difference of more than 10% were shown.

### Figure S20

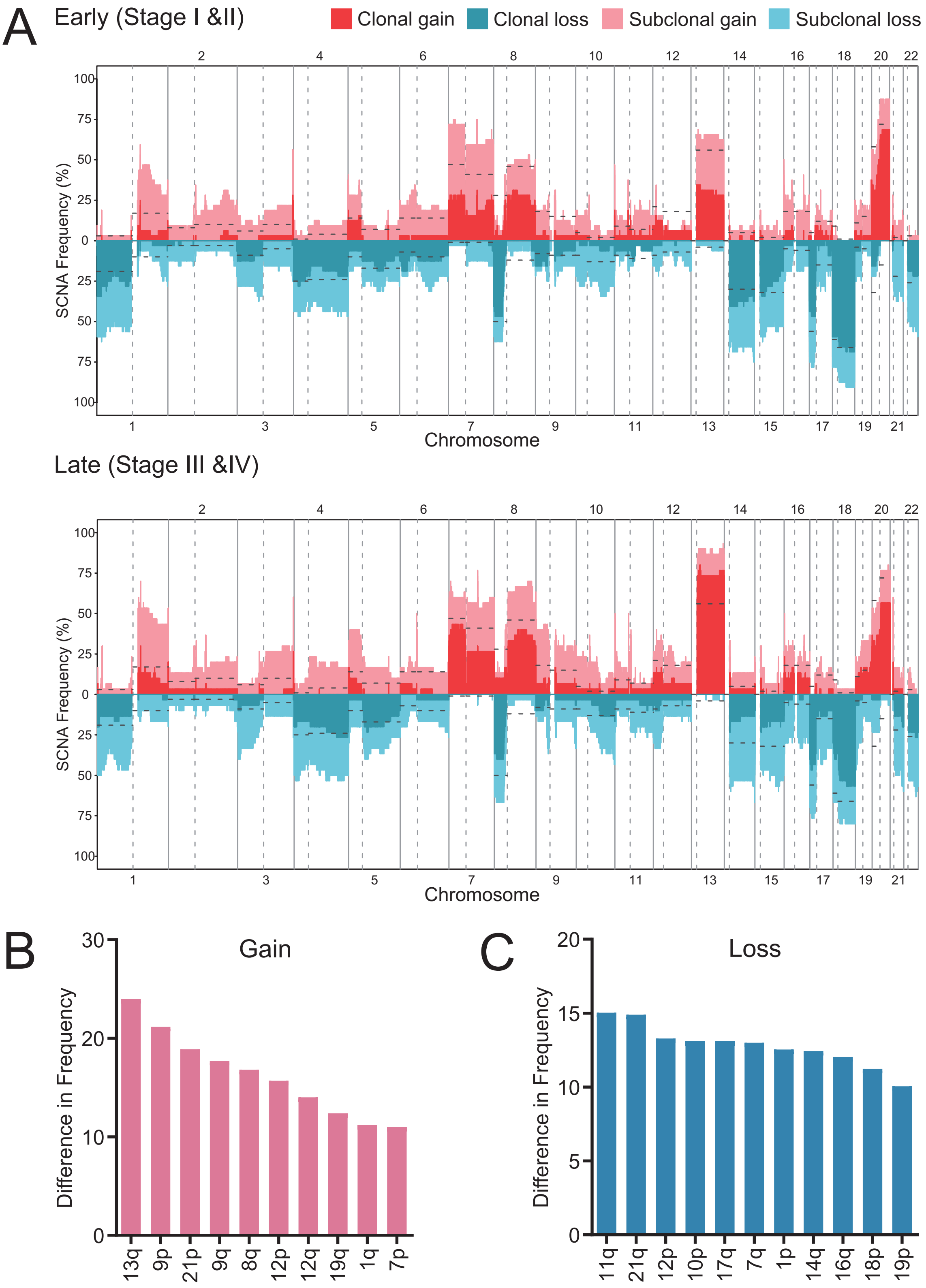

**Figure S20. Difference in SCNA frequencies between early and late stage of CRC tumors.**

(A) SCNA frequency of CRC tumors based on stage. The dotted lines were frequency of SCNAs in TCGA CRC samples.

(B) and (C) The mean frequency of each chromosome arm by early and late stage were calculated and only the chromosome arm with absolute difference of more than 10% were shown.

**Figure S21**

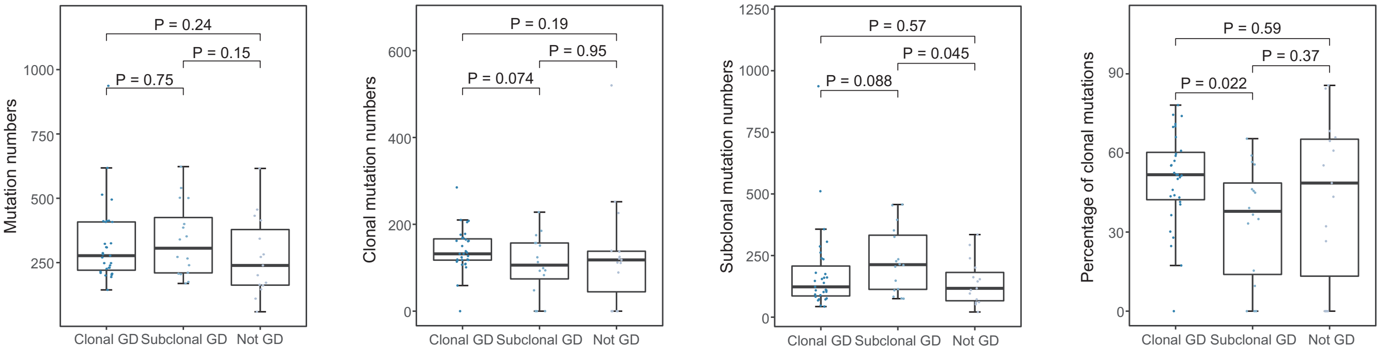

**Figure S21. Intratumor heterogeneity of mutations among different whole genome doubling status.**
Box plots of total, clonal, subclonal and percentage of clonal mutations by different whole genome doubling status: clonal genome doubling, subclonal genome doubling and no genome doubling.

#### Figure S22

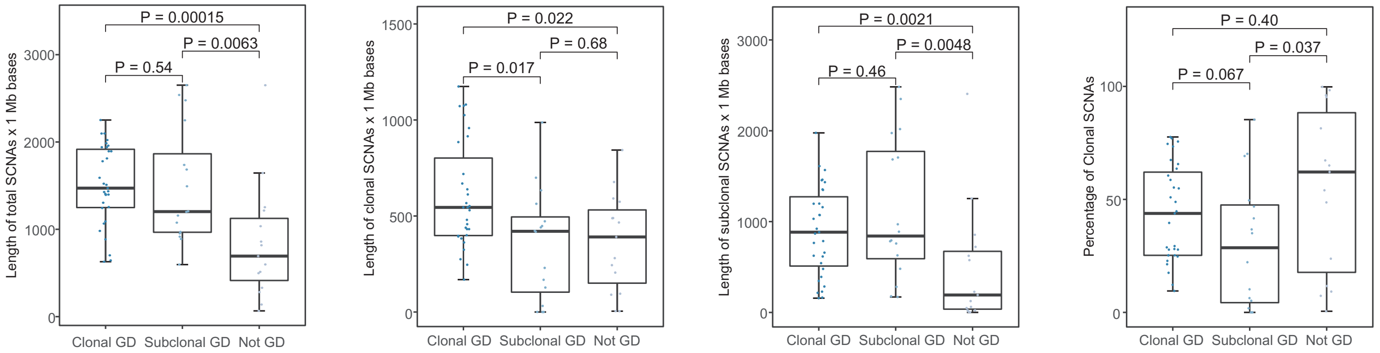

**Figure S22. Intratumor heterogeneity of SCNAs among different whole genome doubling status.**
Box plots of total, clonal, subclonal and percentage of clonal SCNAs by different whole genome doubling status: clonal genome doubling, subclonal genome doubling and no genome doubling.

### Figure S23

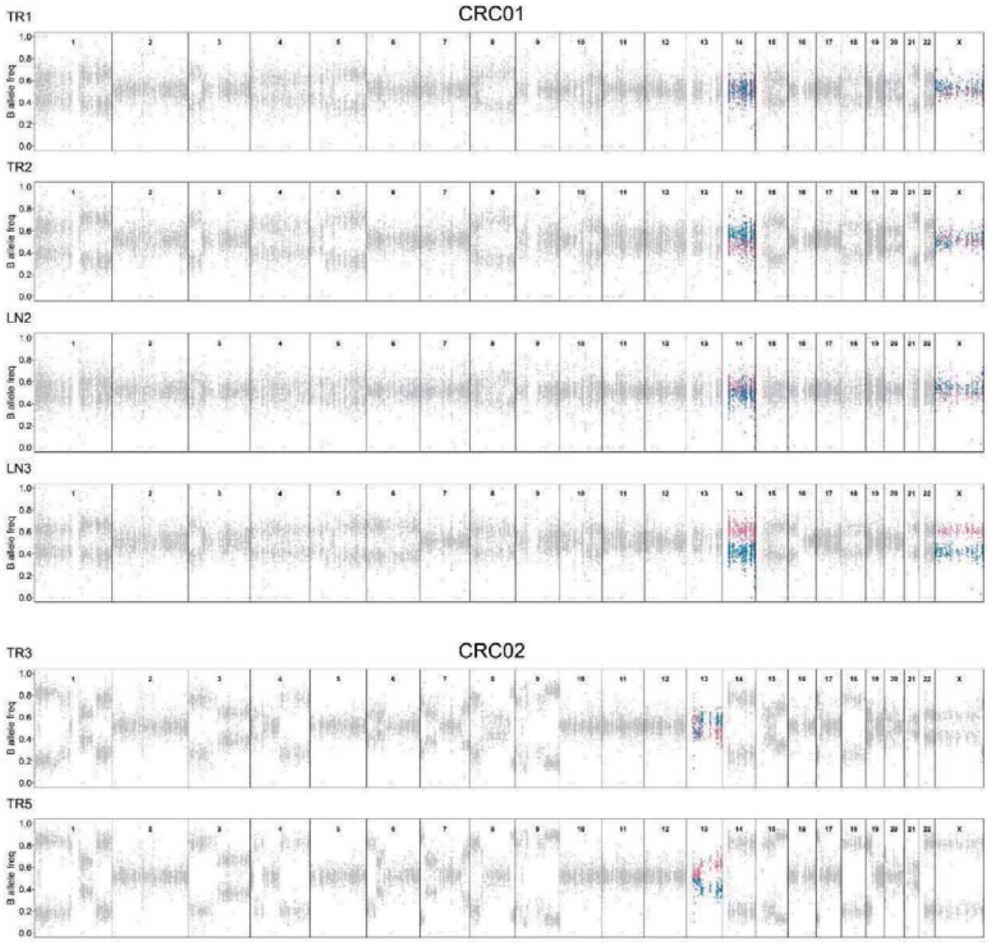

**Figure S23. Mirrored subclonal allelic imbalance (MSAI) events in CRC tumors.**
B-allele frequency (BAF) profile of heterozygous SNPs across the genome (chromosomes 1-22, X) from all regions of different tumors. Sections of BAF in regions that have MSAI were highlighted in blue or red.

CRC06

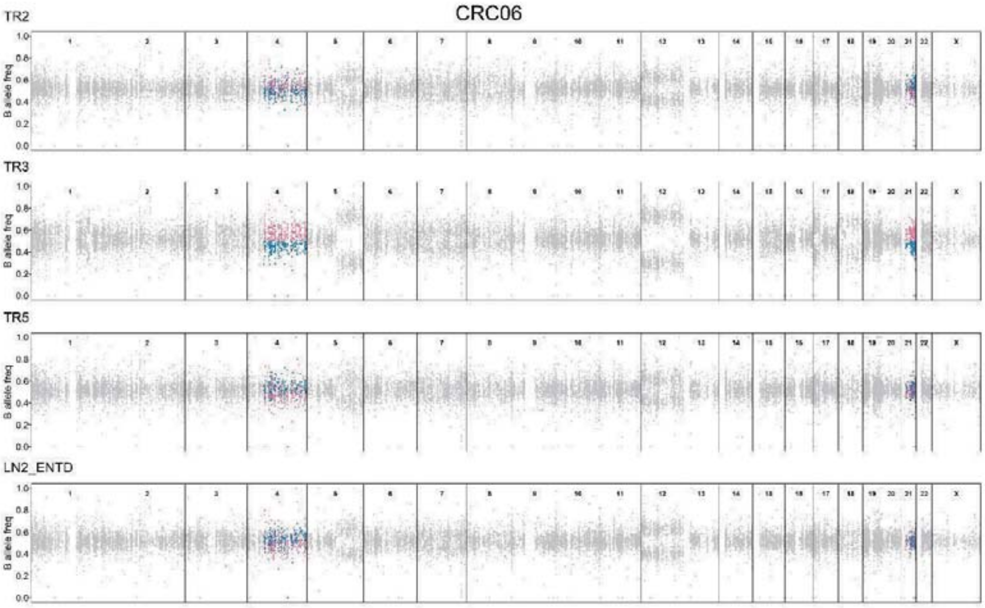

TR2
TR3
TR5
LN2_ENTD
B allele freq
1.0
0.8
0.6
0.4
0.2
0.0
1 2 3 4 5 6 7 8 9 10 11 12 13 14 15 16 17 18 19 20 21 22 X

CRC07

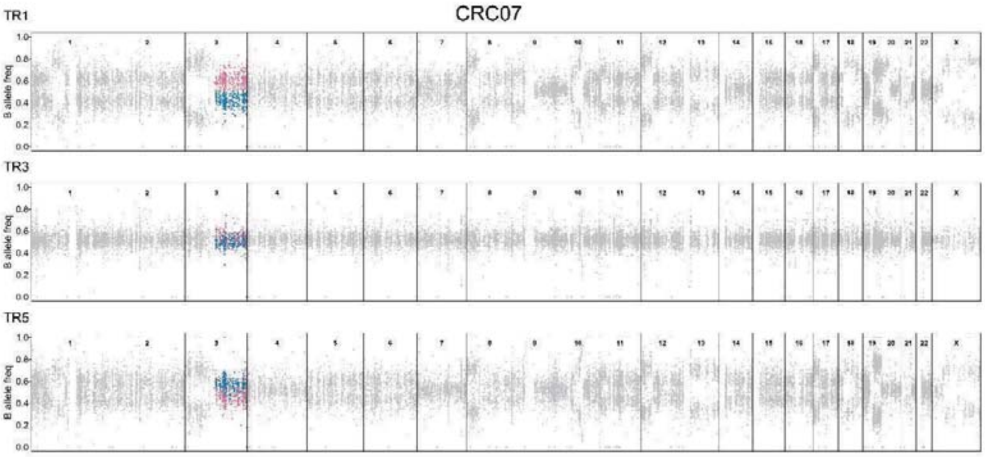

TR1
TR3
TR5
B allele freq
1.0
0.8
0.6
0.4
0.2
0.0
1 2 3 4 5 6 7 8 9 10 11 12 13 14 15 16 17 18 19 20 21 22 X

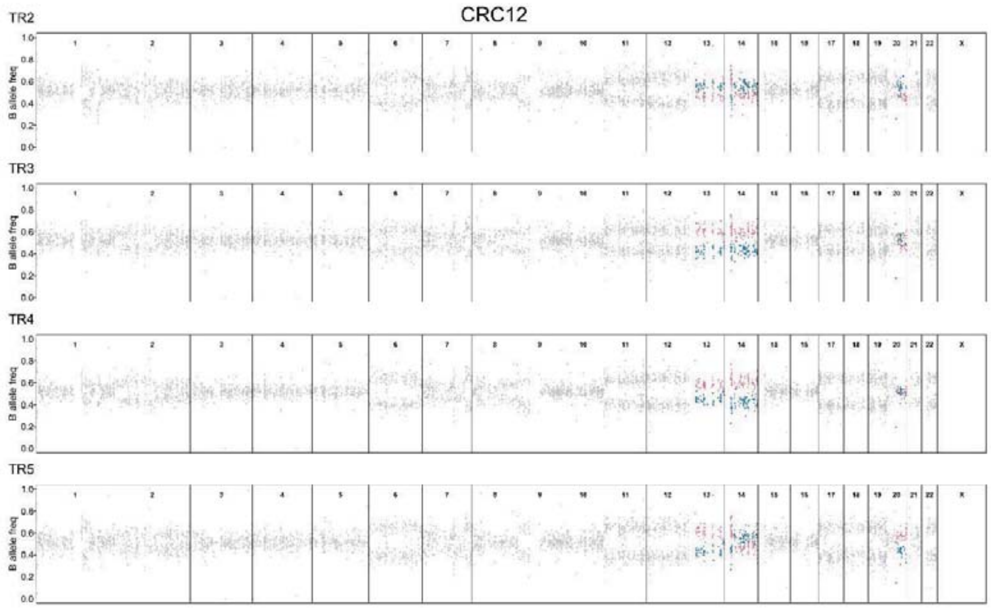
CRC12
TR2
TR3
TR4
TR5
B allele freq

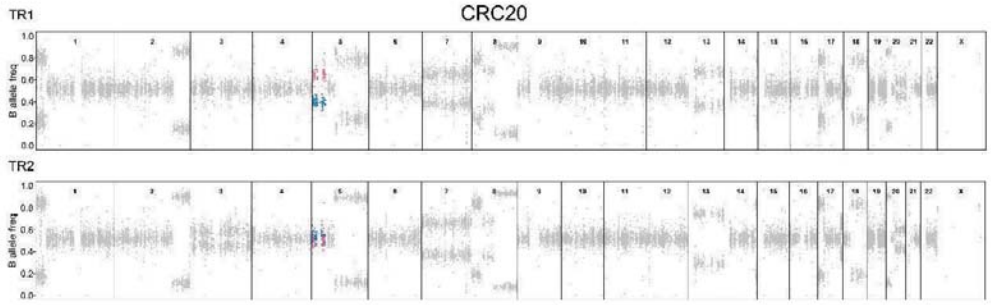
CRC20
TR1
TR2
B allele freq

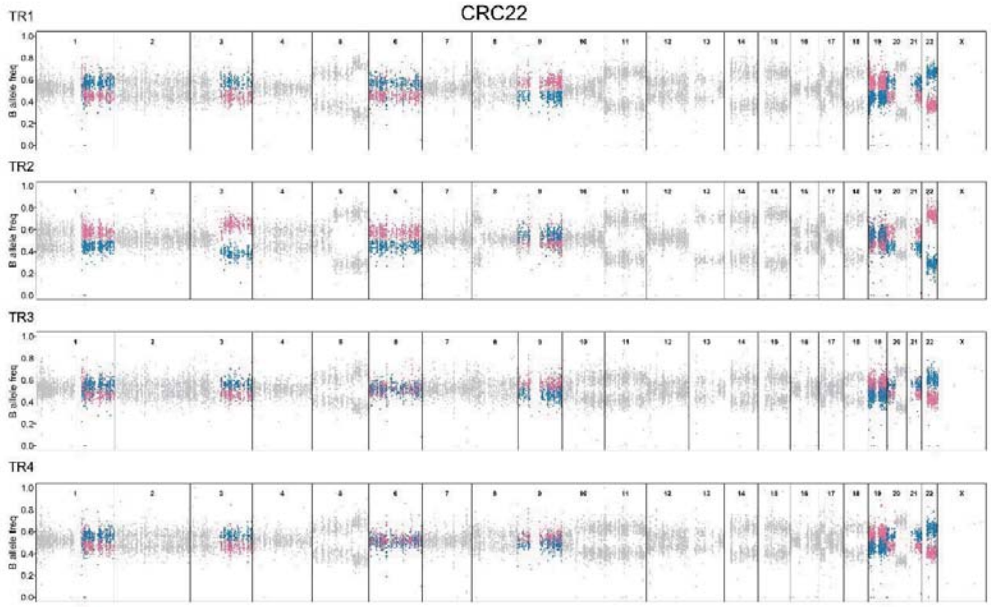
CRC22
TR1
TR2
TR3
TR4
B allele freq

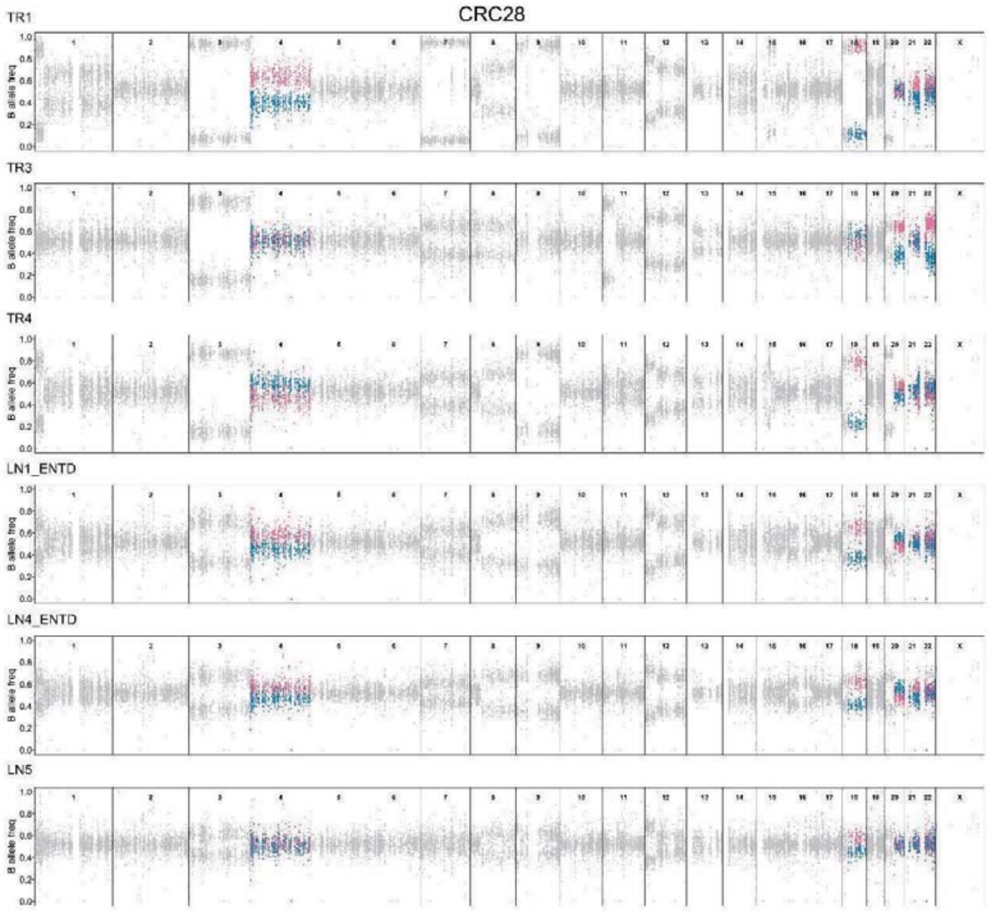
CRC28
TR1
TR3
TR4
LN1_ENTD
LN4_ENTD
LN5
B allele freq

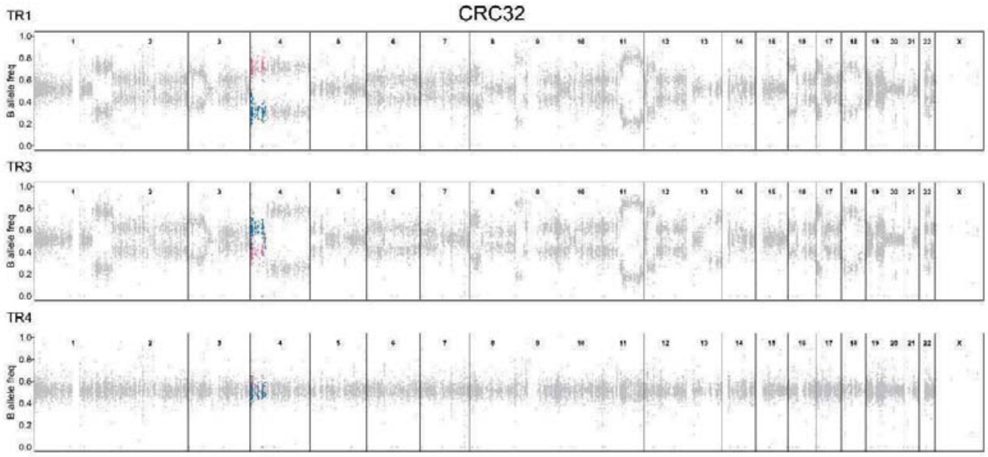
CRC32
TR1
TR3
TR4
B allele freq

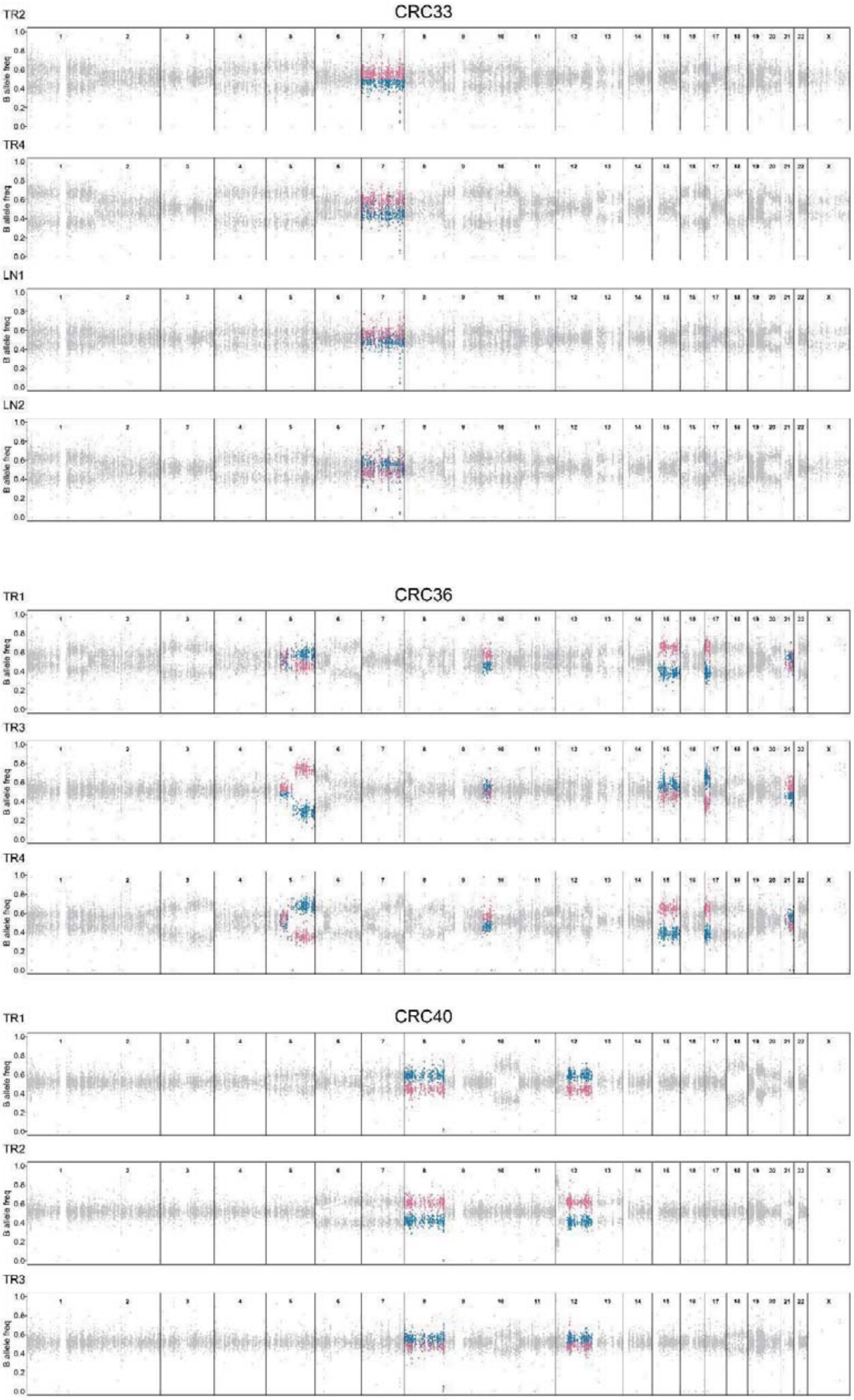
CRC33
TR2
TR4
LN1
LN2
CRC36
TR1
TR3
TR4
CRC40
TR1
TR2
TR3
B allele freq
1 2 3 4 5 6 7 8 9 10 11 12 13 14 15 16 17 18 19 20 21 22 X

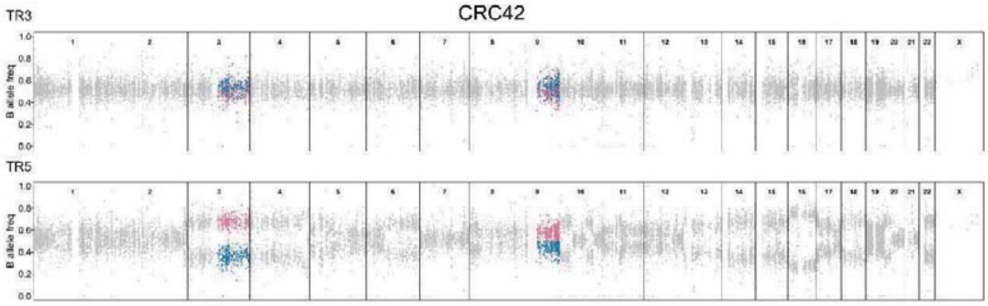
CRC42
TR3
TR5
B allele freq

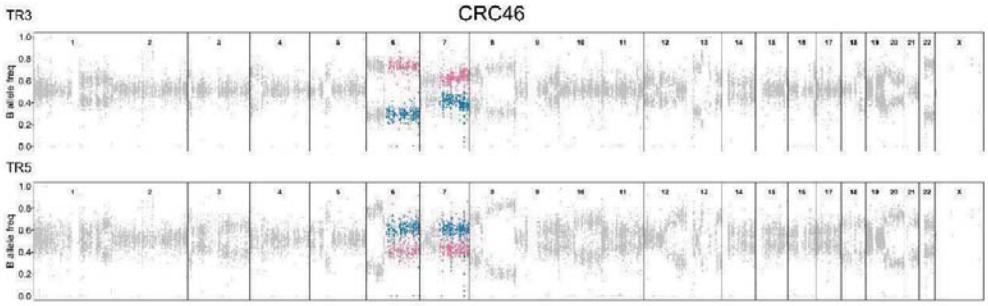
CRC46
TR3
TR5
B allele freq

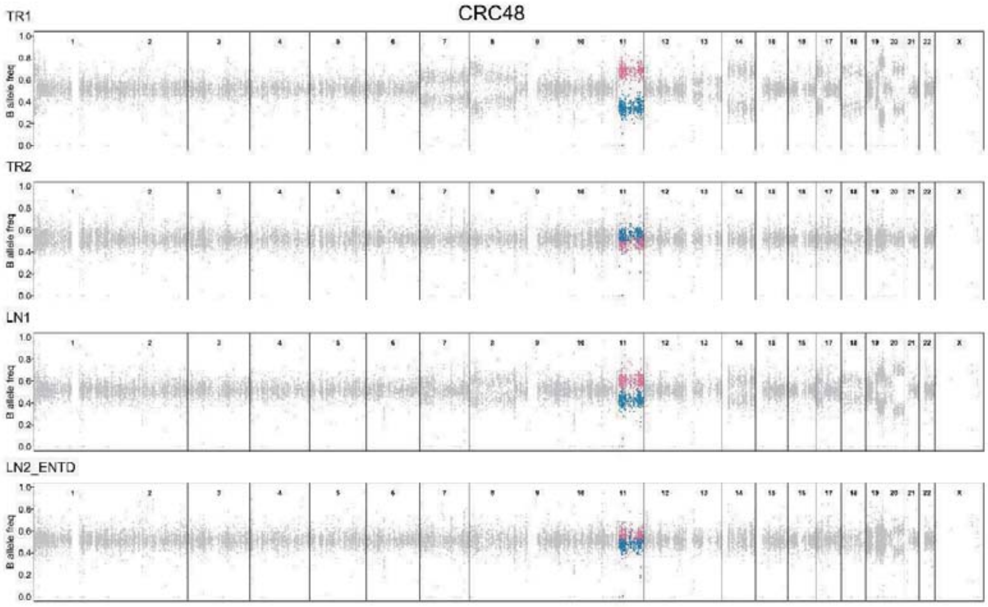
CRC48
TR1
TR2
LN1
LN2_ENTD
B allele freq

CRC50

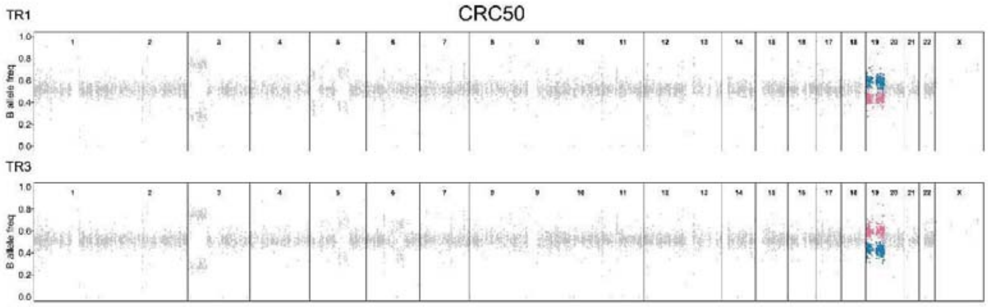

CRC53

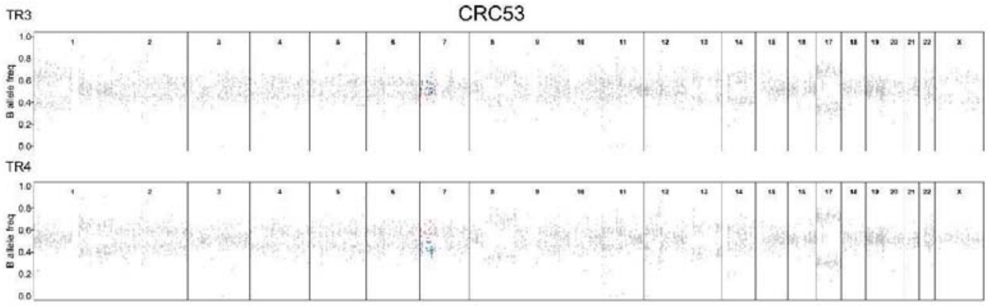

CRC55

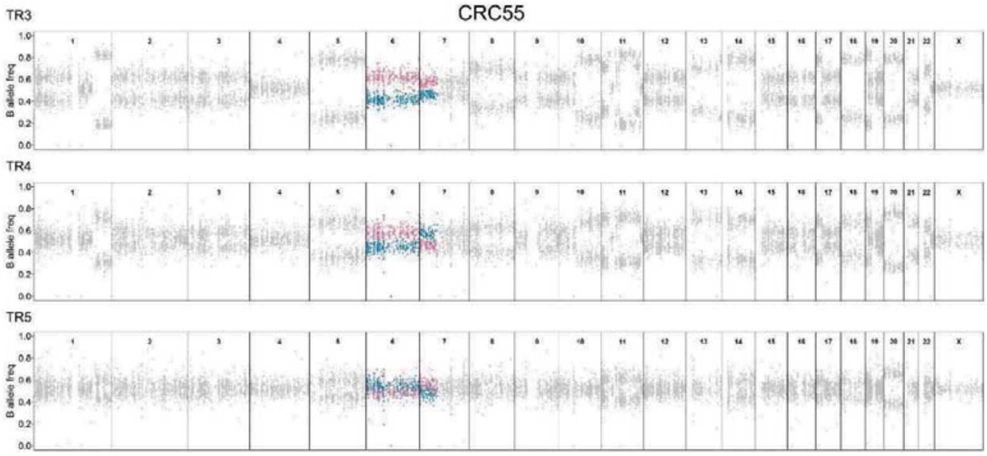

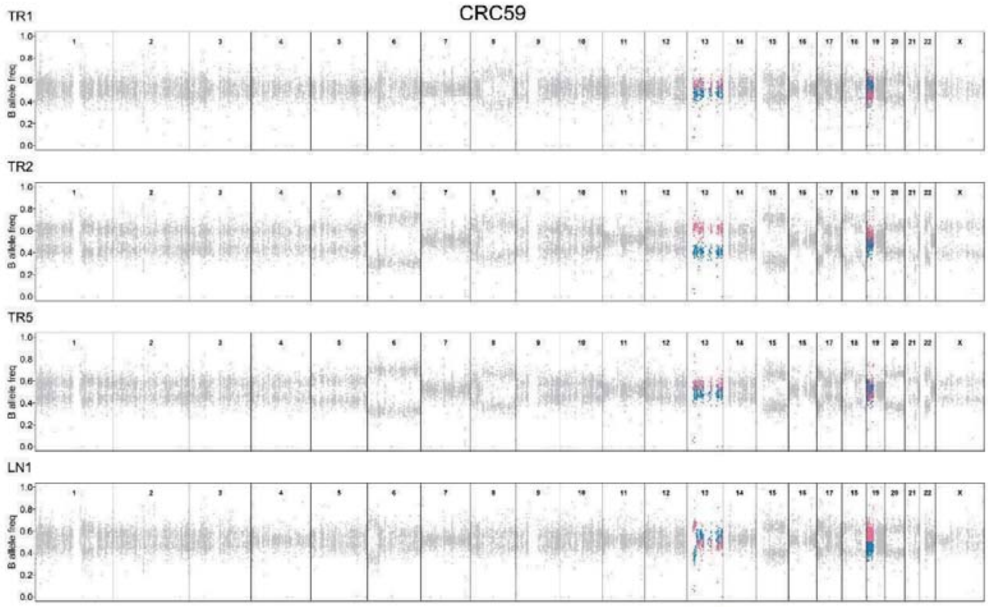
CRC59
TR1
TR2
TR5
LN1
B allele freq
0.0
0.2
0.4
0.6
0.8
1.0
1 2 3 4 5 6 7 8 9 10 11 12 13 14 15 16 17 18 19 20 21 22 X

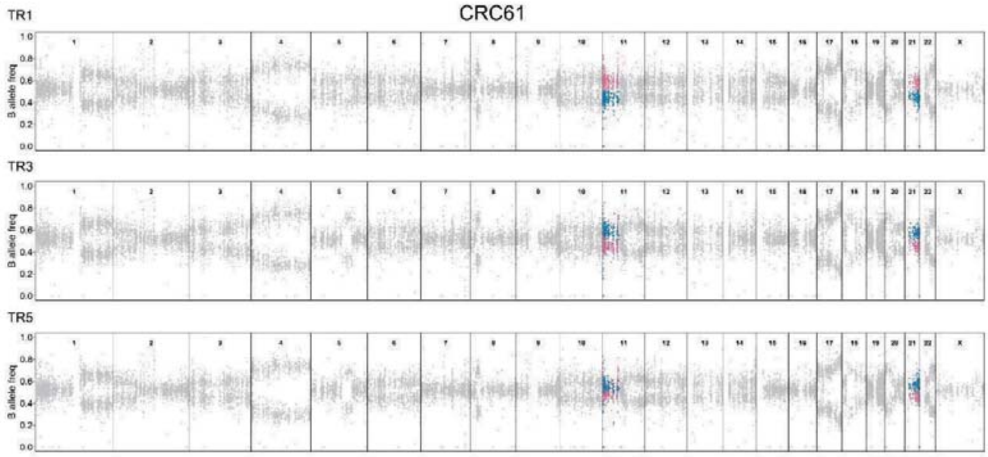
CRC61
TR1
TR3
TR5
B allele freq
0.0
0.2
0.4
0.6
0.8
1.0
1 2 3 4 5 6 7 8 9 10 11 12 13 14 15 16 17 18 19 20 21 22 X

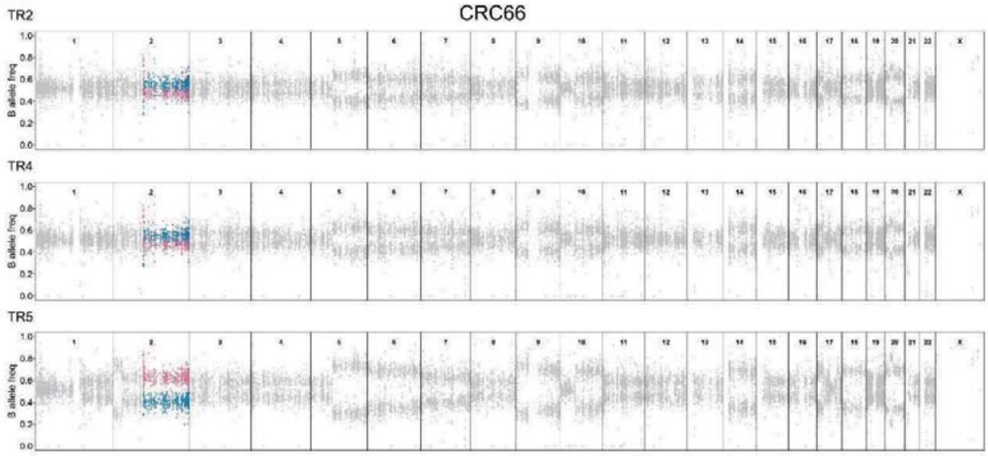
CRC66
TR2
TR4
TR5
B allele freq
0.0
0.2
0.4
0.6
0.8
1.0
1 2 3 4 5 6 7 8 9 10 11 12 13 14 15 16 17 18 19 20 21 22 X

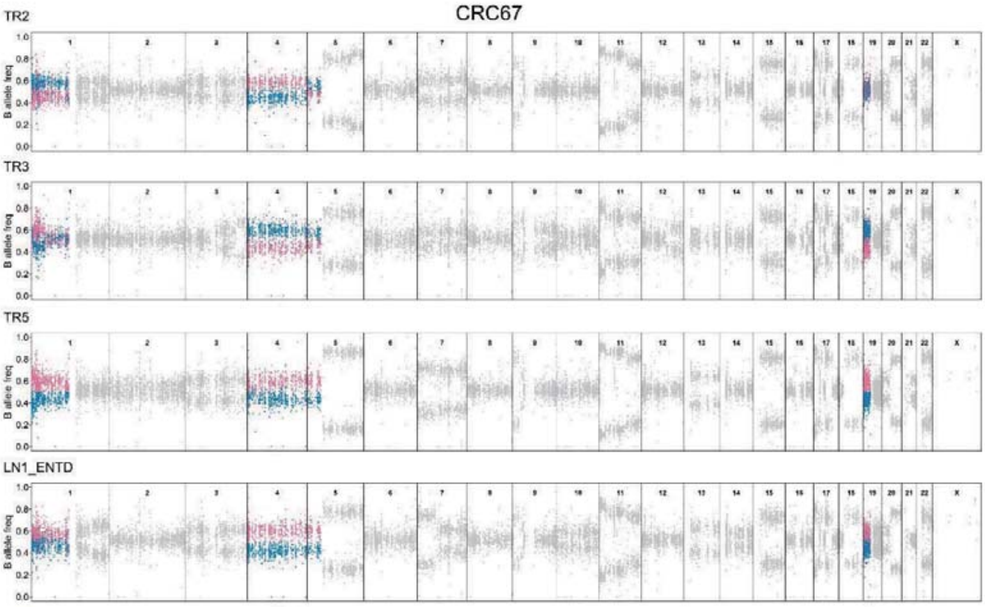
CRC67
TR2
TR3
TR5
LN1_ENTD
B allele freq

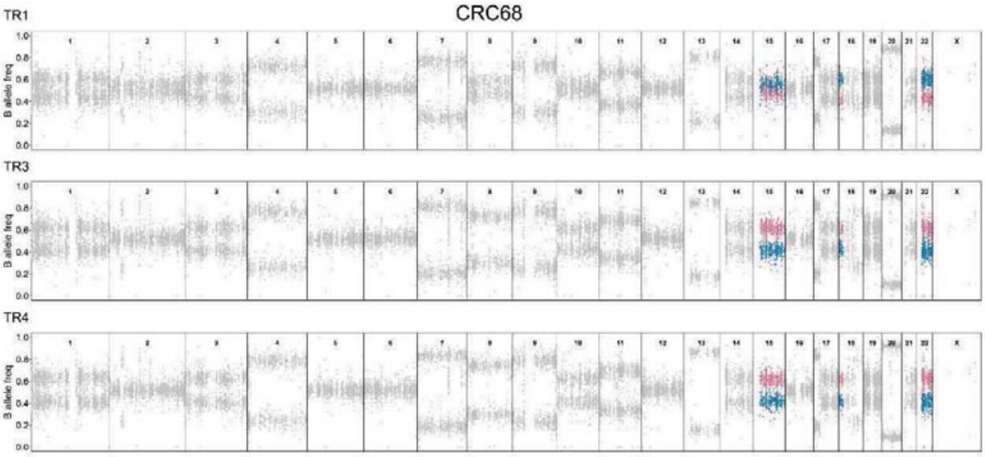
CRC68
TR1
TR3
TR4
B allele freq

#### Figure S24

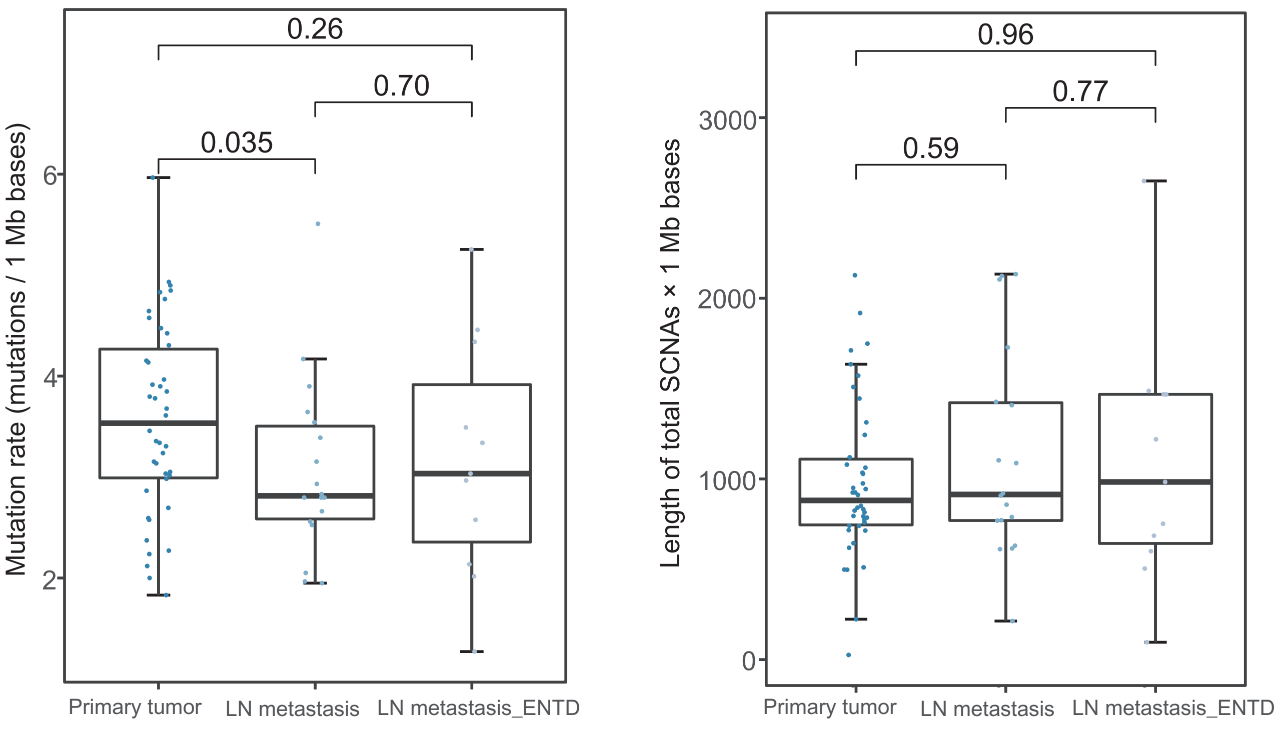

**Figure S24. Intratumor heterogeneity of mutations and SCNAs between different tumor region types in patients with lymph node metastasis or ENTD.**

Box plots of total mutations and SCNAs were statistically analyzed by different tumor regions (primary tumors, lymph node metastasis and ENTD) of patients with lymph node metastasis or ENTD.

#### Figure S25

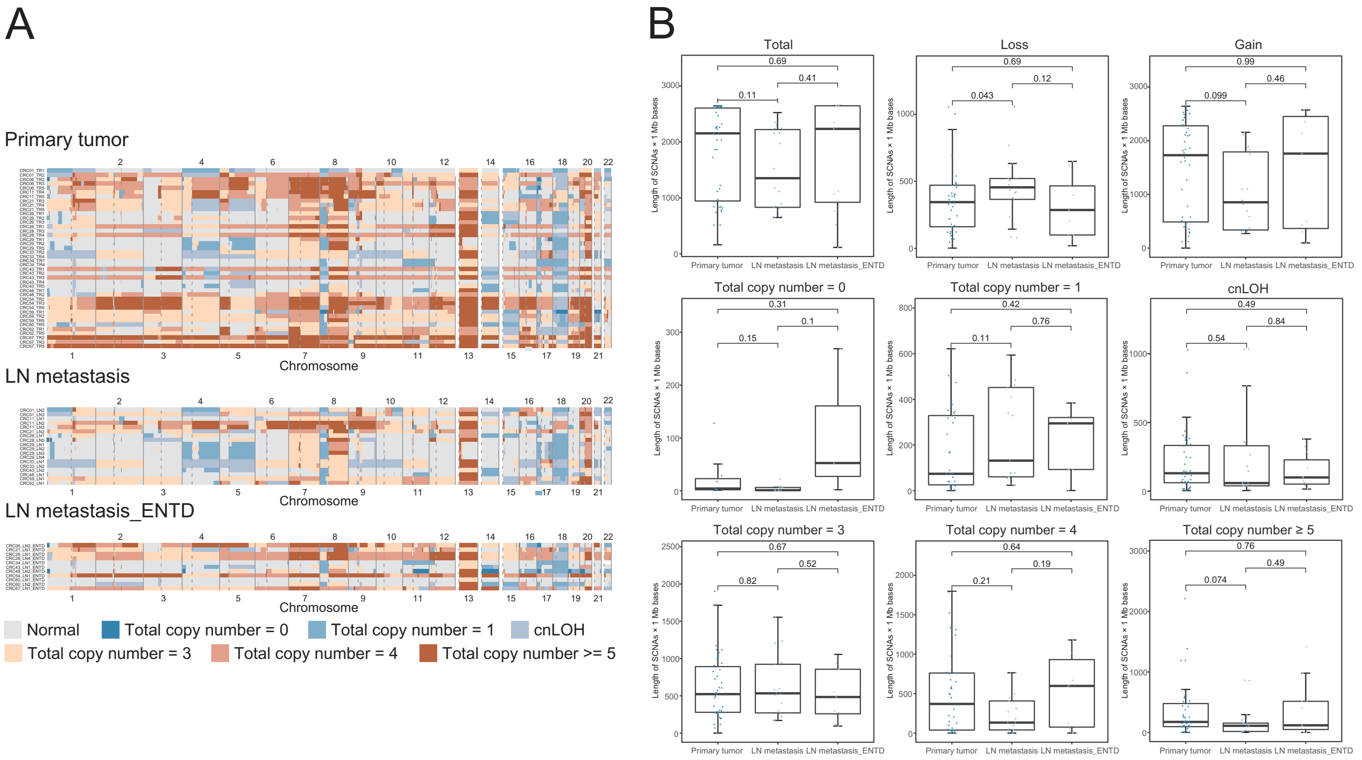

**Figure S25. Heatmap of SCNAs between different tumor region types of patients with lymph node metastasis or ENTD.**

(A) Heatmap of genome-wide view SCNAs were shown for different tumor region types (primary tumor, lymph node metastasis and ENTD) of patients with lymph node metastasis or ENTD.

(B) Box plots of different SCNAs types by different tumor region types of patients with lymph node metastasis or ENTD. Loss included total copy number 0, 1 and cnLOH. Gain included total copy number 3, 4 and ≥5.

### Figure S26

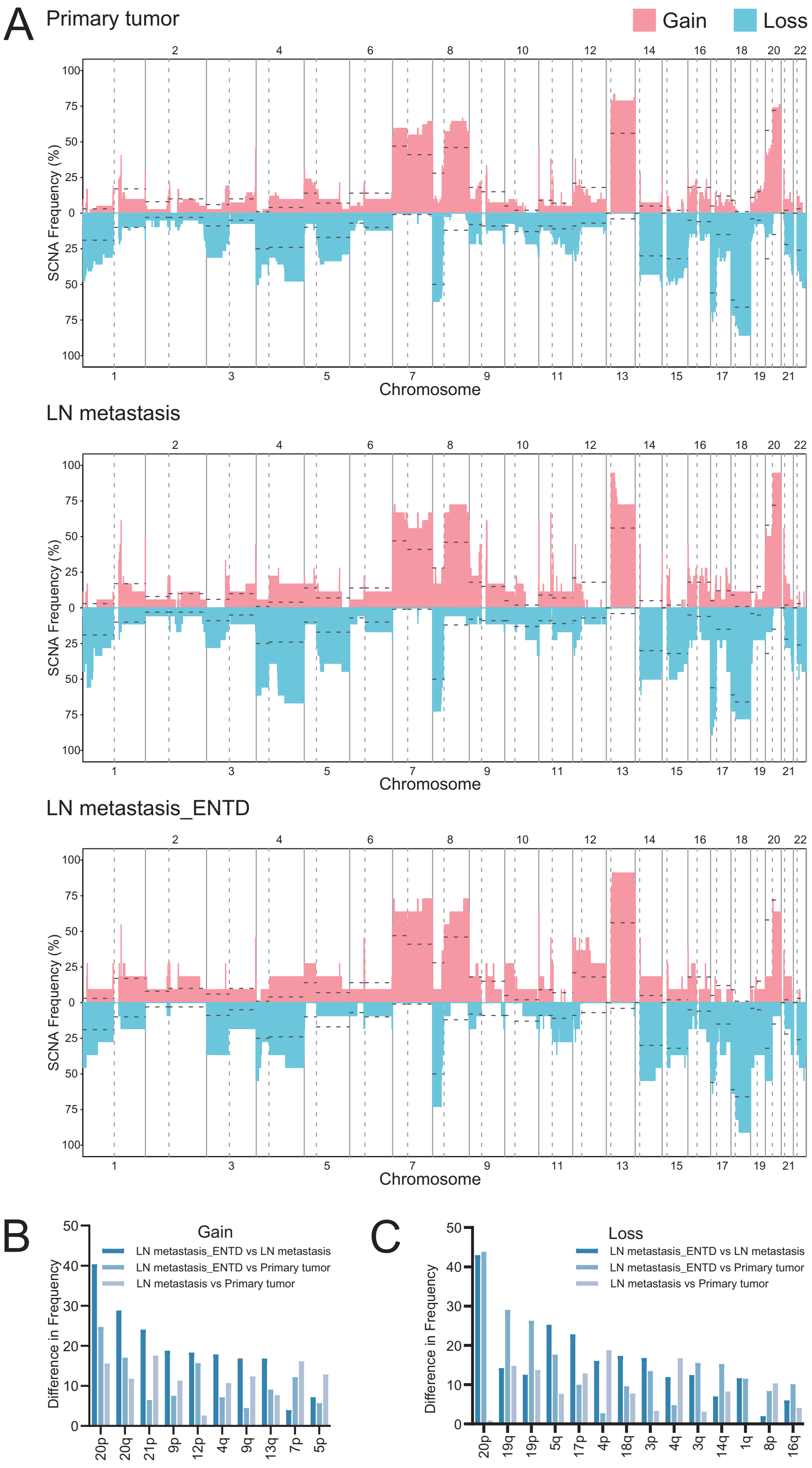

**Figure S26. Difference in SCNA frequency between different tumor region types of patients with lymph node metastasis or ENTD.**

(A) SCNA frequency of CRC tumors by different tumor region types (primary tumor, lymph node metastasis and ENTD) of patients with lymph node metastasis or ENTD. The dotted lines were frequency of SCNAs in TCGA CRC samples.

(B) and (C) The mean frequency of each chromosome arm by tumor region types of patients with lymph node metastasis or ENTD were calculated and only the chromosome arm with absolute difference of more than 10% were shown.

### Figure S27

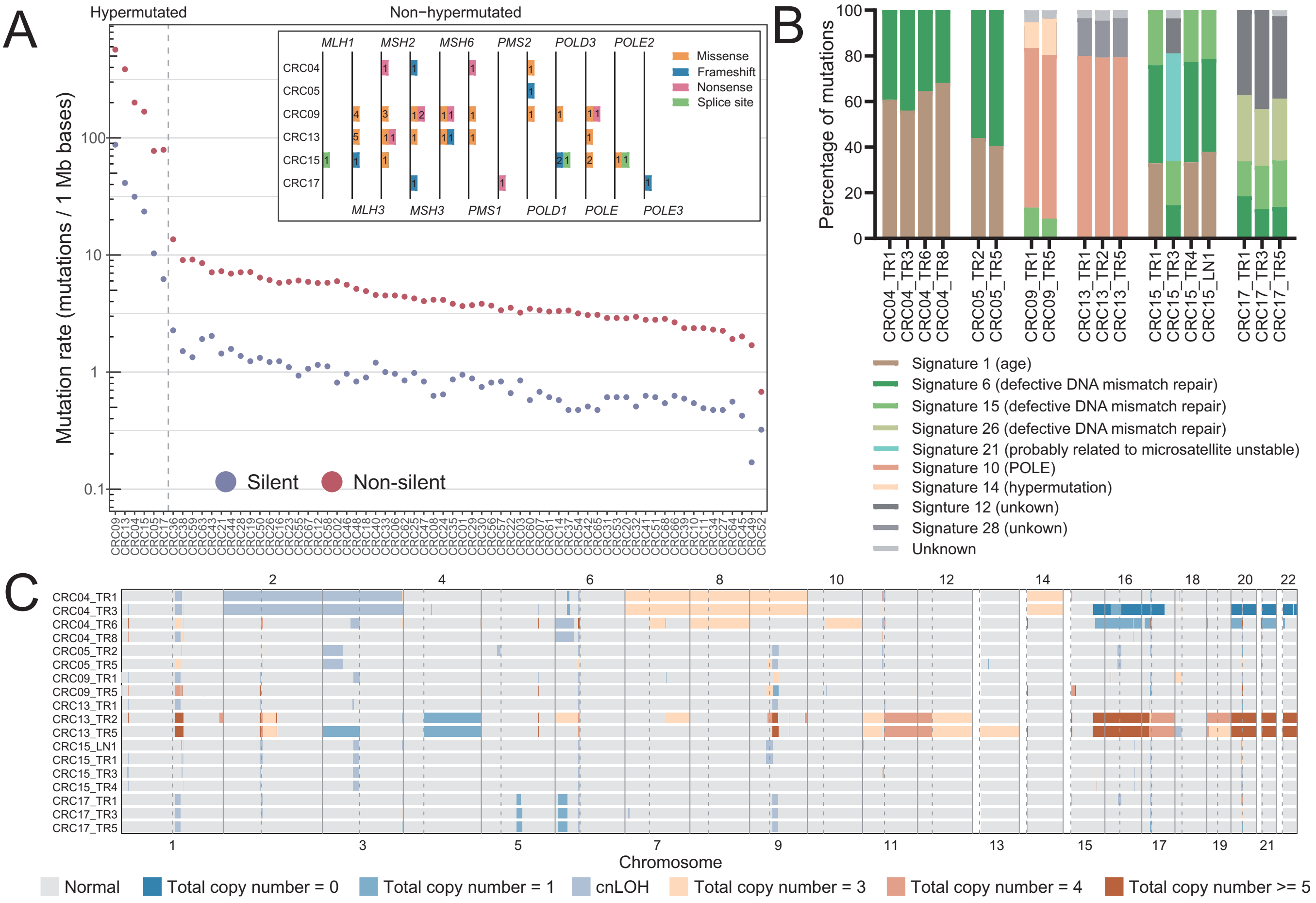

**Figure S27. Intratumor heterogeneity landscape of hypermutated CRC tumors.**

(A) Mutation rate of 68 CRC tumors. Inset, mutations in mismatch-repair genes, *POLE* and *POLD* gene family among the hypermutated tumors. The definition of hypermutated tumors was that all the sample in that tumor had more than 10 mutations/1 Mb bases.

(B) Mutation signature of all the samples from 6 hypermutated CRC tumors.

(C) Heatmap of genome-wide view SCNAs in 6 hypermutated CRC tumors.

### Figure S28

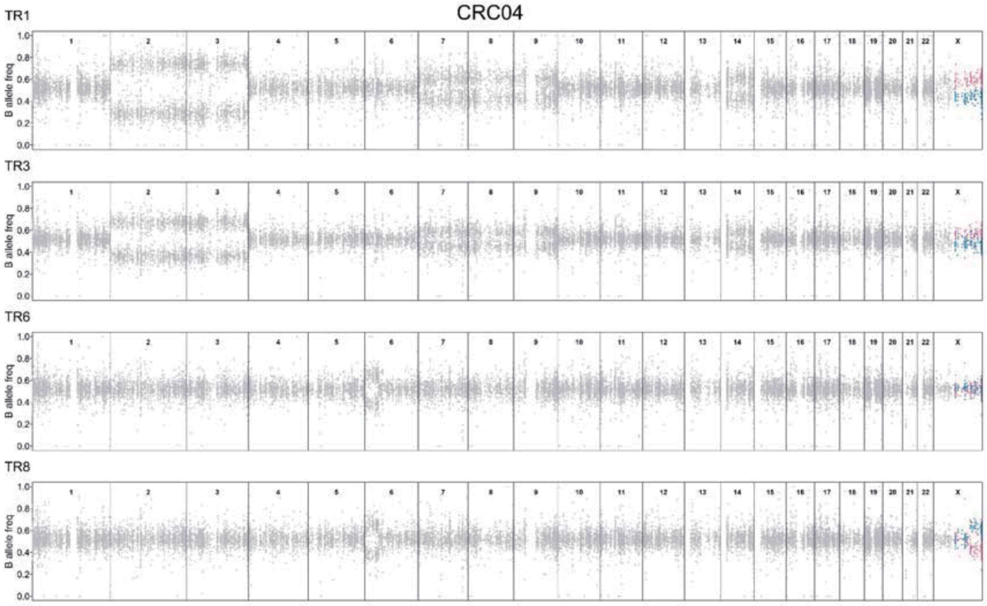

**Figure S28. MSAI events in hypermutated CRC tumor.**

B-allele frequency (BAF) profile of heterozygous SNPs across the genome (chromosomes 1-22, X) from all the regions of CRC04. Sections of BAF in regions that had MSAI were highlighted in blue or red.
